# Rational design of next-generation filovirus vaccines through glycoprotein stabilization, nanoparticle display, and glycan modification

**DOI:** 10.1101/2025.03.02.641072

**Authors:** Yi-Zong Lee, Yi-Nan Zhang, Maddy L. Newby, Garrett Ward, Keegan Braz Gomes, Sarah Auclair, Connor DesRoberts, Joel D. Allen, Andrew B. Ward, Robyn L. Stanfield, Linling He, Max Crispin, Ian A. Wilson, Jiang Zhu

## Abstract

Filoviruses pose a significant threat to human health due to frequent outbreaks and high mortality. Although two vector-based vaccines are available for Ebola virus, a broadly protective filovirus vaccine remains elusive. Here, we evaluate a general strategy for stabilizing glycoprotein (GP) structures from Ebola, Sudan, and Bundibugyo orthoebolaviruses and Ravn orthomarburgvirus. A 3.2 Å crystal structure provides atomic-level details of the redesigned Ebola virus GP, while cryo-electron microscopy reveals how a pan-orthoebolavirus neutralizing antibody targets a conserved site on the stabilized Sudan virus GP (3.13 Å resolution), along with a low-resolution model of antibody-bound Ravn virus GP. A self-assembling protein nanoparticle (SApNP), I3-01v9, is redesigned at the N terminus to enable optimal surface display of filovirus GP trimers. Following detailed in vitro characterization, we examine the lymph node dynamics of Sudan virus GP and GP-presenting SApNPs in mice. Compared with the soluble trimer, SApNPs exhibit ∼112-fold longer retention in lymph node follicles, up to 28-fold greater presentation on follicular dendritic cell dendrites, and up to 3-fold stronger germinal center reactions. Functional antibody responses induced by filovirus GP trimers and SApNPs bearing wild-type and modified glycans are assessed in mice. This study provides a foundation for next-generation filovirus vaccine development.

**ONE-SENTENCE SUMMARY:** Filovirus glycoproteins and nanoparticles were rationally designed and characterized in vitro and in vivo to aid filovirus vaccine development.

## INTRODUCTION

The first documented case of human filovirus infection occurred in 1967 in Marburg and Frankfurt, Germany, when laboratory workers were infected with Marburg virus (MARV) by non-human primates (NHPs) imported from Africa (*1*). The largest filovirus outbreak on record was caused by Ebola virus (EBOV), first identified in 1976 in Zaire, Central Africa (*2, 3*). The 2013–2016 EBOV epidemic resulted in approximately 28,600 cases and 11,325 deaths (*4*). Both MARV and EBOV cause viral hemorrhagic fever (VHF) in humans, with case fatality rates of up to 90% (*5, 6*). Filoviruses are classified into nine genera, of which *Orthoebolavirus* and *Orthomarburgvirus* are the most clinically relevant (*7–9*). *Orthoebolavirus* includes pathogenic species such as EBOV, Bundibugyo virus (BDBV), and Sudan virus (SUDV), as well as other species that are either non-pathogenic or rarely infect humans. *Orthomarburgvirus* comprises a single species containing two members: MARV and Ravn virus (RAVV). The single-stranded, negative-sense RNA genome of filoviruses encodes structural proteins, including nucleoprotein, virion proteins, glycoprotein (GP), and RNA polymerase. As a class I viral fusion protein (*10–13*), GP functions as a trimer of heterodimers, mediating host cell attachment, membrane fusion, and viral entry (*14*). GP is recognized by the host immune system during natural infection (*15–17*) and represents the primary target for the development of prophylactic vaccines and antibody therapeutics (*18–21*).

Substantial progress has been made in filovirus research (*22*). First, our understanding of filovirus pathogenesis and infection mechanisms has advanced considerably (*23–25*). Niemann-Pick C1 (NPC1) was identified as the endosomal receptor for filovirus entry, indicating that NPC1 inhibitors may have antiviral potential (*26–28*). Second, neutralizing antibodies (NAbs) have been established as effective therapeutics for treating filovirus infections (*29–32*). Compared to the monoclonal antibody (mAb) cocktail ZMapp (*33*), which initially showed promise (*34*), REGN-EB3 (a three-mAb cocktail (*35*)) and mAb114 (*36*) demonstrated superior efficacy against EBOV in a randomized controlled trial (*37*). The 2013–2016 EBOV outbreak led to sustained efforts to identify anti-filovirus antibodies (*38*). Panels of mAbs were isolated from human survivors and immunized animals, revealing multiple sites of vulnerability on GP (*16, 17, 36, 39–54*). At the peak of this effort, the Viral Hemorrhagic Fever Immunotherapeutic Consortium analyzed 171 mAbs targeting diverse epitopes on GP, including the base, glycan cap, fusion loop, GP1/core, GP1/head, GP1/2 interface, heptad repeat 2 (HR2), and mucin-like domain (MLD) (*55*). Third, X-ray crystallography and cryo-electron microscopy (cryo-EM) have determined atomic structures of filovirus GP bound to NPC1 (*28*), small-molecule ligands (*56–58*), and mAbs (*16, 39, 40, 42, 43, 47, 49–52, 59, 60*). Notably, negative-stain EM (nsEM) remains an effective tool for validating GP conformation and mapping mAb epitopes (*17, 41, 46–48, 53–55*). These structural insights have advanced our understanding of filovirus neutralization and supported the development of antibody therapeutics and vaccines (*18, 61, 62*). Fourth, two viral vector-based EBOV vaccines have been approved for human use. ERVEBO (*63*) is a single-dose vaccine based on recombinant vesicular stomatitis virus (rVSV) expressing EBOV GP, whereas Zabdeno/Mvabea (*64*) delivers two doses – an Adenovirus 26 (Ad26) vector encoding EBOV GP followed by a modified Vaccinia Ankara (MVA) vector expressing multiple filovirus GPs. Additional EBOV vaccines using DNA (*65*) and viral vectors (*66–71*) have been tested in humans. Recent efforts have expanded to mRNA, protein subunit, and nanoparticle (NP) platforms (*72–75*).

Despite this progress, critical innovations are still needed to advance next-generation filovirus vaccines (*76*). Although correlates of protection remain undefined (*77*), in-depth B cell analysis revealed diverse yet convergent polyclonal NAb responses in a Phase 1 trial of the rVSV-ZEBOV vaccine (*78*). A follow-up study elucidated how NAbs from multiple individuals used the same germline genes to target the receptor-binding site (RBS) on GP (*79*), reinforcing the central role of GP as the primary vaccine antigen. Over the past decade, antibody-guided, structure-based antigen design has transformed vaccine development for important targets such as HIV-1 (*80–83*) and respiratory syncytial virus (RSV) (*84*). Rational design strategies have been applied to stabilize the HIV-1 envelope (Env) (*85–87*), the RSV fusion (F) protein (*88–91*), and, more recently, the SARS-CoV-2 spike (*92, 93*) in their prefusion conformations. Many of these constructs have progressed from preclinical testing to human trials (*94, 95*), with multiple COVID-19 vaccines (*96, 97*) and three RSV vaccines now approved for human use (*98*). By contrast, only two studies have reported rationally stabilized filovirus GP designs, both with limited in vivo evaluation (*99, 100*). Most recently, Xu et al. proposed an immunofocusing approach by engineering additional N-linked glycans onto GP (*74*). Notably, nearly all current filovirus vaccine candidates (*101*) rely on wild-type GP, which contains a highly variable MLD that shields conserved NAb epitopes and facilitates immune evasion (*102, 103*). A mucin-deleted GP (GPΔmuc) could, in principle, elicit NAbs more effectively than wild-type GP, but it has yet to be evaluated in humans. In addition to antigen format, the choice of delivery platform may also influence vaccine-induced immunity. Although viral vectors remain prominent, as reflected in the licensure of two EBOV vaccines, mRNA platforms represent a compelling alternative given their success during the COVID-19 pandemic (*104, 105*). In the realm of protein-based vaccines, NPs have emerged as a promising platform due to their virus-like size, multivalent surface display, and ability to elicit strong and durable NAb responses (*106–111*). Although two EBOV GP-presenting NP vaccines have recently been reported (*74, 100*), neither has been evaluated in animal challenge models. Finally, glycan modification may be a key strategy for optimizing filovirus vaccine antigens. Glycan trimming has improved NAb responses in vaccine candidates targeting influenza hemagglutinin (HA), SARS-CoV-2 spike, and HIV-1 Env (*112–115*). Whether this approach applies to filovirus GP, which is covered with up to 17 surface glycans per protomer, remains to be determined.

In our previous study, we stabilized EBOV GP with mutations in the HR1_C_ bend and HR2 stalk and displayed one of two lead designs, termed GPΔmuc-WL^2^P^2^, on self-assembling protein nanoparticles (SApNPs) as EBOV vaccine candidates (*100*). Here, we first extended the GPΔmuc-WL^2^P^2^ and WL^2^P^4^ construct designs from EBOV to SUDV and BDBV and determined a crystal structure of EBOV GPΔmuc-WL^2^P^4^ at 3.2 Å resolution. We also obtained a 3.13 Å cryo-EM reconstruction of SUDV GPΔmuc-WL^2^P^4^ bound to the CA45 Fab (*47*). The nearly identical backbones of SUDV and EBOV GPΔmuc-WL^2^P^4^ demonstrate a broadly applicable strategy for stabilizing orthoebolavirus GPs. We next displayed EBOV, SUDV, and BDBV GPΔmuc-WL^2^P^4^ trimers on SApNPs derived from E2p and I3-01v9, the latter of which was computationally redesigned at the N terminus. A similar design approach was applied to RAVV GP, resulting in a promising trimer construct termed GPΔmuc-P^2^CT. We then examined the lymph node delivery and immunological responses of SUDV GPΔmuc-WL^2^P^4^ trimer and SApNPs at the tissue, cellular, and subcellular levels in a mouse model. Compared to soluble trimers, 60-meric SApNPs exhibited prolonged retention and sustained presentation on follicular dendritic cell (FDC) dendrites, along with more robust and durable germinal center (GC) reactions in lymph node follicles. Finally, we immunized mice with a panel of filovirus GP vaccines to assess antibody responses. While the EBOV and SUDV GPΔmuc-WL^2^P^4^ trimer and SApNP vaccines elicited serum neutralization limited to orthoebolaviruses, the RAVV GPΔmuc-P^2^CT trimer induced detectable cross-genus NAb responses. Both multivalent display and glycan modification influenced the immunogenicity of GP-based filovirus vaccines. Although no universal glycan modification strategy may apply to all viral GPs, enriching the GP glycan shield with oligomannose-type glycans improved filovirus vaccine-induced NAb responses in mice. Together, these findings provide a foundation for developing broadly effective vaccines to combat future filovirus outbreaks.

## RESULTS

### A universal design strategy for orthoebolavirus GP stabilization

GPs within the *Orthoebolavirus* genus share a high degree of sequence and structural similarity (*59, 116*). These viral GPs consist of conserved functional domains, including GP1, MLD, internal fusion loop (IFL), and GP2, which contains heptad repeat 1 (HR1), heptad repeat 2 (HR2), and the transmembrane domain (TMD). Previously, we stabilized EBOV GPΔmuc by introducing a T577P (P^2^) or L579P (P^4^) mutation in the HR1_C_ bend, a W615L mutation in the neck region of the HR2 stalk, and a C-terminal extension in HR2 (D632–D637) (*100*) (**Fig. 1a**). A foldon trimerization motif was appended to the C terminus of HR2. These modifications enhanced the structural and thermal stability of EBOV GPΔmuc trimers compared with wild-type GP.

**Fig. 1.**
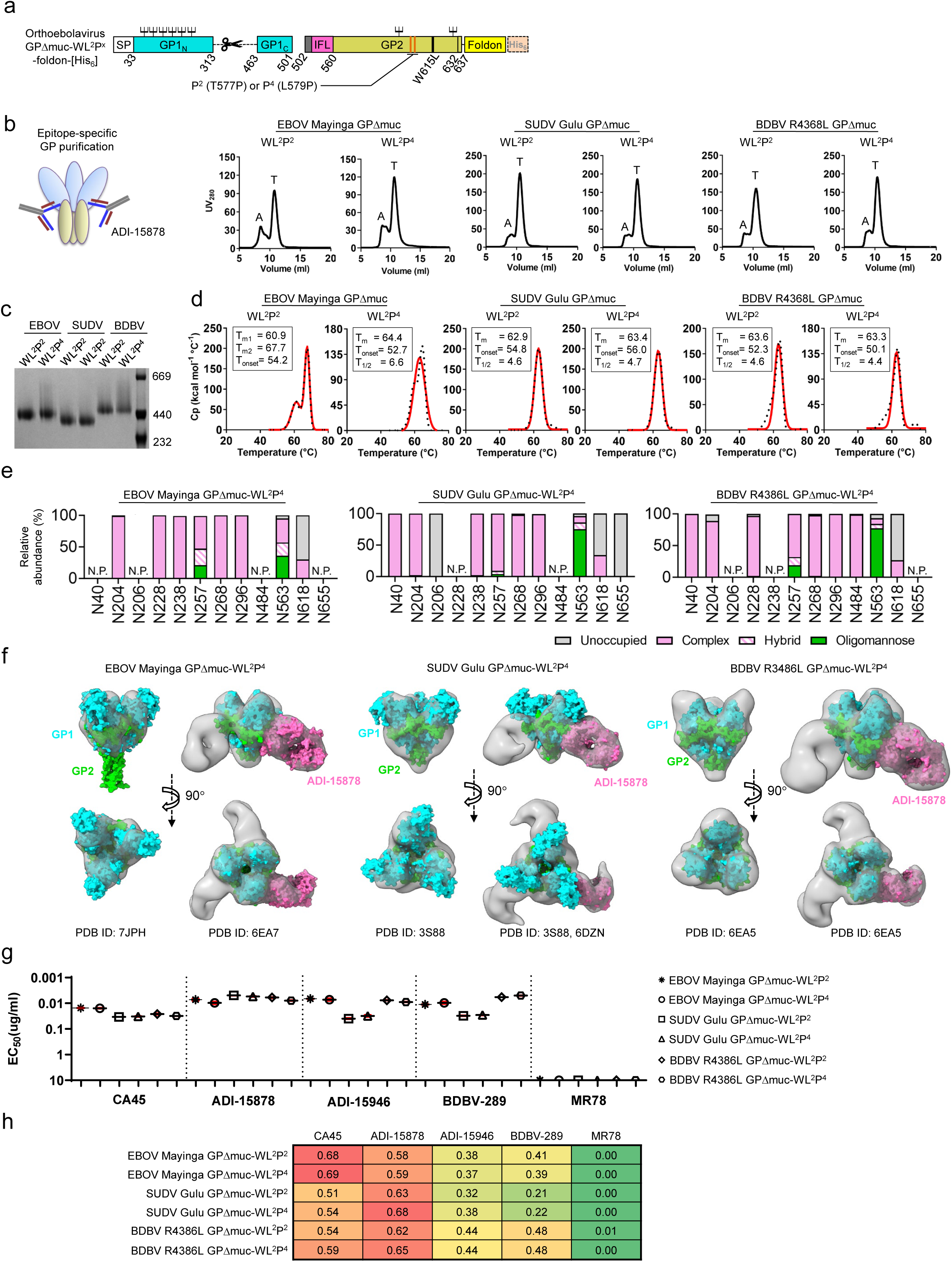
Rational design and in vitro characterization of orthoebolavirus GP trimers. (**a**) Schematic of GP construct design, including MLD deletion (Δmuc, residues G314–N463), a P^2^ (T577P) or P^4^ (L579P) mutation in the HR1c bend, a W615L mutation at the neck of the HR2 stalk, and a C-terminal extension (D632–D637). IFL: internal fusion loop. (**b**) Schematic of an orthoebolavirus GP trimer bound to NAb ADI-15878 (left), and SEC profiles of six GPΔmuc constructs expressed in 500 ml HEK293F cultures and purified using an ADI-15878 IAC column (right). Aggregation (A) and trimer (T) peaks are indicated. (**c**) BN-PAGE analysis of six ADI-15878/SEC-purified GPΔmuc trimers, showing bands consistent with trimer size and molecular weight. (**d**) DSC thermograms of six ADI-15878/SEC-purified GPΔmuc trimers. Experimental data (black dots) and Gaussian fits (red lines) are shown, with key parameters labeled: melting temperature(s) (T_m_, or T_m1_ and T_m2_), onset temperature (T_onset_), and peak half-width (ΔT_1/2_). (**e**) Site-specific glycan profiles of ADI-15878/SEC-purified EBOV, SUDV, and BDBV GPΔmuc-WL^2^P^4^ trimers. Glycan compositions are grouped into four types, including complex (pink), hybrid (pink striped), oligomannose (green), and unoccupied (gray). “N.D.”: not determined; “N.P.”: not present. (**f**) nsEM-derived 3D reconstructions of EBOV, SUDV, and BDBV GPΔmuc-WL^2^P^4^ trimers and their complexes with NAb ADI-15878. Crystal structures of EBOV GPΔmuc-WL^2^P^2^ (PDB ID: 7JPH) and ADI-15878 (PDB ID: 6DZN) were used in the density fitting, with GP1, GP2, and Fab shown in cyan, green, and pink, respectively. (**g**) ELISA-derived EC_50_ (µg/ml) values for the six orthoebolavirus GPΔmuc trimers binding to four pan-orthoebolavirus NAbs and the MARV NAb, MR78. If absorbance at 450 nm was <0.5 at the starting concentration (10 μg/ml), the antigen was considered a non-binder with the EC_50_ value set to 10 μg/ml. (**h**) BLI profiles for the same six GPΔmuc trimers binding to the same NAbs. Sensorgrams were collected using a six-point, 2-fold antigen dilution (starting at 600 nM) from an Octet RED96 instrument and are shown in **Fig. S1i**. Peak responses at the highest concentration are summarized in a matrix, color-coded from green (weak binding) to red (strong binding).

Here, we applied the same design principles to SUDV and BDBV GPs and characterized two constructs, GPΔmuc-WL^2^P^2^ and GPΔmuc-WL^2^P^4^, across all three orthoebolaviruses (**Fig. S1a**). Following our previous protocol (*100*), all six GPΔmuc constructs were transiently expressed in HEK293F cells and purified using two methods. In addition to immobilized nickel affinity chromatography, pan-orthoebolavirus NAbs ADI-15878 (*117*) and CA45 (*118*), expressed in immunoglobulin G (IgG) form in ExpiCHO cells, were used to prepare immunoaffinity chromatography (IAC) columns. The IAC method was established in our prior study, where mAb100 and mAb114 (*36*) columns were used to purify EBOV GP trimers and SApNPs (*100*). The IAC- or nickel-purified orthoebolavirus GP samples were then characterized by size-exclusion chromatography (SEC) on a Superdex 200 column following ADI-15878 (**Fig. 1b)** and CA45 (**Fig. S1b**) purification. All SEC profiles displayed an aggregation peak around 8.0 ml and a consistent trimer peak at ∼11.0 ml, with minimal evidence of dimer or monomer species. Compared to IAC, nickel purification resulted in higher aggregation for EBOV and SUDV GPs (**Fig. S1b**). Of the two broadly neutralizing antibodies (bNAbs) tested, ADI-15878 captured orthoebolavirus GPs more efficiently than CA45, as indicated by stronger ultraviolet absorbance at 280 nm (UV_280_) in the elution fractions (**Fig. 1b** and **S1b**). These results support the use of ADI-15878 as a universal affinity ligand for tag-free purification of orthoebolavirus GP immunogens. Blue native polyacrylamide gel electrophoresis (BN-PAGE) confirmed that SEC fractions eluting at ∼11 ml corresponded to GPΔmuc trimers, with bands consistent with theoretical molecular weights (**Fig. 1c**). Thermostability was assessed by differential scanning calorimetry (DSC). For EBOV, the GPΔmuc-WL^2^P^2^ trimer showed two melting temperatures at 60.9 ⁰C (T_m1_) and 67.7 ⁰C (T_m2_), whereas GPΔmuc-WL^2^P^4^ had a single, broadened melting peak at 64.4 ⁰C (**Fig. 1d**), consistent with our previous findings (*100*). In contrast, all four SUDV and BDBV GPΔmuc trimers generated similar DSC curves with a single denaturing state and tightly clustered T_m_ values (62.9– 63.6 ⁰C). Among the three orthoebolaviruses, SUDV GPΔmuc exhibited the highest onset temperatures (T_onset_, 54.8 ⁰C and 56.0 ⁰C), followed by EBOV and BDBV GP constructs (50.1– 54.2 ⁰C). Based on these biochemical and biophysical profiles, GPΔmuc-WL^2^P^4^ was selected as the lead design candidate for pan-orthoebolavirus GP stabilization.

The HIV-1 Env spike employs a glycan shield as a primary defense mechanism to evade host NAb responses (*113*). Similarly, orthoebolavirus GPs contain a glycan cap that shields the receptor binding site (RBS) (*100*), along with a heavily glycosylated MLD, which is removed in GPΔmuc constructs. These glycans play a critical role in both immune evasion and modulating antibody interactions with GP. To generate site-specific glycosylation profiles, the purified EBOV, SUDV, and BDBV GPΔmuc-WL^2^P^4^ proteins were digested with trypsin, chymotrypsin, and α-lytic protease to produce glycopeptides, each containing a single N-linked glycan site. Site-specific composition and occupancy were analyzed by liquid chromatography-mass spectrometry (LC-MS), as previously described (*113*). All glycosylation sites on the three orthoebolavirus GPΔmuc-WL^2^P^4^ proteins predominantly carried complex-type glycans, except at positions N563 and N257, where mixed glycan forms were detected. At N563, ∼30% of glycans were oligomannose-type for EBOV, compared to ∼60% for both SUDV and BDBV. At N257, ∼15% oligomannose-type glycans were observed for EBOV (**Fig. 1e** and **Fig. S1c**). Due to technical limitations, glycan compositions could not be assessed accurately at all sites.

Negative-stain EM (nsEM) was used to characterize orthoebolavirus GPΔmuc-WL^2^P^4^ structures, either alone or bound to bNAb ADI-15878 (**Fig. S1d**). For EBOV and BDBV, crystal structures of wild-type GPΔmuc in complex with ADI-15878 (Protein Data Bank [PDB] IDs: 7JPH (*100*), 6EA7, and 6EA5 (*116*)) were fitted into three-dimensional (3D) EM reconstructions. For SUDV, the GPΔmuc and ADI-15878 structures from two separate complexes (PDB IDs: 3S88 (*59*) and 6DZM (*117*)) were fitted individually (**Fig. 1f** and **Figs. S1e-g**). Overall, the low-resolution nsEM models of three orthoebolavirus GPΔmuc trimers displayed a three-lobed chalice shape consistent with the native-like structure, although the HR2 stalk was unresolved. The ADI-15878-bound nsEM models were highly similar across all three species, revealing an epitope that spans two GP protomers and overlaps the conserved regions of the IFL and HR1 in GP2 (*48, 117*).

The antigenicity of six stabilized orthoebolavirus GPΔmuc trimers was assessed using four pan-orthoebolavirus bNAbs – CA45 (*47*), ADI-15878, ADI-15946 (*119*), and BDBV-289 (*52*) – alongside MR78, a MARV-specific NAb (*40*). In enzyme-linked immunosorbent assays (ELISA), the proline mutation in HR1c (P^2^ or P^4^) had no effect on antibody binding (**Fig. 1g** and **Fig. S1h**). Among the four bNAbs, ADI-15946 and BDBV-289 showed the lowest binding affinity for SUDV GPΔmuc trimers, 2.5- to 6-fold weaker than those for EBOV and BDBV GPΔmuc trimers, as indicated by the half-maximal effective concentration (EC_50_) values. The NHP-derived bNAb CA45 exhibited 1.5- to 2.5-fold stronger binding to EBOV GPΔmuc than to its SUDV and BDBV counterparts. As expected, the negative control MR78 showed no detectable binding to any of the GPΔmuc trimers from the *Orthoebolavirus* genus. These findings were confirmed by biolayer interferometry (BLI) (**Fig. 1h** and **Fig. S1i**). Among the six orthoebolavirus antigens tested, CA45 bound most strongly to EBOV GPΔmuc, whereas ADI-15946 and BDBV-289 interacted least favorably with SUDV GPΔmuc. Consistent with ELISA, BLI revealed negligible or no binding between orthoebolavirus GPΔmuc trimers and MARV-specific NAb MR78.

### Structural characterization of rationally designed orthoebolavirus GPΔmuc trimers

Previously, we determined the crystal structure of EBOV GPΔmuc-WL^2^P^2^ (*100*). Here, we sought to obtain atomic details for the more stable EBOV GPΔmuc-WL^2^P^4^. To this end, EBOV GPΔmuc-WL^2^P^4^ was transiently expressed in HEK293S cells and purified by IAC (mAb100) and SEC, as previously described (*100*) (**Fig. S2a**). Crystallization under conditions similar to those used for GPΔmuc-WL^2^P^2^ resulted in comparable packing in the P321 space group (**Table S1**). The final structure was resolved at 3.2 Å. Each asymmetric unit contained one GP protomer, with the glycan cap atop GP1 and the belt-forming IFL of GP2 clearly visible in the electron density (**Fig. 2a**). Three GP1-GP2 heterodimers assembled into a chalice-shaped trimer in the crystal lattice, with the GP2 subunits forming a base that cradles the outward-facing GP1 lobes (**Fig. S2b**). Four N-linked glycans (N228, N257, N563, and N618) were identified in the structure. The most complete carbohydrate chain was observed at N563 (with Man_4_GlcNAc_2_ density), part of the ADI-15878 epitope in HR1a (*116*), whereas only a single *N*-acetylglucosamine (GlcNAc) was visible at the other three sites. The backbone of GPΔmuc-WL^2^P^4^ closely resembled that of GPΔmuc-WL^2^P^2^, with a global Cα root-mean-square deviation (RMSD) of 0.48 Å across 243 aligned Cα atoms and a local Cα-RMSD of 0.68 Å for the HR1c bend following protomer superposition (**Fig. 2a** and **Fig. S2c**). In GPΔmuc-WL^2^P^4^, the amine group of the K95 side chain formed hydrogen bonds with the backbone carbonyls of L573 and T576 (∼3.0 Å) – interactions absent in GPΔmuc-WL^2^P^2^. These interprotomer hydrogen bonds at the trimer center likely contributed to the enhanced thermostability of GPΔmuc-WL^2^P^4^. Similar interactions were observed in the EBOV GPΔmuc-WL^2^ structure (PDB ID: 7JPI), where K95 formed hydrogen bonds with L573 and T576 at 2.7 and 2.9 Å, respectively. This structural analysis is consistent with the DSC data, in which EBOV GPΔmuc-WL^2^P^4^ exhibited a single denaturation transition with a T_m_ of 64.4°C (*100*).

**Fig. 2.**
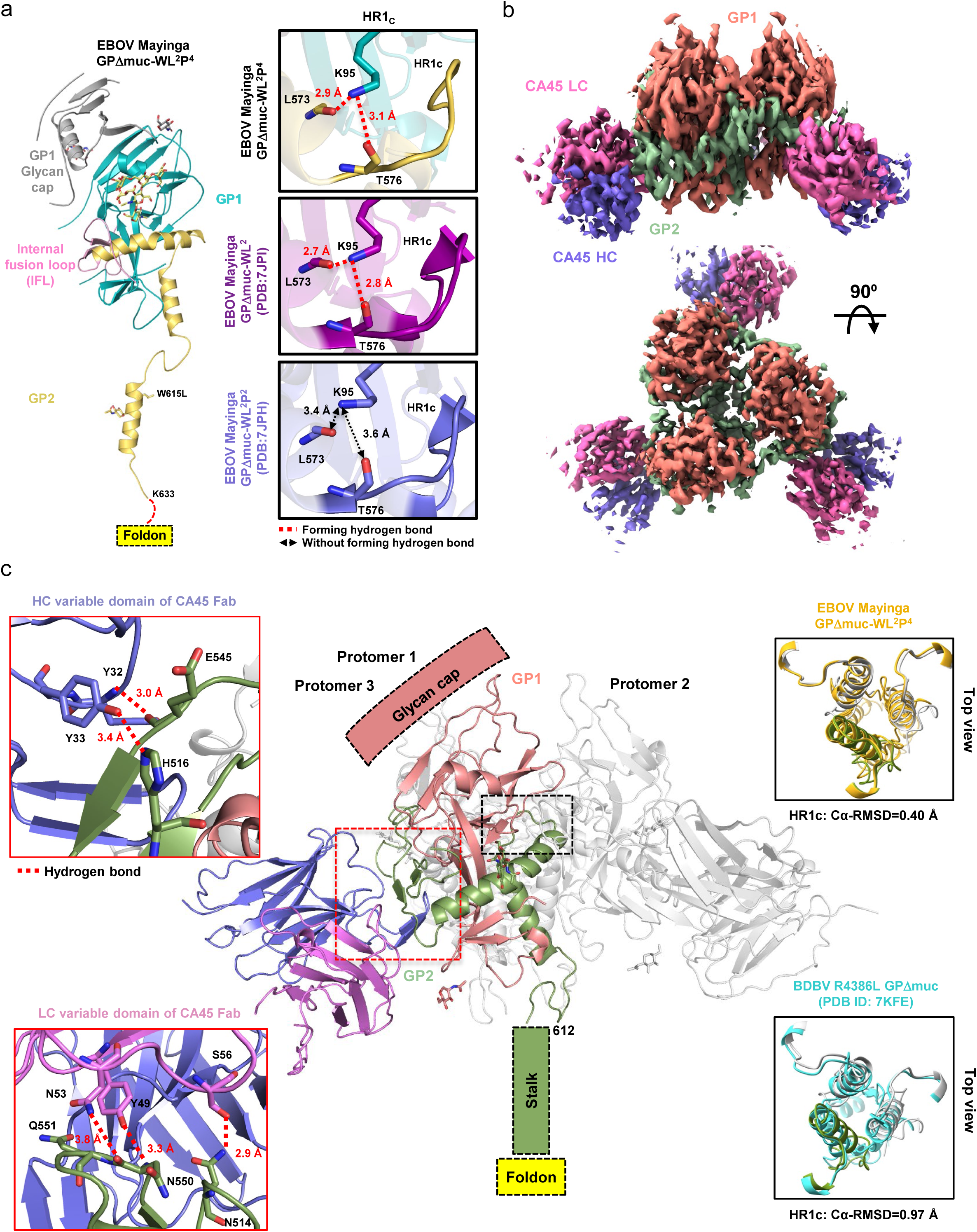
Structural characterization of an unliganded EBOV GPΔmuc-WL^2^P^4^ trimer and a SUDV GPΔmuc-WL^2^P^4^ trimer bound to NAb CA45. (**a**) Left: crystal structure of the stabilized EBOV GPΔmuc-WL^2^P^4^ trimer determined at 3.2 Å resolution, with GP1 in cyan, glycan cap in gray, IFL in pink, and GP2 in gold. Structural details could not be resolved for residues R200-P209, K294-I301, and T634-D637 (C terminus). Right: comparative analysis of the HR1c bend and its local structure for EBOV GPΔmuc constructs bearing WL^2^ (PDB ID: 7JPI), WL^2^P^2^ (PDB ID: 7JPH), and WL^2^P^4^ mutations. Side chains are shown for residues forming cross-protomer interactions (L95, R130, L573, and T576). A 3.3 Å cutoff was used to define donor-acceptor distances for hydrogen bonds. (**b**) Cryo-EM 3D reconstruction of the SUDV GPΔmuc-WL^2^P^4^ trimer bound to CA45 Fab at 3.13 Å resolution, with GP1 in orange, GP2 in green, CA45 heavy chain in blue, and light chain in pink. The glycan cap and HR2 stalk were not resolved and became partially visible at lower density thresholds beyond 4.0 Å (see **Fig. S2e**). (**c**) Ribbon representation of the cryo-EM-derived SUDV GPΔmuc-WL^2^P^4^/CA45 complex using the color scheme. Atomic details of the glycan cap and HR2 stalk could not be resolved due to limited local resolution. Left: atomic details of hydrogen bonds between GP and the CA45 heavy chain (top) and light chain (bottom). Right: superposition of the HR1c bend in SUDV GPΔmuc-WL^2^P^4^ with that of BDBV GPΔmuc (top, cryo-EM) and EBOV GPΔmuc-WL^2^P^4^ (bottom, crystal structure).

Cryo-EM plays an important role in understanding how viral GPs interact with antibodies (*120–122*). Several cryo-EM models have been reported for EBOV and BDBV GPs (*43, 52, 117*), but only two crystal structures (*59, 123*) and no cryo-EM models are currently available for SUDV GP. Here, we determined a cryo-EM structure of SUDV GPΔmuc-WL^2^P^4^ in complex with the CA45 Fab. SUDV GP was transiently expressed in HEK293F cells and purified using an ADI-15878 IAC column, prior to complexation with CA45 Fab and SEC on a Superose 6 16/600 GL column (**Fig. S2d**). The complex fraction at ∼72 ml was imaged on a Krios G4 transmission electron microscope (TEM), with data processed in CryoSPARC (*124*) (**Fig. S2e**). A resolution of 3.13 Å with an EMRinger score of 2.60 (*125*) was obtained for the final reconstruction, in which the SUDV GPΔmuc-WL2P4 trimer flanked by three CA45 Fabs was well resolved in the density map (**Fig. 2b**). Three regions – GP1 (A187-L218), GP2 (E522-N525), and C-terminal stalk (P612-N637) – were excluded from the final model due to limited local resolution (**Fig. S2f**). The glycan cap and GP2 stalk became partially visible when the density threshold was lowered beyond 4.0 Å (**Fig. S2e**). Crystal structures of EBOV GPΔmuc-WL^2^P^2^ (PDB ID: 7JPH) (*100*) and CA45 variable domains (PDB ID: 6EAY) (*118*) were used as starting models for EM fitting, followed by manual refinement. The final SUDV GPΔmuc-WL^2^P^4^ model, lacking the glycan cap and GP2 stalk, was superimposed onto the EBOV GPΔmuc-WL^2^P^4^ crystal structure, yielding a Cα-RMSD of 0.66 Å for 154 matching Cα atoms (**Fig. S2g**, left). The three HR1_C_ bends of SUDV GPΔmuc-WL^2^P^4^were also superimposed onto those of EBOV GPΔmuc-WL^2^P^4^ from this study and BDBV GPΔmuc (PDB ID: 7KFE) (*52*), with Cα-RMSDs of 0.34 and 0.53 Å for the 24 aligned Cα atoms, respectively (**Fig. 2c**, right). Compared with the crystal structure of the EBOV GPΔmuc/CA45 complex (PDB ID: 6EAY) (*118*), cryo-EM revealed a highly similar antibody epitope in the SUDV GPΔmuc-WL^2^P^4^/CA45 complex, with a Cα-RMSD of 0.69 Å for 147 aligned Cα atoms after superposition of a single GP/CA45 component from each complex (**Fig. S2g**, right). Specifically, the CA45-GP2 interface was stabilized by a backbone hydrogen bond between CA45 HC-Y32 and GP2-E545 (∼3.0 Å), a side chain hydrogen bond between the CA45 HC-Y33 carboxyl group and the GP2-H516 imidazole ring (∼3.4 Å), and additional hydrogen bonds involving the CA45 LC-Y49 and LC-N53 side chains with the GP2-N550 backbone (∼3.3 Å and ∼3.8 Å, respectively), as well as the CA45 LC-S56 backbone with the GP2-N514 side chain (∼2.9 Å) (**Fig. 2c**). Although interpretation of these interactions may be limited by the cryo-EM resolution, similar contacts were observed in the 3.7 Å-resolution crystal structure of the EBOV GPΔmuc/CA45 complex (PDB ID: 6EAY) (**Fig. S2h**) (*118*), including those between CA45 HC-Y32 and GP2-E545 (∼3.2 Å), CA45 HC-D101 and GP2-H549 (∼3.9 Å), and the same CA45 LC-GP2 residue pairs with nearly identical distances. Snapshots of the cryo-EM density at the GP-CA45 interface were inspected fragment by fragment for anisotropy, demonstrating that the modeled side chain contacts are supported by well-defined density (**Fig. S2i and S2j**). Furthermore, we compared our SUDV GPΔmuc-WL^2^P^4^/CA45 complex with previously reported cryo-EM models of EBOV GPΔmuc bound to ADI-15878 (PDB ID: 6DZL and EMD-8935 (*117*)) (**Fig. S2k**), rEBOV-515 and rEBOV-442 (PDB ID: 7M8L and EMD-23719 (*126*)) (**Fig. S2l**), and rEBOV-520 and rEBOV-548 (PDB ID: 6PCI and EMD-20301 (*50*)) (**Fig. S2m**), demonstrating comparable model quality.

### Rational design and characterization of orthoebolavirus GPΔmuc-presenting nanoparticles

In our previous EBOV study, a 10GS linker was used to connect GPΔmuc to the N-termini of I3-01v9, which formed a triangle of 55.0 Å (*100*). This SApNP exhibited variability in yield, purity, and stability, suggesting that further optimization might be necessary (*100*). In a more recent study, we redesigned the I3-01v9 SApNP to display monomeric antigens (*127*). The resulting I3-01v9a scaffold featured a smaller triangle of 43.9 Å between the N-termini (**Fig. 3a**, left). Here, using I3-01v9a as a template, we aimed to improve the design of an I3-01v9 scaffold optimized for surface display of filovirus GP trimers, which contain a narrow stalk. After removing the first 4 amino acids (aa) of I3-01v9a, we fused a 9-aa helix to the truncated I3-01v9a *N*-terminal helix via a 4-aa turn. The resulting model was computationally optimized – first in structure and then in sequence – to generate two I3-01v9 variants, termed I3-01v9b and I3-01v9c (**Fig. S3a**), in which the N-termini form a triangle of 12.9 Å (**Fig. 3a**, right). The trimeric forms of I3-01v9b/c, termed I3-01v9b/c-T, were generated by interrupting the particle-forming interface based on the structure of the aldolase (PDB ID: 1VLW), from which I3-01 was derived. Two “scaffolded” EBOV GP constructs, termed GPΔmuc-WL^2^P^4^-I3-01v9b-T and GPΔmuc-WL^2^P^4^-I3-01v9c-T, were designed to test whether the I3-01v9b/c N-termini could accommodate the GP2 stalk (**Fig. S3b**). Both constructs were transiently expressed in HEK293S cells, purified using a mAb100 column, and analyzed by SEC (**Fig. S3c**). The trimer fractions were validated by BN-PAGE (**Fig. S3d**) prior to nsEM analysis, which yielded two-dimensional (2D) classes consistent with native-like GPΔmuc-WL^2^P^4^ trimers anchored to I3-01v9b/c-T (**Figs. S3e** and **S3f**). The GP2 stalk, although not visible by EM, was likely well-formed, as the GPΔmuc trimer displayed the classic chalice shape (**Fig. 3b** and **Fig. S3g**). Computational modeling suggested that EBOV GPΔmuc-WL^2^P^4^-I3-01v9b would form a particle with a diameter of 49.1 nm, measured at GP1-T269 (**Fig. 3c**).

**Fig. 3.**
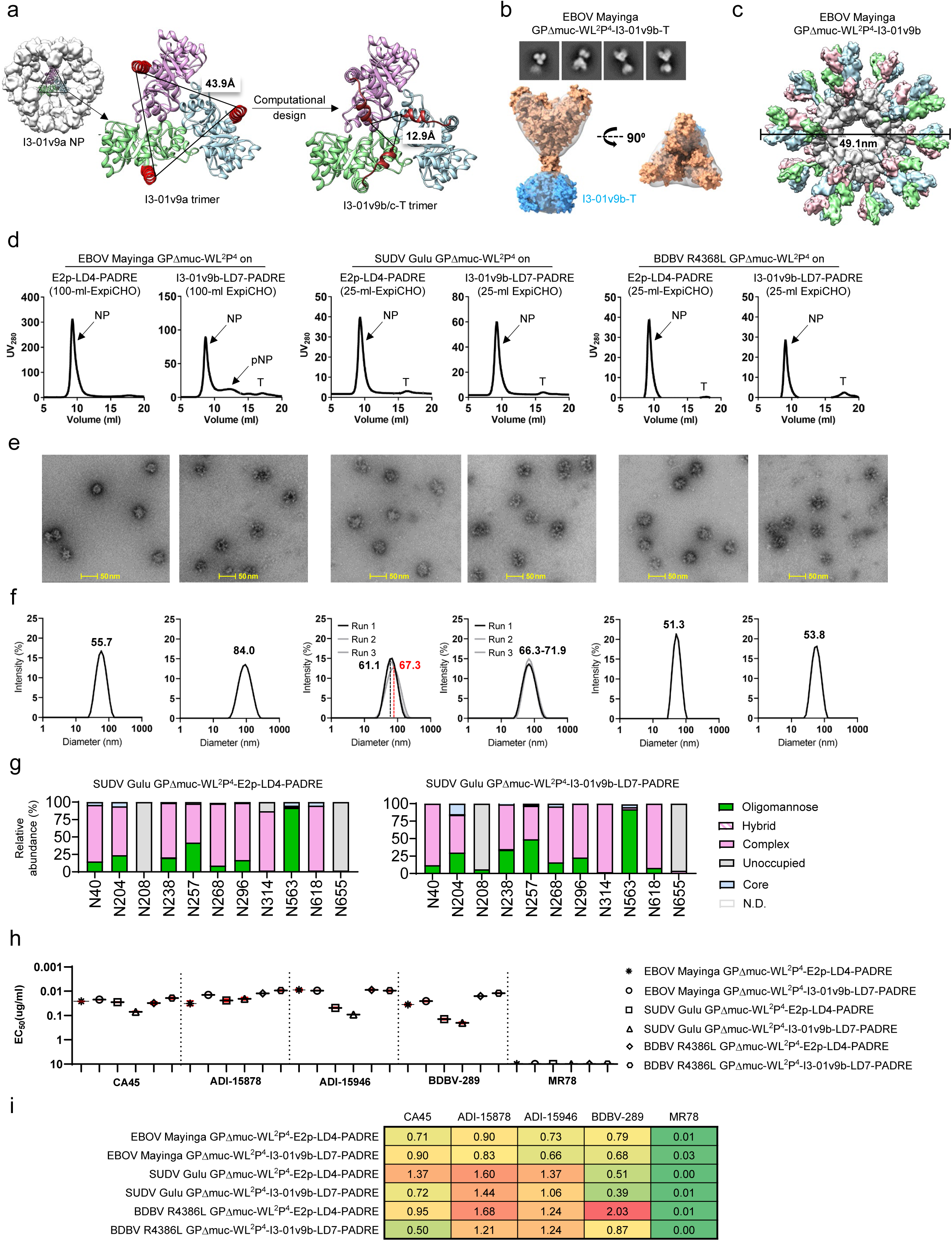
Rational design and in vitro characterization of orthoebolavirus GP-presenting SApNPs. (**a**) SApNP optimization for surface display of trimeric antigens with a narrow stalk. Twenty I3-01v9a trimers assemble into an SApNP (left), with the N-termini forming a triangle of 43.9 Å (middle). Using computational design, a helical extension was added to the I3-01v9a N-terminus to pack against the protein core, reducing the triangle to 12.9 Å and yielding I3-01v9b/c (right). (**b**) nsEM analysis of EBOV GPΔmuc-WL^2^P^4^-I3-01v9b-T, in which I3-01v9b-T is a trimeric version of I3-01v9b with mutations that interrupt particle assembly. Top: representative 2D class averages; Bottom: side and top views of the 3D reconstruction with fitted EBOV GPΔmuc-WL^2^P^4^ and aldolase (PDB: 1VLW) structures. (**c**) Schematic of EBOV GPΔmuc-WL2P4-presenting I3-01v9b/c SApNP. The predicted particle diameter is 49.1 nm, measured at GP1 residue T269. (**d**) SEC profiles of six orthoebolavirus GPΔmuc-WL^2^P^4^-presenting SApNPs expressed in ExpiCHO cells and purified using an ADI-15878 IAC column. Peaks corresponding to full NPs, partially assembled NPs (pNP), and trimers (T) are indicated. (**e**) Representative nsEM images of the six SEC-purified SApNPs. Scale bar: 50 nm. (**f**) Particle size distributions of the six SEC-purified SApNPs measured by DLS. Three independent measurements are shown for each construct, with hydrodynamic diameters (Dₕ) labeled. (**g**) Site-specific glycan profile of SUDV GPΔmuc-WL^2^P^4^-presenting E2p and I3-01v9b SApNPs. Glycan types are colored as in Fig. 1e. (**h**) ELISA-derived EC_50_ (µg/ml) values for six orthoebolavirus GPΔmuc-WL^2^P^4^-presenting SApNPs binding to five NAbs. Absorbance <0.5 at 10 µg/ml was used as a non-binding cutoff, as in Fig. 1g. (**i**) BLI profiles for the same six SApNPs binding to the same five NAbs. Sensorgrams were obtained using the same protocol as in Fig. 1h (see **Fig. S3k**). Peak responses at the highest concentration are summarized in a matrix as in Fig. 1h.

Six SApNP constructs – displaying EBOV, SUDV, and BDBV GPΔmuc-WL^2^P^4^ trimers on the E2p and I3-01v9b scaffolds (**Fig. S3h**) – were transiently expressed in ExpiCHO cells and purified by IAC using an ADI-15878 column, followed by SEC on a Superose 6 increase 10/300 GL SEC column. All constructs exhibited a peak at ∼8.7 ml, corresponding to assembled protein particles (**Fig. 3d**). For EBOV and BDBV, higher yields were observed for the E2p SApNPs, whereas for SUDV, 3-01v9b produced a more prominent particle peak in the SEC profile. The structural integrity of these SApNPs was validated by nsEM (**Fig. 3e**). Most micrographs showed well-formed protein particles displaying arrays of GP trimers on the outer surface. Particle size distributions were analyzed in solution by dynamic light scattering (DLS) (**Fig. 3f**). For EBOV SApNPs, E2p and I3-01v9b exhibited sizes of 55.7 nm and 84.0 nm, respectively (**Fig. 3f**). The unusually large DLS size for I3-01v9b may reflect SApNP aggregation, although this form was rarely observed by EM. For SUDV and BDBV SApNPs, particle sizes ranged from 61.1 to 71.9 nm and from 51.3 to 53.8 nm, respectively. This variation may be attributed to the intrinsic structural flexibility of the glycan cap that likely affects the hydrodynamic radius measured by DLS. Site-specific glycosylation profiles were generated for SUDV GPΔmuc-presenting SApNPs (**Fig. 3g** and **Fig. S3i**). An increase in oligomannose-type glycans was observed on SApNPs compared to soluble trimers. At site N563, both the soluble GPΔmuc trimer and SApNPs presented predominantly oligomannose-type glycans (**Fig. 3g**).

The antigenicity of orthoebolavirus GPΔmuc-WL^2^P^4^-presenting SApNPs was assessed against the same antibody panel. In ELISA, SApNPs exhibited broadly similar antibody-binding patterns to the corresponding soluble trimers (**Fig. 3h** and **Fig. S3j**). For example, SUDV SApNPs bound less favorably to ADI-15946 and BDBV-289 than EBOV and BDBV SApNPs, with 5.0- to 16.8-fold higher EC_50_ values. Between the two NP scaffolds, E2p outperformed I3-01v9b, showing 1.4- to 1.9-fold lower EC_50_ values for both bNAbs. A similar trend was observed in SUDV SApNP binding to CA45, with a 2.5-fold difference in EC_50_ between the two NP platforms. Interestingly, the opposite pattern was seen for EBOV and BDBV SApNPs, where I3-01v9b bound more tightly than E2p. BLI revealed a somewhat different antigenic profile (**Fig. 3i** and **Fig. S3k**). Most notably, SUDV and BDBV SApNPs exhibited stronger binding signals to ADI-15878 and ADI-15946 than EBOV SApNPs, whereas SUDV SApNPs showed the weakest binding to BDBV-289 among the three orthoebolaviruses. For each GP antigen, binding signals to individual bNAbs were generally similar between E2p and I3-01v9b, except in the cases of SUDV SApNPs with CA45 and BDBV SApNPs with CA45 and BDBV-289, where scaffold-dependent differences were apparent.

In this study, we displayed the stabilized GPΔmuc-WL^2^P^4^ trimer on two SApNP scaffolds, E2p and the newly designed I3-01v9b, for three representative orthoebolaviruses. SEC, nsEM, and DLS were used to assess particle properties, which were largely comparable to those of our previously reported HIV-1 Env trimer-presenting SApNPs (*113*). However, site-specific glycan analysis and antigenic profiling revealed distinct features unique to the SApNP platform, including an increased proportion of oligomannose-type glycans.

### In vitro effect of glycan modification on orthoebolavirus GPΔmuc trimers and SApNPs

Previously, we investigated the effect of glycan modification on rationally designed HIV-1 Env trimer and SApNP vaccines (*113*). Supplementing ExpiCHO cultures with kifunensine (Kif) – a class I α-mannosidase inhibitor that blocks ER glycan processing and leads to accumulation of high mannose – did not affect NAb responses. In contrast, glycan trimming with endoglycosidase H (Endo H) significantly enhanced NAb elicitation, as reflected by a higher vaccine responder rate (*113*). The benefit of glycan trimming has also been reported for influenza HA and SARS-CoV-2 spike vaccines (*112, 114*). These findings highlight the importance of examining how glycan modification may affect filovirus vaccines in vivo. However, the glycan-modified filovirus GP immunogens first needed to be generated and characterized in vitro.

Here, we adopted a similar strategy (*113*) to produce orthoebolavirus GPΔmuc trimers carrying either oligomannose-only glycans or a single layer of mono-*N*-acetylglucosamine (mono-GlcNAc) glycans. EBOV and SUDV GPΔmuc-WL^2^P^4^ were transiently expressed in HEK293F cells in the presence of Kif, which yielded oligomannose-only glycans, and were purified using an ADI-15878 column followed by SEC. The SEC profiles showed a trimer peak at ∼10.7 ml, similar to wild-type trimers (**Fig. 4a**). SEC-purified, Kif-treated GPΔmuc (e.g., 1 mg) was then treated with Endo H (termed Kif/Endo H) to reduce the glycan shield to mono-GlcNAc stumps, with the enzyme subsequently removed by SEC. The resulting profiles showed a trimer peak at ∼11.1 ml and an Endo H peak at ∼13.9 ml (**Fig. 4a**). Sodium dodecyl sulfate (SDS)-PAGE and BN-PAGE confirmed that Kif/Endo H-treated GPΔmuc had a lower molecular weight than both wild-type and Kif-treated GPΔmuc (**Fig. S4a**). DSC revealed a differential effect of glycan modification on GPΔmuc thermostability (**Fig. 4b**). The Kif-treated EBOV GPΔmuc-WL^2^P^4^ trimer exhibited two denaturation states, with T_m1_ at 58.2 ⁰C and T_m2_ at 64.3 ⁰C. Endo H treatment further reduced T_m1_ and T_m2_ by 5.2 ⁰C and 2.6 ⁰C, respectively. In contrast, the Kif-treated SUDV GPΔmuc-WL^2^P^4^ trimer remained thermostable, displaying the same T_m_ (63.6 ⁰C) as the wild-type trimer, with T_onset_ measured at 54.7 ⁰C. Endo H treatment slightly reduced the T_m_ to 62.6 ⁰C and lowered the T_onset_ to 48.9 ⁰C. Antigenicity was assessed using the same antibody panel. In the ELISA (**Fig. 4c and Fig. S4b**), Endo H treatment enhanced trimer binding to CA45, ADI-15878, and ADI-15946, which recognize a quaternary epitope near the trimer base with few glycan contacts, but reduced binding to BDBV-289, which targets the glycan cap and relies on glycans at N238 and N268. However, the EC_50_ differences were small (1.5- to 1.9-fold) and not specific to either orthoebolavirus. In the BLI analysis, glycan-modified trimers generated slightly higher binding signals than wild-type trimers (**Fig. 4d** and **Fig. S4c**). Overall, glycan modification had only a modest impact on the biochemical, biophysical, and antigenic properties of stabilized orthoebolavirus GPΔmuc trimers, except for thermostability in the EBOV constructs.

**Fig. 4.**
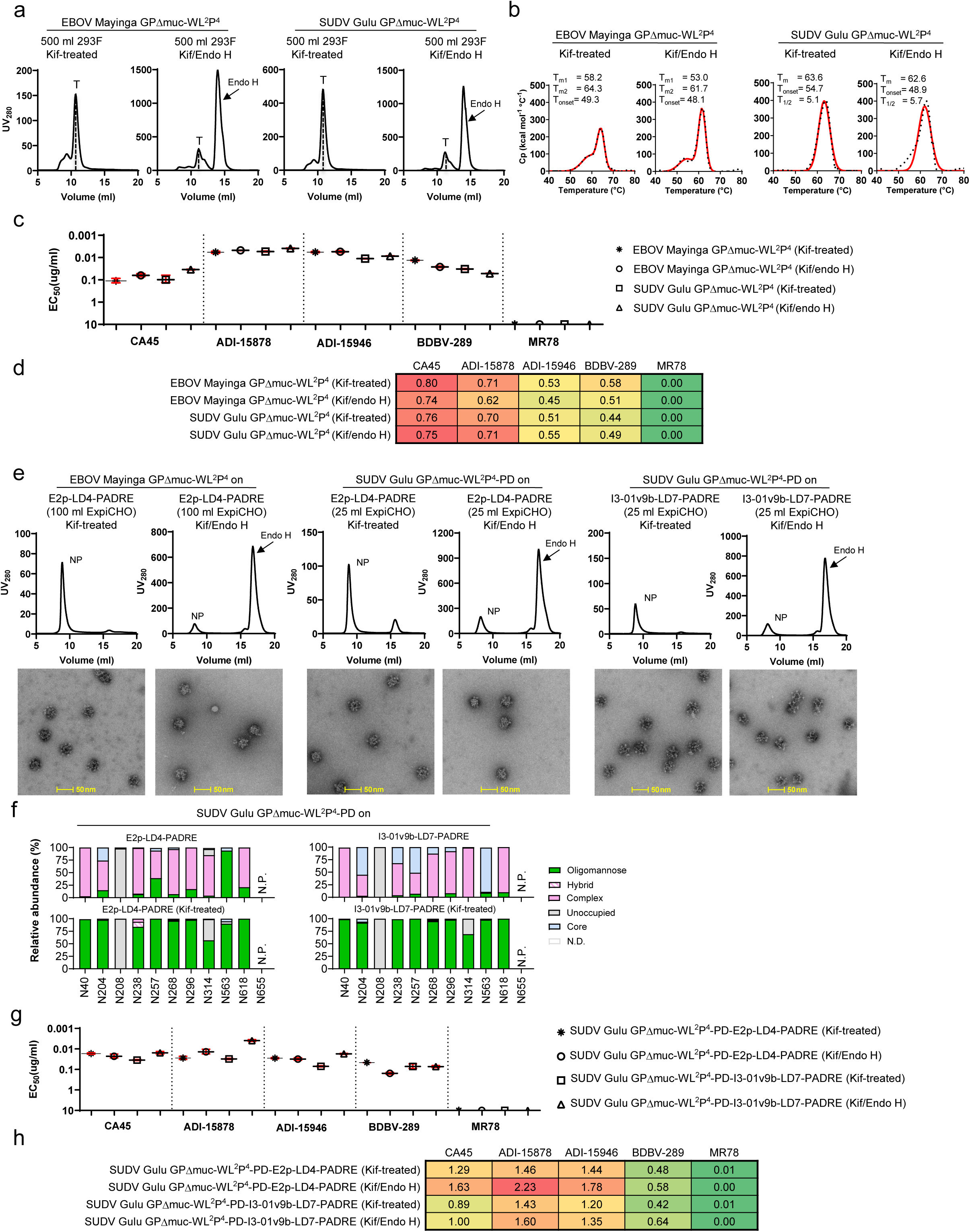
Glycan modification of EBOV and SUDV GPΔmuc-WL^2^P^4^ trimers and SApNPs. (**a**) SEC profiles of EBOV and SUDV GPΔmuc-WL^2^P^4^ constructs expressed in 500 ml HEK293F cultures and purified using an ADI-15878 IAC column. For each construct, two types of glycan modifications were performed: kifunensine (Kif) treatment and Kif plus Endo H (Kif/Endo H) treatment. HEK293F expression in the presence of Kif enriches oligomannose-type glycans, and subsequent Endo H treatment yields GlcNAc stumps. The SEC peak at 14.7 ml corresponds to Endo H. (**b**) DSC thermograms of ADI-15878/SEC-purified EBOV and SUDV GPΔmuc-WL^2^P^4^ trimers after glycan modification. Experimental data and Gaussian fits are shown as black dots and red lines, respectively. Melting temperatures (T_m_ or T_m1_/T_m2_), onset temperature (T_onset_), and peak half-width (ΔT_1/2_) are labeled. (**c**) ELISA-derived EC_50_ (µg/ml) values of glycan-modified EBOV and SUDV GPΔmuc-WL^2^P^4^ trimers binding to five NAbs. Absorbance <0.5 at 10 µg/ml was used as a non-binding cutoff, as in Fig. 1f. (**d**) BLI profiles of the same glycan-modified trimers binding to five NAbs. Sensorgrams were obtained using the same protocol as in Fig. 1h (see **Fig. S4c**) Peak responses at the highest concentration are summarized in a matrix, as in Fig. 1h. (**e**) SEC profiles (top) and nsEM images (bottom) of glycan-modified EBOV and SUDV GPΔmuc-WL^2^P^4^- presenting SApNPs. Glycan modification was performed as in panel (a). EBOV GPΔmuc I3-01v9bs are not shown due to low yield. Scale bar: 50 nm. (**f**) Site-specific glycan profiles of SUDV GPΔmuc-WL^2^P^4^-presenting E2p and I3-01v9b SApNPs before and after Kif treatment, with glycan types colored as in Fig. 1e. (**g**) ELISA-derived EC_50_ (µg/ml) values for glycan-modified SUDV GPΔmuc-WL^2^P^4^-presenting E2p and I3-01v9b SApNPs binding to five NAbs. Absorbance <0.5 at 10 µg/ml was used as a non-binding cutoff, as in Fig. 1g. (**h**) BLI profiles of the same glycan-modified SApNPs binding to five NAbs. Sensorgrams were obtained using the same protocol as in Fig. 1h (see **Fig. S4h**). Peak responses are summarized in a matrix, as in Fig. 1h.

We next characterized glycan-modified GPΔmuc-WL^2^P^4^-presenting SApNPs. For SUDV, both E2p and I3-01v9 SApNPs were included in this analysis, whereas for EBOV, only the E2p scaffold was tested due to the low yield of I3-01v9b SApNPs. Notably, an N655D mutation was introduced to eliminate an N-glycan site at the base of the SUDV GP2 stalk that may interfere with SApNP assembly. This design variant, termed SUDV GPΔmuc-WL^2^P^4^-PD, will be used hereafter (**Fig. S4d**). EBOV and SUDV SApNPs were transiently expressed in ExpiCHO cells with Kif and purified by IAC (ADI-15878) followed by SEC. SEC-purified, Kif-treated SApNP was trimmed with Endo H and further purified by SEC. The resulting SEC profiles showed a main peak at ∼8.5-9.0 ml corresponding to SApNPs, and following Endo H treatment, an additional peak at ∼16.8 ml corresponding to the enzyme (**Fig. 4e**). The structural integrity of glycan-modified EBOV and SUDV GPΔmuc-presenting SApNPs was validated by nsEM (**Fig. 4e**). SDS-PAGE confirmed that Kif/Endo H-treated SApNPs had a lower molecular weight than Kif-treated SApNPs (**Fig. S4e**). Because of the low yield of EBOV SApNPs, only SUDV SApNPs were subjected to glycan profiling and antigenic characterization. Site-specific glycan profiles confirmed that all complex glycans in SUDV GPΔmuc were replaced by oligomannose-type glycans in the Kif-treated material (**Fig. 4f** and **Fig. S4f**). As expected, the N655 glycan was absent in the N655D mutant construct. In the ELISA (**Fig. 4g** and **Fig. S4g**), the antigenic profiles of SApNPs appeared to be largely consistent with those of the trimers, though a greater difference in EC_50_ values between Kif and Kif/Endo H-treated samples (1.0- to 7.9-fold) was observed for CA45, ADI-15878, and ADI-15946. In the BLI analysis, all pan-orthoebolavirus bNAbs showed increased binding signals to Kif/Endo H-treated SApNPs, which displayed only a single layer of mono-GlcNAc stumps on the GPΔmuc trimer (**Fig. 4h** and **Fig. S4h**). In summary, these results suggest that glycan trimming enhances antibody access to conserved GP epitopes on the SApNP surface.

### Rational design of a prefusion-stabilized RAVV GPΔmuc trimer

RAVV is one of the two known members of the *Orthomarburgvirus* genus in the filovirus family (*40, 49*). Similar to orthoebolavirus GPs, RAVV GP contains GP1 with a heavily glycosylated MLD and GP2 (**Fig. 5a**). Here, we applied a similar strategy to stabilize the prefusion RAVV GP trimer, using a mucin-deleted construct (ΔN257-T425), termed GPΔmuc. Additional mutations – T578P (P^2^) or E580P (P^4^) in the HR1c bend, and E631F/Q632V at the C terminus (termed CT) – were also evaluated. The five RAVV GP constructs tested were: (1) GPΔTM (truncation at residue 637), (2) GPΔmuc, (3) GPΔmuc-P^2^, (4) GPΔmuc-P^4^, and (5) GPΔmuc-P^2^CT, each containing a C-terminal foldon motif and His_6_ tag (**Fig. S5a**). All constructs were expressed in ExpiCHO cells and purified using a nickel column followed by SEC. Both GPΔTM and GPΔmuc yielded low expression levels with multiple GP species, including aggregates (∼9-10 ml), trimers (∼10.5 ml), dimers (∼13.6 ml), and monomers (∼14.7 ml) (**Fig. 5b**). The P^2^ mutation substantially improved the SEC profiles of GPΔmuc constructs, producing a predominant trimer peak at ∼10.5 ml, along with visible dimer and monomer peaks. In contrast, the P^4^ mutation resulted in significant aggregation, indicated by a peak at ∼8.6 ml (**Fig. S5b**). The GPΔmuc-P^2^CT construct demonstrated an optimal SEC profile, comparable in yield to GPΔmuc-P^2^ but with reduced dimer and monomer content. When MARV NAbs MR78 and MR191 (*49*) were used in IAC purification, they failed to capture GPΔmuc-P^2^ and GPΔmuc-P^2^CT (**Fig. S5c**). Nevertheless, distinct trimer bands were observed by SDS-PAGE after nickel/SEC purification (**Fig. 5c** and **Fig. S5d**). In DSC, GPΔmuc-P^2^ exhibited two melting transitions at 51.0 ⁰C and 60.0 ⁰C, while GPΔmuc-P^2^CT displayed three transitions at 54.5 ⁰C, 58.3 ⁰C, and 66.6 ⁰C (**Fig. 5d**). Although CT appeared to enhance trimer thermostability, this double mutation introduced an additional unfolding intermediate.

**Fig. 5.**
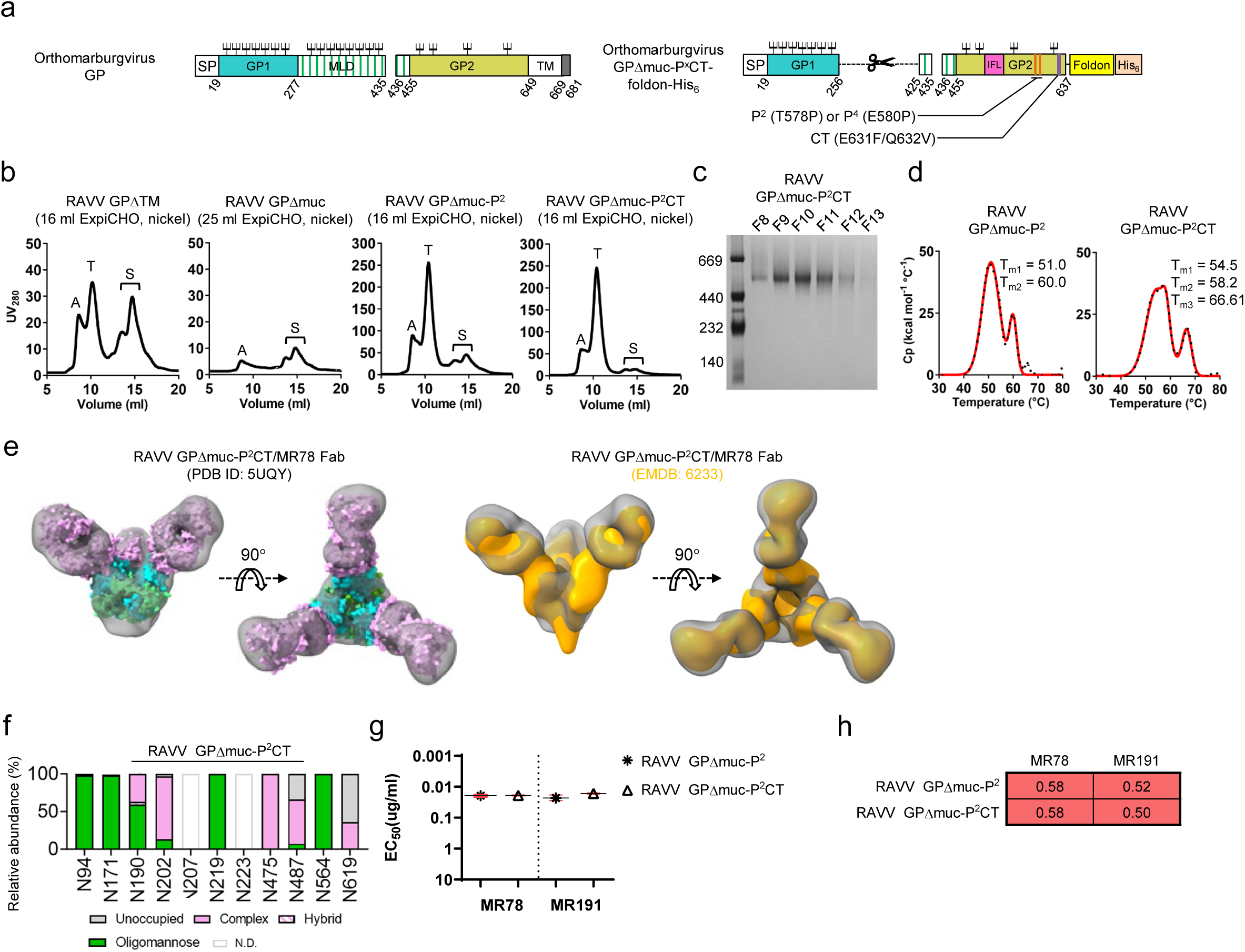
Rational design and in vitro characterization of orthomarburgvirus GP trimers. (**a**) Schematic of GP construct design based on the RAVV backbone, including MLD deletion (Δmuc, residues T256–T424), a P^2^ (T578P) or P^4^ (E580P) mutation in the HR1c bend, and a C-terminal double mutation (E631F/Q632V). (**b**) SEC profiles of four RAVV GP constructs expressed in 16 ml ExpiCHO cultures and purified using a nickel column. Aggregation (A), trimer (T), and subunit (S) peaks are indicated. (**c**) BN-PAGE analysis of RAVV GPΔmuc-P^2^CT trimer fractions purified by SEC. (**d**) DSC thermograms of nickel/SEC-purified RAVV GPΔmuc-P^2^ and GPΔmuc-P^2^CT trimers. Experimental data (black dots) and Gaussian fits (red lines) are shown, with melting temperatures (T_m1_, T_m2_, and T_m3_) labeled. (**e**) nsEM analysis of RAVV GPΔmuc-P^2^CT/MR78 Fab complex. Left: crystal structure (PDB ID: 5UQY) of RAVV GPΔmuc/MR78 Fab complex fitted into the nsEM density. Right: nsEM density map of RAVV GPΔmuc/MR78 Fab complex (EMDB: 6233, orange surface) overlaid with the nsEM map obtained in this study. (**f**) Site-specific glycan profile of RAVV GPΔmuc-P^2^CT trimer. Glycan types are colored as in Fig. 1e. (**g**) ELISA-derived EC_50_ (µg/ml) values for RAVV GPΔmuc-P^2^ and GPΔmuc-P^2^CT trimers binding to MARV NAbs MR78 and MR191. Absorbance <0.5 at 10 µg/ml was used as a non-binding cutoff, as in Fig. 1f. (**h**) BLI profiles of RAVV GPΔmuc-P^2^ and GPΔmuc-P^2^CT trimers binding to MARV NAbs MR78 and MR191. Sensorgrams were obtained using the same protocol as in Fig. 1h (see **Fig. S5h**). Peak responses are summarized in a matrix, as in Fig. 1h.

The lead design, RAVV GPΔmuc-P^2^CT, was complexed with MR78 for nsEM analysis (**Fig. 5e** and **Fig. S5e**). We hypothesized that if the nsEM model, despite its low resolution, matched the crystal structure (PDB ID: 5UQY) (*40*), it would validate our design and confirm presentation of a conserved NAb epitope. Indeed, 2D class averages revealed well-formed RAVV GPΔmuc/MR78 complexes, with the crystal structure fitting well into the nsEM density map (**Fig. 5e**, left). Notably, our model also aligned closely with a previously reported nsEM reconstruction of the same complex (*42*) (**Fig. 5e**, right). Site-specific glycan analysis revealed distinct patterns relative to orthoebolavirus GPs (**Fig. 5f** and **Fig. S5f**). Specifically, N-glycan sites N94, N171, N219, and N564 carried primarily oligomannose-type glycans. In contrast, sites N190, N202, and N487 presented either oligomannose- or complex-type glycans, while N475 and N619 were mostly occupied by complex-type glycans. In ELISA (**Fig. 5g** and **Fig. S5g**), RAVV GPΔmuc-P^2^ and GPΔmuc-P^2^CT trimers bound to MR78 and MR191 with similar EC_50_ values, consistent with the BLI data (**Fig. 5h** and **Fig. S5h**). RAVV GPΔmuc-P^2^CT was subsequently displayed on E2p and I3-01v9b SApNPs and analyzed by nsEM following ExpiCHO expression and IAC purification (**Fig. S5i**). However, well-formed protein particles were not observed in the micrographs. Taken together, these findings support the use of RAVV GPΔmuc-P^2^CT as a native-like, prefusion-stabilized GP construct for orthomarburgvirus vaccine development, although further optimization is needed for multivalent display on SApNPs.

### Distribution and retention of SUDV GPΔmuc trimers and SApNPs in lymph nodes

In our previous studies, we observed a positive correlation between the thermostability of a viral antigen (i.e., T_m_) displayed on E2p or I3-01v9 SApNPs and the retention time of the resulting SApNP vaccine antigens in lymph node follicles (*113, 127, 128*). Given the comparable T_m_ values (∼63-66 °C) of SUDV GPΔmuc and HIV-1 Env trimers, we anticipated that SUDV GPΔmuc-presenting SApNPs would exhibit similar in vivo behavior. Following our previously established protocol (*113, 127, 128*), we immunized mice with SUDV GPΔmuc-WL^2^P^4^ trimer and SApNPs (including both E2p and I3-01v9b) to investigate their interaction with cellular components in lymph nodes and analyze vaccine-induced immunological responses.

To induce a robust humoral response, vaccine antigens must be transported through the lymphatic system, accumulate in lymph node follicles, and effectively engage B cell receptors (BCRs) to stimulate B cell activation (*129–132*). We first examined the distribution of E2p and I3-01v9b SApNPs in lymph nodes. SApNPs were administered intradermally to mice via footpads (4 footpads, 10 μg/footpad). The brachial and popliteal sentinel lymph nodes were isolated from both sides of the body of the mouse 12 h after a single-dose injection for immunohistological studies. Four NAbs – ADI-15878 (*116*), ADI-15946 (*119*), CA45 (*118*), and mAb100 (*133*) – were used to stain lymph node sections for antigen detection (**Fig. S6a**). Overall, the immunostaining images from CA45 exhibited the best signal-to-noise ratio. Based on these data, CA45 was selected to study the distribution and trafficking of SUDV GPΔmuc vaccines in lymph nodes. Consistent with our previous studies of SARS-CoV-2 spike (*128*), HIV-1 Env (*113*), and influenza M2ex3 (*127*) SApNPs, SUDV GPΔmuc-presenting E2p and I3-01v9b SApNPs accumulated in the centers of lymph node follicles (**Figs. 6a** and **6b**, images on the left; schematics on the right).

**Fig. 6.**
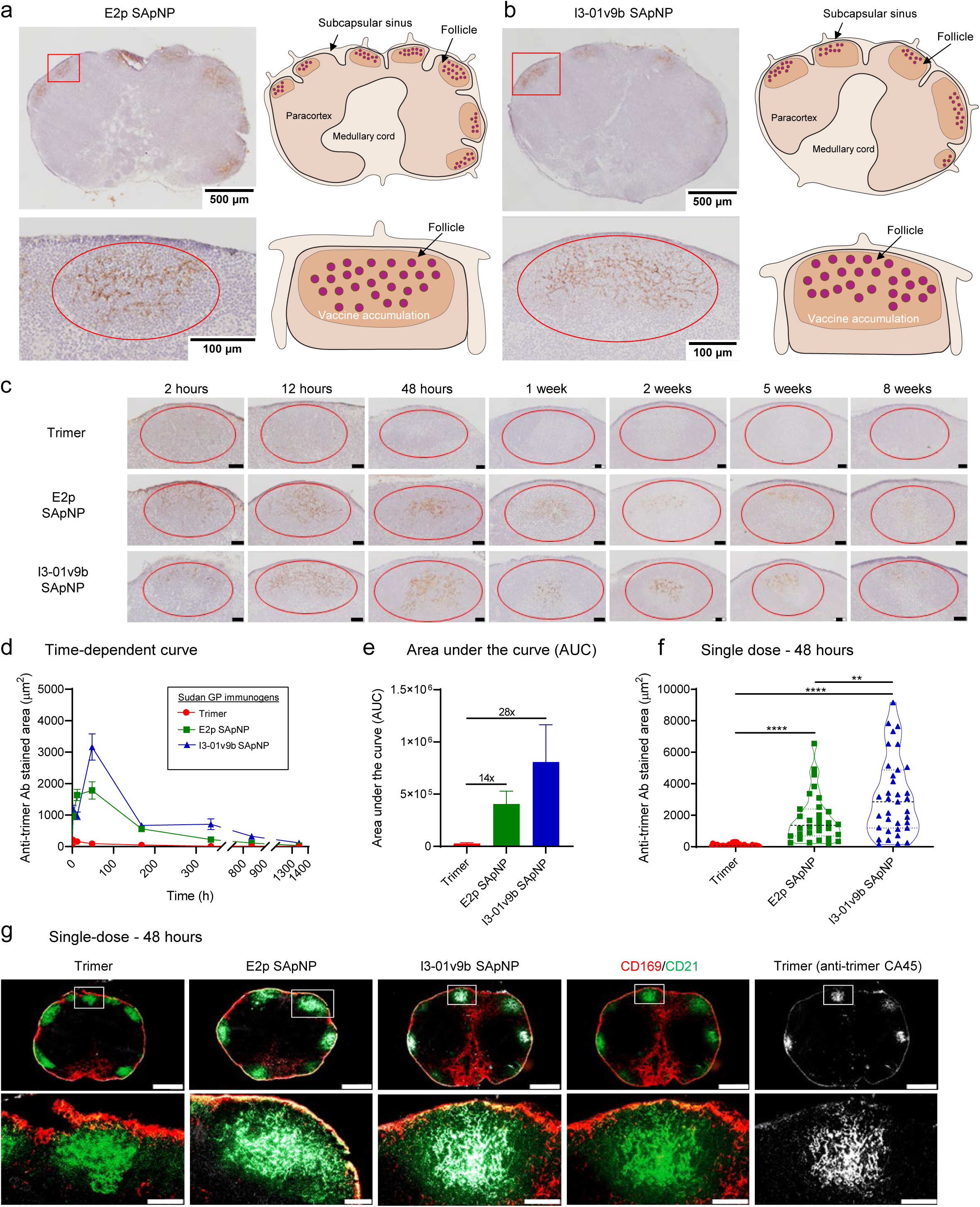
Retention of SUDV GPΔmuc-presenting SApNPs in lymph node follicles. (**a, b**) Left: distribution of (**a**) E2p and (**b**) I3-01v9b SApNPs displaying SUDV GPΔmuc-WL^2^P^4^ trimers in lymph nodes 12 h after a single-dose injection (10 μg/injection, 40 μg/mouse). NAb CA45 was used to stain the sentinel lymph node sections. Right: schematic of SUDV SApNP accumulation in lymph node follicles. (**c**) Trafficking and retention of the SUDV GPΔmuc trimer and SApNPs in lymph node follicles 2 h to 8 weeks after a single-dose injection. Scale bar: 50 μm. (**d**) Time-dependent curve and (**e**) area under the curve (AUC) of CA45-stained areas in immunohistological images of SUDV vaccine retention in lymph node follicles over 8 weeks. (**f**) Quantification of SUDV vaccine accumulation in lymph node follicles 48 h after a single-dose injection. Data were collected from > 10 lymph node follicles (n = 3-5 mice/group). (**g**) Interaction of the SUDV GPΔmuc trimer and SApNPs with FDC networks in lymph node follicles 48 h after a single-dose injection. Both E2p and I3-01v9b SApNPs colocalized with FDC networks. Immunofluorescent images are pseudo-color-coded (CD21^+^, green; CD169^+^, red; CA45, white). Scale bars: 500 (whole lymph node) and 100 μm (enlarged follicle). Data are presented as mean ± SEM in (d), and mean ± SD in (e, f). Statistical analysis was performed using one-way ANOVA followed by Tukey’s multiple comparison *post hoc* test. **p < 0.01, ****p < 0.0001.

We next assessed the trafficking and retention patterns of SUDV GPΔmuc-WL^2^P^4^ trimer and SApNPs in lymph node follicles over a period of 8 weeks following a single-dose injection (4 footpads, 10 μg/footpad) (**Fig. 6c**). Histological images showed that all SUDV immunogens were transported into lymph nodes and accumulated in the subcapsular sinus within 2 h (**Fig. 6c**). While the soluble trimer was transported into lymph node follicles within 2 h and completely cleared by 48 h, both E2p and I3-01v9b SApNPs appeared in follicles at 2 h, reached peak accumulation at 48 h, and were retained in follicles over the entire 8-week period (**Fig. 6c**). Next, the CA45-stained area was quantified in a time-dependent manner, revealing a ∼112-fold longer retention for E2p and I3-01v9b SApNPs compared with the soluble trimer (**Figs. 6c** and **6d**). The area under the curve (AUC) indicated that the exposure of SUDV GPΔmuc presented on the SApNP surface was 14-28 times higher than the same antigen in soluble trimer form (**Fig. 6e**). At 48 h, the two large SApNPs displaying 20 GPΔmuc trimers also showed 19-33 times greater accumulation compared with the soluble GPΔmuc trimer (**Fig. 6f**). These findings are consistent with our previous studies (*113, 127, 128*), in which individual antigens were cleared from follicles within 48 h, whereas large antigen-presenting SApNPs exhibited prolonged follicular retention lasting 2-8 weeks. Notably, the retention time of SUDV GPΔmuc SApNPs fell within the same range as HIV-1 Env and influenza M2ex3 SApNPs, all of which had T_m_ values of 62-66°C. Together, these data support the conclusion that stabilized antigens with high thermostability (T_m_ > 62°C) result in prolonged SApNP retention in lymph node follicles for at least 8 weeks.

FDCs are resident stromal cells that form a network structure in lymph node follicles and play an essential role in antigen retention and presentation to stimulate B cell responses (*129–131, 134*). FDC networks retain soluble antigens, immune complexes, viruses, and bacteria on their surfaces and dendrites to initiate and maintain GC responses (*135–137*). Our previous studies demonstrated that FDC networks are the primary follicular components responsible for retaining SARS-CoV-2 spike, HIV-1 Env, and influenza M2ex3 SApNPs (*113, 127, 128*). To test whether this was also true for SUDV GPΔmuc SApNPs, we isolated sentinel lymph nodes at the peak of antigen accumulation (48 h), as well as a series of timepoints (2 h to 8 weeks) after intradermal injection (**Fig. 6g** and **Figs. S6b-g**). Lymph node sections were stained with the NAb CA45 (white) (*118*) for SUDV GPΔmuc, anti-CD21 antibodies (green) for FDCs, and anti-CD169 antibodies (red) for subcapsular sinus macrophages. To confirm the identity of FDC networks, co-staining was performed using the more specific FDC marker, anti-FDC-M1. Immunofluorescence imaging showed that SApNPs colocalized with FDC networks labeled by both CD21 and FDC-M1 at 48 h post-injection (**Fig. 6g** and **Fig. S6h**), supporting the conclusion that FDC networks mediate the accumulation and retention of SUDV GPΔmuc-presenting SApNP vaccines.

### Interaction of SUDV GPΔmuc SApNPs with FDCs and phagocytic cells in lymph nodes

FDC networks form a reservoir in lymph node follicles for antigen sequestration, alignment, and presentation to effectively crosslink BCRs, activate naive B cells, and stimulate GC reactions (*129–131, 138*). Our previous TEM analysis showed that FDC networks retained SARS-CoV-2 spike and HIV-1 Env SApNPs on their surfaces and dendrites through interactions with complement protein 3 (C3) and complement receptor 2 (CR2) (*113, 128*). Here, we visualized the interface between FDC dendrites and B cells to understand how FDC networks present SUDV GPΔmuc SApNPs adjuvanted with aluminum phosphate (AP) to engage B cells. To this end, we injected adjuvanted E2p and I3-01v9b SApNPs into the mouse hind footpads (2 footpads, 50 μg/footpad). Fresh popliteal sentinel lymph nodes were isolated 2, 12, and 48 h after a single injection. The lymph node tissues were sectioned and processed for TEM analysis. TEM images revealed the expected, characteristic morphology of long FDC dendrites interacting with B cells in lymph node follicles (**Figs. 7a-c**). Intact SApNPs (round-shaped granules, yellow arrows) were aligned on FDC dendrites and B cell surfaces at all timepoints, and AP particles did not appear to colocalize with SApNPs (**Figs. 7a-c** and **Fig. S7a-f**). These findings demonstrate FDC networks collect and present vaccine antigens (i.e., SUDV GPΔmuc SApNPs) and use their long dendrites to maximize interactions between antigens and BCRs. At the later timepoints (12 and 48 h), SUDV GPΔmuc SApNPs were also found within endolysosomes of B cells (**Figs. S7c** and **e**).

**Fig. 7.**
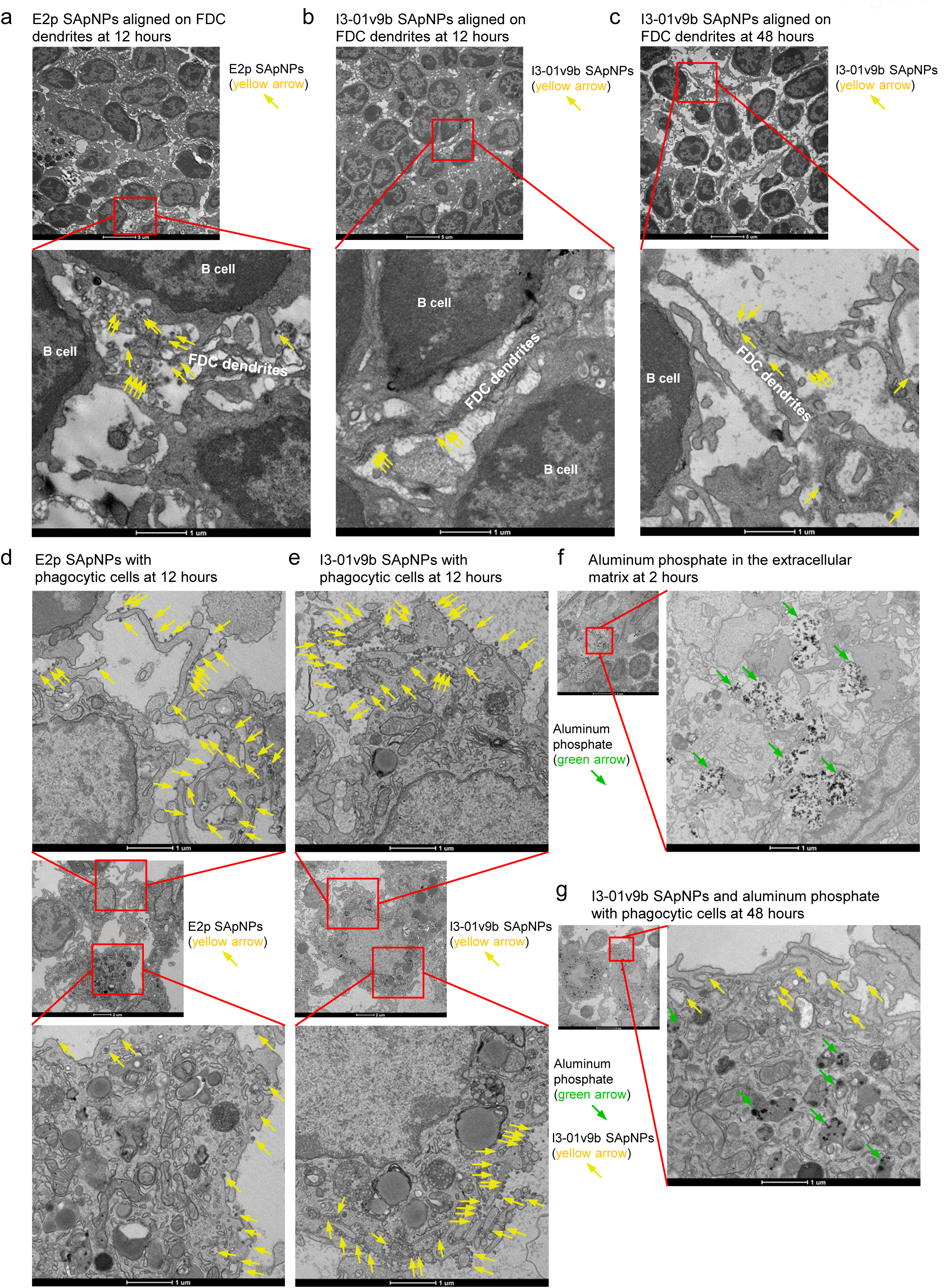
Interaction of SUDV GPΔmuc-presenting SApNPs with FDCs and phagocytic cells in lymph nodes. **(a-c)** TEM images of (**a**) E2p and (**b, c**) I3-01v9b SApNPs (yellow arrows) displaying SUDV GPΔmuc-WL^2^P^4^ trimers aligned on FDC dendrites in lymph node follicles at (**a, b**) 12 h and (**c**) 48 h after a single-dose injection. Each mouse received 200 μl of antigen/aluminum phosphate (AP) adjuvant mix (100 μg SApNP + 100 μl adjuvant), administered into both hind footpads (2 footpads, 50 μg/footpad). Fresh popliteal lymph nodes were collected. No AP adjuvant was observed on FDC dendrites. (**d, e**) TEM images of (**d**) E2p and (**e**) I3-01v9b SApNPs (yellow arrows) displaying SUDV GPΔmuc-WL^2^P^4^ trimers on the surface and within endolysosomes of phagocytic cells at 12 h post-injection. (**f**) TEM images showing AP adjuvant (green arrows) aggregated in the extracellular matrix (ECM) at 2 h post-injection. (**g**) TEM images of Sudan GPΔmuc-presenting I3-01v9b SApNPs and AP adjuvant localized inside endolysosomes of phagocytic cells at 48 h post-injection.

Phagocytic cell populations, such as macrophages and dendritic cells, take up and process vaccine antigens to facilitate adaptive immune responses (*132, 139, 140*). These innate immune cells, located in the subcapsular sinus and medullary sinus of lymph nodes, can capture vaccine antigens, transport them to migrating B cells, and eventually deposit them on FDCs through a complement-dependent transport mechanism (*130, 131, 141–145*). In this study, we assessed the interaction between phagocytic cells and AP-adjuvanted SApNPs. SUDV GPΔmuc SApNPs, both E2p and I3-01v9b, were found either on the surface or within endolysosomes of phagocytic cells (**Figs. 7d** and **e** and **Figs. S7g-l**), consistent with our previous findings for SARS-CoV-2 spike and HIV-1 Env SApNPs (*113, 128*). AP particles tended to aggregate in the extracellular matrix (ECM) 2 h after injection (**Fig. 7f** and **Fig. S7m**). Both I3-01v9b SApNPs and AP particles were also observed inside endolysosomes of phagocytic cells at all timepoints (**Fig. 7g** and **Figs. S7n-p**). Overall, E2p and I3-01v9b SApNPs exhibited similar patterns of interaction with FDC networks, B cells, and phagocytic cells. FDC networks retained SApNPs on their dendrites and presented native antigens to B cells in lymph node follicles. A substantial amount of SApNPs was also processed inside endolysosomes of phagocytic cells in the presence of AP adjuvant.

### Assessment of GC reactions induced by SUDV GPΔmuc trimers and SApNPs in lymph nodes

In lymph node follicles, GCs are the sites where B cell somatic hypermutation, selection, affinity maturation, and class switching occur (*135, 146, 147*). GC reactions lead to the formation of immune memory and the development of NAb responses following vaccination (*129, 132*). As the essential components supporting GC initiation and maintenance, FDC networks and T follicular helper (T_fh_) cells present vaccine antigens and stimulate B cells, respectively (*148–150*). Here, we hypothesized that SUDV GPΔmuc SApNPs, due to prolonged retention by FDC networks, would induce more robust and long-lived GC reactions in lymph node follicles compared with the soluble trimers. We first characterized GC reactions induced by I3-01v9b SApNPs after a single-dose injection (4 footpads, 10 μg/footpad). As previously described (*113, 127, 128*), vaccine-induced GC B cells (GL7^+^, red) and T_fh_ cells (CD4^+^ Bcl6^+^, co-labeled with cyan and red) in lymph nodes were assessed by immunohistological analysis. Robust GCs were formed and attached to FDC networks (CD21^+^, green), with well-organized dark zone (DZ) and light zone (LZ) compartments in B cell follicles (B220+, blue) (**Fig. 8a**, left). T_fh_ cells were observed in the LZ of GCs, sustaining B cell affinity maturation (**Fig. 8a**, right). We then applied this immunohistological assessment to the SUDV GPΔmuc trimer and SApNPs (E2p and I3-01v9b) at 2, 5, and 8 weeks after a single-dose injection (**Fig. 8b** and **Figs. S8a-c**), and at 2 and 5 weeks after the boost (**Fig. 8c** and **Figs. S8d** and **e**). GC reactions were quantified using two metrics: GC/FDC ratio (the frequency of GCs associated with FDC networks) and size of GCs (area occupied in follicles) as previously described (*113, 127, 128*). All three SUDV GPΔmuc immunogens induced robust GCs, with the I3-01v9b SApNP group showing the largest GCs at 2 weeks after a single-dose vaccination (**Fig. 8b** and **Fig. S8a**). While the GC/FDC ratio decreased significantly over 8 weeks following a single trimer dose, both SApNPs supported long-lived GCs with high GC/FDC ratios (e.g., > 50%) lasting for 8 weeks (**Fig. 8b**). GC sizes declined over time in both trimer and SApNP groups (**Figs. 8b** and **d**). After boosting, all vaccine groups showed restored GC formation, and the GC/FDC ratio in the GPΔmuc trimer group remained comparable (all > 75%) to those in the two SApNP groups (**Fig. 8c**). Overall, the two SApNPs generated larger GCs than the soluble trimer, 1.3 to 2.5 times larger after the prime (**Figs. 8b** and **d**) and 1.4 to 1.8 times larger after the boost (**Figs. 8c** and **e**).

**Fig. 8.**
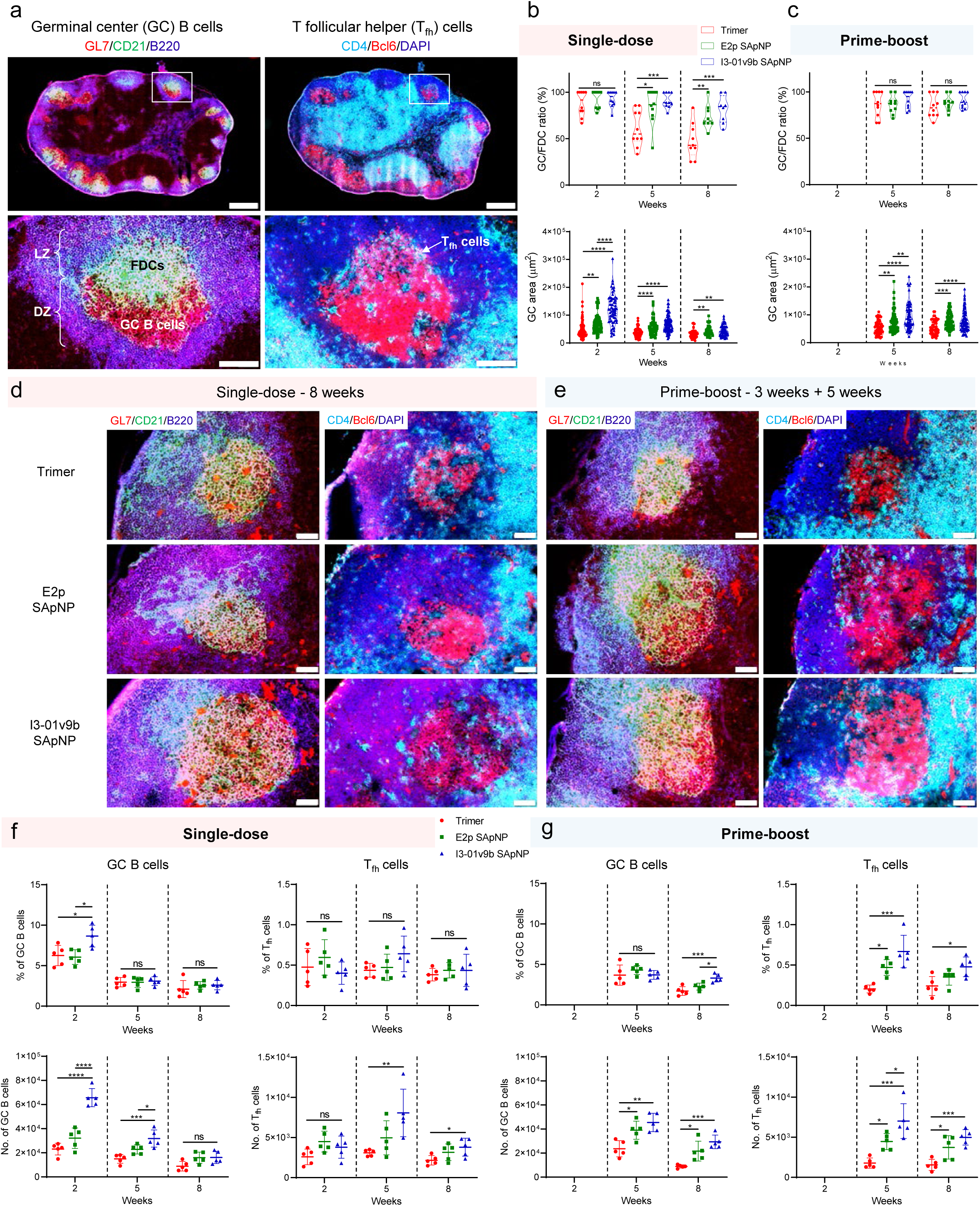
Induction of germinal center reactions by SUDV GPΔmuc-presenting SApNPs in lymph nodes. (**a**) Top: representative immunofluorescent images of germinal centers (GCs) induced by the SUDV GPΔmuc-presenting SApNP vaccine 2 weeks after a single-dose injection (10 μg/injection, 40 μg/mouse). Each dose consisted of 80 μl of antigen/aluminum phosphate (AP) adjuvant mix. Bottom: robust GC formation with well-organized light zone (LZ) and dark zone (DZ) compartments in lymph node follicles. GC B cells (GL7^+^, red) attached to FDCs (CD21^+^, green) and T_fh_ cells located in the LZ. Scale bars: 500 μm (whole lymph node), 100 μm (enlarged follicle). (**b**, **c**) Assessment of GC responses from immunofluorescent images at 2, 5, and 8 weeks after a single-dose injection, or 2 and 5 weeks after a boost administered 3 weeks post-prime (n = 5 mice/group). The GC/FDC ratio and GC size were measured and plotted. (**d**, **e**) Representative GC immunofluorescent images from lymph nodes collected at week 8 after single-dose or prime-boost immunization with SUDV GPΔmuc vaccines. Scale bar: 50 μm. (**f, g**) Quantification of GCs by flow cytometry after single-dose or prime-boost regimens (n = 5 mice/group). Frequency and number of GC B cells and T_fh_ cells were characterized and plotted. Data are shown as mean ± SD. Statistical analysis was performed using one-way ANOVA followed by Tukey’s multiple comparison *post hoc* test at each timepoint. ns (not significant), *p < 0.05, **p < 0.01, ***p < 0.001, ****p < 0.0001.

Next, we assessed GC reactions using flow cytometry. We intradermally immunized mice with three vaccines, and sentinel lymph nodes were isolated at 2, 5, and 8 weeks after a single-dose injection (**Fig. 8f** and **Fig. S9**) and 2 and 5 weeks after the boost (**Fig. 8g**) (four footpads, 10 μg/injection). Fresh lymph node tissues were disaggregated into a single-cell suspensions and stained with an antibody cocktail. GC reactions were quantified based on the percentage and number of GC B and T_fh_ cells (**Fig. S9**). The I3-01v9b SApNP group yielded the highest percentage and number of GC B cells at 2 weeks post-prime (**Fig. 8f**), consistent with the immunohistological findings. Among all vaccine groups, GC B cells declined over time, whereas T_fh_ cell levels remained stable over the 8-week period. A boost enhanced both the frequency and number of GC B and T_fh_ cells (**Fig. 8g**). Notably, GC B and T_fh_ cells induced by I3-01v9b SApNP persisted at relatively high levels over 8 weeks post-boost. Compared with the soluble trimer, the E2p and I3-01v9b SApNPs elicited 1.8/2.4-3.3 times more GC B cells, and 1.5-1.8/2.4-3.2 times more T_fh_ cells, at 8 weeks after the single-dose/boost injection, respectively (**Figs. 8f** and **g**). In summary, the two SUDV GPΔmuc SApNPs generated more robust and long-lived GC reactions than the soluble GPΔmuc trimer, leading to more potent and durable humoral immune responses.

### Antibody responses induced by rationally designed filovirus GPΔmuc vaccines in mice

We assessed the immunogenicity of rationally designed filovirus GPΔmuc constructs, including trimers and SApNPs (E2p and I3-01v9b), in mice (**Fig. 9a**). The filovirus antigen (10 µg) was adjuvanted with aluminum hydroxide (AH) and a Toll-like receptor 9 (TLR9) agonist, CpG ODN 1826. Mice were immunized intraperitoneally at weeks 0, 3, 9, and 15 with 6-week intervals between the second and third doses and between the third and fourth doses. Serum was collected two weeks after each dose for serological analysis. EBOV vaccine-induced binding antibody (bAb) responses were measured by ELISA using an EBOV GPΔmuc-WL^2^P^4^(1TD0) antigen (**Fig. 9b** and **Figs. S10a**, **b**). The use of 1TD0, rather than foldon, as the trimerization motif in the coating antigen enabled the detection of GPΔmuc-specific bAb responses. Overall, the EBOV GPΔmuc E2p SApNP elicited the highest bAb titers at all four timepoints, with EC_50_ values 8.2- and 5.3- fold higher than those of the soluble trimer at weeks 2 and 5, respectively. All vaccine groups reached plateau EC_50_ titers by week 11, after three immunizations. The EBOV GPΔmuc I3-01v9b SApNP achieved an EC_50_ titer of 182,253 at week 11, identical to its E2p counterpart and 2.5-fold higher than that elicited by the soluble trimer. Week-17 sera (after four doses) were tested against a SUDV GPΔmuc-WL^2^P^4^(1TD0) antigen (**Fig. 9c** and **Fig. S10c**). The EBOV GPΔmuc E2p SApNP again elicited the highest EC_50_ titer (32,585), 3.5-fold higher than that of the soluble trimer. Cross-reactive bAb responses were also evaluated against a RAVV GPΔmuc-P^2^CT(1TD0) antigen (**Fig. 9d** and **Fig. S10d**). Both E2p and I3-01v9b SApNP groups showed significantly higher serum binding signals, indicated by absorbance at 450 nm (A450), compared with the soluble trimer group. NAb responses were evaluated using a filovirus pseudoparticle (pp) neutralization assay (*36*), with 50% inhibitory dilution (ID_50_) values calculated for comparison. Week-2 sera served as negative controls for longitudinal analysis. None of the EBOV GPΔmuc vaccines elicited NAb responses at week 2, but ID_50_ titers were detected at weeks 5, 11, and 17, increasing steadily against EBOV-Makona pseudovirus (**Fig. 9e** and **Figs. S10e** and **f**). Consistent with the bAb results, EBOV GPΔmuc E2p SApNP produced the highest NAb titers at all timepoints, although no statistical significance was observed across groups. Week-17 sera were also evaluated against the SUDV-Gulu pseudovirus (**Fig. 9f** and **Fig. S10g**). The EBOV GPΔmuc E2p SApNP yielded the highest ID_50_ titer (761) among all groups. Notably, the EBOV GPΔmuc trimer group elicited the highest ID_50_ titer (2,759) at week 17 against a BDBV-Uganda pseudovirus, 4.5- and 4.9-fold higher than those elicited by the E2p and I3-01v9b SApNPs, respectively (**Fig. 9g** and **Fig. S10h**). Thus, while soluble trimers can achieve high NAb titers, they may require more doses than SApNPs.

**Fig. 9.**
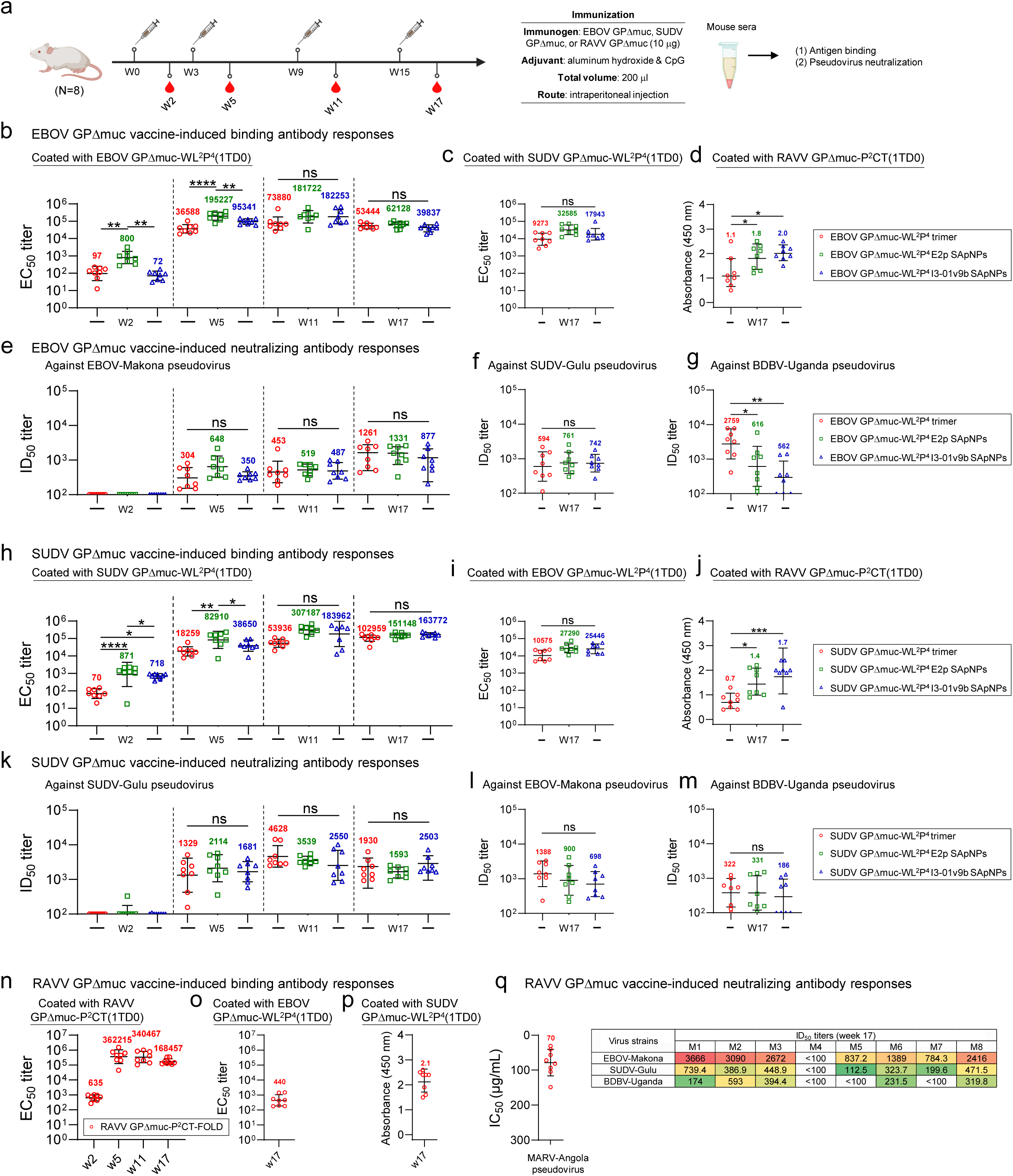
Antibody responses to rationally designed filovirus vaccines in mice. (a) Schematic of the mouse immunization regimen for in vivo evaluation of EBOV, SUDV, and RAVV GPΔmuc vaccines (n = 8 mice/group). Each dose consisted of 200 μl of antigen/AH + CpG adjuvant mix containing 10 μg of immunogen and 100 μl of adjuvant. Mice were immunized at weeks 0, 3, 9, and 15 via the intraperitoneal (i.p.) route, with 6-week intervals between the second and third, and third and fourth doses. (**b-d**) EBOV vaccine-induced binding antibody (bAb) responses against (**b**) EBOV GPΔmuc-WL^2^P^4^(1TD0), (**c**) SUDV GPΔmuc-WL^2^P^4^(1TD0), and (**d**) RAVV GPΔmuc-P^2^CT(1TD0). (**e-g**) EBOV vaccine-induced NAb responses against (**e**) EBOV Makona, (**f**) SUDV Gulu, and (**g**) BDBV Uganda pseudoviruses. (**h-j**) SUDV vaccine-induced bAb responses against (**h**) SUDV GPΔmuc-WL^2^P^4^(1TD0), (**i**) EBOV GPΔmuc-WL^2^P^4^(1TD0), and (**j**) RAVV GPΔmuc-P^2^CT(1TD0). (**k-m**) SUDV vaccine-induced NAb responses against (**k**) SUDV Gulu, (**l**) EBOV Makona, and (**m**) BDBV Uganda pseudoviruses. (**n-p**) RAVV vaccine-induced bAb responses against (**n**) RAVV GPΔmuc-P^2^CT(1TD0), (**o**) EBOV GPΔmuc-WL^2^P^4^(1TD0), and (**p**) SUDV GPΔmuc-WL^2^P^4^(1TD0). (**q**) RAVV vaccine-induced NAb responses against MARV Angola pseudovirus (using purified IgG) and pseudotyped orthoebolaviruses (using serum). EC_50_ titers were derived from serum ELISA against coating antigens, and ID_50_/IC_50_ titers were determined from pseudovirus neutralization assays. Geometric means are labeled on the plots. ID_50_/IC_50_ values were calculated using a % neutralization range of 0.0–100.0%. Color coding indicates ID_50_ magnitude (green to red: low to high neutralization). Data are presented as the mean ± SD. Statistical analysis was performed using one-way ANOVA followed by Tukey’s multiple comparison *post hoc* test at each timepoint. ns (not significant), \**p* < 0.05, \*\**p* < 0.01, \*\*\**p* < 0.001, and \*\*\*\**p* < 0.0001. The schematic in (a) was created with BioRender.com.

We assessed SUDV vaccine-induced serum bAb responses by ELISA against a SUDV GPΔmuc-WL^2^P^4^(1TD0) antigen (**Fig. 9h** and **Figs. S10i**, **j**). Consistent with the bAb responses elicited by EBOV vaccines, SUDV GPΔmuc E2p SApNP generated the highest EC_50_ titers of 871, 82,910, and 307,187 at weeks 2, 5 and 11, respectively, 12.4-, 4.5-, and 5.7-fold higher than those induced by the soluble trimer. EC_50_ titers in both E2p and I3-01v9 SApNP groups plateaued after three doses by week 11, whereas the soluble trimer group reached a peak of 102,959 after four doses at week 17, which remained slightly lower than the titers induced by SApNPs. These results correlated well with the GC reactions generated by SUDV GPΔmuc trimer and SApNPs (**Fig. 8**). Week-17 sera were then evaluated against an EBOV GPΔmuc-WL^2^P^4^(1TD0) antigen (**Fig. 9i** and **Fig. S10k**). The SUDV GPΔmuc E2p and I3-01v9b SApNPs elicited EC_50_ titers of 27,290 and 25,446, respectively, which were 2.6- and 2.4-fold higher than that of the soluble GPΔmuc trimer. Cross-reactive bAb responses were further tested using week-17 sera against a RAVV GPΔmuc-P^2^CT(1TD0) antigen (**Fig. 9j** and **Fig. S10l**). Both E2p and I3-01v9b SApNP groups exhibited significantly higher serum binding signals (A_450_) compared with the soluble trimer group. Next, we evaluated SUDV GPΔmuc vaccine-induced autologous NAb responses using a SUDV-Gulu pseudovirus assay. Week-2 sera were used as a negative control for the longitudinal analysis. At this timepoint, none of the vaccines elicited detectable NAb responses, except for one mouse in the E2p group. ID_50_ titers became detectable at week 5, plateaued at week 11 (after three doses), and remained stable through week 17 (**Fig. 9k** and **Figs. S10m**, **n**). Overall, the SUDV GPΔmuc trimer and SApNP vaccines generated comparable ID_50_ titers (1,329–4,628) at each timepoint after two doses. Finally, week-17 sera were assessed for cross-neutralizing activity against EBOV-Makona (**Fig. 9l** and **Fig. S10o**) and BDBV-Uganda (**Fig. 9m** and **Fig. S10p**) pseudoviruses. No statistically significant differences in cross-NAb responses were observed between the SUDV trimer and SApNP groups, as measured by ID_50_ titers.

Lastly, we assessed RAVV GPΔmuc trimer-induced serum bAb responses in ELISA using a RAVV GPΔmuc-P^2^CT(1TD0) antigen (**Fig. 9n** and **Fig. S10q**). Similar to EBOV and SUDV GPΔmuc trimers, RAVV GPΔmuc trimer generated a strong bAb response after two doses, with the peak EC_50_ titer of 36,2215 at week 5. Cross-reactive bAb responses were evaluated using week-17 sera against two orthoebolavirus antigens: EBOV GPΔmuc-WL^2^P^4^(1TD0) (**Fig. 9o** and **Fig. S10r**) and SUDV GPΔmuc-WL^2^P^4^(1TD0) (**Fig. 9p** and **Fig. S10s**). Low bAb titers, indicated by A_450_ values, were observed for SUDV GPΔmuc in most mice. Vaccine-induced NAb responses were evaluated against pseudotyped MARV-Angola and orthoebolaviruses using purified IgG and week-17 sera, respectively (**Fig. 9q** and **Figs. S10t**, **u**). Notably, IgG purification was required to eliminate nonspecific serum background in the MARV pseudovirus assay (**Fig. S10t**, left), and neutralization potency was measured by the 50% inhibitory concentration (IC_50_). The RAVV GPΔmuc trimer group yielded IC_50_ values of 30.9-148.4 μg/ml, with a geometric mean of 70.2 μg/ml, when IgG was tested against MARV-Angola (**Fig.9q**, left). RAVV GPΔmuc trimer-induced sera were then assessed against EBOV-Makona, SUDV-Gulu, and BDBV-Uganda pseudoviruses (**Fig. 9q**, right). Interestingly, cross-genus neutralization activity was demonstrated by detectable ID_50_ titers, although confirmation by IgG-based assays will be needed.

### Antibody responses induced by glycan-modified filovirus GPΔmuc vaccines in mice

We characterized the immunogenicity of filovirus GPΔmuc vaccines bearing oligomannose-rich (Kif) and trimmed (Kif/Endo H) glycans in mice (**Fig. 10a**). The same immunization protocol used in our previous vaccine studies (*100, 113, 151, 152*) was applied here, which differed from the longer-interval (6-week) regimen used for wild-type GPΔmuc vaccines (**Fig. 9a**). Mice were immunized intraperitoneally at weeks 0, 3, 6, and 9, with 3-week intervals between doses. Each injection contained 10 μg of glycan-modified antigen formulated with 100 μl of CpG/AH adjuvant, resulting in a final volume of 200 μl. Serum was collected two weeks after each vaccination to enable longitudinal comparison of bAb and NAb responses.

**Fig. 10.**
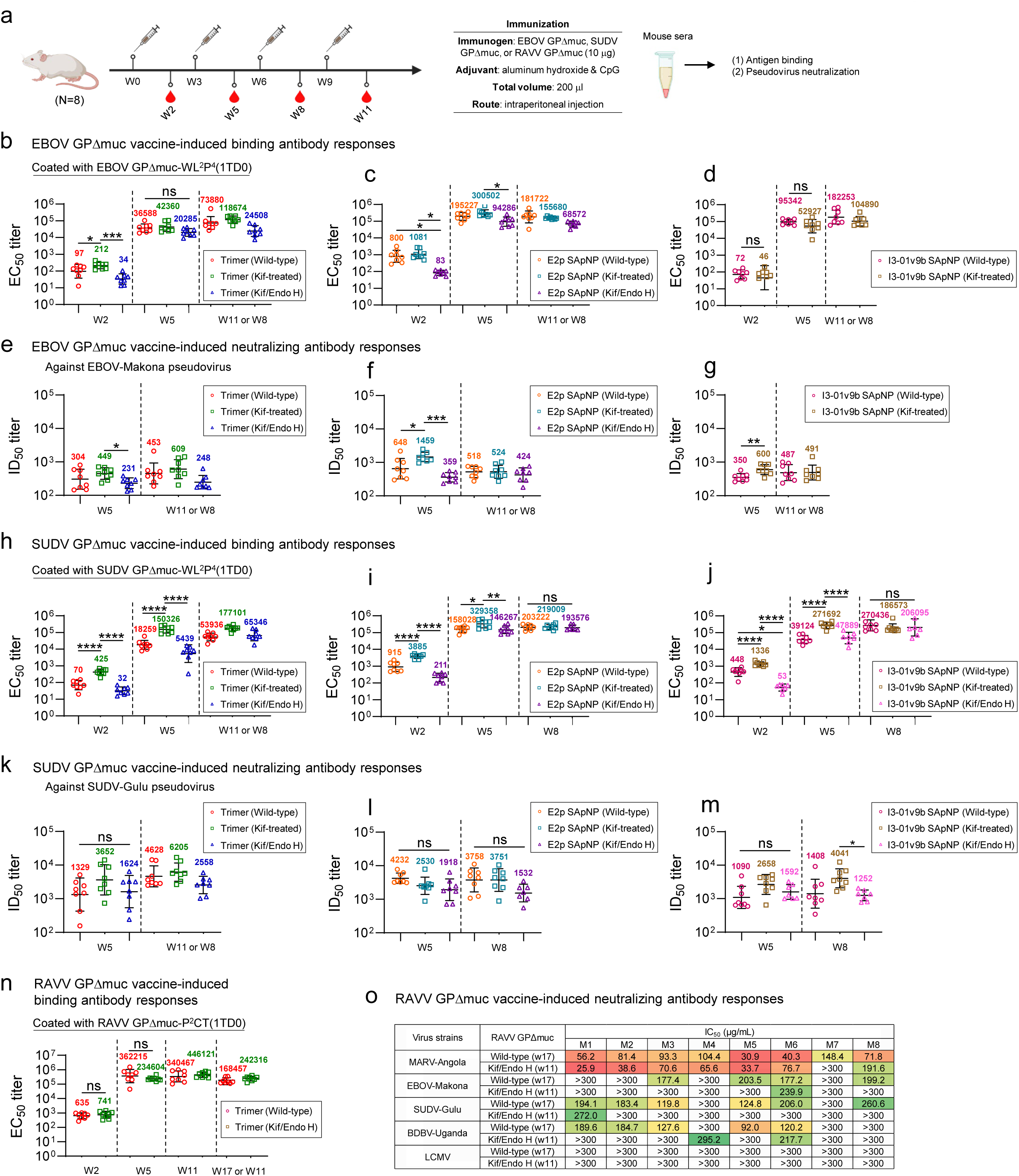
Effect of glycan modification on antibody responses elicited by filovirus vaccines. Schematic of the mouse immunization regimen for in vivo evaluation of glycan-modified EBOV, SUDV, and RAVV vaccines (*n* = 8 mice/group). Each dose consisted of 200 μl of antigen/AH + CpG adjuvant mix containing 10 μg of glycan-modified immunogen and 100 μl of adjuvant. Mice were immunized at weeks 0, 3, 6, and 9 via the intraperitoneal (i.p.) route, with 3-week intervals between doses. Vaccine immunogenicity was compared across three conditions: wild-type, Kif-treated (oligomannose-enriched), and Kif/Endo H-treated (a monolayer of GlcNAc stumps). (**b**-**d**) Glycan-modified EBOV vaccine-induced bAb responses against EBOV GPΔmuc-WL^2^P^4^(1TD0). (**e-g**) Glycan-modified EBOV vaccine-induced NAb responses against EBOV Makona pseudovirus. (**h-j**) Glycan-modified SUDV vaccine-induced bAb responses against SUDV GPΔmuc-WL^2^P^4^(1TD0). (**k-m**) Glycan-modified SUDV vaccine-induced NAb responses against Gulu pseudovirus. (**n**) Glycan-trimmed RAVV vaccine-induced bAb responses against RAVV GPΔmuc-P^2^CT(1TD0). (**o**) Glycan-trimmed RAVV vaccine-induced NAb responses against MARV Angola and pseudotyped orthoebolaviruses using purified IgG. EC_50_ titers were derived from serum ELISA against coating antigens, and ID_50_/IC_50_ titers were determined from the pseudovirus neutralization assays. Geometric means are labeled on the plots. ID_50_/IC_50_ values were calculated using a % neutralization range of 0.0–100.0%. Color coding indicates IC50 magnitude (green to red: low to high neutralization). Data for wild-type EBOV GPΔmuc trimer and SApNPs, SUDV GPΔmuc trimer, and RAVV GPΔmuc trimer from the long-interval immunization study were included for comparison. Boost timepoints (W11 and W17) were labeled but were excluded from statistical analysis. Data are presented as the mean ± SD. Statistical analysis was performed using one-way ANOVA, followed by Tukey’s multiple comparison *post hoc* test at each timepoint. Two-tailed unpaired t-tests were used to compare geometric means between two groups. ns (not significant), \**p* < 0.05, \*\**p* < 0.01, \*\*\**p* < 0.001, and \*\*\*\**p* < 0.0001. The schematic in (a) was created with BioRender.com.

We first measured serum bAb responses induced by glycan-modified EBOV GPΔmuc trimers by ELISA using an EBOV GPΔmuc-WL^2^P^4^(1TD0) antigen (**Fig. 10b** and **Figs. S11a, b**). Among the three groups, the glycan-trimmed trimer elicited the lowest EC_50_ titers at all timepoints, while the Kif-treated trimer generated the highest EC_50_ titers, 212 and 42,360 at weeks 2 and 5, respectively. In terms of EC_50_ titers, the Kif-treated group was 2.2- and 6.2-fold higher at week 2, and 1.2- and 2.1-fold higher at week 5, compared with the wild-type and glycan-trimmed groups, respectively. All trimer groups reached peak EC_50_ titers after three doses. Following a similar trend, the Kif-treated EBOV GPΔmuc E2p SApNP group showed the highest bAb titers at all three timepoints (**Fig. 10c** and **Figs. S11a**, **b**). Responses appeared to plateau at week 5, with the Kif-treated E2p SApNP yielding an EC_50_ titer of 300,502, 1.5- and 3.2-fold higher than its wild-type and glycan-trimmed counterparts, respectively. Wild-type and Kif-treated EBOV GPΔmuc I3- 01v9 SApNPs induced comparable bAb responses, both reaching peak EC_50_ titers after three doses (**Fig. 10d** and **Figs. S11a, b**). We next evaluated NAb responses induced by glycan-modified EBOV GPΔmuc vaccines against EBOV-Makona. Overall, the Kif-treated vaccines were the most immunogenic, while the glycan-trimmed vaccines elicited the lowest NAb titers at all three timepoints (**Fig. 10e-g** and **Figs. S11c, d**). Specifically, the Kif-treated EBOV GPΔmuc trimer yielded an ID_50_ titer of 449 at week 5, 1.5- and 1.9-fold higher than its wild-type and glycan-trimmed counterparts, respectively (**Fig. 10e**). The Kif-treated EBOV GPΔmuc E2p SApNP elicited an ID_50_ titer of 1,459 at week 5, 2.3- and 4.1-fold higher than those of the wild-type and glycan-trimmed groups, respectively (**Fig. 10f**). A similar pattern was observed in the I3-01v9b SApNP groups (**Fig. 10g**). These results demonstrated the beneficial effect of an oligomannose-rich glycan shield on bAb and NAb responses induced by EBOV GPΔmuc vaccines, in marked contrast to the glycan effects observed in our recent HIV-1 Env vaccine study (*113*).

We then used a SUDV GPΔmuc-WL^2^P^4^(1TD0) antigen in ELISA to assess serum bAb responses induced by glycan-modified SUDV GPΔmuc trimers (**Fig. 10h** and **Figs. S11e, f**). At all three timepoints, the Kif-treated SUDV GPΔmuc trimer elicited the highest bAb titers, whereas the glycan-trimmed trimer elicited the lowest, consistent with results observed for EBOV. Regardless of glycan modification, all vaccine groups reached peak EC_50_ titers after three doses. The Kif-treated SUDV GPΔmuc trimer yielded EC_50_ titers of 425 and 150,326 at weeks 2 and 5, respectively. In terms of EC_50_ titers, this trimer achieved 6.1- and 13.3-fold increases at week 2, and 8.2- and 27.6-fold increases at week 5, compared with the wild-type and glycan-trimmed counterparts, respectively. Similarly, for both SUDV GPΔmuc E2p and I3-01v9b SApNPs, the Kif-treated groups showed the highest bAb responses at weeks 2 and 5 (**Figs. 10i, j** and **Figs. S11e, f**). For both SApNPs, Kif treatment led to plateaued EC_50_ titers after two doses (week 5), whereas the wild-type and glycan-trimmed groups required three doses to reach their EC_50_ titers at week 8. The Kif-treated SUDV GPΔmuc E2p and I3-01v9b SApNPs elicited EC_50_ titers of 329,358 and 271,692 at week 5, respectively, 2.1- and 6.9-fold higher than their wild-type counterparts and 2.3-and 5.7-fold higher than their glycan-trimmed counterparts, respectively. We then evaluated NAb responses induced by glycan-modified SUDV GPΔmuc vaccines against SUDV-Gulu. All three SUDV GPΔmuc trimer groups reached peak ID_50_ titers after three doses (**Fig. 10k** and **Figs. S11g, h**). The Kif-treated SUDV GPΔmuc trimer group showed the highest ID_50_ titer of 3,652 at week 5, 2.7- and 2.2-fold higher than those of the wild-type and glycan-trimmed groups, respectively (**Fig. 10k**). While all three E2p SApNPs exhibited comparable ID_50_ titers (**Fig. 10l**), the Kif-treated I3- 01v9 SApNP elicited the highest ID_50_ titer of 4,041 at week 8, 2.9- and 3.2-fold higher than its wild-type and glycan-trimmed counterparts, respectively (**Fig. 10m**). Therefore, oligomannose enrichment conferred a similar advantage to SUDV as to EBOV vaccines (**Figs. 10e–g**).

We assessed serum bAb responses induced by the wild-type and glycan-trimmed RAVV GPΔmuc trimers by ELISA using a wild-type RAVV GPΔmuc-P^2^CT(1TD0) antigen (**Fig. 10n** and **Figs. S11i**, **j**). The wild-type group reached a peak EC_50_ titer of 362,215 after two doses, whereas the glycan-trimmed group required three doses to plateau at 446,121. Overall, no significant differences were observed between the two groups at any timepoint. We also compared NAb responses by testing purified IgG against pseudotyped MARV-Angola and orthoebolaviruses (**Fig. 10o** and **Fig. S10k**). Against MARV-Angola, both groups yielded similar IC_50_ values, ranging from 25.9 to 191.6 μg/ml. Cross-orthoebolavirus neutralization was evaluated using mouse IgG against EBOV-Makona, SUDV-Gulu, and BDBV-Uganda. The wild-type trimer induced stronger cross-genus activity than its glycan-trimmed counterpart, although the longer dosing interval may have contributed to this difference. Neither group showed nonspecific IgG neutralization against lymphocytic choriomeningitis virus pseudoparticles (LCMV-pps), included as a negative control. These findings were largely consistent with serum neutralization data (**Fig. 9q**).

### Sex-dependent antibody responses induced by stabilized SUDV GPΔmuc trimers

We studied the impact of sex on NAb responses induced by wild-type and Kif-treated SUDV GPΔmuc trimers. Briefly, mice were injected at weeks 0, 3, 6, and 9 via intradermal injection into four footpads, with 3-week intervals between doses (**Fig. S12a**). Each administered dose consisted of 80 μl of antigen/AH adjuvant mix containing 10 μg of trimer antigen and 40 μl of adjuvant. Serum samples collected at week 11, after four injections, were assessed against SUDV Gulu pseudovirus. For the wild-type SUDV GPΔmuc trimer, a significant sex-specific difference was observed: the ID_50_ titer in female mice (607) was 2.3-fold higher than in male mice (**Fig. S12b**). For the Kif-treated SUDV GPΔmuc trimer, female mice developed a slightly higher ID_50_ titer (593) than males, although the difference was not statistically significant (**Fig. S12c**). In both vaccine groups, female mice mounted greater NAb responses than males. This result is consistent with our previous study on RSV F trimer vaccines (*153*) and with broader consensus that females tend to generate stronger humoral immune responses than males following vaccination (*154, 155*).

Our detailed analysis of vaccine-induced antibody responses underscores the critical role of multivalent display and glycan modification in enhancing filovirus vaccine immunogenicity. Overall, Kif-treated GPΔmuc SApNPs elicited the strongest bAb and NAb responses. However, due to differences in immunization protocols (e.g., dosing intervals), additional in vivo studies are needed to confirm these results. Another key observation is the induction of cross-orthoebolavirus NAbs by the RAVV GPΔmuc-P^2^CT trimer, detected at both serum and IgG levels (**Fig. 9q**, right; **Fig. 10o**). This finding suggests that this GP possesses intrinsic immunogenic potential, warranting further validation in animal models, and could be incorporated into a pan-filovirus vaccine. Finally, sex is an important variable in evaluating filovirus vaccine responses.

## DISCUSSION

Filoviruses are a major cause of lethal VHF in humans and NHPs (*5, 6*). The viral GP mediates host cell entry by facilitating attachment and membrane fusion (*14*). Structural studies of filovirus GPs bound to NAbs from human survivors have shown that GP harbors neutralizing epitopes and is the primary target for vaccine development (*18–21*). Two vector-based EBOV vaccines, rVSV-ZEBOV (ERVEBO) (*63*) and a two-dose regimen (Zabdeno + Mvabea) (*64*), were approved by the U.S. FDA and the European Commission in 2019 and 2020, respectively. However, no licensed vaccines are available to prevent VHF caused by other filoviruses. Over the past decade, structure-based rational design has emerged as a powerful approach to accelerate vaccine development (*84, 85, 87, 88*). Like other class I viral fusion proteins, filovirus GP is metastable (*99, 100*), and vaccines expressing wild-type GP may not elicit optimal immune responses. Two recent studies identified molecular features contributing to GP metastability and stabilized the GPΔmuc trimer in a native-like prefusion conformation (*99, 100*). While native-like GP trimers have remained the focus of filovirus vaccine design, engineered protein NPs have emerged as a promising platform to enhance antigen presentation and immunogenicity (*74, 100*). In parallel, glycan trimming has been shown to improve NAb responses to vaccines targeting influenza, HIV-1, and SARS-CoV-2 (*112–114*), suggesting a broadly applicable strategy for viral GP-based immunogens. However, the combined effects of GP stabilization, NP display, and glycan modification have not been evaluated systematically across diverse filoviruses in the context of rational vaccine design.

We previously reported that the HR2 stalk and HR1c bend in GP2 contribute to EBOV GP metastability, and that a W615L mutation combined with a C-terminal extension (targeting HR2), along with a proline substitution at T577 or L579 (targeting HR1c), yielded a stable GPΔmuc trimer with high yield and homogeneity in mammalian cells (*100*). In this study, we extended this rational GP design strategy to SUDV and BDBV of the *Orthoebolavirus* genus and to RAVV of the *Orthomarburgvirus* genus to support vaccine development against future filovirus outbreaks – a priority of the U.S. Biomedical Advanced Research and Development Authority (BARDA) (*76, 156, 157*). Although the same GP2 elements (HR1c and HR2 stalk) appear to drive metastability across both filovirus genera, the stabilization strategies differ. For EBOV, SUDV, and BDBV, comparative analysis of GPΔmuc-WL^2^P^2^ and -WL^2^P^4^ constructs revealed that the P^4^ mutation conferred a more favorable thermostability profile, while trimer yield, purity, and antigenicity remained comparable between P^2^ and P^4^. The crystal structure of EBOV GPΔmuc-WL^2^P^4^ revealed a stabilizing interprotomer hydrogen bond (R130-A575), while the cryo-EM structure of SUDV GPΔmuc-WL^2^P^4^ resolved atomic details of the GP-CA45 interface, together confirming GPΔmuc-WL^2^P^4^ as a broadly applicable design for stabilizing orthoebolavirus GP in a native-like prefusion conformation. In contrast, for RAVV, the P^2^ (T578P) construct yielded substantially more trimer than the P^4^ (E580P) variant. A double mutation at the base of the HR2 stalk (E631F/Q632V) further improved trimer stability, resulting in the GPΔmuc-P^2^CT design. Notably, the W615L mutation used to stabilize orthoebolavirus GPs was originally derived from MARV (*100*), whereas the E631F/Q632V mutation used here to stabilize RAVV GP was introduced from EBOV. This reciprocal exchange suggests that filoviruses may have evolved distinct mechanisms for regulating GP metastability, via a widened neck in orthoebolaviruses or a charged tail in orthomarburgviruses, within the HR2 stalk. These findings suggest that triggers of metastability may also reside in the stalk regions of other class I viral fusion proteins, such as RSV F and SARS-CoV-2 spike.

We previously focused our design efforts on the NP interior, incorporating an inner shell of locking domains (LDs) and a core of helper T-cell epitopes (*100*). The resulting multilayered E2p and I3-01v9 SApNPs were used to present antigens from EBOV, SARS-CoV-2, HIV-1, and influenza (*100, 113, 127, 151, 158*). We also observed that EBOV GP and RSV F (*100, 153*), both containing a C-terminal coiled-coil stalk, dissociated into monomers in the absence of a C-terminal trimerization motif. This suggests that SApNPs with small, rigid anchoring sites may be better suited for displaying trimeric antigens with extended stalks. Compared with E2p, which features a compact triangular anchoring site (∼9 Å) on the NP surface (*159*), I3-01v9 presents a wider spacing of 50.5 Å between N-termini (*160*), which may destabilize filovirus GPs when displayed on the NP surface. To address this, we redesigned I3-01v9 using a previously developed I3-01v9a variant (*127*) as a template. An N-terminal helix was modeled and packed against the protein core in silico, yielding I3-01v9b and I3-01v9c. When the I3-01v9b/c-T variants were used as scaffolds to present EBOV GPΔmuc-WL^2^P^4^, the resulting constructs showed native-like GP trimer structures by EM. Notably, while all three orthoebolavirus GPΔmuc-WL^2^P^4^ trimers were successfully displayed on E2p and I3-01v9b SApNPs, a similar attempt with RAVV GPΔmuc-P^2^CT resulted in low yield and aggregation. It is plausible that specific regions within RAVV GP may interact with the NP backbone and cause misfolding. Nonetheless, our findings suggest that I3-01v9b/c can be used to display filovirus GPs, as well as other class I viral GPs such as RSV F and SARS-CoV-2 spike, with improved trimer stability. Glycan modification adds another dimension to rational vaccine design (*113*), but its biochemical, biophysical, structural, and antigenic effects must be carefully assessed. Although glycan-trimmed GPΔmuc trimers exhibited lower thermostability (T_m_) than both wild-type and Kif-treated forms, glycan trimming enhanced NAb epitope recognition in both trimer and SApNP formats, consistent with our previous HIV-1 study (*113*).

In recent studies, we established a vaccine strategy that integrates detailed mechanistic analysis to characterize the in vivo behavior of immunogens and provide insights into vaccine-induced NAb responses (*113, 127, 128*). In the analysis of SUDV GPΔmuc vaccines, multivalent display on SApNPs enhanced GP interactions with FDC networks, B cells, and phagocytic cells in lymph nodes. The size and intrinsic thermostability of SApNPs contributed to prolonged vaccine retention and sustained antigen presentation in lymph node follicles (over 8 weeks), resulting in more robust and durable GC reactions compared with the soluble GPΔmuc trimer. Nearly identical trafficking and retention patterns were observed for SApNPs presenting SUDV GPΔmuc, HIV-1 Env (*113*), and influenza M2e×3 (*127*), despite their distinct glycan profiles. These results support our previous conclusion that glycans are not a major determinant of follicular localization for NP vaccines in lymph nodes (*113*). Collectively, these findings suggest that NP size and antigen thermostability are key factors influencing the lymph node behavior of protein NPs (*113, 127, 128*). Mouse immunization studies further elucidated antibody responses to diverse filovirus vaccines. Overall, longitudinal analysis revealed that NAb responses closely tracked with bAb responses. Among wild-type vaccines, E2p SApNPs presenting EBOV and SUDV GPΔmuc elicited the highest EC_50_ and ID_50_ titers at most timepoints, consistent with the strong GC reactions observed in lymph nodes. The two glycan modification strategies, however, yielded unexpected results. Glycan-trimmed trimers and SApNPs consistently elicited the lowest EC_50_ and ID_50_ titers, while oligomannose-rich immunogens – produced in the presence of kifunensine (Kif) – induced the strongest bAb and NAb responses at most timepoints. This pattern contrasts with findings from the previous studies of influenza HA, SARS-CoV-2 spike, and HIV-1 Env vaccines, where glycan trimming enhanced the breadth and potency of NAb responses (*112–115*). Although the reasons for these differences remain unclear, our results suggest that there may not be a single glycan modification strategy applicable to all viral GP vaccines.

Our future research will follow several directions. First, antibodies induced by GPΔmuc trimers and SApNPs, with or without glycan modification, will be isolated for structural epitope mapping. This will help determine how SApNP display enhances NAb function, as demonstrated for SARS-CoV-2 (*128*), how cross-NAbs recognize diverse GPs, and how glycan modification influences epitope targeting. Such analyses may explain why improved epitope exposure following glycan trimming did not translate into stronger NAb responses in vivo. Second, alternative glycan modification strategies will be explored. While it is unclear whether oligomannose enrichment improves NAb elicitation for other viral GPs, various approaches that alter glycan profiles can be tested using filovirus vaccine constructs. Non-mammalian systems, such as fungal (*161, 162*) and insect (*163*) cell lines, may provide simpler oligomannose-type glycans and reduce the need for chemical modification. Third, pan-filovirus vaccine strategies merit further investigation. EBOV and SUDV GPΔmuc SApNPs elicited cross-orthoebolavirus NAbs, and combining these with an optimized SApNP presenting RAVV GPΔmuc-P^2^CT trimers may, in principle, induce a cross-genus, pan-filovirus bNAb response. Finally, filovirus challenge studies in mice and guinea pigs will be essential to evaluate protection and support advancement to NHP models.

## METHODS

### Construct design, expression, and purification of filovirus GP immunogens

For orthoebolaviruses, the amino acid sequences of EBOV GP (Mayinga strain, GenBank Accession: NP_066246) with a T42A substitution, SUDV GP (Gulu strain, GenBank Accession: AAU43887), and BDBV GP (R4386L strain, GenBank Accession: AYI50307) were used to design GPΔmuc trimers (**Fig. S1a**) and SApNPs (**Fig. S3h**). Notably, an N637D mutation was later introduced to the SUDV GPΔmuc-WL^2^P^4^ construct to eliminate a potential N-linked glycosylation site at N637 due to the enzymatic site “AS” between the GP and SApNP backbone. For orthomarburgviruses, the amino acid sequence of RAVV GP (Ravn-87 strain, GenBank Accession: ABE27071) was used to design GPΔTM and various GPΔmuc trimers (**Fig. S5a**).

Rationally designed EBOV, SUDV, and BDBV GPΔmuc trimers were transiently expressed in HEK293F cells as previously described (*100*). Briefly, HEK293F cells were thawed and suspended in FreeStyle 293 Expression Medium (Life Technologies, Carlsbad, CA) and placed in a shaker incubator at 37°C, 135 rotations per minute (rpm), with 8% CO_2_. When the cells reached a density of 2.0 × 10^6^/ml, they were diluted to a concentration of 1.0 × 10^6^/ml in 1 L of FreeStyle 293 Expression Medium for transfection using a polyethyleneimine (PEI) (Polysciences) transfection protocol. In brief, 900 μg of plasmid DNA in 25 ml of Opti-MEM transfection medium (Life Technologies) was mixed with 5 ml of PEI-MAX (1.0 mg/ml) in 25 ml of Opti-MEM and incubated for 30 min at room temperature. Cells were then transfected with the DNA–PEI-MAX mixture and incubated in a shaker incubator at 37°C, 135 rpm, with 8% CO_2_. Kifunensine (10 mg/L, Tocris Bioscience) was added at the time of transfection to inhibit α-mannosidase I and promote oligomannose-type glycosylation. Five days later, the supernatants containing the target protein were harvested, clarified by centrifugation at 1,126 × g for 22 min, and filtered through a 0.45 μm membrane (Thermo Scientific). Orthoebolavirus GP was purified with an ADI-15878 IAC column, except for EBOV GPΔmuc-WL^2^P^4^ used in X-ray crystallography and EBOV GPΔmuc-WL^2^P^4^-I3-01v9b/c-T used in nsEM, which were purified with an mAb100 IAC column. Orthoebolavirus GPs were eluted three times with 5 ml of 0.2 M glycine (pH 2.2), neutralized with 0.5 ml of 2 M Tris base (pH 9.0), and buffer-exchanged into phosphate-buffered saline (PBS; pH 7.2). IAC-purified orthoebolavirus GPs were further purified by SEC using a Superdex 200 Increase 10/300 GL column (Cytiva).

RAVV GPΔmuc trimers and all orthoebolavirus GPΔmuc-presenting SApNPs were produced in ExpiCHO cells (Thermo Fisher). Briefly, ExpiCHO cells were thawed and incubated in ExpiCHO Expression Medium (Thermo Fisher) in a shaker incubator at 37°C, 135 rpm, with 8% CO_2_. When cell density reached 10 × 10^6^/ml, ExpiCHO Expression Medium was added to adjust the cell density to 6 × 10^6^/ml for transfection. ExpiFectamine CHO–plasmid DNA complexes were prepared for 100-ml transfections according to the manufacturer’s instructions. Kifunensine (10 mg/L) was added at the time of ExpiCHO transfection to inhibit α-mannosidase I. For these constructs, 100 μg of plasmid and 320 μl of ExpiFectamine CHO reagent were mixed in 7.7 ml of cold OptiPRO™ medium (Thermo Fisher). Following the first feed on day 1, cells were cultured in a shaker incubator at 33°C, 115 rpm, with 8% CO_2_ using the Max Titer protocol, with an additional feed on day 5 (Thermo Fisher). Culture supernatants were harvested 13–14 days post-transfection, clarified by centrifugation at 3724 × g for 25 min, and filtered through a 0.45 μm membrane (Thermo Fisher). RAVV GPΔmuc trimers were purified using Ni Sepharose excel resin (Nickel, Cytiva), eluted twice with 15 ml of 0.5 M imidazole, and buffer-exchanged into Tris-buffered saline (TBS; pH 7.2). Orthoebolavirus GPΔmuc-presenting SApNPs were purified using an ADI-15878 IAC column, eluted three times with 5 ml of 0.2 M glycine (pH 2.2), neutralized with 0.5 ml of 2 M Tris base (pH 9.0), and buffer-exchanged into PBS (pH 7.2). Final polishing was performed by SEC using a Superdex 200 Increase 10/300 GL column (Cytiva) for nickel-purified RAVV GPΔmuc trimers and a Superose 6 Increase 10/300 GL column (Cytiva) for IAC-purified orthoebolavirus GPΔmuc-presenting SApNPs. Selected SEC fractions were pooled, aliquoted, and stored in liquid nitrogen or at −80 ⁰C until use.

### Expression and purification of neutralizing antibodies (NAbs)

Antibodies in the IgG form were transiently expressed in ExpiCHO cells (Thermo Fisher). At 12– 14 days post-transfection, cells were centrifuged at 3,724 × *g* for 25 min, and the supernatants were filtered through a 0.45-μm membrane (Millipore). IgGs were purified using protein A affinity resin (Cytiva) and eluted with 0.3 M citric acid (pH 3.0). The eluate was immediately titrated to neutral pH (7.0) by adding 2 M Tris base (pH 9.0). IgGs were then concentrated and buffer-exchanged into PBS using Amicon 10 kDa filters (Millipore). Concentrations were determined by UV_280_ absorption using theoretical extinction coefficients.

### Glycan trimming by Endo H treatment and enzyme removal

The protocol for Endo H treatment and removal was described in our previous study (*113*). Briefly, surface glycans of filovirus GPΔmuc trimers and SApNPs were trimmed using endoglycosidase H (Endo Hf), a fusion of Endo H and MBP (NEB, catalog no. P0703L). In each reaction, 1 mg of antigen was mixed with 200 μl of 10× GlycoBuffer 3, 250 μl of Endo Hf, and H_2_O (if necessary) to a final volume of 2 ml. The mixture was incubated at room temperature (25°C) for 5 h to allow enzymatic processing of the GP glycans. After incubation, the reaction mixtures were purified by SEC using a Superdex 200 Increase 10/300 GL column (for trimers) or a Superose 6 Increase 10/300 GL column (for SApNPs) to remove the MBP-tagged enzyme. Notably, glycan trimming by Endo H requires expression of filovirus GPΔmuc immunogens in the presence of kifunensine. In our previous study (*113*), amylose resin (NEB, catalog no. E8021S) was used to remove residual Endo Hf from SEC-purified HIV-1 Env trimers and SApNPs. In the present study, this additional step of Endo Hf removal was omitted to preserve vaccine antigens for animal studies. SEC-purified fractions were pooled, aliquoted, and stored in liquid nitrogen or at −80 ⁰C until use.

### SDS-PAGE and BN-PAGE

Filovirus GPΔmuc trimers and SApNPs were analyzed by sodium dodecyl sulfate-polyacrylamide gel electrophoresis (SDS-PAGE) and blue native-polyacrylamide gel electrophoresis (BN-PAGE). Proteins were mixed with loading dye and loaded onto a 10% Tris-Glycine Gel (Bio-Rad) or a 4-12% Bis-Tris NativePAGE gel (Life Technologies). For SDS-PAGE under reducing conditions, proteins were mixed with 6× Laemmli SDS sample buffer (Thermo Scientific) and boiled for 5 min at 100 °C. SDS-PAGE gels were run at 250 V for 25 min using SDS running buffer (Bio-Rad) and stained with InstantBlue (Abcam). For BN-PAGE, proteins were mixed with 4× native dye and loaded onto a BN-PAGE gel. The gel was run at 150 V for 2–2.5 h with NativePAGE running buffer (Life Technologies), following the manufacturer’s instructions. BN-PAGE gels were stained with Coomassie Brilliant Blue R-250 (Bio-Rad) and destained using a solution of 6% ethanol and 3% glacial acetic acid.

### Differential scanning calorimetry (DSC)

The melting temperature (T_m_) and other thermal parameters of filovirus GPΔmuc trimers purified by an IAC or nickel column followed by SEC were measured using a MicroCal PEAQ-DSC Man instrument (Malvern). Briefly, purified proteins in PBS buffer were diluted to 0.5-5 μM. Melting was assessed at a scan rate of 60 °C/h from 20 °C to 100 °C. Data processing, including buffer correction, normalization, and baseline subtraction, was conducted using MicroCal PEAQ-DSC software. Gaussian fitting was carried out using GraphPad Prism version 10.3.1.

### Site-specific glycan analysis

To generate site-specific glycan profiles, 100 µg aliquots of each antigen were denatured for 1 h in 50 mM Tris/HCl (pH 8.0) containing 6 M urea and 5 mM dithiothreitol (DTT). The denatured antigens were reduced and alkylated by adding 20 mM iodoacetamide (IAA) and incubating for 1 h in the dark, followed by 1h of incubation with 20 mM DTT to eliminate residual IAA. Alkylated samples were buffer-exchanged into 50 mM Tris/HCl (pH 8.0) using Vivaspin columns (10 kDa cutoff). Three aliquots were then digested separately overnight with trypsin (Mass Spectrometry Grade, Promega), chymotrypsin (Mass Spectrometry Grade, Promega), or alpha lytic protease (Sigma Aldrich) at a ratio of 1:30 (w/w). The next day, peptides were dried and extracted using an Oasis HLB µElution Plate (Waters).

Peptides were dried again, resuspended in 0.1% formic acid, and analyzed by nanoLC-electrospray ionization MS using an Ultimate 3000 HPLC (Thermo Fisher Scientific) system coupled to an Orbitrap Eclipse mass spectrometer (Thermo Fisher Scientific). Stepped higher-energy collisional dissociation (HCD) fragmentation was applied. Peptides were separated using an EasySpray PepMap RSLC C18 column (75 µm × 75 cm). A trapping column (PepMap 100 C18 3μM 75μM × 2cm) was used in line with liquid chromatography (LC) prior to separation with the analytical column. The LC conditions were as follows: 280-minute linear gradient consisting of 4-32% acetonitrile in 0.1% formic acid over 260 minutes followed by 20 minutes of alternating 76% acetonitrile in 0.1% formic acid and 4% ACN in 0.1% formic acid, used to ensure all the sample had eluted from the column. The flow rate was set to 300 nl/min. The spray voltage was set to 2.7 kV and the temperature of the heated capillary was set to 40 °C. The ion transfer tube temperature was set to 275 °C. The scan range was 375−1500 m/z. The stepped HCD collision energies were set to 15%, 25%, and 45% and the MS2 for each energy was combined. Precursor and fragment detection were performed using an Orbitrap at a resolution of MS^1^ = 120,000 and MS^2^ = 30,000. The AGC target for MS1 was set to standard and injection time set to auto which involves the system setting the two parameters to maximize sensitivity while maintaining cycle time. Full LC and MS methodology can be extracted from the appropriate Raw file using XCalibur FreeStyle software or upon request.

Glycopeptide fragmentation data were extracted from the raw file using Byos 4.6 (Protein Metrics). The glycopeptide fragmentation data were evaluated manually for each glycopeptide; the peptide was scored as true-positive when the correct b and y fragment ions were observed along with oxonium ions corresponding to the glycan identified. The MS data was searched using the Protein Metrics 38 insect N-glycan library. The relative amounts of each glycan at each site as well as the unoccupied proportion were determined by comparing the extracted chromatographic areas for different glycotypes with an identical peptide sequence. All charge states for a single glycopeptide were summed. The precursor mass tolerance was set at 4 ppm and 10 ppm for fragments. A 1% false discovery rate (FDR) was applied. The relative amounts of each glycan at each site as well as the unoccupied proportion were determined by comparing the extracted ion chromatographic areas for different glycopeptides with an identical peptide sequence. Glycans were categorized according to the composition detected.

HexNAc(2)Hex(9−3) was classified as M9 to M3. Any of these structures containing a fucose were categorized as fucosylated mannose (FM). Complex-type glycans were classified according to the number of HexNAc subunits and the presence or absence of fucose. Core glycans refer to truncated structures smaller than M3. As this fragmentation method does not provide linkage information, compositional isomers are grouped.

### Biolayer interferometry

Antigenic profiles of filovirus GPΔmuc trimers and SApNPs were measured on an Octet RED96 (FortéBio, Pall Life Sciences) against a panel of antibodies, including five pan-orthoebolavirus NAbs and two MARV NAbs, all in the IgG form. All assays were performed with agitation set to 1000 rpm in FortéBio 1× kinetic buffer. The final volume for all solutions was 200 μl per well. Experiments were conducted at 30 °C in solid black 96-well plates (Greiner Bio-One). For all antigens, 5 μg/ml antibody in 1× kinetic buffer was loaded onto the surface of anti-human IgG Quantitation (AHQ) Biosensors for 300 s. Next, a 2-fold concentration gradient of antigen, starting at 500 nM for trimers and 10 nM for SApNPs, was used in a 6-point dilution series. A 60-s biosensor baseline step was applied before measuring association of antigen in solution with antibody-loaded biosensors for 200 s. Dissociation of the interaction was monitored for 300 s. The correction of baseline drift was performed by subtracting the mean value of shifts recorded for a sensor loaded with antibody but not incubated with antigen, and for a sensor without antibody but incubated with antigen. Octet data were processed using FortéBio’s data acquisition software (version 8.1). Peak response signals at the highest antigen concentration were summarized in a matrix and color-coded for comparison across constructs. Experimental data for each antigen-antibody pair were fitted to a 1:1 binding model, and the three best-fitting datasets were grouped to calculate K_on_ and K_D_ values. Antibody-antigen binding curves were plotted using GraphPad Prism version 10.3.1.

### Enzyme-linked immunosorbent assay (ELISA)

Costar 96-well, high-binding, flat-bottom, half-area plates (Corning) were coated with 50 µl of PBS containing 0.1 μg of the appropriate filovirus GPΔmuc trimer or SApNP antigen. Plates were incubated overnight at 4 °C and then washed five times with PBST wash buffer (PBS containing and 0.05% (v/v) Tween 20). Each well was then blocked for 1 h at room temperature with 150 µl of blocking buffer consisting of 4% (w/v) blotting-grade blocker (Bio-Rad) in PBS. Plates were washed again five times with PBST wash buffer. To evaluate antibody binding to the coating antigens, antibodies were diluted in blocking buffer to a maximum concentration of 10 µg/ml, followed by a 10-fold dilution series. For each antibody dilution, a total volume of 50 µl was added to the appropriate wells. For serum sample analysis, mouse sera were diluted 1:40 in blocking buffer and subjected to a 10-fold dilution series. For each sample dilution, 50 µl of each dilution was added to the wells. Plates were incubated for 1 h at room temperature and then washed five times with PBST wash buffer. For secondary antibody binding, a 1:5000 dilution of goat anti-human IgG antibody (Jackson ImmunoResearch Laboratories) or, for mouse sample analysis, a 1:3000 dilution of horseradish peroxidase (HRP)-conjugated goat anti-mouse IgG antibody (Jackson ImmunoResearch Laboratories) was prepared in PBST wash buffer. Then, 50 µl of the diluted secondary antibody was added to each well. Plates were incubated for 1 h at room temperature and then washed six times with PBST wash buffer. Finally, wells were developed with 50 µl of 3,3’,5,5’-tetramethylbenzidine (Life Sciences) for 3-5 min before the reaction was stopped with 50 µl of 2 N sulfuric acid. Plates were immediately read on a BioTek Synergy plate reader at a wavelength of 450 nm. EC_50_ values were calculated from full curves using GraphPad Prism version 10.3.1. For samples with OD_450_ values below 0.5, EC_50_ values were set to 10 µg/ml in **Figs. 1g**, **3h**, **4c**, and **4h** to facilitate EC_50_ plotting and comparison.

### Protein crystallization and data collection

EBOV GPΔmuc-WL^2^P^4^ was transiently expressed in HEK293S cells using the same protocol as for HEK293F cells, followed by IAC and SEC purification. The mAb100/SEC-purified EBOV GPΔmuc-WL^2^P^4^ trimer protein at 9.3 mg/ml was subjected to crystallization by sitting-drop vapor diffusion using an automated CrystalMation robotic system (Rigaku) at 20 °C at The Scripps Research Institute (*164*). The reservoir solution consisted of 12% PEG-6000 (*v/v*) and 0.1 M sodium citrate-citric acid (pH 4.51). Diffraction-quality crystals were obtained after 2 weeks of incubation at 20°C. Crystals were cryoprotected in well solution containing 25% (*v/v*) ethylene glycol, mounted on a nylon loop, and flash-cooled in liquid nitrogen. Diffraction data were collected at beamline 12-1 of the Stanford Synchrotron Radiation Lightsource (SSRL) and processed with HKL-2000 (*165*). The data were indexed in space group P321.

### Structure determination and refinement

The crystal structure of EBOV GPΔmuc-WL^2^P^4^ was determined by molecular replacement using Phaser-MR in Phenix (version 1.19.2-4158), with the coordinates of the previously reported EBOV GPΔmuc-WL^2^P^2^ structure (PDB ID: 7JPH) (*100*) as the search model. Polypeptide chains were manually adjusted into electron density using Coot (*166*), refined with *phenix.refine* (*167*), and validated using the wwPDB Validation System (*168*). Carbohydrates were further validated using Privateer in CCP4i (*169*), both before and after refinement. Final data processing and refinement statistics are described in **Table S1**. All images for crystal structures shown in the figures were generated using PyMOL version 2.3.4. Notably, due to weak electron density, structural models were not built for residues K633-D637 in the GP2 stalk region of EBOV GPΔmuc-WL^2^P^4^, nor for the C-terminal foldon trimerization motif. These omissions result in apparent gaps in crystal packing when visualizing the GP structure within the electron density map.

### Negative-stain electron microscopy (nsEM)

The nsEM analysis of filovirus GPΔmuc trimers, their antibody-bound complexes, and GPΔmuc-presenting SApNPs, was performed by the Core Microscopy Facility at The Scripps Research Institute. Briefly, samples were prepared at concentrations of 0.008 and 0.01 mg/ml, respectively. Carbon-coated copper grids (400 mesh) were glow-discharged, and 8 µl of each sample was adsorbed for 2 min. Excess samples were removed, and grids were negatively stained with 2% uranyl formate for 2 min. Excess stain was wicked away, and the grids were allowed to air-dry. Trimer samples were analyzed at 120 kV with a Talos L120C transmission electron microscope (Thermo Fisher), and images were acquired using a CETA 16 M CMOS camera under 73,000× magnification at a resolution of 1.93 Å/pixel and a defocus range of 0.5-2 µm. For SApNP samples, the images were collected under 52,000× magnification. The resulting pixel size at the specimen plane was 2.05 Å, and the defocus was set to −1.50 µm. Computational analysis of the images was performed using the high-performance computing core facility at The Scripps Research Institute. nsEM images were converted to MRC format using EMAN2 (*170*) and further processed with CryoSPARC version 4.3.0 (*124*). Micrographs were contrast-transfer function (CTF)-corrected by patch CTF estimation. Particles were selected using a blob/template picker and later extracted with a box size of 130 pixels for 2D classification, with 50 and 230 Å used as the minimum and maximum particle sizes for blob picking. 3D models were generated by *ab initio* reconstruction and optimized using heterogeneous and homogeneous refinement procedures. All nsEM and fitted structure images were generated using UCSF Chimera (*171*) and ChimeraX (*172–174*).

### Cryo-electron microscopy (cryo-EM) for SUDV GP-CA45 complexes

The SUDV GPΔmuc-WL^2^P^4^ construct was expressed in HEK293F cells and purified using an ADI-15878 column. The eluate was mixed with CA45 Fab at a 1:1.5 molar ratio. The mixture was further purified using a Superose 6 Increase 10/300 GL column, and complex fractions were pooled. Cryo-EM grid preparation and data collection were performed at the UC San Diego Cryo-EM Facility. Vitrification was carried out using a Vitrobot Mark IV (FEI) equilibrated to 4 ⁰C and 100% humidity. Quantifoil Cu 1.2/1.3 300 grids were plasma cleaned using a Solarus II (Gatan) with a mixture of Ar/O_2_ for 10 s. Following plasma cleaning, the grid was blotted on one side with 3.5 µl of the SUDV GPΔmuc-WL^2^P^4^/CA45 Fab complex at a concentration of 0.55 mg/ml for 4 s, and frozen in liquid ethane. The frozen grids were stored in liquid nitrogen until imaging. Cryo-EM data were collected on a Titan Krios G4 (Thermo Fisher) operated at 200 keV, equipped with a Falcon 4 camera and a Selectris-X energy filter (Thermo Fisher). Automated data collection was performed using Smart EPU software (Thermo Fisher). Images were acquired under 130,000× magnification, corresponding to a pixel size of 0.89 Å, with a defocus range of −1 to −2.5 µm. All datasets were processed using CryoSPARC version 3.3. Movie frames were aligned using Patch Motion Correction, and CTF estimation was performed with Patch CTF Estimation (**Fig. S2d**). Initial particle picking was done with Blob Picker with the particle size set to 100-350 Å, and particles were extracted using a box size of 360 pixels. After 2D classification, selected particle classes were used as input for Template Picking and re-extracted with a box size of 360 pixels. After multiple rounds of 2D classification, 249,139 particles were selected for 3D reconstruction. Three starting reference models were generated using Ab-Initio Reconstruction and used for Heterogeneous Refinement. Particles in the largest cluster (163,924 particles) were refined using Homogeneous Refinement with a focusing mask and C3 symmetry. The refined model was further subjected to 3D Classification, resulting in 10 models. The class associated with the most complete model (41,233 particles) was subjected to another round of Ab-Initio Reconstruction. This intermediate model was further optimized using Non-uniform Refinement and Local Refinement, both using 163,924 particles and a focusing mask. The final GSFSC resolution was 3.13 Å, and the final structural model was built using the crystal structures of EBOV GPΔmuc-WL^2^P^2^ (PDB ID: 7JPH) (*100*) and EBOV GPΔmuc-bound CA45 Fab (PDB ID: 6EAY) (*118*) as templates. The structures were fitted into the density map using ChimeraX (*172–174*), followed by real-space refinement in Phenix (*167*), manual model correction in Coot (*166*), and a final round of real-space refinement in Phenix (*167*). Statistics for the cryo-EM reconstruction and model refinement are provided in **Table S2**. The coordinates and 3D density map have been deposited in Protein Data Bank (PDB) and Electron Microscopy Data Bank (EMDB), respectively.

### Rational design of trimer-presenting I3-01v9b/c SApNP scaffolds

We previously redesigned the I3-01v9 scaffold for optimal display of monomeric antigens (*127*). In the resulting I3-01v9a SApNP, the *N*-terminal helix was extended such that its first residue was positioned just above the NP surface. Here, we further redesigned I3-01v9a to optimize NP display of trimeric antigens with a long coiled-coil stalk (**Fig. S3**). First, the 11-aa *N*-terminal helix in I3-01v9a was truncated to 7 aa. Then, a 13-aa helix-turn fragment (9-aa helix + 4-aa turn, all alanine) was fused to this truncated N-terminus. The newly added 9-residue helix was designed to pack within the groove formed by two helices in the I3-01 core. This 9-residue helix was derived from the structure of a human transcription factor (PDB ID: 6G6L). To accommodate this new helix, four mutations were introduced into the I3-01 core helices to remove steric clashes. The helix-turn backbone was first relaxed using iterative modular optimization (IMO) (*175*), followed by structural sampling with the CONCOORD program (*176*) to generate 1,000 slightly perturbed backbone conformations. An ensemble-based protein design protocol (*87, 152*) was then used to identify the optimal amino acid sequence for the 9-residue N-terminal helix based on Cα and Cβ-based RAPDF (*177*) scoring functions. The 4-residue turn was fixed as “GSGS” during design. The final I3-01v9b design was selected by combining the results from both scoring functions. A second design, I3-01v9c, was derived by mutating the turn sequence from “GSGS” to “GPPS” to increase rigidity. The I3-01v9b structural model was further relaxed using IMO with a minimum backbone perturbation angle. The distance between the N-termini of the I3-01v9b/c trimer model (I3-01v9b/c-T) was 43.9 Å. EBOV GPΔmuc-WL^2^P^4^-I3-01v9b/c-T constructs (**Fig. S3b**) were designed, transiently expressed in HEK293F cells, and structurally validated by nsEM.

### Dynamic Light Scattering (DLS)

Particle size distributions of orthoebolavirus GPΔmuc-presenting SApNPs were measured using a Zetasizer Ultra instrument (Malvern). Briefly, ADI-15878/SEC-purified SApNPs produced from ExpiCHO cells were diluted to 0.2 mg/ml in 1× PBS, and 30 μl of each sample was added to a quartz batch cuvette (Malvern, catalog no. ZEN2112). Measurements were performed at 25 °C in backscatter mode. Data processing was carried out using the Zetasizer particle size analyzer with the Ultra instrument software. The resulting particle size distributions were plotted using GraphPad Prism version 10.3.1.

### Mouse immunization and sample collection

Mouse immunization protocols were adapted from our previous studies (*113, 127, 128*). All animal experiments were conducted in accordance with the guidelines of the Association for the Assessment and Accreditation of Laboratory Animal Care (AAALAC) and approved by the Institutional Animal Care and Use Committee (IACUC) of The Scripps Research Institute. Female and male BALB/c mice, aged 6–8 weeks, were purchased from The Jackson Laboratory and housed in ventilated cages in environmentally controlled rooms at 20°C with 50% humidity, under a 12 h/12 h light-dark cycle. For mechanistic studies of vaccine transport and immune responses in lymph nodes, mice were intradermally inoculated via four footpads using a 29-gauge insulin needle under 3% isoflurane anesthesia with oxygen. The injection dose was 80 μl of antigen and AP adjuvant mix, containing 40 μg of vaccine antigen per mouse or 10 μg per footpad. In the first immunogenicity study, mice were injected via the intraperitoneal (i.p.) route at weeks 0, 3, 9, and 15. A 6-week interval was maintained between the second and third, and the third and fourth doses.

In the second immunogenicity study, our standard 4-dose screening regimen (*113, 127, 128, 158*) was used, with mice injected at weeks 0, 3, 6, and 9 at 3-week intervals. Each injection consisted of 200 μl of antigen/adjuvant mix containing 10 μg of vaccine antigen and 100 μl of adjuvant, including 40 µg/50 µl of CpG (oligonucleotide 1826, a TLR9 agonist from InvivoGen) adsorbed onto 50 µl of AH (InvivoGen). In the sex-dependent antibody response study, mice were injected intradermally via the footpads at weeks 0, 3, 6, and 9, with 3-week intervals between doses. Each mouse received 80 μl of an antigen/AH adjuvant mixture containing 10 μg of immunogen and 40 μl of adjuvant. Blood samples were collected 2 weeks after each injection through the retro-orbital sinus using heparinized capillary tubes. Samples were centrifuged at 12,000 rpm for 10 min to separate serum (top layer) and the rest of the whole blood layer. Upon heat inactivation at 56°C for 30 min, serum was centrifuged again at 12000 rpm for 10 min to remove precipitates. For the last timepoint, the rest of the whole blood layer was diluted with an equal volume of PBS and then overlaid on 4.5 ml of Ficoll in a 15 ml SepMate tube (STEMCELL Technologies) and centrifuged at 1,200 rpm for 10 min at 20 °C to separate peripheral blood mononuclear cells (PBMCs). Cells were washed once in PBS and then resuspended in 1 ml of ACK red blood cell lysis buffer (Lonza). After washing with PBS, PBMCs were resuspended in 1 ml of freezing media (10% dimethyl sulfoxide [DMSO]/90% fetal calf serum [FCS]). Two weeks after the last bleed, spleens were harvested and ground against a 70-μm cell strainer (BD Falcon) to release splenocytes into a cell suspension. Splenocytes were centrifuged, washed in PBS, treated with 5 ml of ACK lysing buffer (Lonza), and frozen with 3 ml of freezing media (10% DMSO/90% FCS). Serum samples from week 11 in the second study were purified using a CaptureSelect™ IgG-Fc (Multispecies) Affinity Matrix (Thermo Scientific) following the manufacturer’s instructions. Purified IgG from individual mice was analyzed in filovirus-pp assays to assess vaccine-induced NAb responses.

### Histology, immunostaining, and imaging for lymph node tissues

To study vaccine distribution, trafficking, retention, cellular interaction, and GC reactions in sentinel lymph nodes, SUDV GPΔmuc trimer and SApNPs (E2p and I3-01v9b) adjuvanted with AP were intradermally injected into four footpads of mice. A similar protocol was described in our previous studies (*113, 127, 128*). Briefly, each injection consisted of 80 μl of antigen/adjuvant mix containing 40 μg of filovirus antigen (10 μg/footpad; 40 μg/mouse). Mice were euthanized at time points ranging from 2 h to 8 weeks after a single-dose injection. Fresh sentinel lymph nodes were collected and embedded in frozen section compound (VWR International, catalog no. 95057-838). Sample molds were immersed in liquid nitrogen and stored at −80°C before shipping to The Centre for Phenogenomics (Toronto, Canada). Lymph node tissues were sectioned at 8 μm thickness using a cryostat (Cryostar NX70). Primary antibodies were applied to lymph node sections and incubated overnight at 4°C. After washing with TBS containing 0.1% Tween-20 (TBST), biotin- or fluorophore-conjugated secondary antibodies were added, and sections were incubated at 25 °C for 1 h. Lymph node sections were stained with NAbs ADI-15878 (*116*), ADI-15946 (*119*), CA45 (*118*), or mAb100 (*133*) (1:100), followed by biotinylated goat anti-human secondary antibody (Abcam, catalog no. ab7152, 1:300), streptavidin-horseradish peroxidase (HRP) reagent (Vectastain Elite ABC-HRP Kit, Vector, catalog no. PK-6100), and ImmPACT diaminobenzidine (DAB; Vector, catalog no. SK-4105).

To study the interaction between SUDV GPΔmuc immunogens with lymph node cell populations, FDCs were labeled using anti-CD21 primary antibody (Abcam, catalog no. ab75985, 1:1800), followed by anti-rabbit secondary antibody conjugated with Alexa Fluor 555 (Thermo Fisher, catalog no. A21428, 1:200). FDCs were also co-stained using anti-FDC-M1 primary antibody (BD Biosciences, catalog no. 551320, 1:20), followed by anti-rat secondary antibody conjugated with Alexa Fluor 488 (Abcam, catalog no. ab150165, 1:200). Subcapsular sinus macrophages were labeled with anti-sialoadhesin (CD169) antibody (Abcam, catalog no. ab53443, 1:600), followed by anti-rat secondary antibody conjugated with Alexa Fluor 488 (Abcam, catalog no. ab150165, 1:200). B cells were labeled with anti-B220 antibody (eBioscience, catalog no. 14-0452-82, 1:100), followed by anti-rat secondary antibody conjugated with Alexa Fluor 647 (Thermo Fisher, catalog no. A21247, 1:200). GC responses were assessed by immunohistology. GC B cells were stained with FITC-conjugated rat anti-GL7 antibody (BioLegend, catalog no. 144604, 1:250). T_fh_ cells were stained with anti-CD4 antibody (BioLegend, catalog no. 100402, 1:100), followed by anti-rat secondary antibody conjugated with Alexa Fluor 488 (Abcam, catalog no. ab150165, 1:1000). GC cells were stained with Bcl6 antibody (Abcam, catalog no. ab220092, 1:300), followed by anti-rabbit secondary antibody conjugated with Alexa Fluor 555 (Thermo Fisher, catalog no. A21428, 1:1000). Nuclei were stained with 4’,6-diamidino-2-phenylindole (DAPI; Sigma-Aldrich, catalog no. D9542, 100 ng/ml).

Stained lymph node tissues were scanned using an Olympus VS-120 slide scanner equipped with a Hamamatsu ORCA-R2 C10600 digital camera or an Akoya Vectra Polaris imaging system with a 20×/0.45NA objective. The transport of SUDV GPΔmuc trimer and SApNP immunogens, as well as the GCs they induced in lymph nodes, were quantified from bright-field and fluorescent images using ImageJ software (*178*).

### Electron microscopy analysis of protein nanoparticles in lymph node tissues

To assess interactions between SUDV GPΔmuc immunogens and stromal and immune cells in lymph nodes, TEM analysis was performed by the Core Microscopy Facility at The Scripps Research Institute (TSRI). The protocol for lymph node isolation, processing, and analysis followed our previous studies (*113, 128*). To visualize AP-adjuvanted SUDV GPΔmuc SApNPs (E2p and I3-01v9b) interacting with FDCs and phagocytic cells, mice were injected with 200 μl of antigen/adjuvant mix containing 100 μg of SUDV immunogen and 100 μl of AP adjuvant, administered into two hind footpads (50 μg/footpad). Fresh popliteal lymph nodes were collected at 2, 12, and 48 h after a single-dose injection. Lymph node tissues were bisected and immersed in oxygenated fixative containing 2.5% glutaraldehyde and 4% paraformaldehyde in 0.1 M sodium cacodylate buffer (pH 7.4) overnight at 4°C. Lymph node samples were then fixed and stained with 0.5% uranyl acetate overnight at 4°C, infiltrated with LX-112 (Ladd) epoxy resin, and polymerized at 60 °C. Ultrathin tissue sections (70 nm) were cut and mounted on copper grids for imaging. TEM was performed at 80 kV using a Talos L120C transmission electron microscope (Thermo Fisher), and images were acquired with a CETA 16M CMOS camera for further analysis.

### Lymph node disaggregation, cell staining, and flow cytometry

To assess vaccine-induced humoral immune responses, the frequency and number of GC B cells (GL7^+^B220^+^) and T_fh_ cells (CD3^+^CD4^+^CXCR5^+^PD-1^+^) were measured by flow cytometry to quantify GC reactions (**Fig. S9**). The protocol for lymph node tissue collection, processing, and analysis followed our previous studies (*113, 127, 128*). Briefly, mice were administered with 80 μl of antigen/adjuvant mix containing 40 μg of vaccine immunogen (10 μg/injection, 40 μg/mouse) into four footpads. Mice were euthanized at 2, 5, and 8 weeks after a single-dose injection, or at 2 and 5 weeks after the boost. Fresh sentinel lymph nodes were isolated and immediately processed by mechanical disaggregation. Lymph node tissues were then incubated in enzyme digestion solution at 37 °C for 30 min on a rotator, with the resulting cells filtered through a 70 μm cell strainer to obtain a single-cell suspension. Anti-CD16/32 antibody (BioLegend, catalog no. 101302, 1:50) was added to the cell samples to block the nonspecific binding of Fc receptors and the samples were kept on ice for 30 min. Next, cells were transferred to 96-well V-shaped-bottom microplates with a cocktail of antibodies and stains, including: Zombie NIR live/dead stain (BioLegend, catalog no. 423106, 1:100), Brilliant Violet 510 anti-mouse/human CD45R/B220 antibody (BioLegend, catalog no. 103247, 1:300), FITC anti-mouse CD3 antibody (BioLegend, catalog no. 100204, 1:300), Alexa Fluor 700 anti-mouse CD4 antibody (BioLegend, catalog no. 100536, 1:300), PE anti-mouse/human GL7 antibody (BioLegend, catalog no. 144608, 1:500), Brilliant Violet 605 anti-mouse CD95 (Fas) antibody (BioLegend, catalog no. 152612, 1:500), Brilliant Violet 421 anti-mouse CD185 (CXCR5) antibody (BioLegend, catalog no. 145511, 1:500), and PE/Cyanine7 anti-mouse CD279 (PD-1) antibody (BioLegend, catalog no. 135216, 1:500). Cells were incubated with the antibody cocktail on ice for 30 min and centrifuged at 400 × *g* for 10 min. They were then fixed with 1.6% paraformaldehyde (Thermo Fisher Scientific, catalog no. 28906) in Hank’s Balanced Salt Solution (HBSS) on ice for 30 min, centrifuged again at 400 × *g* for 10 min, resuspended in HBSS blocking solution at 4 °C. Sample acquisition was performed on a 5-laser AZE5 flow cytometer (Yeti, Bio-Rad) with Everest software at the Core Facility of The Scripps Research Institute. Data analysis was conducted with FlowJo version 10.

### Pseudovirus production and neutralization assays

A filovirus-pseudoparticle (pp) neutralization assay (*100*) was performed to assess the neutralizing activity of vaccine-induced mouse sera and purified mouse IgG. Filovirus-pps were generated by co-transfecting HEK293T cells with the envelope-deficient HIV-1 pNL4-3.lucR^−^E^−^ plasmid (NIH AIDS Reagent Program: https://www.aidsreagent.org/) and an expression plasmid encoding the GP gene from EBOV Makona (GenBank accession no. KJ660346), SUDV Gulu (GenBank accession no. AY729654), BDBV Uganda (GenBank accession no. KR063673), or MARV Angola (GenBank accession no. DQ447653) at a 4:1 ratio using the Lipofectamine 3000 transfection protocol (Thermo Fisher). After 72 h, pseudoviruses were harvested from the supernatant by centrifugation at 3,724 × *g* for 10 min, aliquoted, and stored at −80°C until use. For pseudovirus neutralization assays, mouse sera (starting dilution 1:100) or purified IgG (starting at 300 μg/ml) were serially diluted 3-fold and incubated with pseudoviruses at 37°C for 1 h in white solid-bottom 96-half-well plates (Corning). Then, 0.8 × 10^4^ Huh-7 cells were added to each well, and the plates were incubated at 37°C for 60-72 h. After incubation, the supernatant was removed, and cells were lysed. Luciferase reporter gene expression was quantified by adding Bright-Glo Luciferase substrate (Promega) according to the manufacturer’s instructions. Luciferase activity (in relative light units, RLU) was measured on a BioTek microplate reader using Gen 5 software. Values from experimental wells were compared to virus-only wells, with background luminescence from a series of uninfected wells subtracted from both. Dose-response neutralization curves were fit by nonlinear regression (0-100% constraints) in GraphPad Prism version 10.3.1, from which IC_50_ and ID_50_ were calculated. The same protocols were used to produce and assay pseudoparticles displaying the envelope glycoprotein of lymphocytic choriomeningitis virus (LCMV-pps) as a negative control for IgG neutralization.

### Statistical analysis

Data were collected from *n* = 8 mice per group in the immunization studies and *n* = 3-5 mice per group in the mechanistic study. All ELISA and pseudovirus neutralization assays were performed in duplicate. Different vaccine constructs and adjuvant-formulated filovirus vaccine antigens were compared using one-way analysis of variance (ANOVA), followed by Tukey’s multiple comparison *post hoc* test. Two-tailed unpaired *t-*tests were used to compare two independent vaccine groups. Statistical significance is indicated in the figures as follows: ns (not significant), \**p* < 0.05, \*\**p* < 0.01, \*\*\**p* < 0.001, and \*\*\*\**p* < 0.0001. Graphs were generated and statistical analyses were performed using GraphPad Prism version 10.3.1.

## ACKNOWLEDGEMENTS

The authors acknowledge the facilities of the UC San Diego cryo-EM facility, part of the Goeddel Family Technology Sandbox, and the scientific and technical assistance of Dr. Mariusz Matyszewski. We thank G. Ossetchkine, K. Duffin, M. Ganguly, and V. Bradaschia at The Centre for Phenogenomics for their expertise and technical support in immunohistology. We also thank M. Livneh, K. Spencer, and S. Henderson of the Histology and Microscopy Core Facility at The Scripps Research Institute for their technical support in immunostaining and imaging. We acknowledge K. Vanderpool, T. Fassel, and S. Henderson of the Core Microscopy Facility at The Scripps Research Institute for their expert assistance with TEM imaging. We thank A. Saluk, B. Seegers, and B. Monteverde of the Flow Cytometry Core Facility at The Scripps Research Institute for their technical support in flow cytometry. Diffraction data were collected at beamline 12-1 of the Stanford Synchrotron Radiation Lightsource (SSRL). Use of the Stanford Synchrotron Radiation Lightsource and Stanford Linear Accelerator Center (SLAC), National Accelerator Laboratory, is supported by the U.S. Department of Energy, Office of Science, Office of Basic Energy Sciences under Contract No. DE-AC02-76SF00515. The SSRL Structural Molecular Biology Program is supported by the Department of Energy (DOE) Office of Biological and Environmental Research, and by the National Institutes of Health (NIH), National Institute of General Medical Sciences (NIGMS) (P30GM133894). The contents of this publication are solely the responsibility of the authors and do not necessarily represent the official views of NIGMS or NIH. We thank M. Arends for proofreading the manuscript.

## Author contributions

Project design by Y.-Z.L., Y.-N.Z., L.H., and J.Z.; protein design by J.Z.; protein expression and purification by Y.-Z.L., G.W., C.D., and L.H.; glycan analysis by M.N., J.D.A., and M.C.; negative-stain EM and cryo-EM by Y.-Z.L., A.B.W., and J.Z.; x-ray crystallographic analysis by Y.-Z.L., R.L.S., and I.A.W; mouse immunization, lymph node isolation, immunohistology, TEM, flow cytometry, and serum ELISA by Y.-N.Z.; pseudovirus neutralization assays by G.W., K.B.G., C.D., and L.H.; manuscript written by Y.-Z.L., Y.-N.Z., M.N., K.B.G., S.A., L.H., M.C., and J.Z. All authors reviewed and provided feedback on the manuscript.

## Data and material availability

All data needed to evaluate the conclusions of this study are available in the main text and Supplementary Materials. Structural data obtained from X-ray crystallography and cryo-EM have been deposited in the Protein Data Bank (PDB, https://www.rcsb.org/) under accession codes 9N8E and 9N8F, respectively. Cryo-EM density maps have been deposited in the Electron Microscopy Data Bank (EMDB, https://www.ebi.ac.uk/emdb/) under accession code EMD-49127. Mass spectrometry data have been deposited in the MassIVE server (https://massive.ucsd.edu) under accession code MassIVE MSV000097145. All data supporting the findings of this study are available within the article and its Supplementary Information. Source data are provided with this paper.

---

**Fig. S1.**
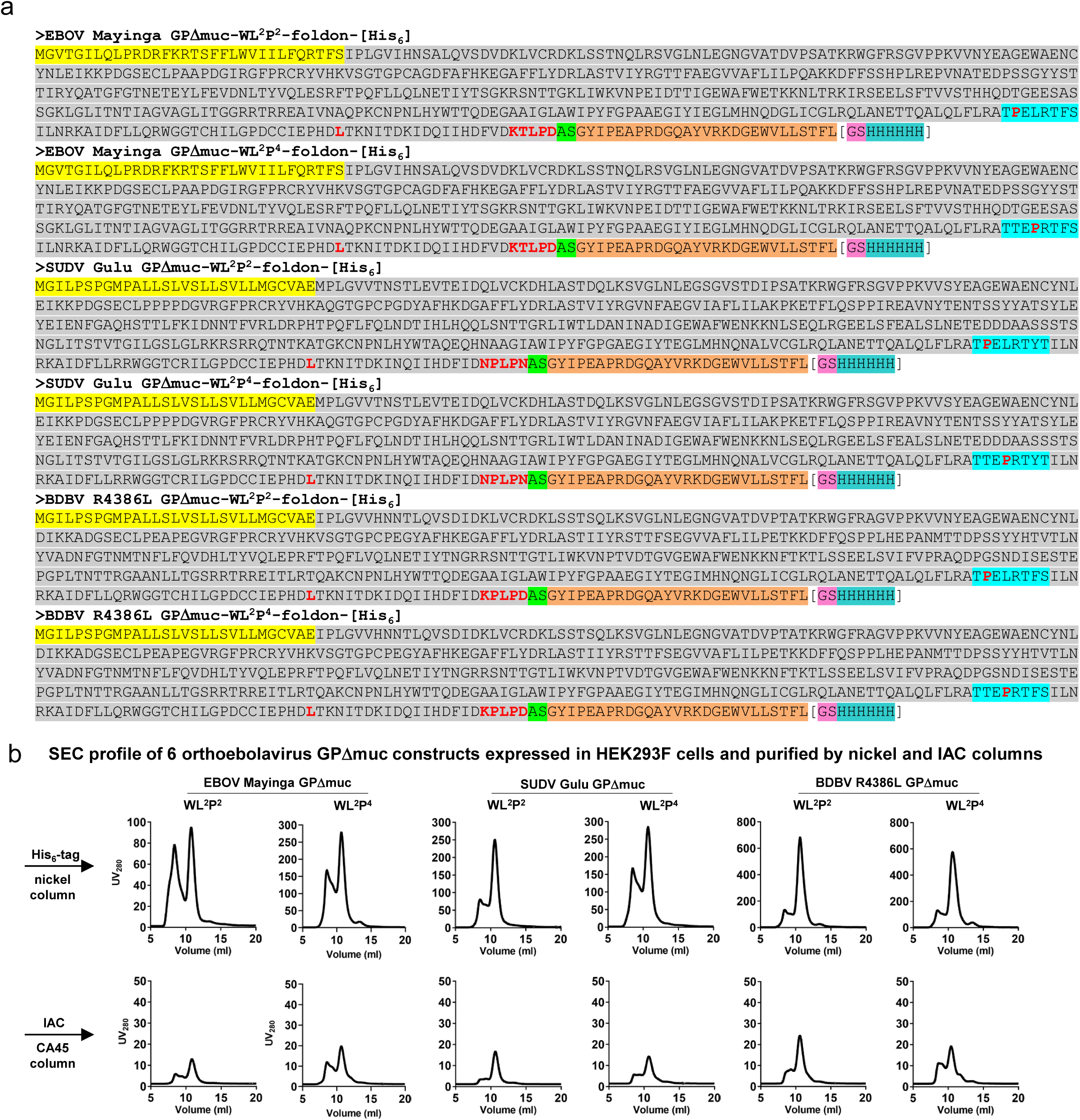
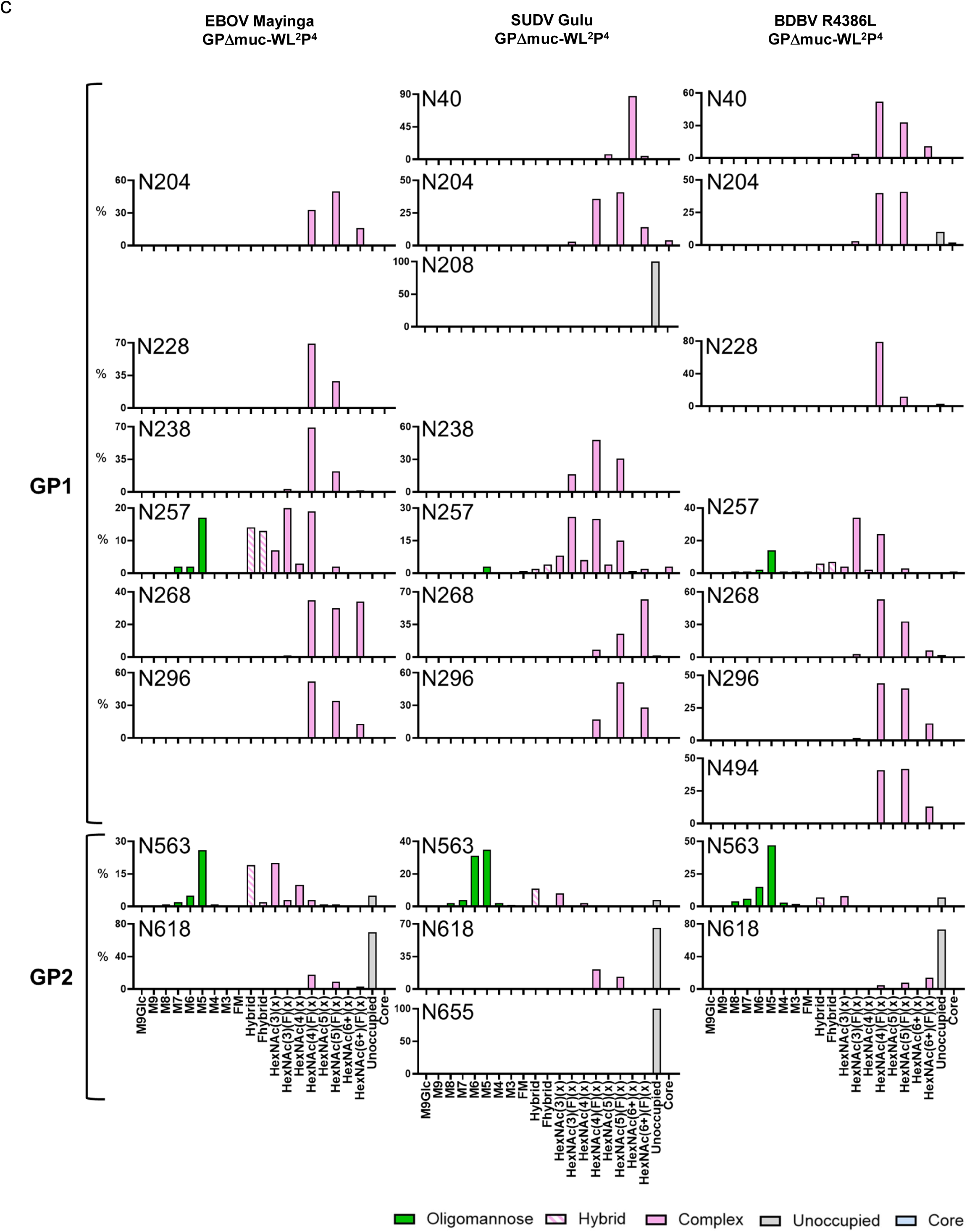
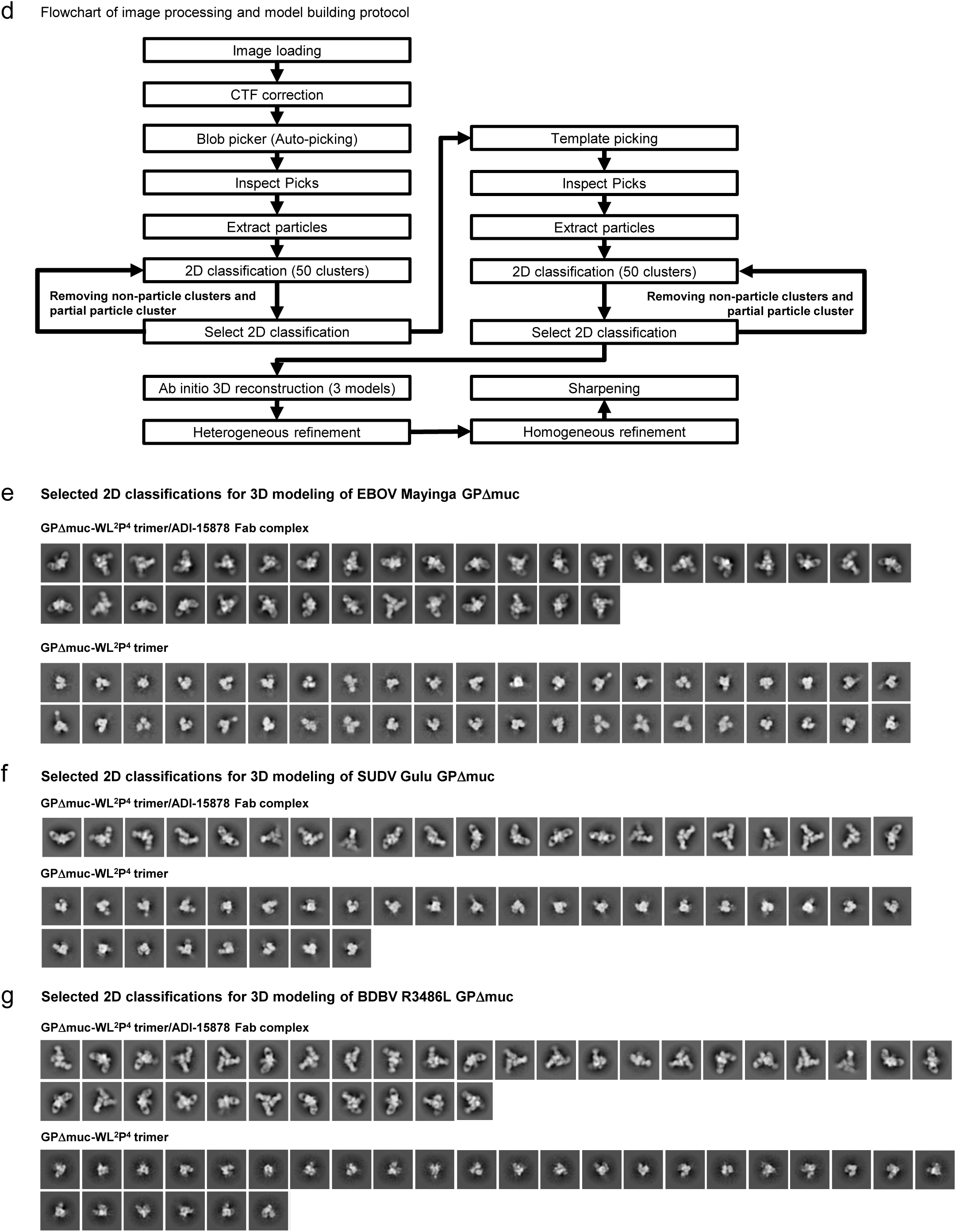
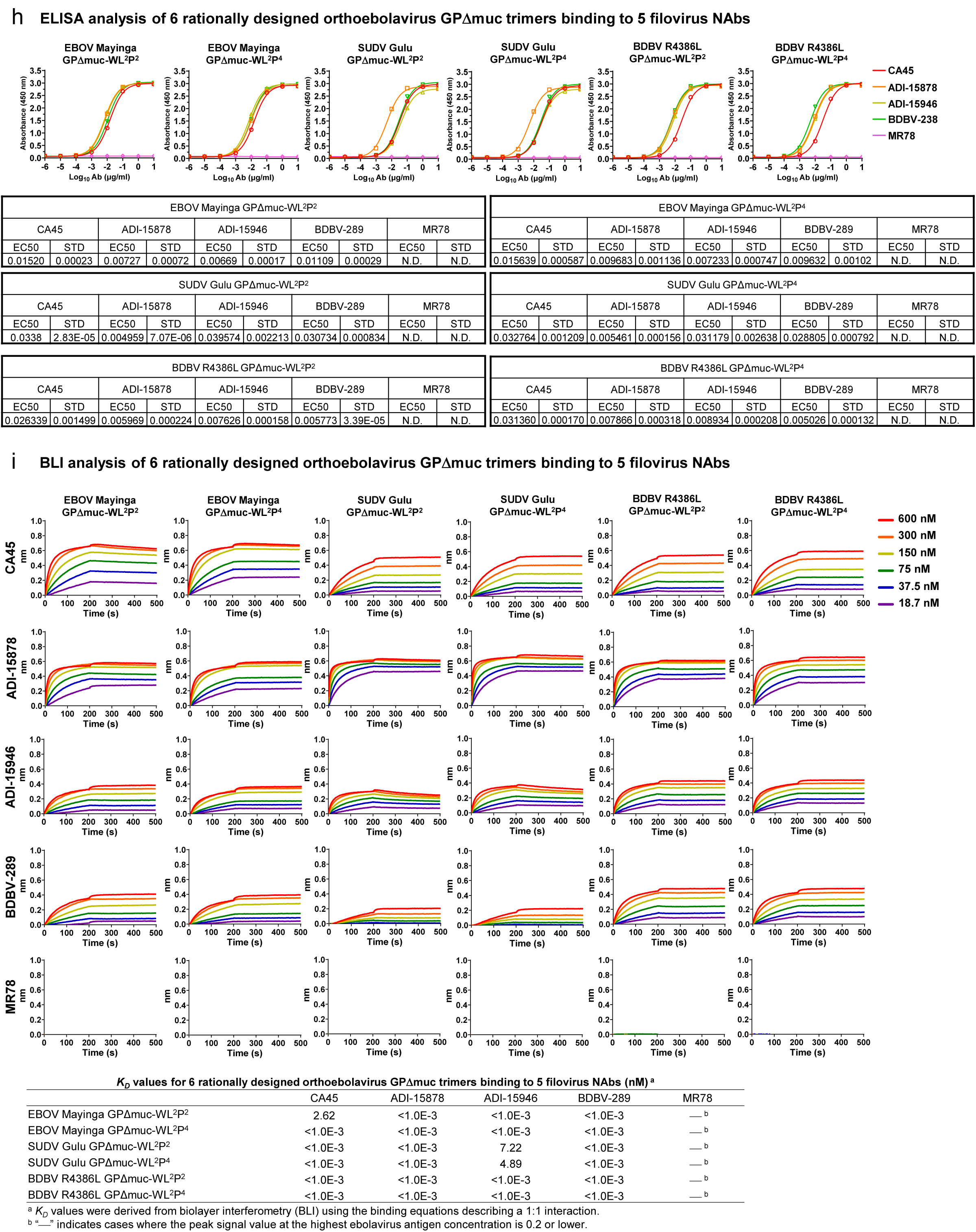
Construct design and in vitro characterization of orthoebolavirus GPΔmuc trimers. (**a**) Amino acid sequences of EBOV Mayinga, SUDV Gulu, and BDBV R3486L GPΔmuc-WL^2^P^x^-foldon-[His_6_] constructs containing either the P^2^ or P^4^ mutation. Signal peptide, GP, restriction site (AS), foldon, linker (GS), and His_6_ tag are highlighted in yellow, grey, green, orange, pink, and teal, respectively. The WL^2^P^x^ mutations are shown in red. Note: Foldon is a C-terminal trimerization motif used in all orthoebolavirus GPΔmuc constructs and therefore will not be included in the construct names (except here in the sequence definition) to avoid redundancy. The GS linker and His_6_ tag are enclosed in [ ] to indicate that they are included only in a subset of constructs used for evaluating nickel-based purification. (**b**) SEC profiles of EBOV Mayinga, SUDV Gulu, and BDBV R3486L GPΔmuc proteins produced in 165 ml HEK293F cultures and purified using either a nickel affinity column or a CA45 immunoaffinity column. (**c**) Compositional site-specific glycan analysis of three orthoebolavirus GPΔmuc-WL^2^P^4^ trimers. The graphs summarize quantitative mass spectrometric analysis of the glycan population present at individual N-linked glycosylation sites simplified into categories of glycans. The oligomannose-type glycan series (M9 to M5; Man_9_GlcNAc_2_ to Man_5_GlcNAc_2_) is colored green, afucosylated and fucosylated hybrid-type glycans (hybrid and F hybrid) are dashed pink, and complex glycans are grouped according to the number of antennae and presence of core fucosylation and are colored pink. Unoccupancy of an N-linked glycan site is represented in gray. Glycan sites that could not be determined are denoted as “N.D.”. (**d**) Flowchart illustrating image processing, 2D classification, and 3D reconstruction of negative stain EM (nsEM) data for EBOV, SUDV, and BDBV GPΔmuc trimers and their complexes with ADI-15878 Fab, using CryoSPARC. (**e**)-(**f**) Representative 2D classification images of EBOV, SUDV, and BDBV GPΔmuc-WL^2^P^4^ trimers and their complexes with ADI-15878 Fab. (**g**) ELISA analysis of EBOV, SUDV, and BDBV GPΔmuc trimers binding to 5 filovirus NAbs in IgG form. Briefly, each well was coated with 0.1 μg of the appropriate antigen, and IgG was diluted in a 10-fold dilution series from a starting concentration of 10 μg/ml for all tested antibodies. Error bars represent the difference between duplicate measurements at each concentration for each sample. (**h**) BLI analysis of EBOV, SUDV, and BDBV GPΔmuc trimers binding to 5 filovirus NAbs in IgG form. Sensorgrams were obtained from an Octet RED96 instrument using AHC biosensors. A two-fold concentration gradient of antigen, starting at 600 nM, was used in a dilution series of six. *K_D_* values derived from a 1:1 fitting model are summarized in a table.

**Fig. S2.**
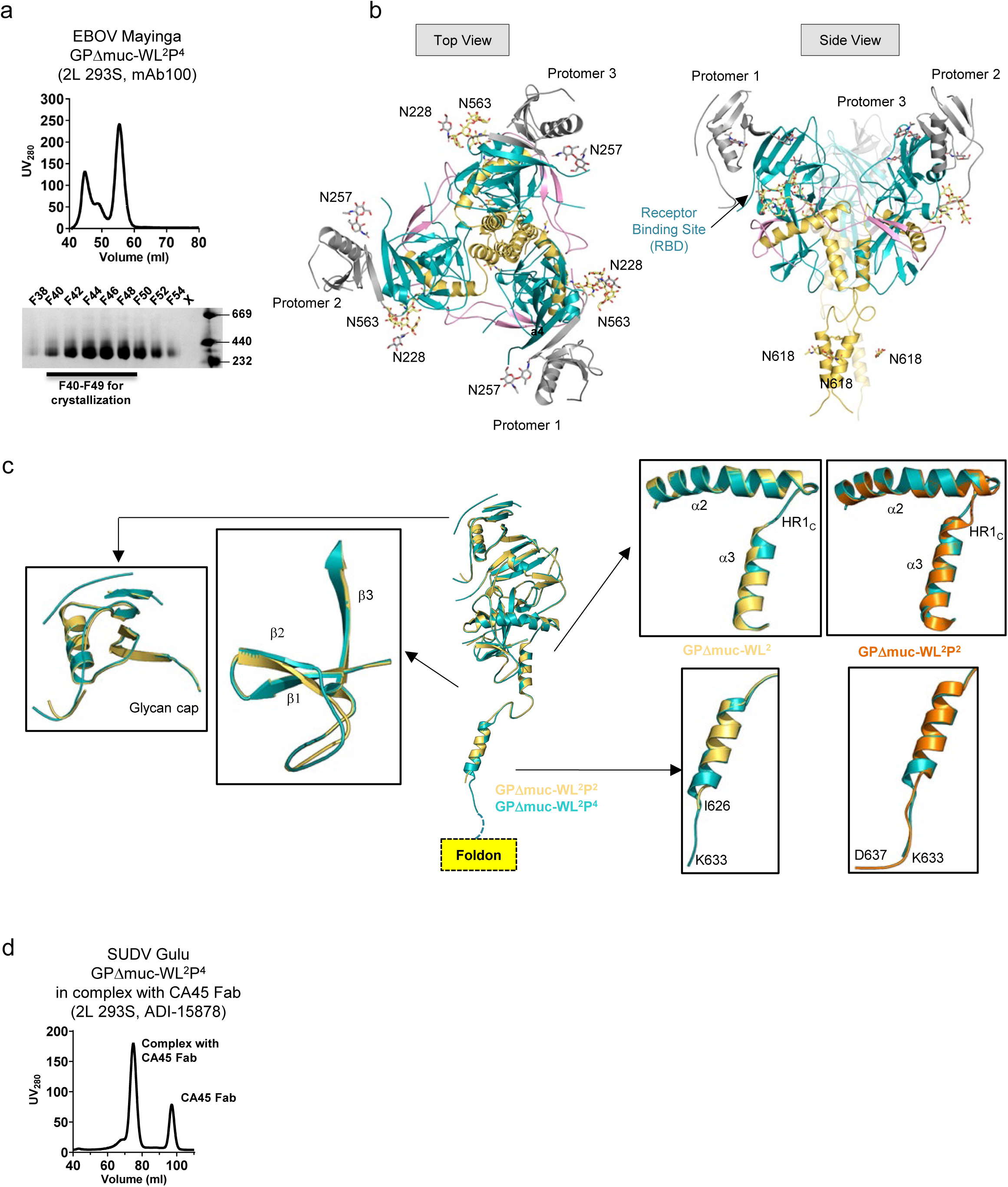
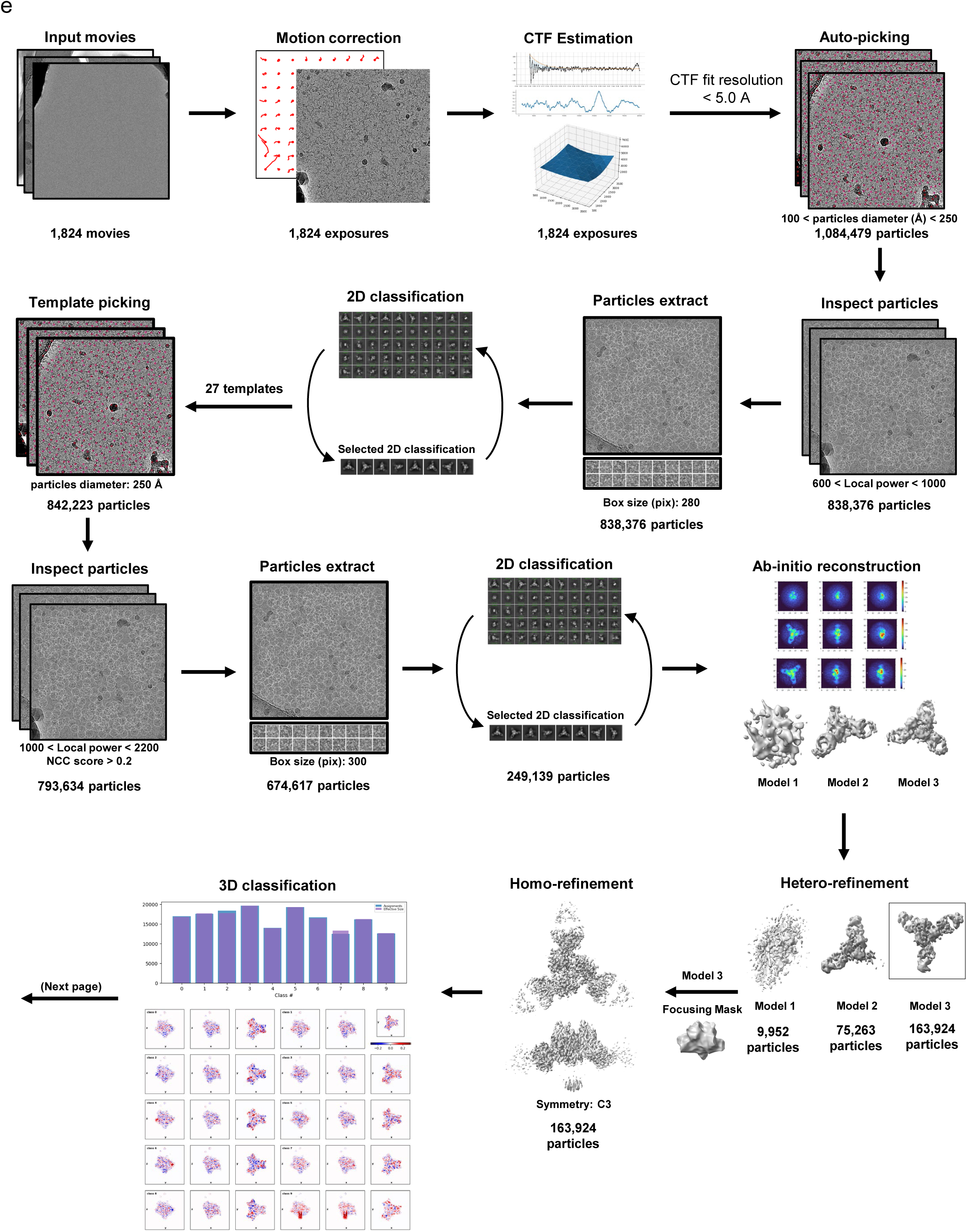
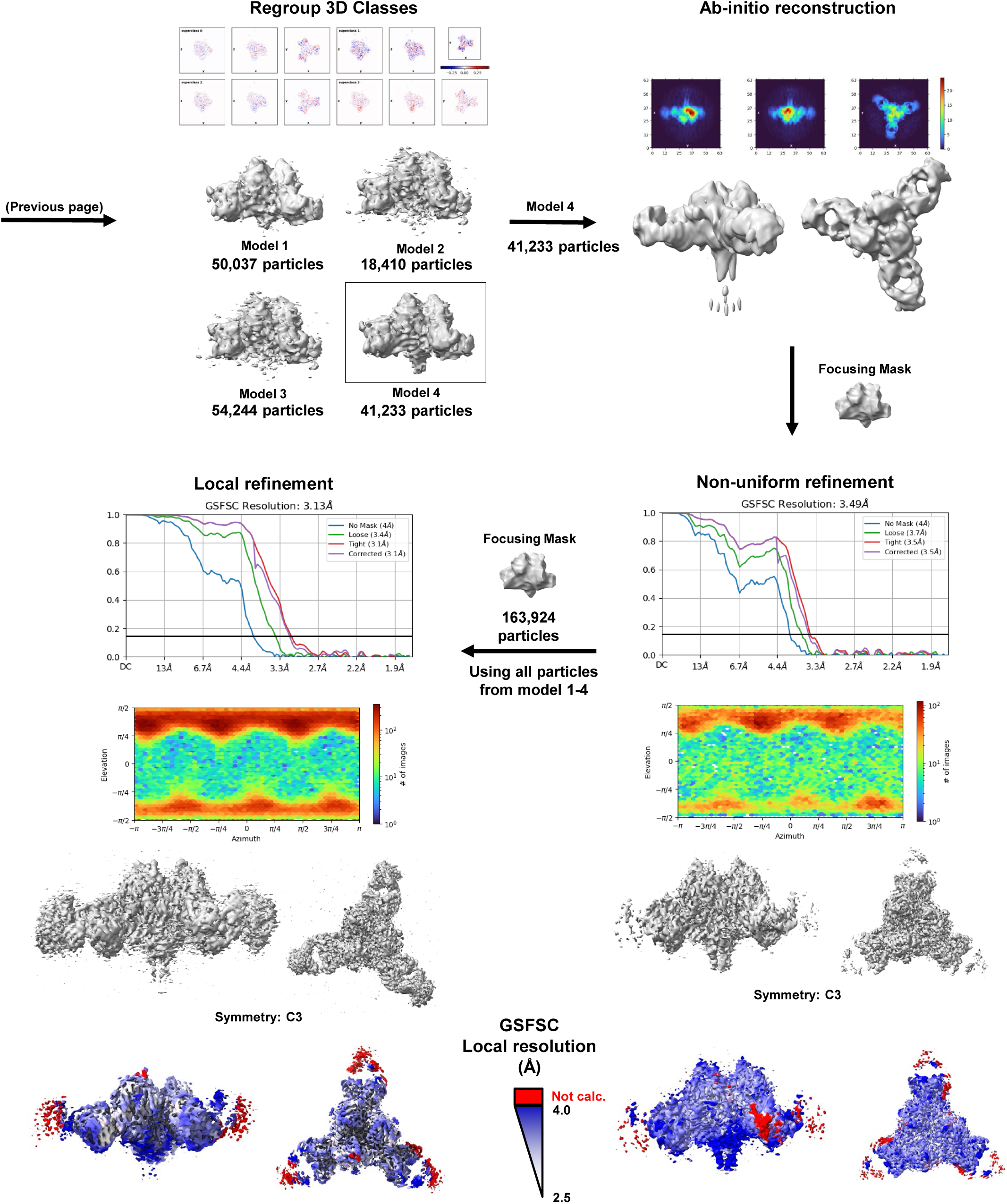
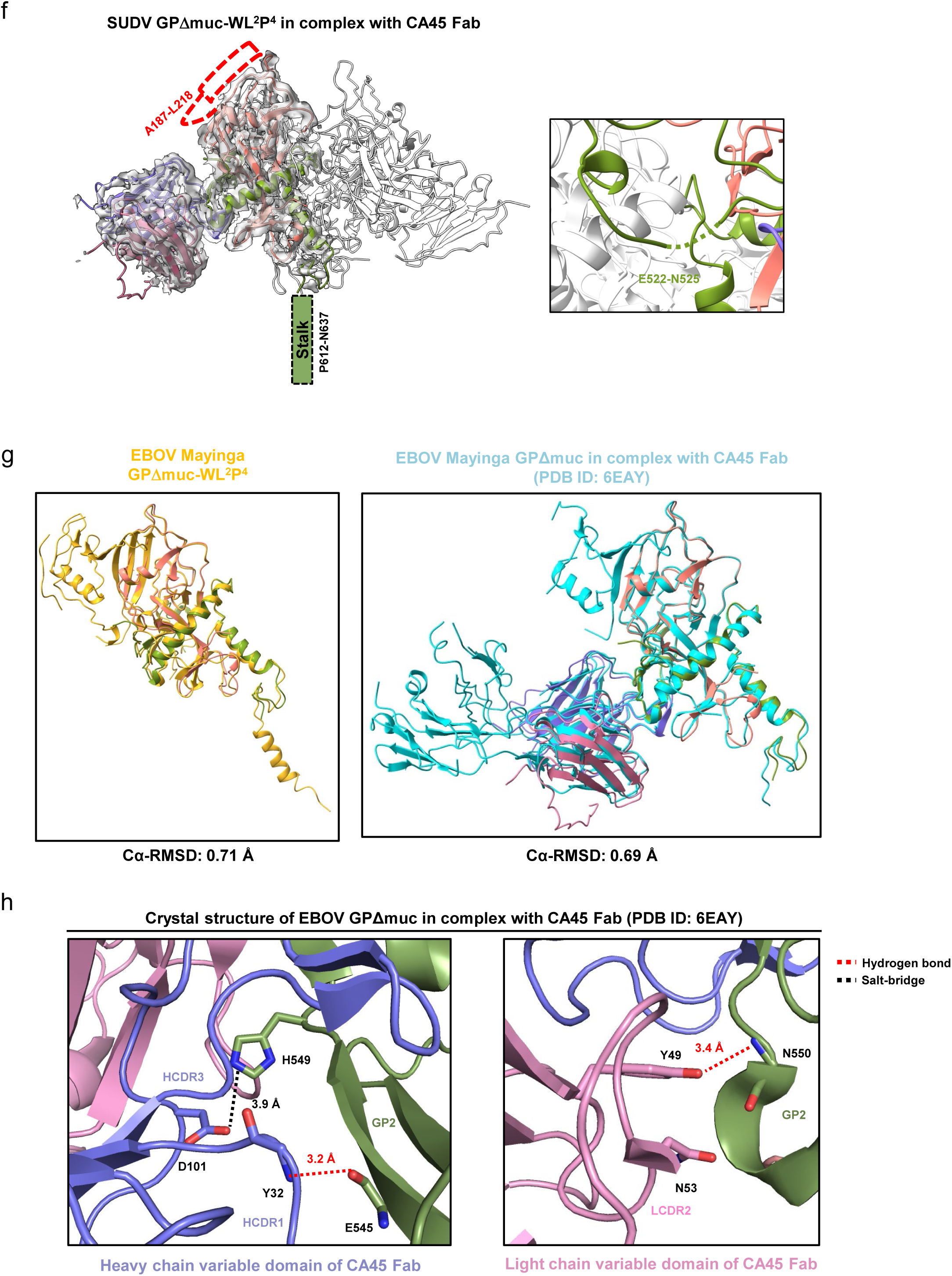
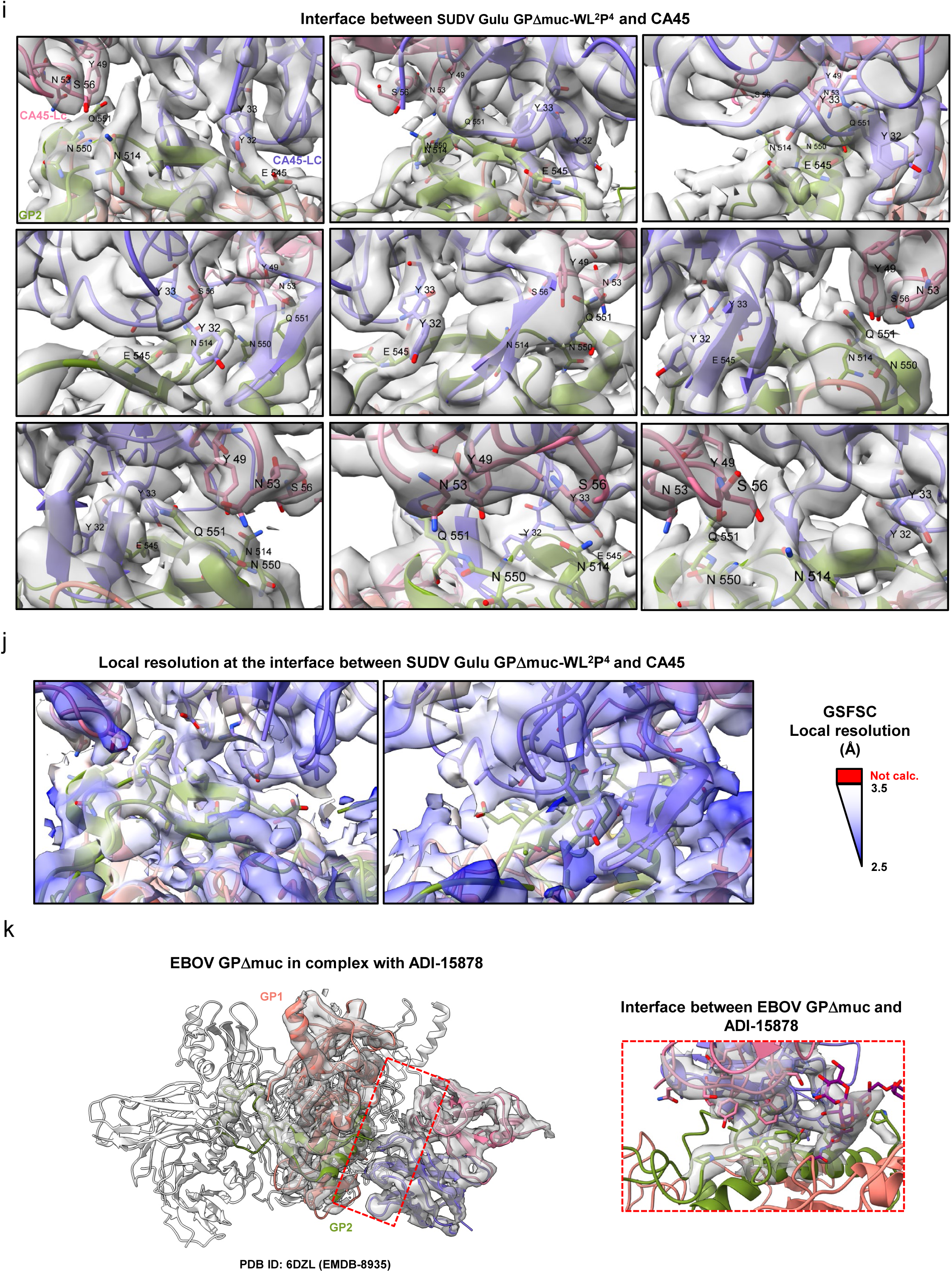
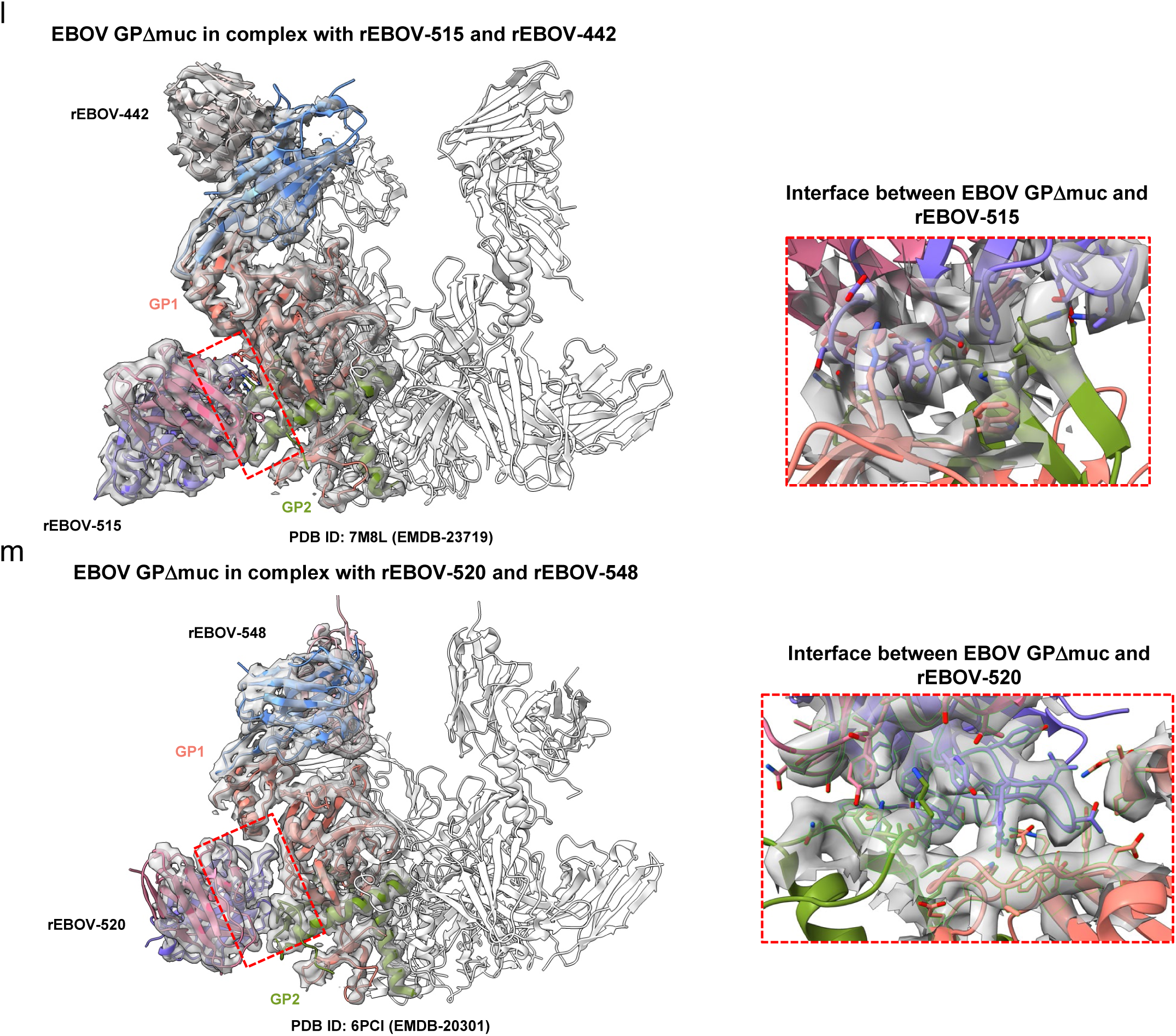
Structural characterization of ligand-free EBOV GPΔmuc-WL^2^P^4^ and CA45 Fab-bound SUDV GPΔmuc-WL^2^P^4^ trimers. (**a**) SEC profile of EBOV GPΔmuc-WL^2^P^4^ trimer expressed in 2L HEK293S cells and BN-PAGE of ADI-15878/SEC-purified EBOV GPΔmuc-WL^2^P^4^ trimer. Fractions F40 to F49 were used for crystallization screening. The SEC profile was generated from a HiLoad Superdex 200 16/600 PG column (Cytiva). (**b**) Ribbon representation of EBOV GPΔmuc-WL^2^P^4^ trimer structure (Left: top view; Right: side view). The structure is colored as follows: glycan cap (gray), RBS (cyan), GP2 (yellow), and IFL (pink). *N*-linked glycans at N228, N257, N563, and N618 are shown as color-coded (by atom type) sticks. (**c**) Structural comparison of GPΔmuc-WL^2^P^4^ and previously reported GPΔmuc-WL^2^ and GPΔmuc-WL^2^P^2^ trimers. The ribbon models of three GPΔmuc trimers containing WL^2^P^4^, WL^2^, and WL^2^P^2^ mutations are shown in cyan, gold, and orange, respectively. Regions that are critical to the GP structure and exhibit appreciable differences in structure are superposed and shown as enlarged insects. (**d**) SEC profile of ADI-15878/SEC-purified SUDV GPΔmuc-WL^2^P^4^ trimer expressed in 2L HEK293S cells and complexed with ADI-15878 Fab. The SEC profile was generated from a Superose 6 10/300 increase column. (**e**) Schematic representation of image processing, 2D classification, and 3D reconstruction of cryo-EM data obtained for SUDV GPΔmuc-WL^2^P^4^ trimer in complex with ADI-15878 Fab using CryoSPARC. (**f**) Regions not modeled in the GP–CA45 complex structure due to limited density resolution. (**g**) Left: cryo-EM structure of SUDV GPΔmuc-WL^2^P^4^ superposed with the crystal structure of EBOV GPΔmuc-WL^2^P^4^ for comparison at the monomer level. The Cα root mean square deviation (Cα-RMSD) is 0.66 Å for 154 matching Cα atoms. Right: cryo-EM structure of SUDV GPΔmuc-WL^2^P^4^ in complex with ADI-15878 Fab superposed with the previously reported crystal structure of EBOV GPΔmuc in complex with CA45 Fab (PDB ID: 6EAY). Cα-RMSD is 0.63 Å for 149 matching Cα atoms. (**h**) Structural details of the GP-CA45 interface as illustrated in the crystal structure of EBOV GPΔmuc in complex with CA45 Fab (PDB ID: 6EAY). Hydrogen bonds are shown as red dashed lines, and the salt bridge is indicated with a black dashed line. Distances are labeled. (**i**) Snapshots of the GP-CA45 interface from different views with the density map shown. Interacting residues are displayed in ribbon representation. (**j**) Local resolution map overlaid with the built model at the GP-CA45 interface. (**k**) Published cryo-EM structure of EBOV GPΔmuc in complex with ADI-15878 (PDB ID: 6DZL; EMDB-8935), highlighting structural details at the interface. (**l**) Published cryo-EM structure of EBOV GPΔmuc in complex with rEBOV-515 and rEBOV-442 (PDB ID: 7M8L; EMDB-23719), with interface details shown. (**m**) Published cryo-EM structure of EBOV GPΔmuc in complex with rEBOV-520 and rEBOV-548 (PDB ID: 6PCI; EMDB-20301), highlighting interface details.

**Fig. S3.**
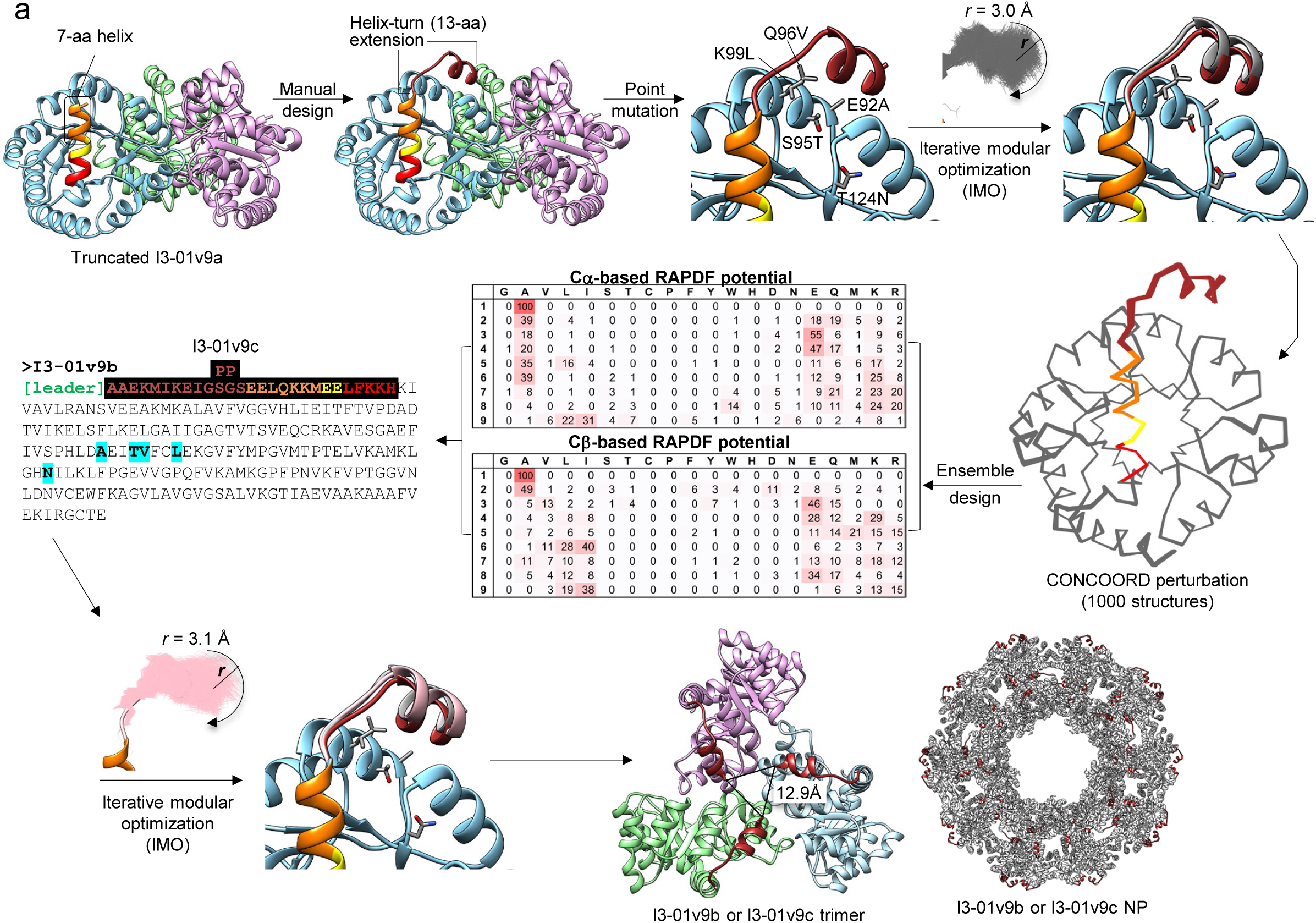
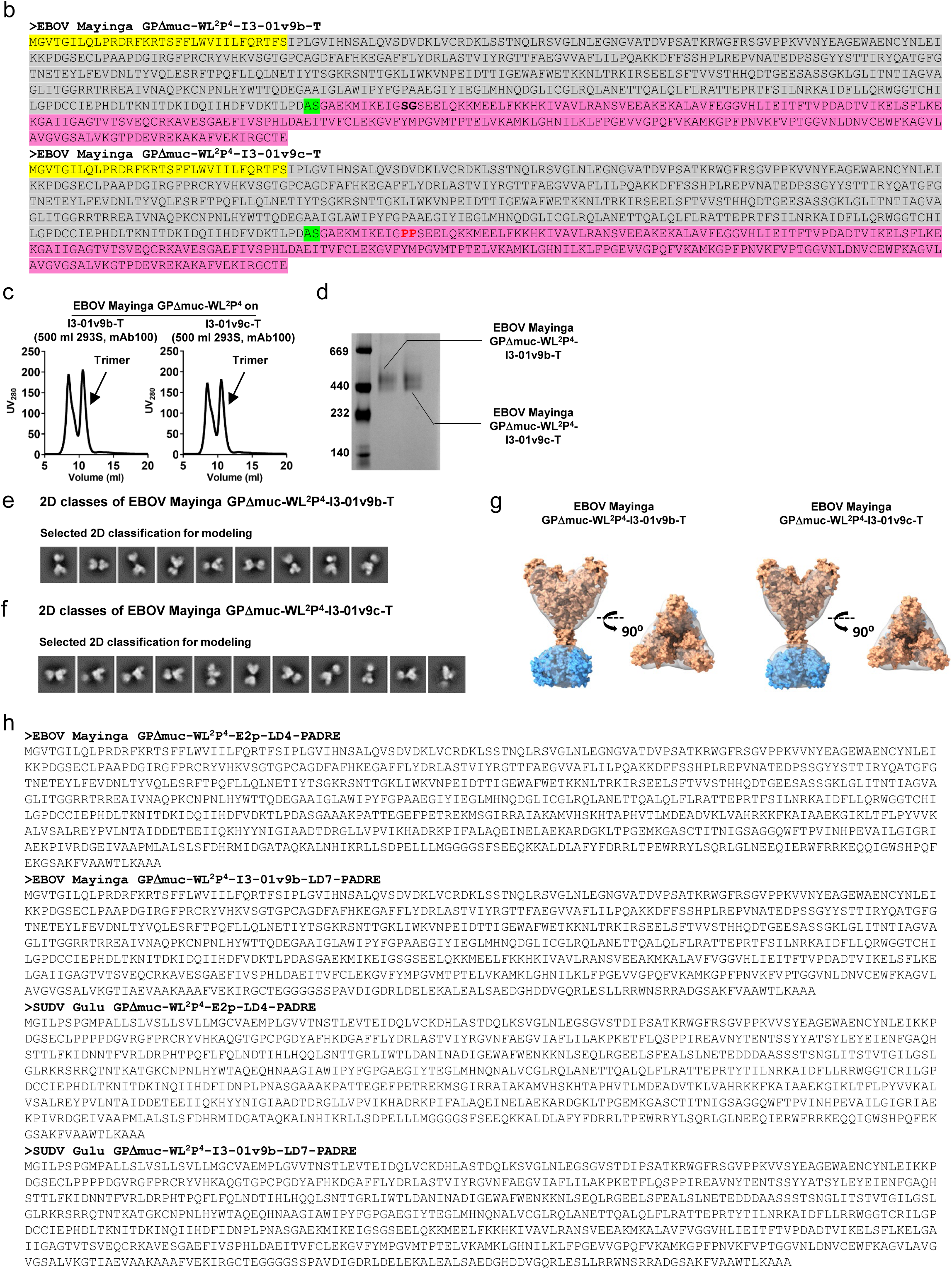
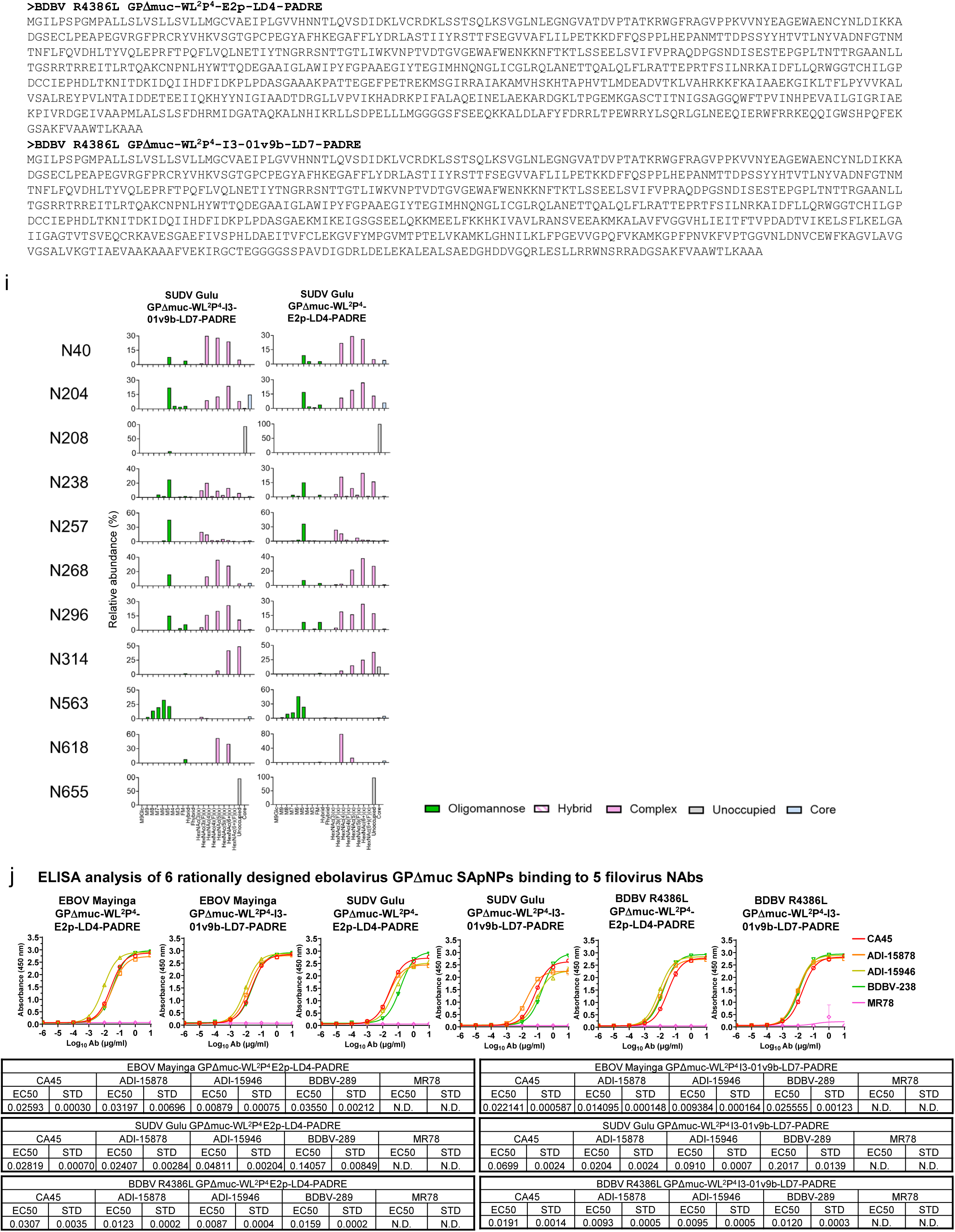
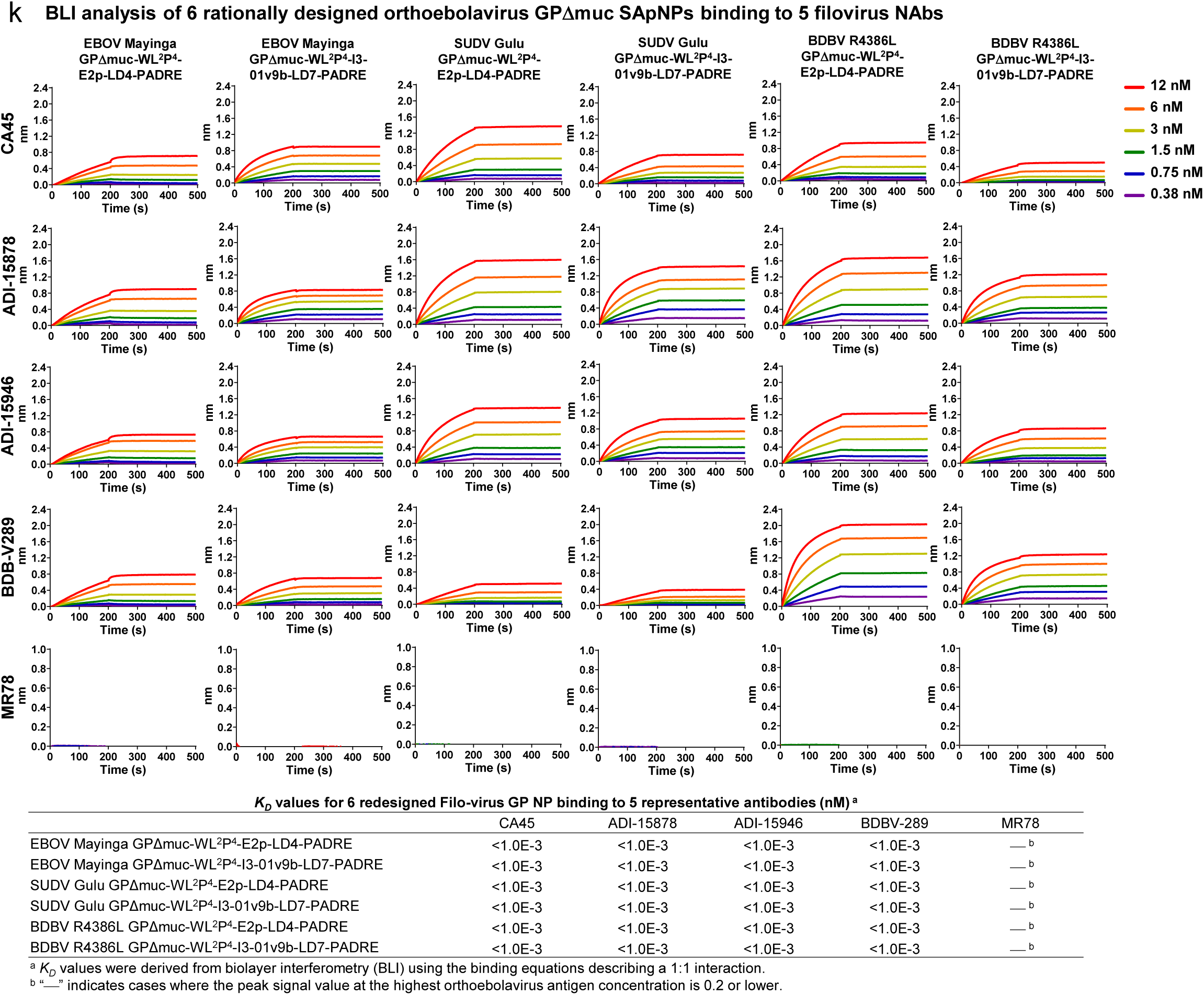
Rational design of I3-01v9b/c and in vitro characterization of orthoebolavirus GPΔmuc-presenting SApNPs. (**a**) Schematic representation of computational design of I3-01v9b/c. Top left: structural model of I3-01v9a with the 11-aa N-terminal helix truncated to 7 aa; Top middle left: a 13-aa helix-turn fragment (9-aa helix + 4-aa turn, all alanine) was fused to the 7-aa N-terminal helix of the truncated I3-01v9a in such a way the new 9-aa N-terminal helix from the fused fragment would pack within the groove of two helices that are part of the I3-01 core; Top middle right: four mutations, E92A, S95T, Q96V, and K99L, were introduced to the I3-01 core helices to remove their steric clashes with the new N-terminal helix; Top right: iterative modular optimization (IMO) of the helix-turn backbone, with a radius of 3.0 Å measured for the first amino acid at the N-terminus; Middle right: 1000 slightly perturbed backbone conformations generated by CONCOORD, a torsion-space sampling program; Middle middle: results from an ensemble-based protein design program used to predict the optimal sequence for the 9-aa N-terminal helix within the fragment using Cα and Cβ-based RAPDF potentials. The 4-aa turn was set to “GSGS”; Middle left: sequence of I3-01v9b designed by combining data from both RAPDF potentials, with the difference between I3-01v9b and I3-01v9c (the “PP” mutation) labeled on the sequence; Bottom left: further backbone relaxation using IMO with a minimum perturbation angle; Bottom right: a structural model of the trimeric I3-01v9b/c, termed I3-01v9b/c-T, in which the N-termini form a triangle of 12.9 Å, and a structural model of fully assembled I3-01v9b/c NP. (**b**) Amino acids sequences of EBOV GPΔmuc-WL2P4-I3-01v9b/c-T, which are EBOV Mayinga GPΔmuc-WL^2^P^4^ anchored to the trimeric I3-01v9b/c-T scaffolds. Signal peptide, GP, restriction site (AS), and I3-01v9b/c-T are highlighted in yellow, grey, green and pink, respectively. The double proline mutation in the N-terminus of I3-01v9c-T that differs from I3-01v9b-T is highlighted in red. (**c**) SEC profiles of EBOV GPΔmuc-WL2P4-I3-01v9b/c-T expressed in 500 ml HEK293S cells and purified using an ADI-15878 column. (**d**) BN-PAGE of ADI-15878/SEC-purified EBOV GPΔmuc-WL2P4-I3-01v9b/c-T. (**e**) and (**f**) Representative 2D classification images of ADI-15878/SEC-purified EBOV GPΔmuc-WL2P4-I3-01v9b/c-T trimers. (**g**) 3D reconstructions of ADI-15878/SEC-purified EBOV GPΔmuc-WL2P4-I3-01v9b/c-T trimers. Crystal structures of EBOV GPΔmuc-WL^2^P^2^ (PDB ID:7JPH) and the bacterial enzyme from which I3-01 was derived (PDB ID:1VLW) were fitted into the model and shown in orange and blue, respectively. (**h**) Amino acids sequences of EBOV, SUDV, and BDBV GPΔmuc-WL^2^P^4^-E2P-LD4-PADRE and -I3-01v9b-LD7-PADRE SApNP constructs. LD stands for locking domain (LD), which is fused to the C-terminus of the NP backbone and forms an inner layer to stabilize the NP shell from the inside. PADRE is a helper T-cell epitope and forms a hydrophobic core at the center of the assembled NP to further stabilize the NP structure and to induce a strong T-help response upon vaccination. (**i**) Compositional site-specific glycan analysis of SUDV GPΔmuc-WL^2^P^4^-E2P-LD4-PADRE and I3-01v9b-LD7-PADRE SApNPs, as in **Fig. S1c**. (**j**) ELISA analysis of EBOV, SUDV, and BDBV GPΔmuc-WL^2^P^4^-E2P-LD4-PADRE and I3-01v9b-LD7-PADRE SApNPs binding to 5 filovirus NAbs, as in **Fig. S1h**. (**k**) BLI analysis of EBOV, SUDV, and BDBV GPΔmuc-WL^2^P^4^-E2P-LD4-PADRE and I3-01v9b-LD7-PADRE SApNPs binding to 5 filovirus NAbs. A two-fold concentration gradient of antigen, starting at 12 nM, was used in a dilution series of six. *K_D_* values derived from a 1:1 fitting model are summarized in a table.

**Fig. S4.**
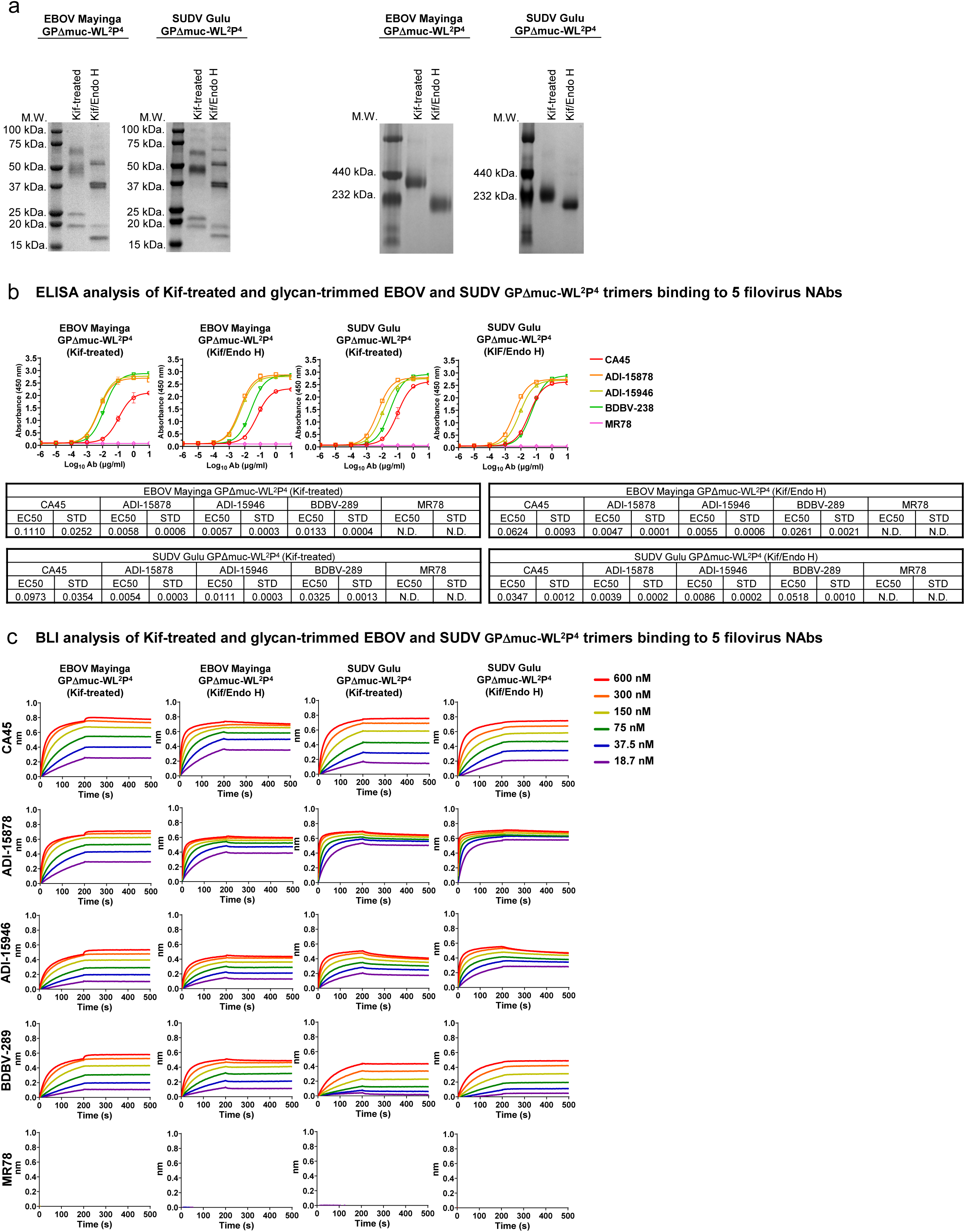
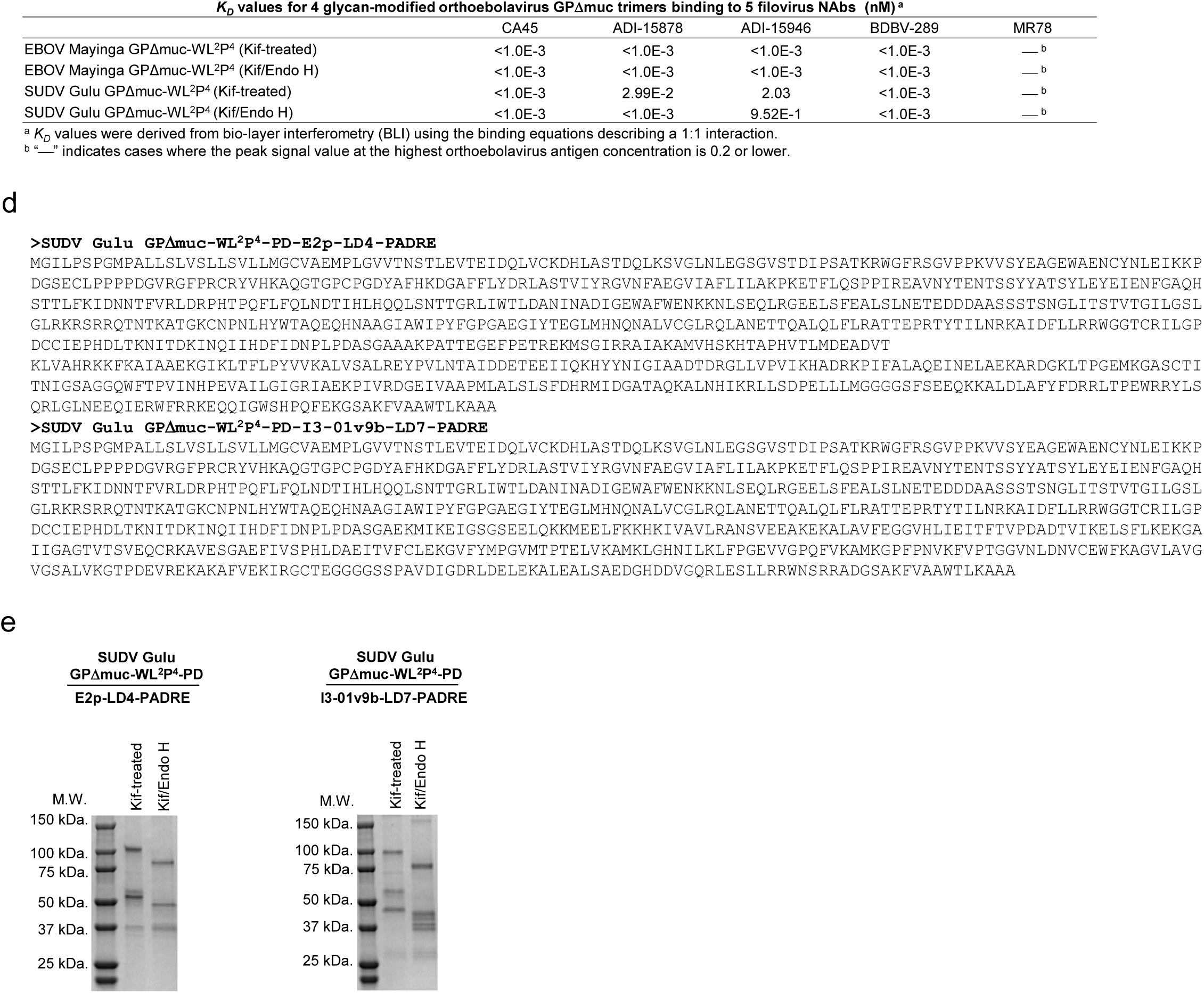
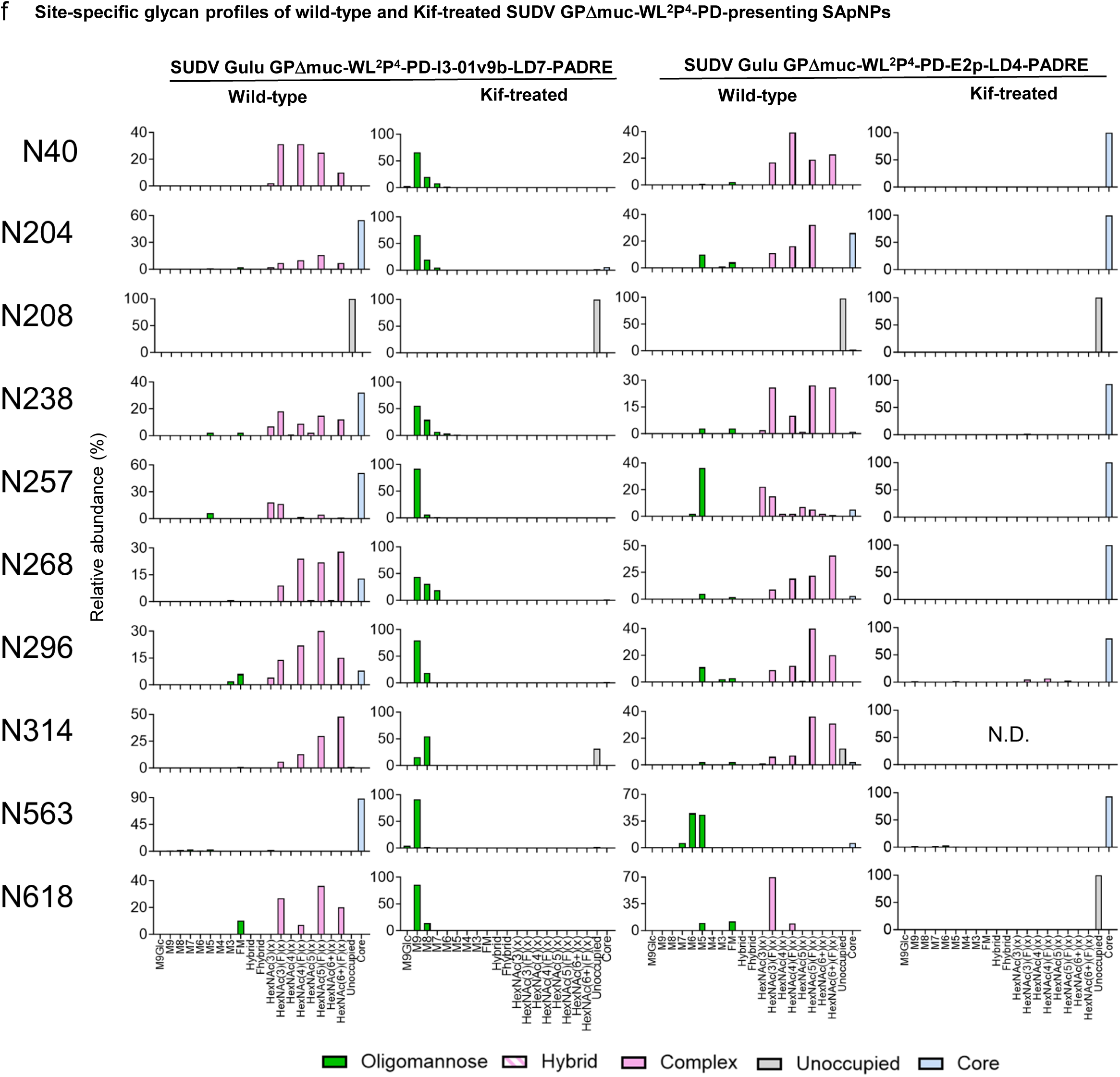
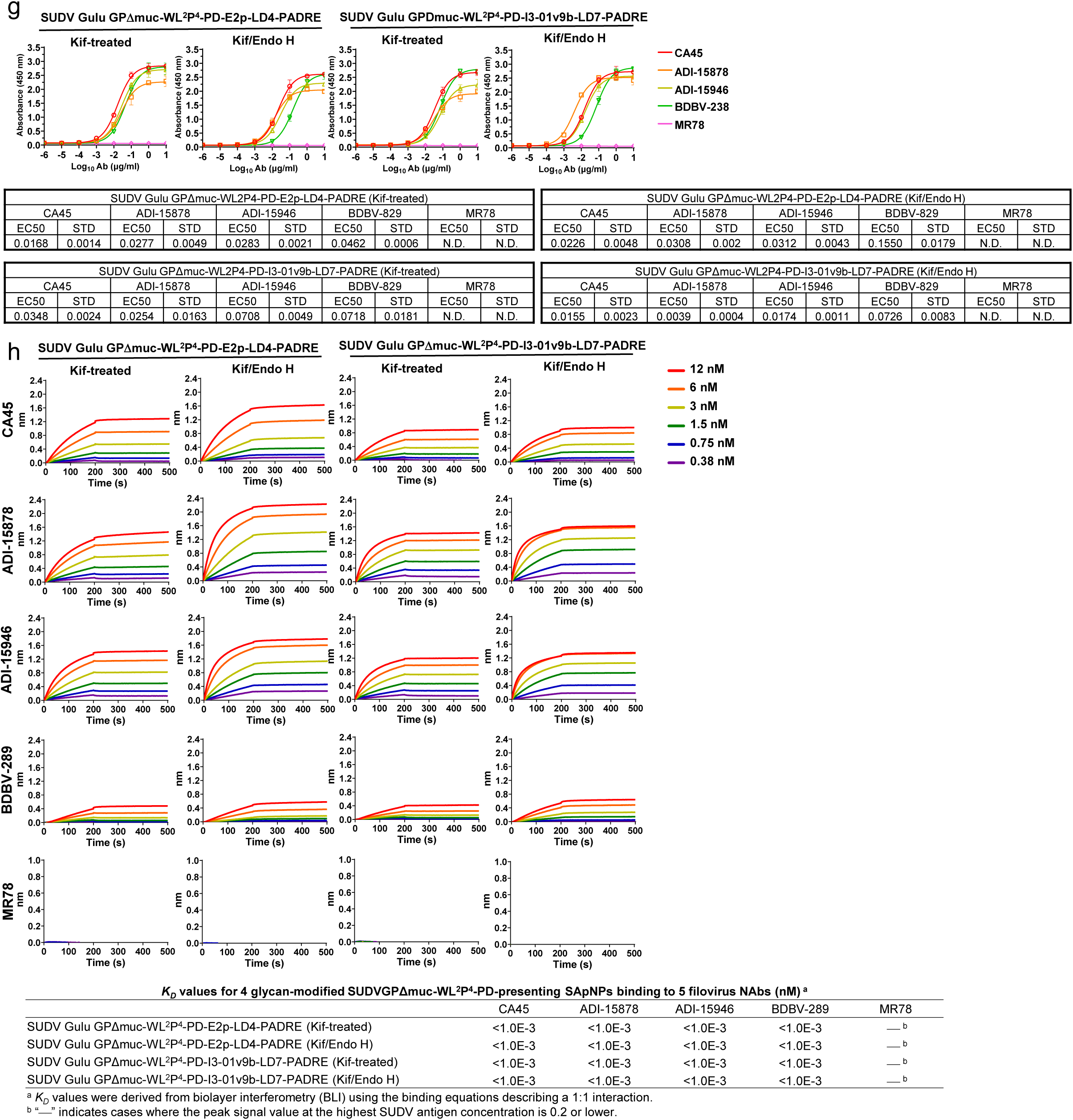
In vitro characterization for glycan-modified orthoebolavirus GPΔmuc trimers and SApNPs. (**a**) SDS-PAGE and BN-PAGE of glycan-modified EBOV Mayinga and SUDV Gulu GPΔmuc-WL^2^P^4^ trimers. They were either expressed in the presence of kifunensine (Kif-treated) to obtain oligomannose-only glycans or further trimmed using endo H (Kif/Endo H) to obtain a mono-layer of GlcNAc stumps. (**b**) ELISA analysis of glycan-modified EBOV Mayinga and SUDV Gulu GPΔmuc-WL^2^P^4^ trimers binding to 5 filovirus NAbs in the IgG form, as in **Fig S1h.** (**c**) BLI analysis of glycan-modified EBOV Mayinga and SUDV Gulu GPΔmuc-WL^2^P^4^ trimers binding to 5 filovirus NAbs in the IgG form, as in **Fig S1i**. (**d**) Construct sequences for “multilayered” E2p (E2p-LD4-PADRE) and I3-01v9b (I3-0v9b-LD7-PADRE) SApNPs presenting 20 copies of a SUDV Gulu GPΔmuc-WL^2^P^4^-PD trimer, which contains a mutation (N637D) at the C-terminus to remove a potential glycosylation site between GP and NP. (**e**) SDS-PAGE of 3 glycan-modified SApNPs corresponding to EBOV Mayinga GPΔmuc-WL^2^P^4^ on the E2p-LD4-PADRE scaffold and SUDV Gulu GPΔmuc-WL^2^P^4^-PD on the E2p-LD4-PADRE and I3-01v9b-LD7-PADRE scaffolds. (**f**) Compositional site-specific glycan analysis of SUDV GPΔmuc-WL^2^P^4^-PD-presenting E2p and I3-01v9b SApNPs, as in **Fig. S1c**. (**g**) ELISA analysis of glycan-modified SUDV Gulu GPΔmuc-WL2P4-PD-presenting E2P and I3-01v9b SApNPs binding to 5 filovirus NAbs in the IgG form, as in **Fig S1h**. (**h**) BLI analysis of glycan-modified SUDV Gulu GPΔmuc-WL^2^P^4^-PD-presenting E2P and I3-01v9b SApNPs binding to 5 filovirus NAbs in the IgG form, as in **Fig S3k**.

**Fig. S5.**
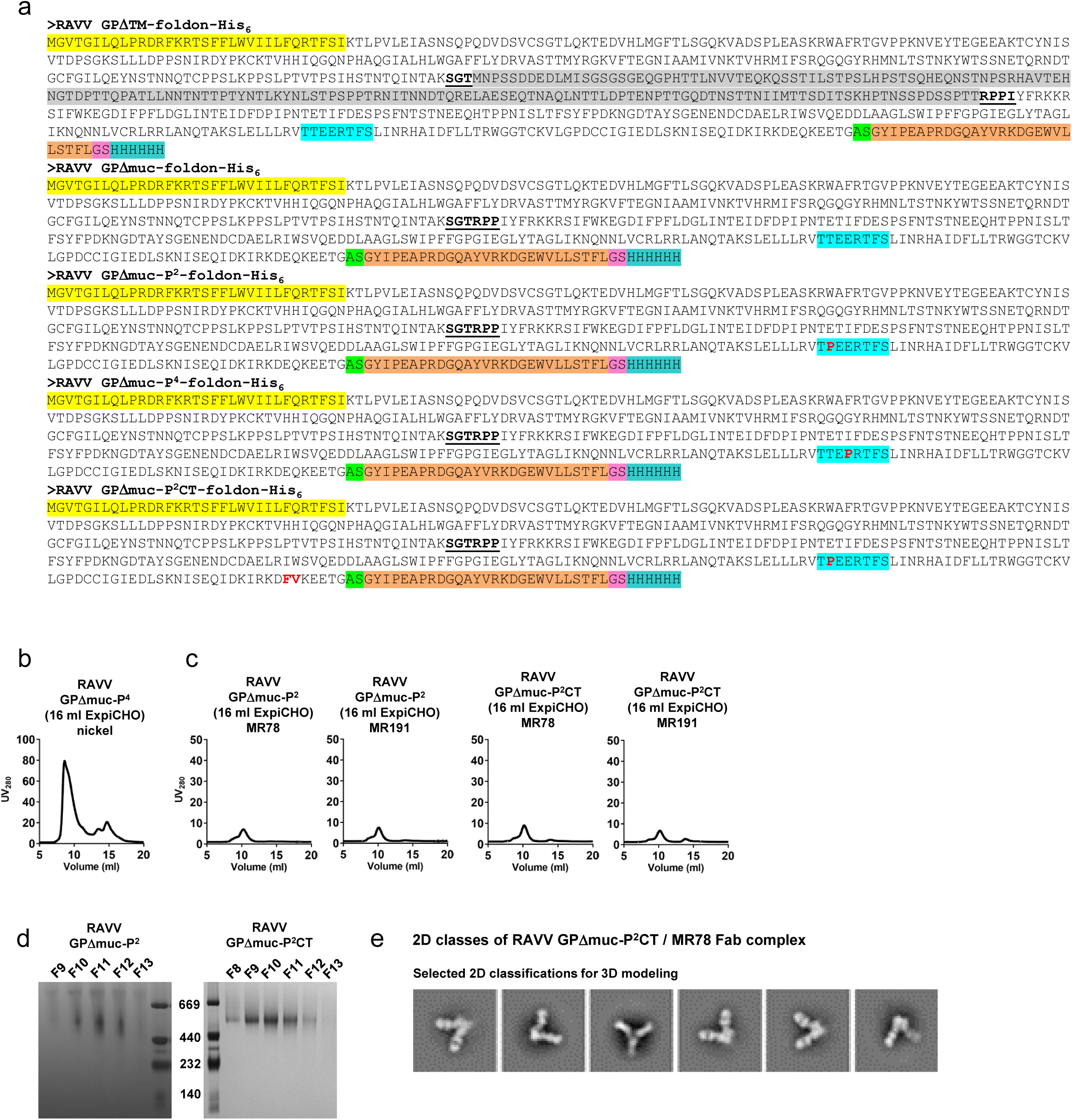
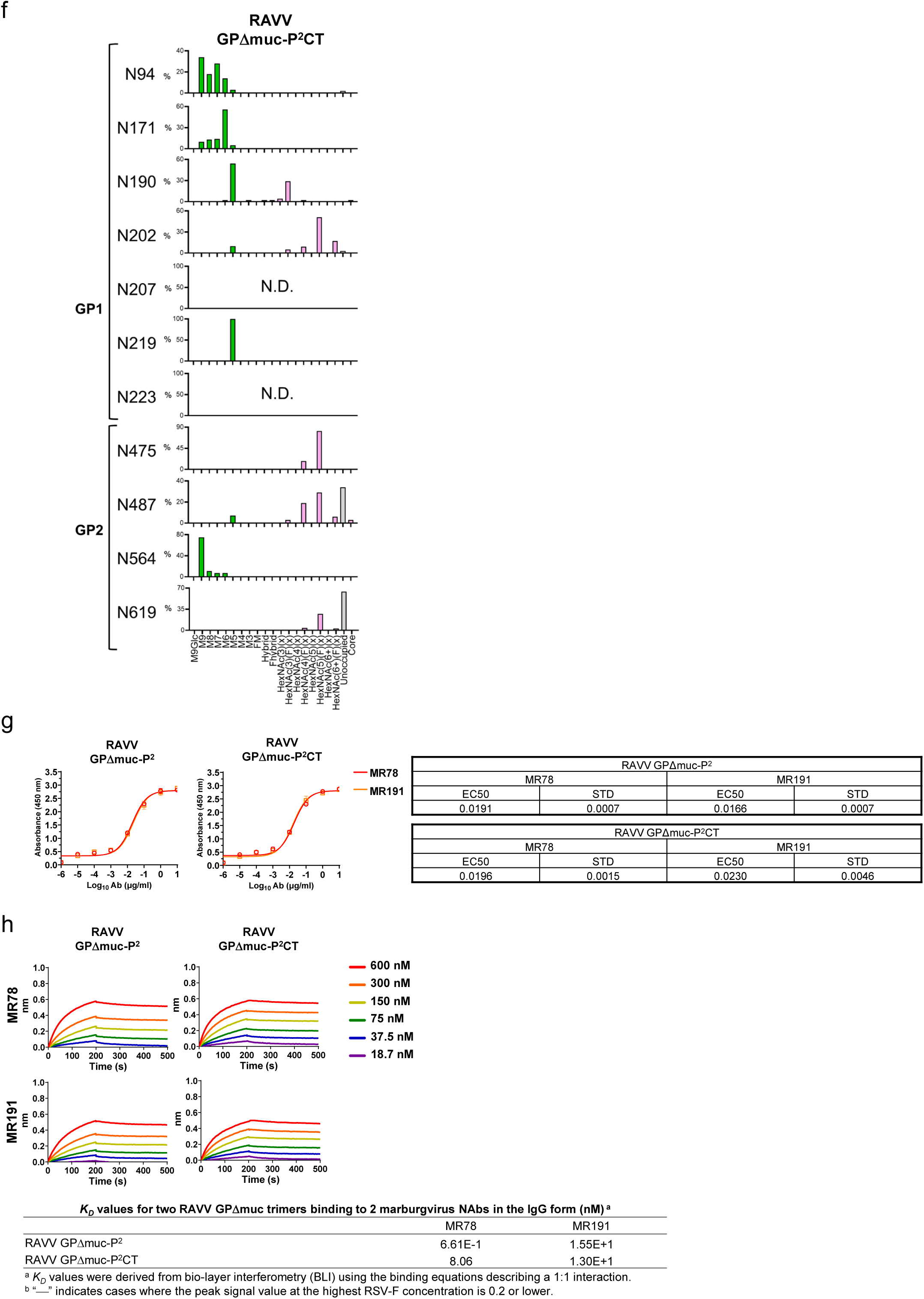
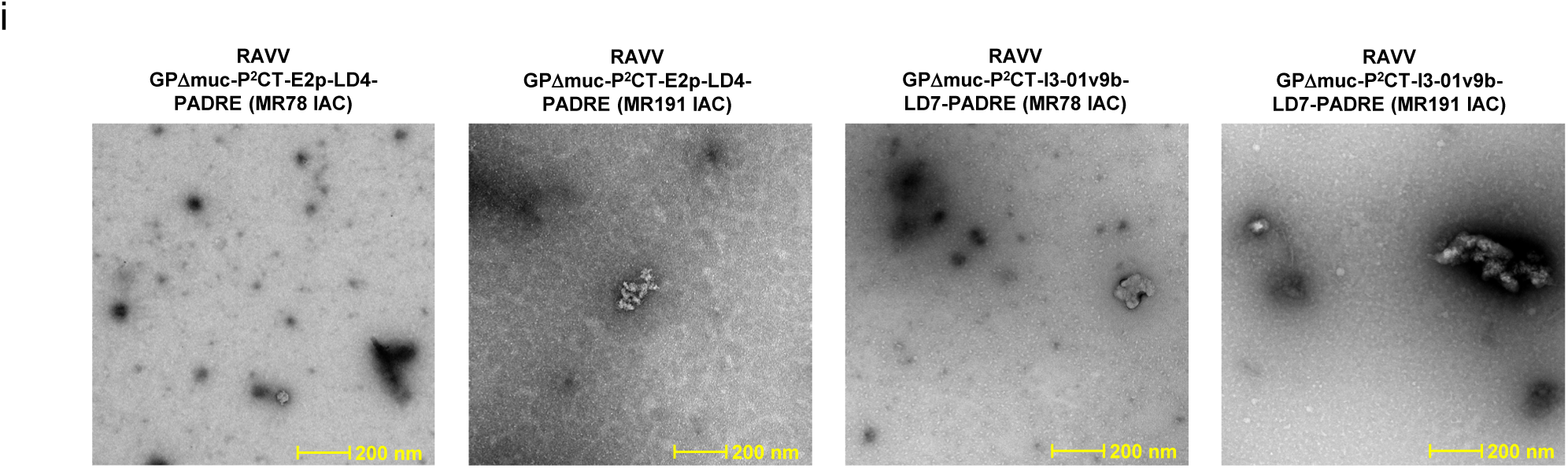
Construct design and in vitro characterization of RAVV GP trimers. (**a**) Constructs sequences of RAVV GP designs. Signal peptide, mucin-like domain (MLD), HR1c, restriction site (AS), foldon, linker (GS), and His_6_-tag are highlighted in yellow, gray, cyan, green, orange, light pink, and light sea green, respectively. The proline mutation on HR1c and CT are shown in red. Notably, the foldon motif, GS linker, and His_6_ tag are present in all RAVV GP trimers tested in this study and will not be included in the construct names to avoid redundancy. (**b**) SEC profile of nickel-purified RAVV GPΔmuc-P^4^ expressed in 16 ml ExpiCHO cells. (**c**) SECs profile of NAb MR78/MR191-purified RAVV GPΔmuc-P^2^ and GPΔmuc-P^2^CT each expressed in 16 ml ExpiCHO cells. (**d**) BN-PAGE of the trimer fractions from the SEC analysis of nickel-purified RAVV GPΔmuc-P^2^ and GPΔmuc-P^2^CT. (**e**) Representative 2D classification images of the RAVV GPΔmuc-P^2^CT trimer in complex with MR78 Fab. (**f**) Compositional site-specific glycan analysis for RAVV GPΔmuc-P^2^CT trimer, as in **Fig. S1c**. (**g**) ELISA analysis of RAVV GPΔmuc-P^2^ and GPΔmuc-P^2^CT trimers binding to NAbs MR78 and MR191 in the IgG form. Briefly, each well was coated with 0.1 μg of the appropriate antigen and IgG antibodies were diluted in a 10-fold dilution series from a starting concentration of 10 μg/ml for all antibodies. Error bars represent the difference between duplicate measurements at each concentration for each sample. (**h**) BLI analysis of RAVV GPΔmuc-P^2^ and GPΔmuc-P^2^CT trimers binding to NAbs MR78 and MR191 in the IgG form, as in **Fig. S1i**. (i) nsEM analysis of RAVV GPΔmuc-P^2^CT E2p and I3-01v9b constructs following ExpiCHO expression and MR78- or MR191-based IAC purification. Scale bar is labeled in yellow on the EM micrographs. No discernible protein NPs are shown in any of the EM micrographs.

**Fig. S6.**
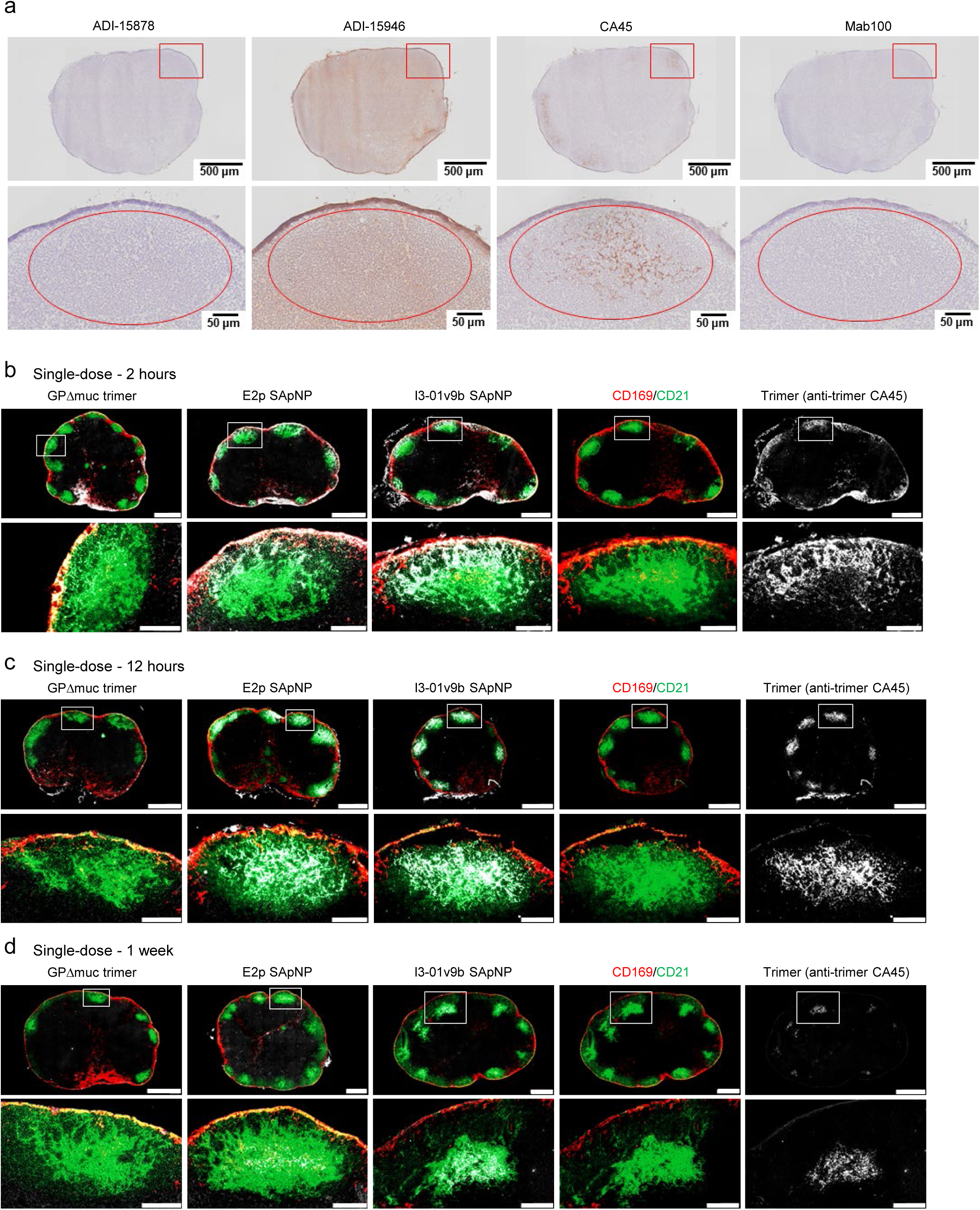
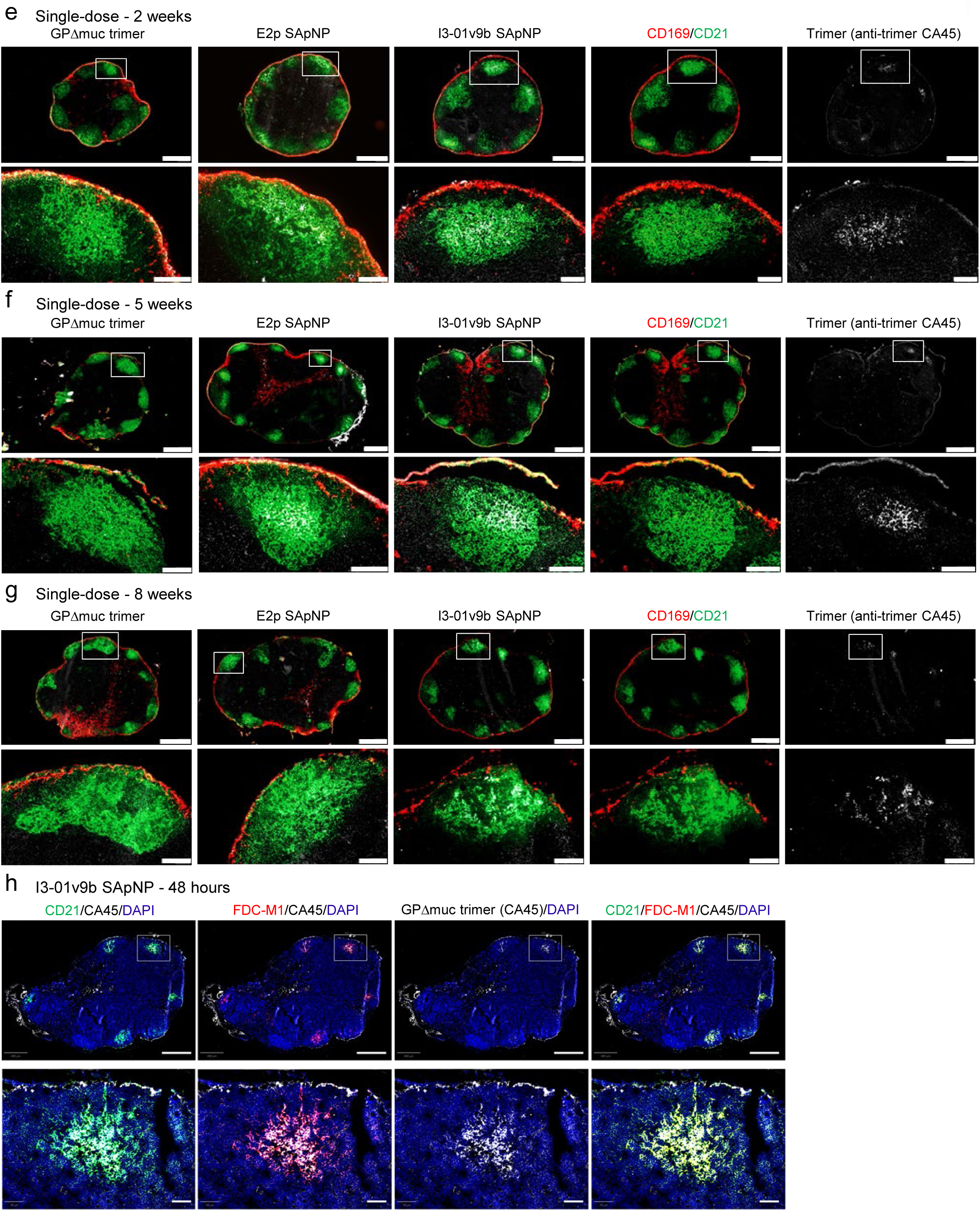
Immunohistological images of SUDV GPΔmuc trimer and SApNPs in lymph nodes. (**a**) Immunostaining images of lymph node tissues from mice injected with SUDV GPΔmuc-presenting SApNPs, stained using antibodies ADI-15878, ADI-15946, CA45, and mAb100. **(b–g)** Colocalization of SUDV GPΔmuc trimer, E2p SApNP, and I3-01v9b SApNP with FDC networks in lymph node follicles at various time points following a single-dose injection (10 μg per injection, 40 μg total per mouse): (**b**) 2 hours, (**c**) 12 hours, (**d**) 1 week, (**e**) 2 weeks, (**f**) 5 weeks, and (**g**) 8 weeks. Immunofluorescent images are pseudo-color-coded as follows: CD21⁺ (green), CD169⁺ (red), and CA45 (white). (**h**) Colocalization of SUDV GPΔmuc-presenting I3-01v9b SApNPs (labeled by CA45, white) with FDC networks labeled by CD21⁺ (green) and FDC-M1⁺ (red) at 48 hours post-injection. Scale bars: 500 μm (entire lymph node) and 100 μm (enlarged follicle image).

**Fig. S7.**
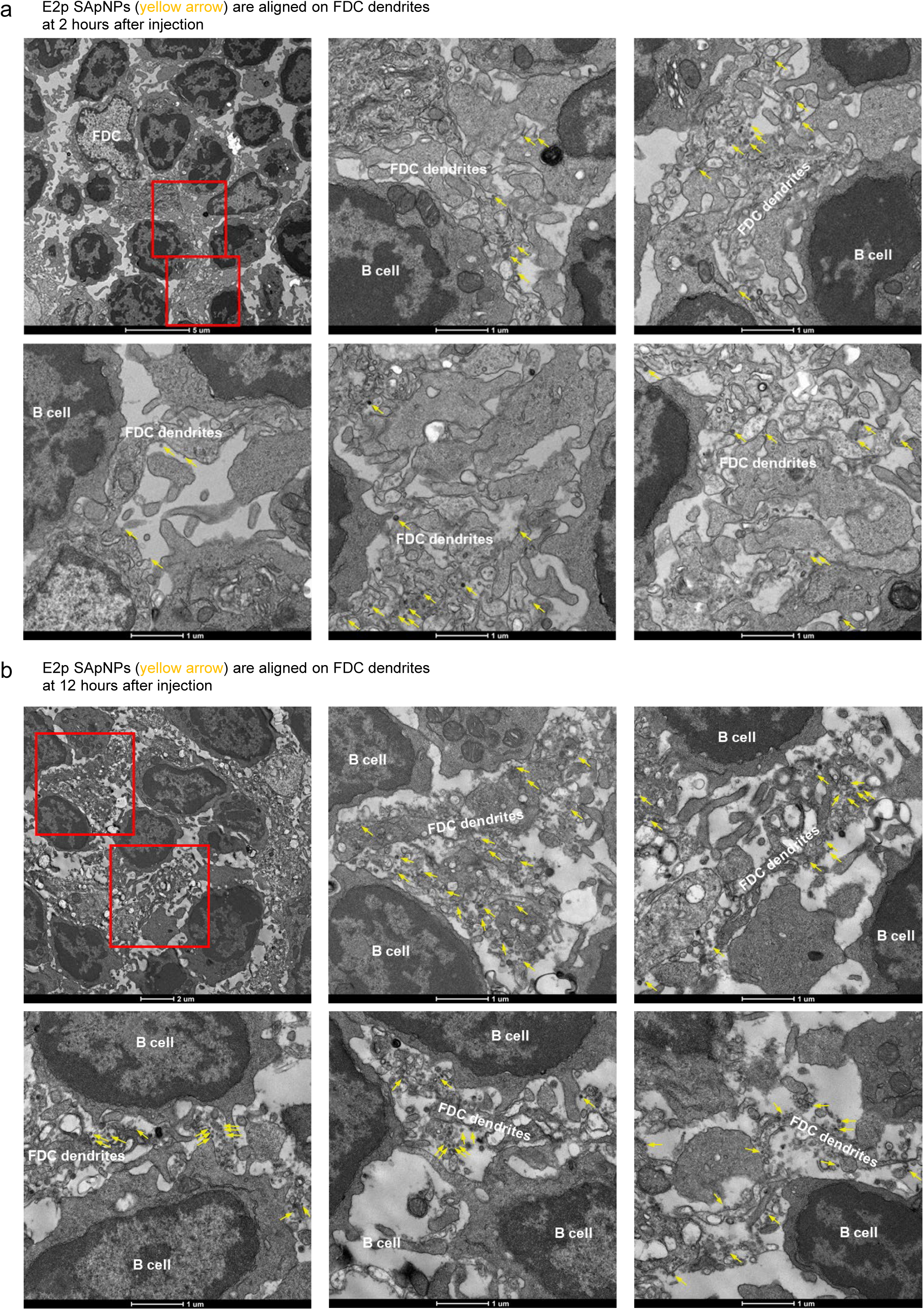
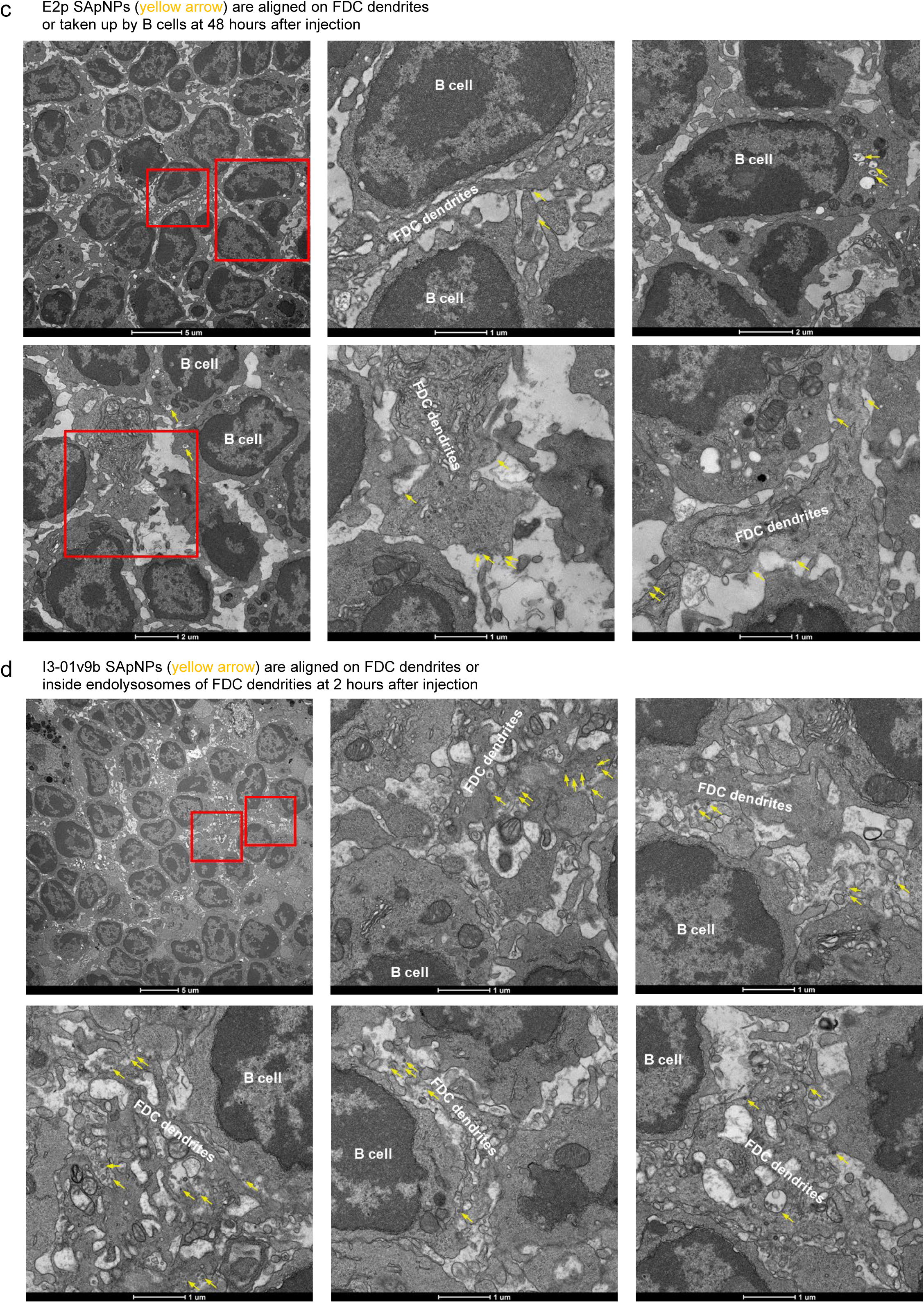
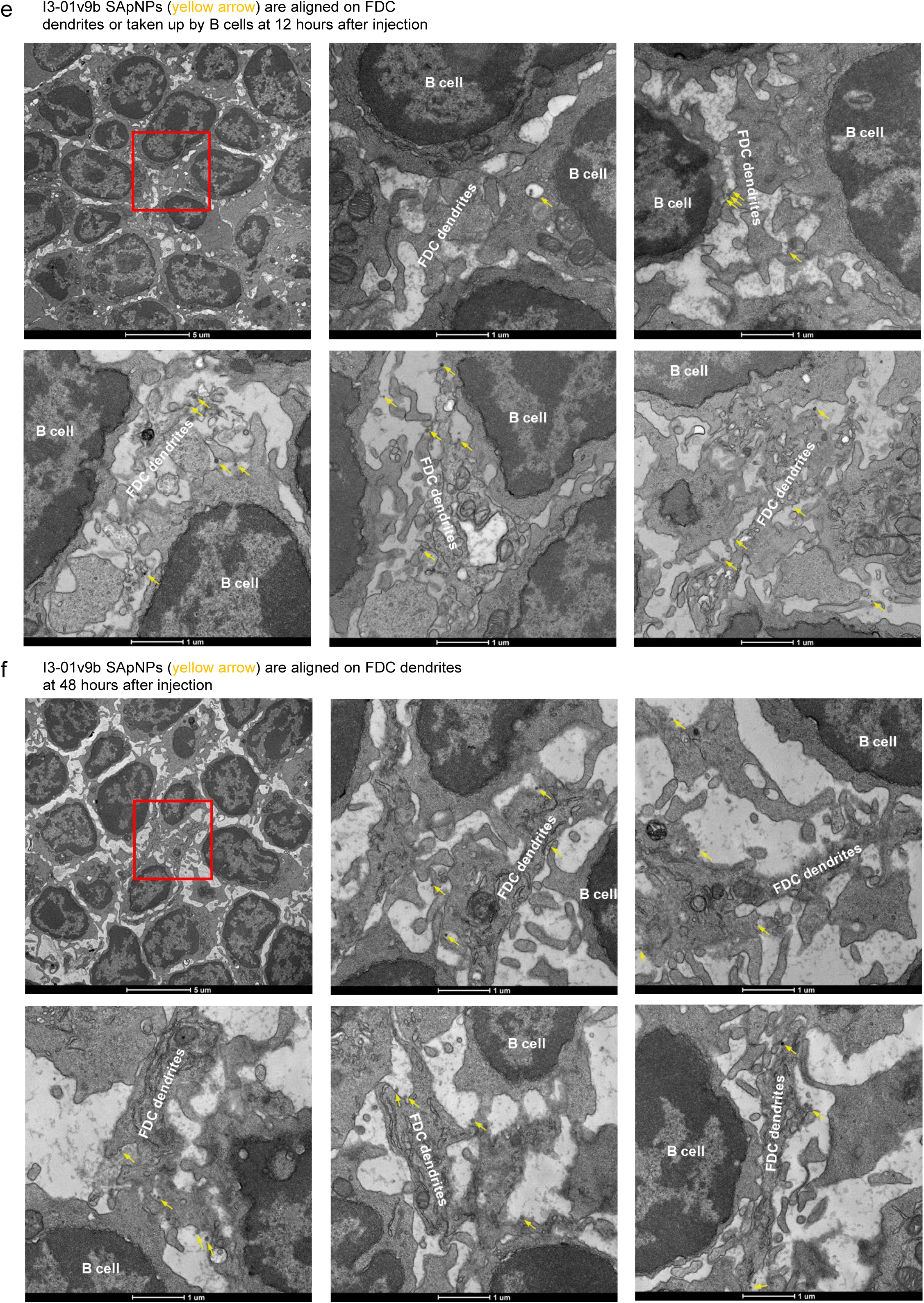
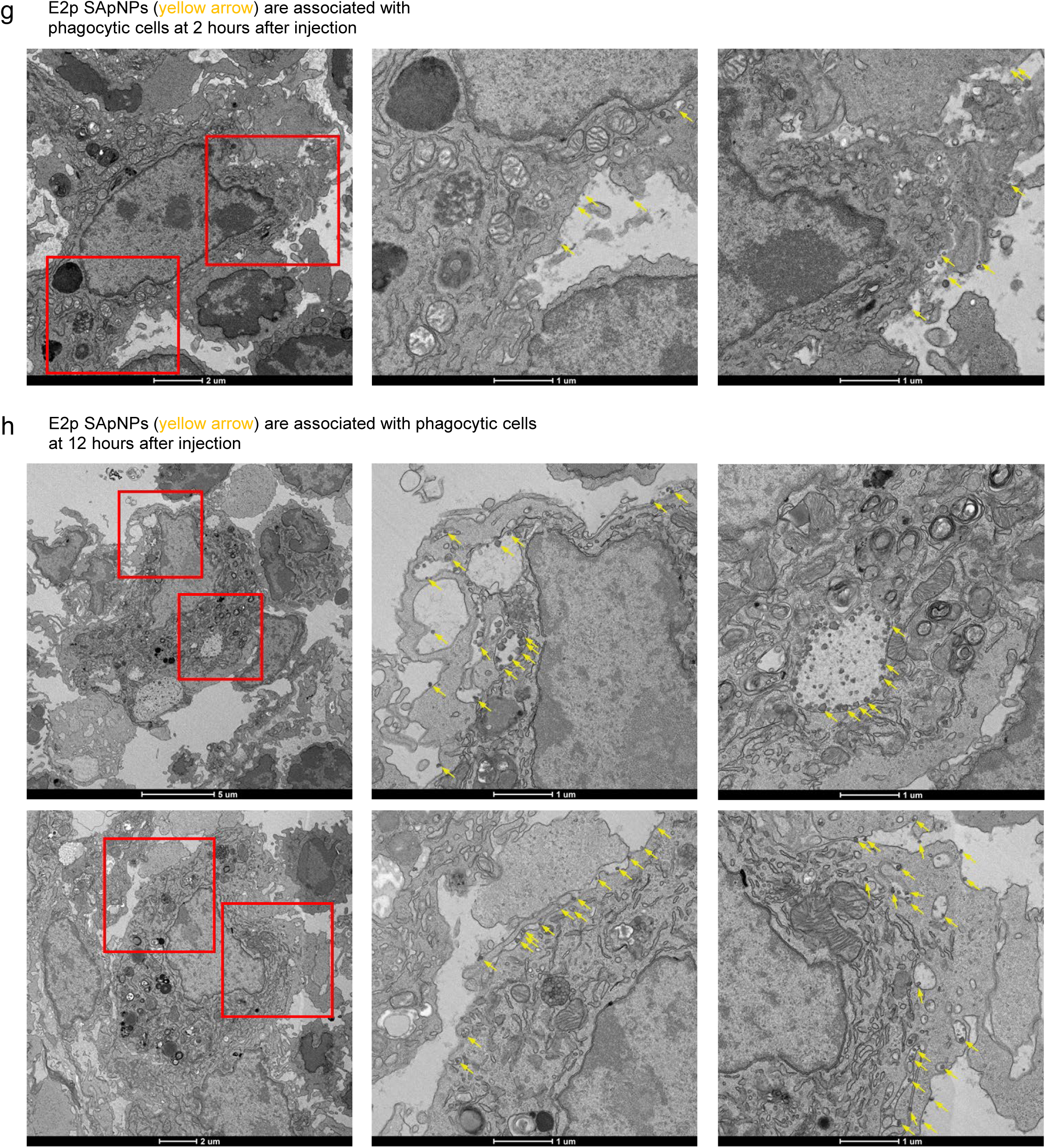
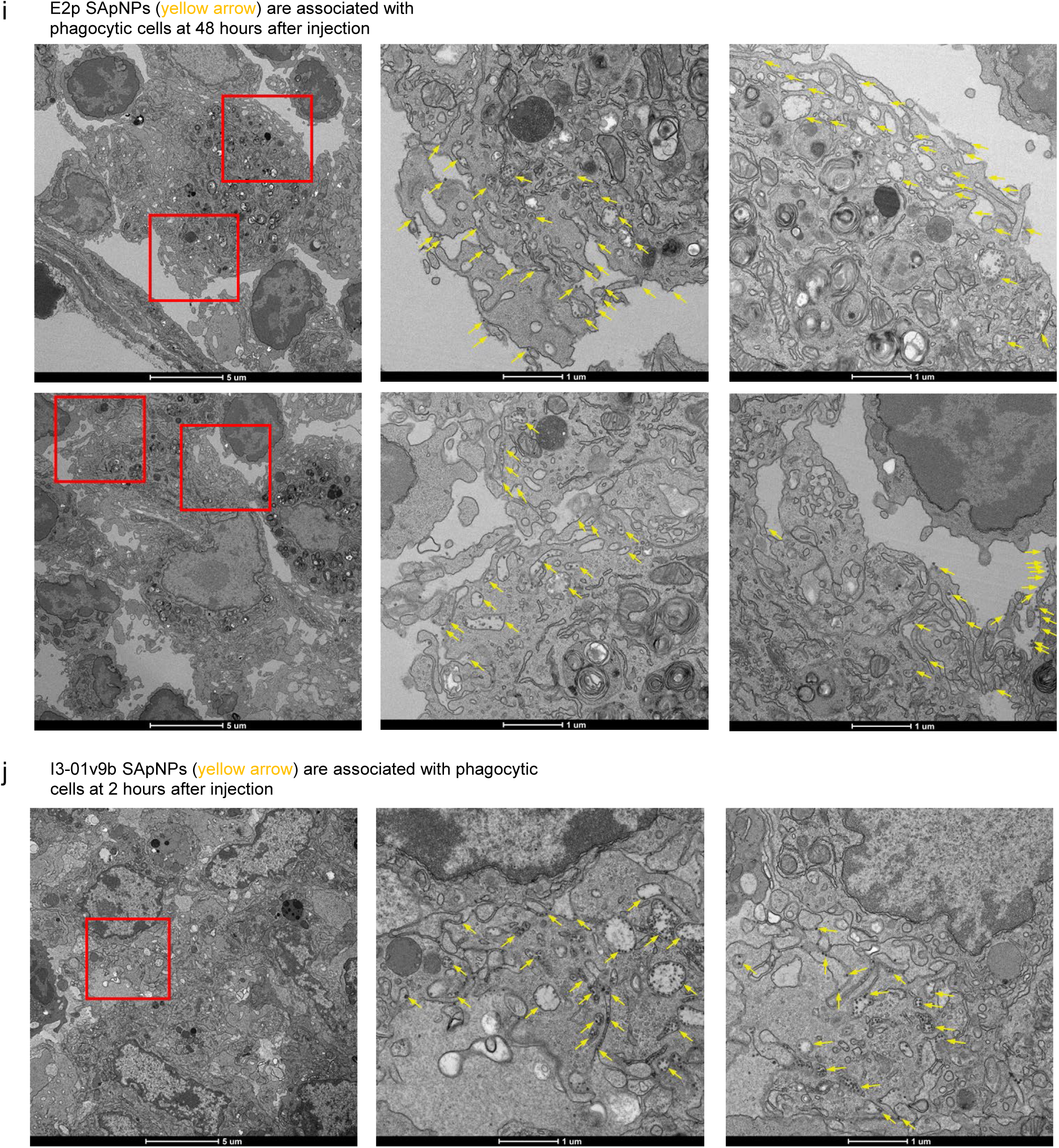
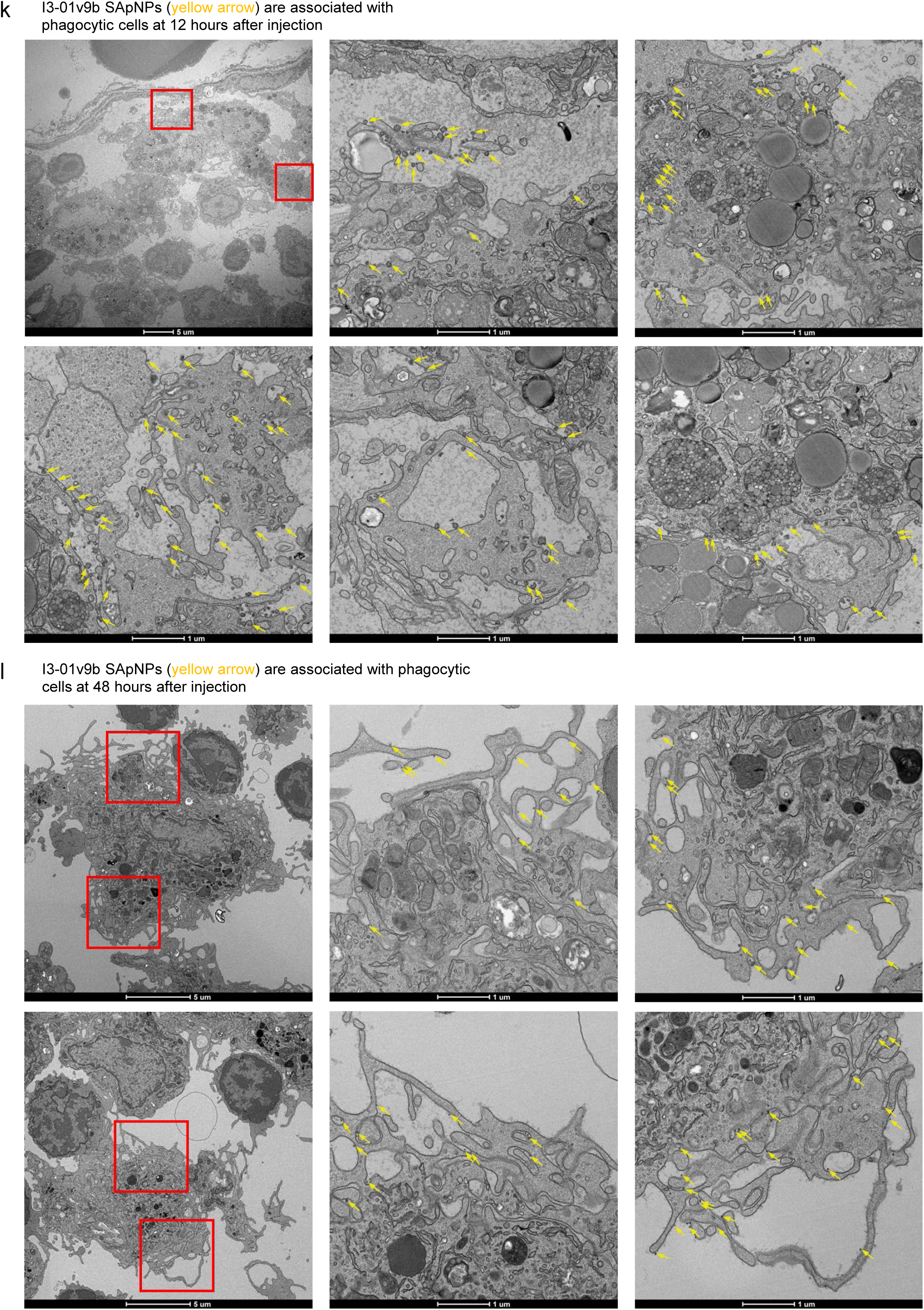
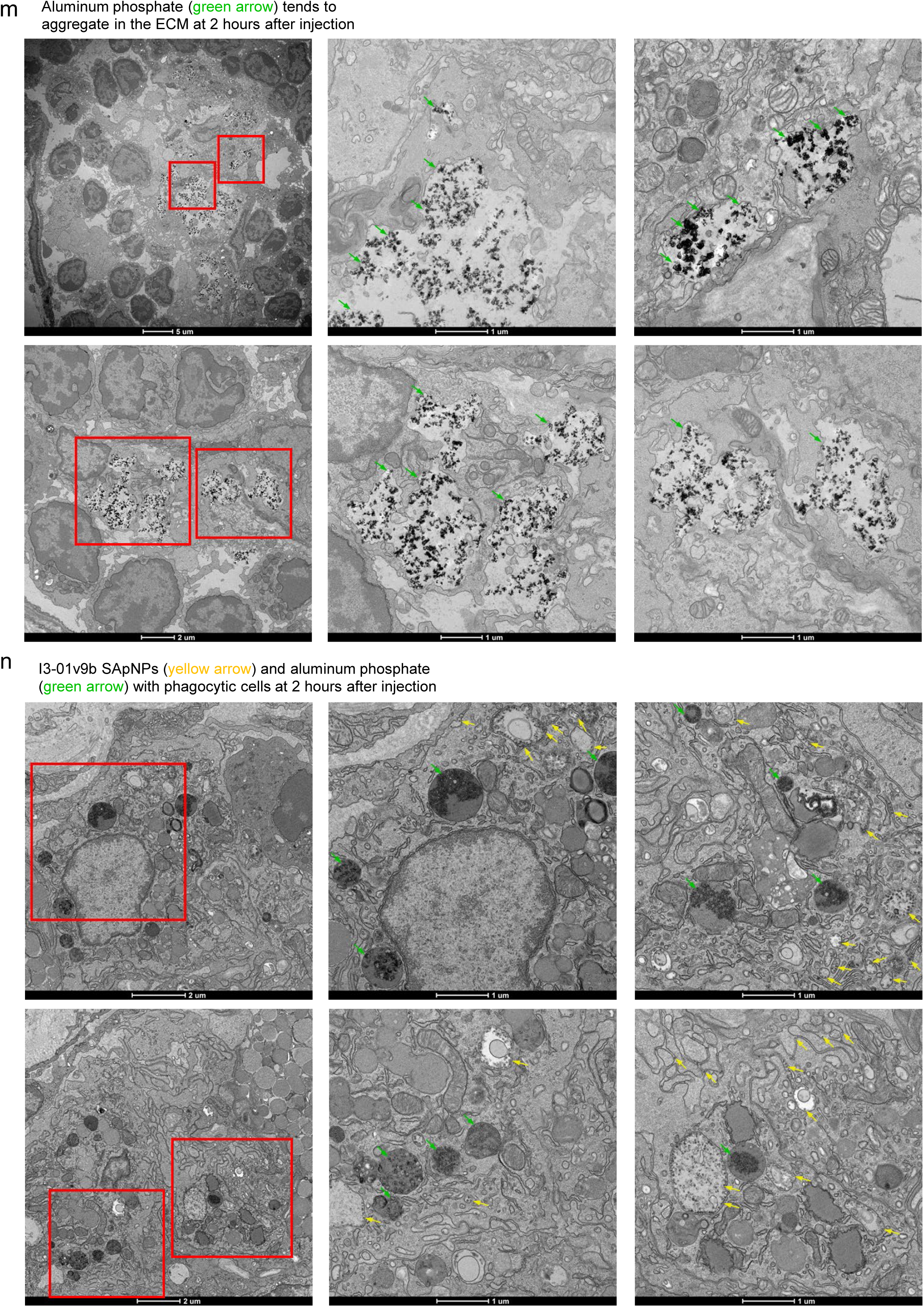
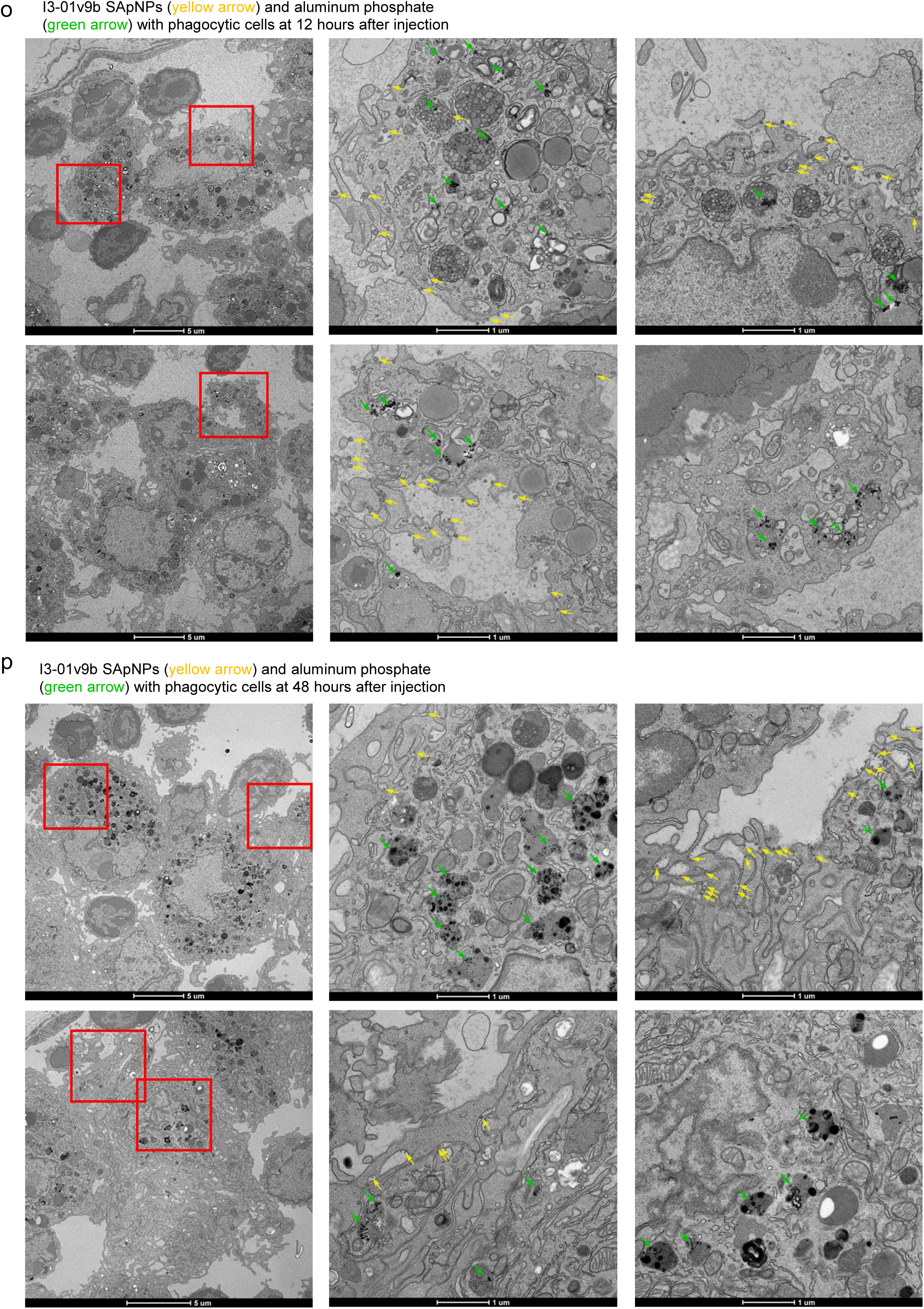
TEM analysis of SUDV GPΔmuc-presenting SApNPs interacting with FDCs and phagocytic cells in lymph nodes. TEM images show the distribution and cellular interactions of SUDV GPΔmuc-presenting SApNPs following a single-dose injection (2 footpads, 50 μg/footpad). (**a–c**) E2p SApNPs aligned along FDC dendrites at 2 hours, 12 hours, and 48 hours post-injection, respectively. (**d–f**) I3-01v9b SApNPs aligned on FDC dendrites at the same time points. Aluminum phosphate (AP) adjuvant particles are not observed in association with SApNPs on FDC dendrites. (**g–i**) E2p SApNPs located on the surface or within endolysosomes of phagocytic cells at 2 hours, 12 hours, and 48 hours post-injection, respectively. (**j–l**) I3-01v9b SApNPs associated with phagocytic cells at the same time points. (**m**) AP adjuvant particles aggregating in the extracellular matrix 2 hours after injection. (**n–p**) I3-01v9b SApNPs formulated with AP observed within phagocytic cells at 2 hours, 12 hours, and 48 hours post-injection. TEM imaging was performed on two popliteal lymph nodes per SApNP construct. E2p and I3-01v9b SApNPs are indicated by yellow arrows; AP adjuvant particles are indicated by green arrows.

**Fig. S8.**
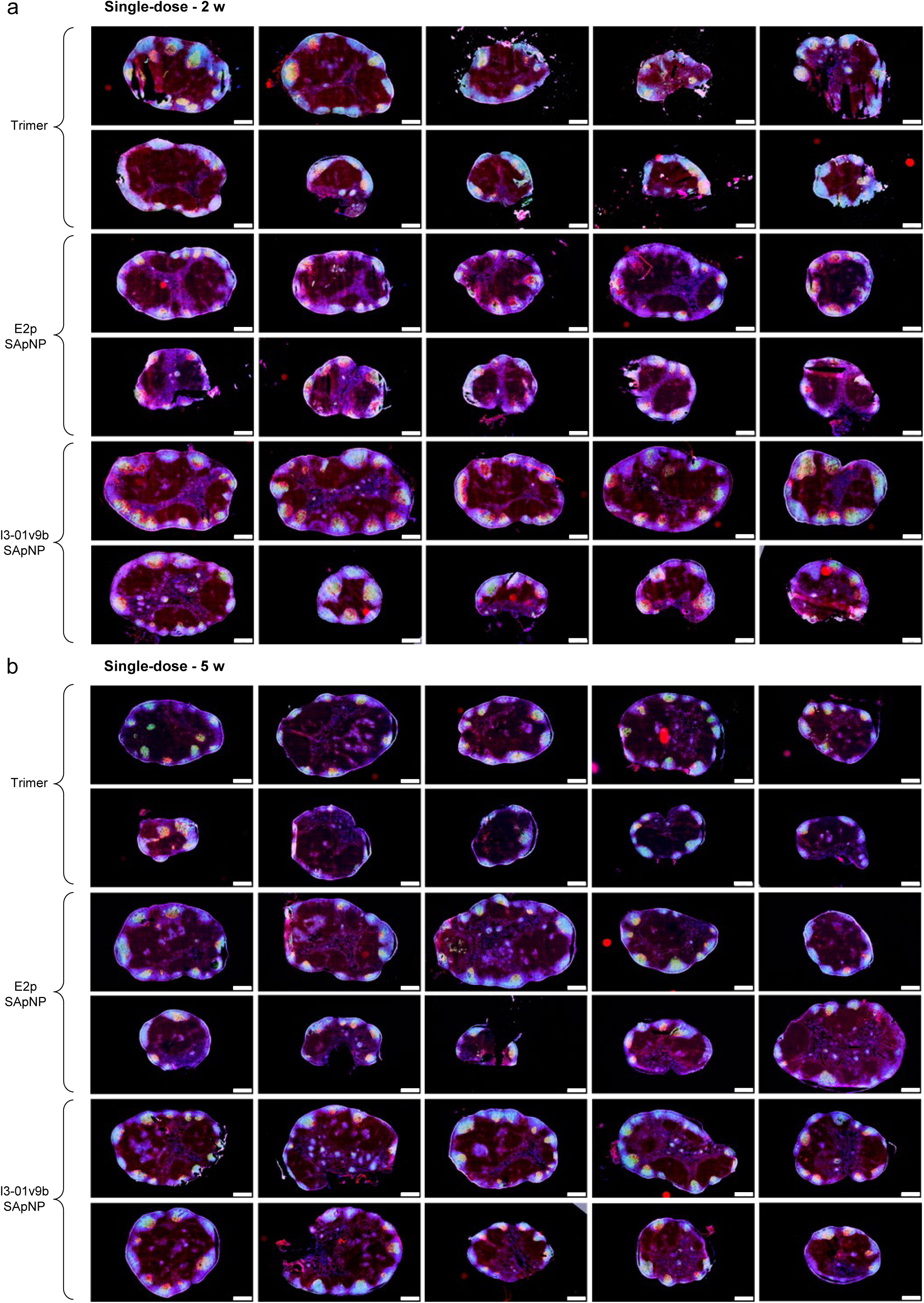
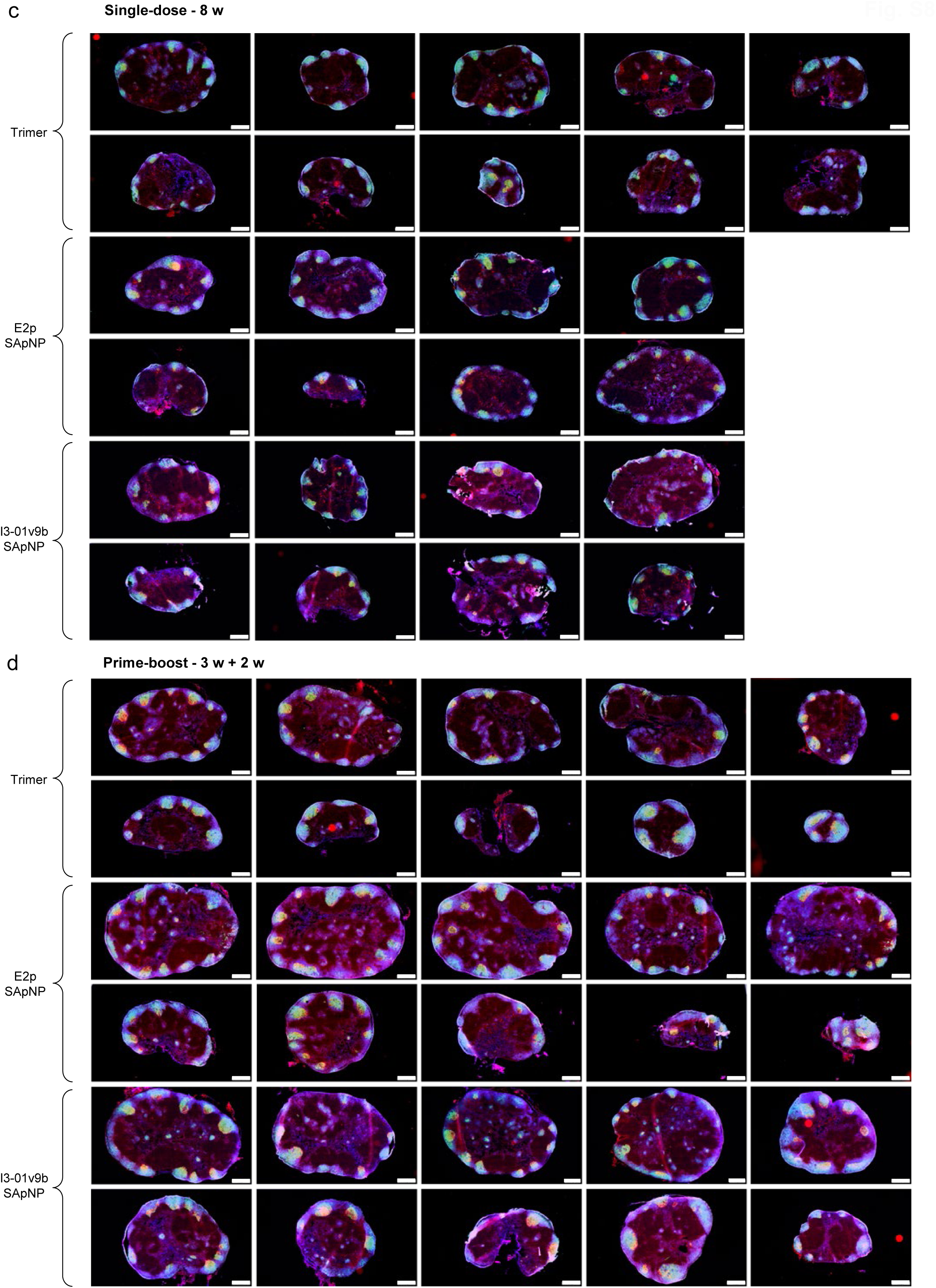
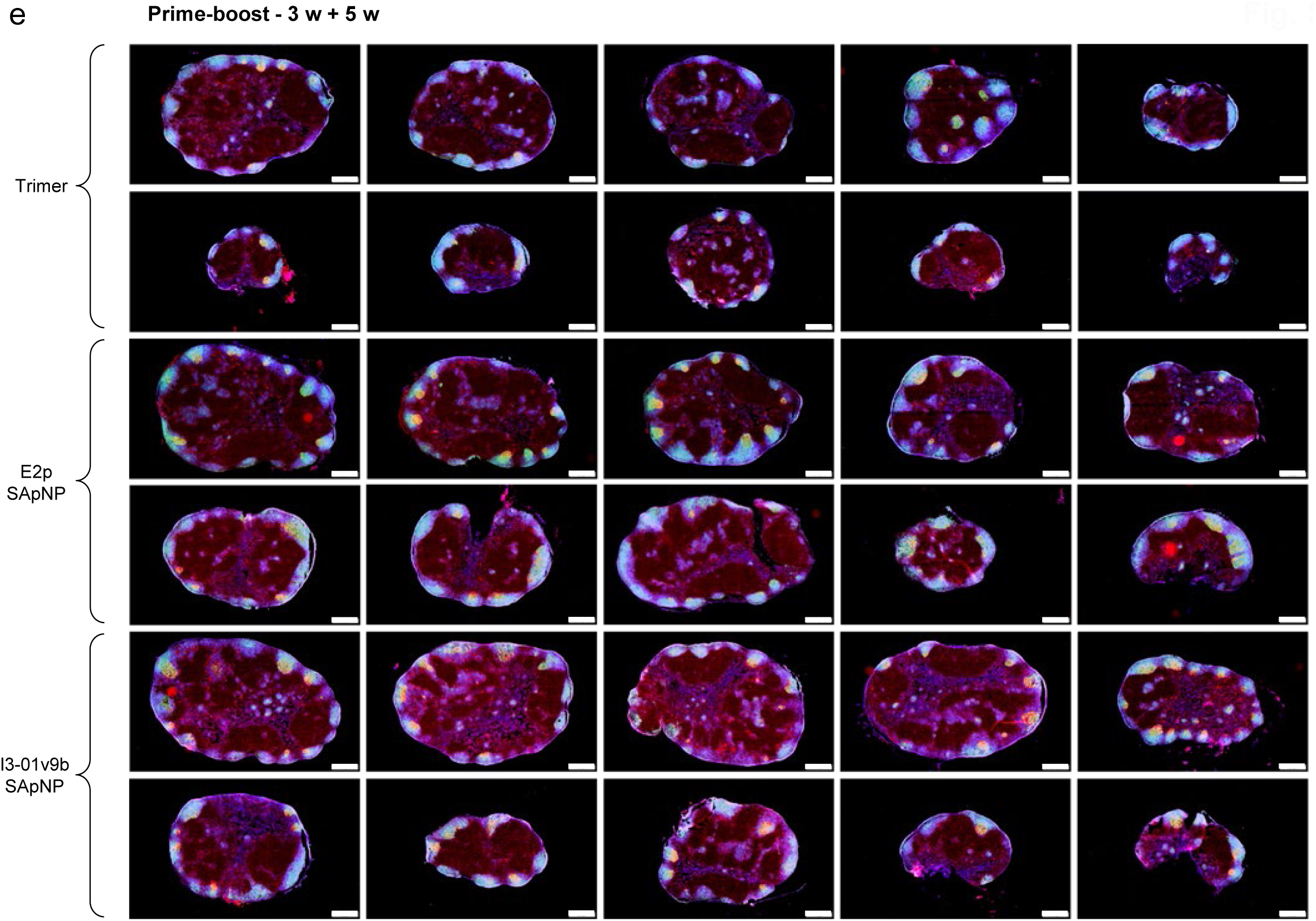
Immunohistological analysis of SUDV GPΔmuc trimer and SApNP vaccine-induced germinal centers (GCs). Immunohistological images of GCs at (**a**) 2, (**b**) 5, and (**c**) 8 weeks following a single-dose injection of GPΔmuc trimer or GPΔmuc-presenting E2p and I3-01v9b SApNPs (10 μg per injection; total of 40 μg per mouse). GC images at (**d**) 2 and (**e**) 5 weeks after a booster dose, administered 3 weeks after the initial immunization (n = 5 mice/group). Immunofluorescent images are pseudo-color coded (GL7^+^, red; CD21^+^, green; B220, blue). Scale bars = 500 μm for each lymph node image.

**Fig. S9.**
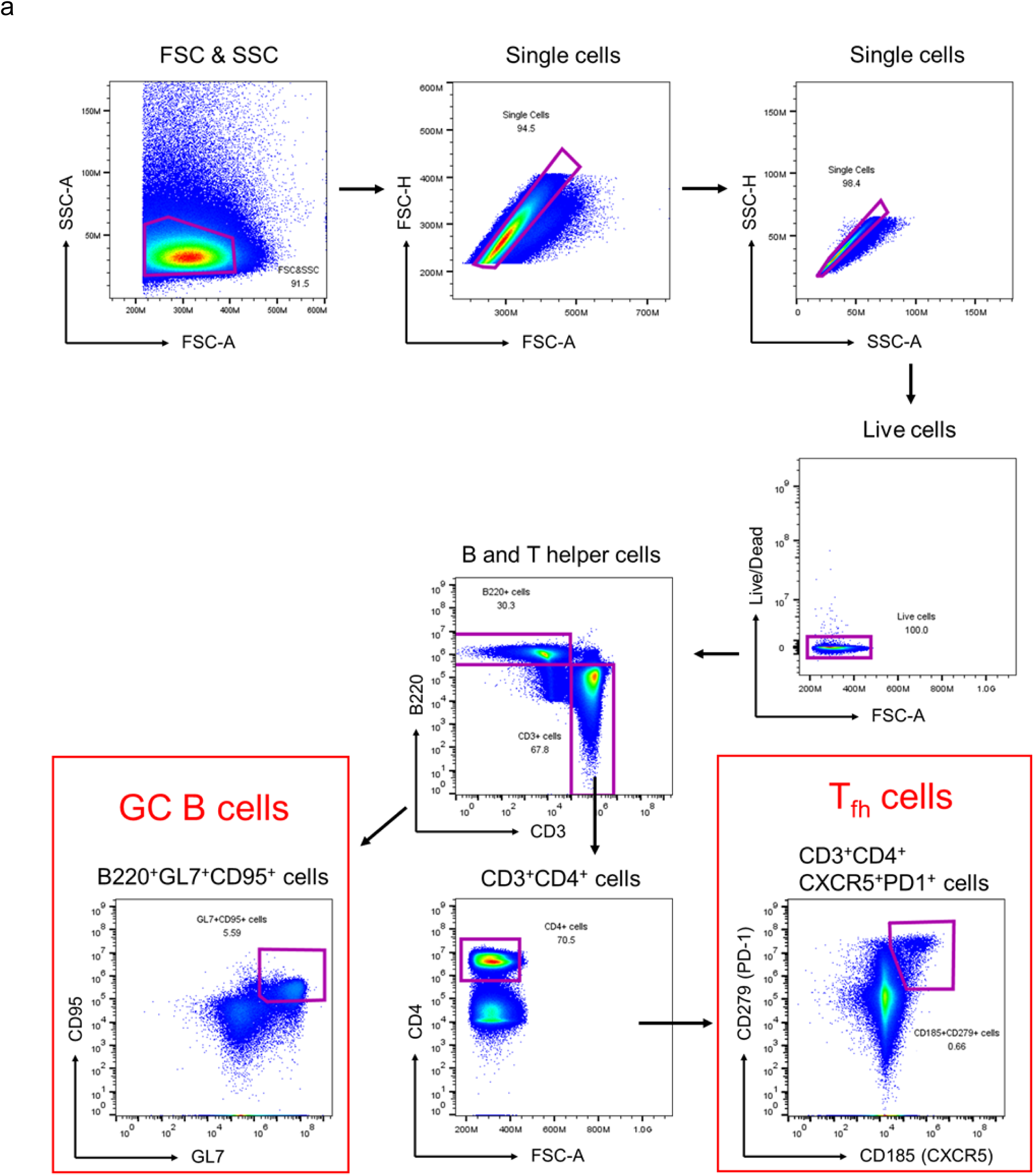
Flow cytometry analysis of germinal centers (GCs) induced by SUDV GPΔmuc trimer and SApNPs vaccines. Gating strategy for identifying GC B cells and T follicular helper (Tfh) cells by flow cytometry (n = 5 mice per group).

**Fig. S10.**
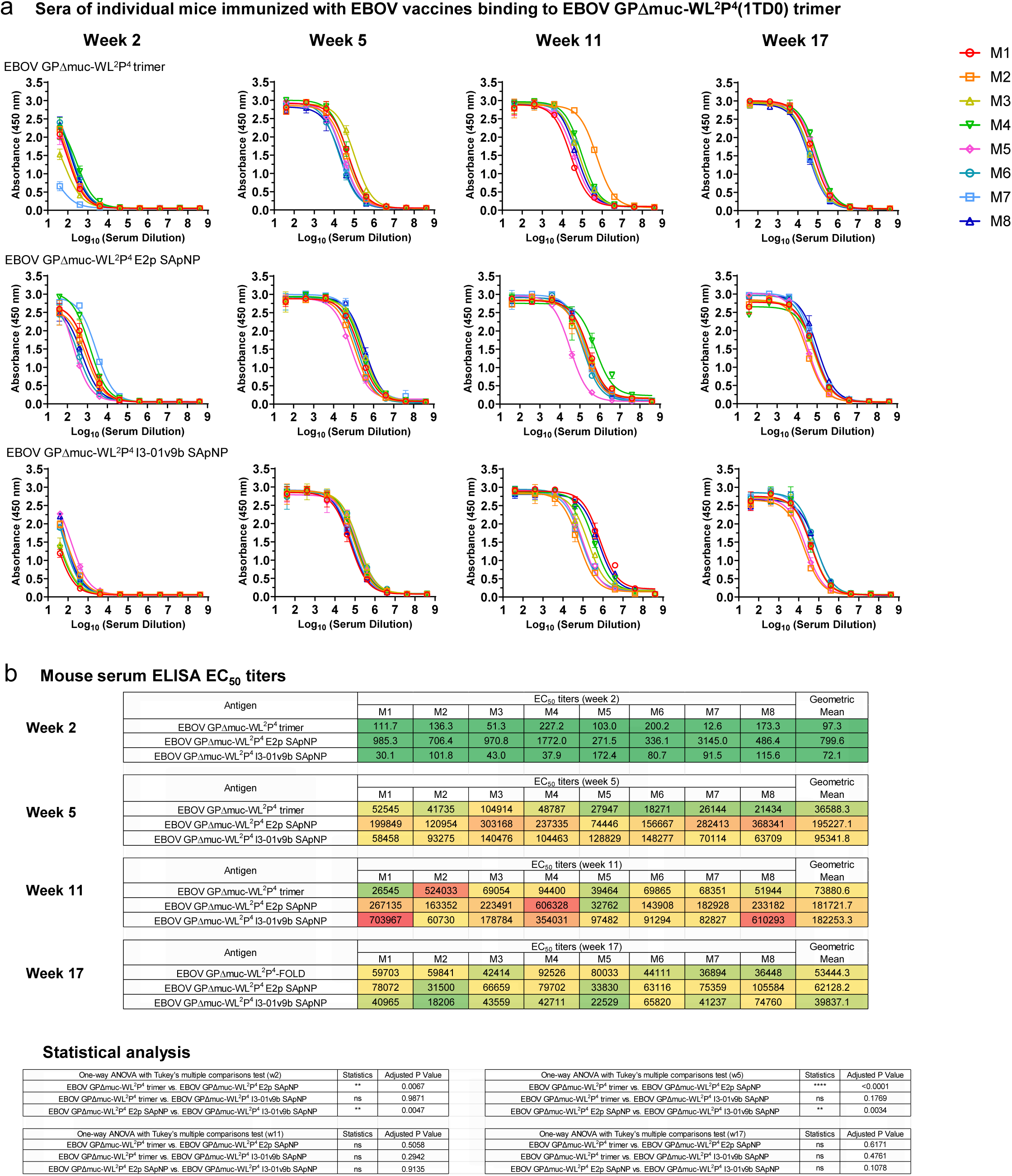
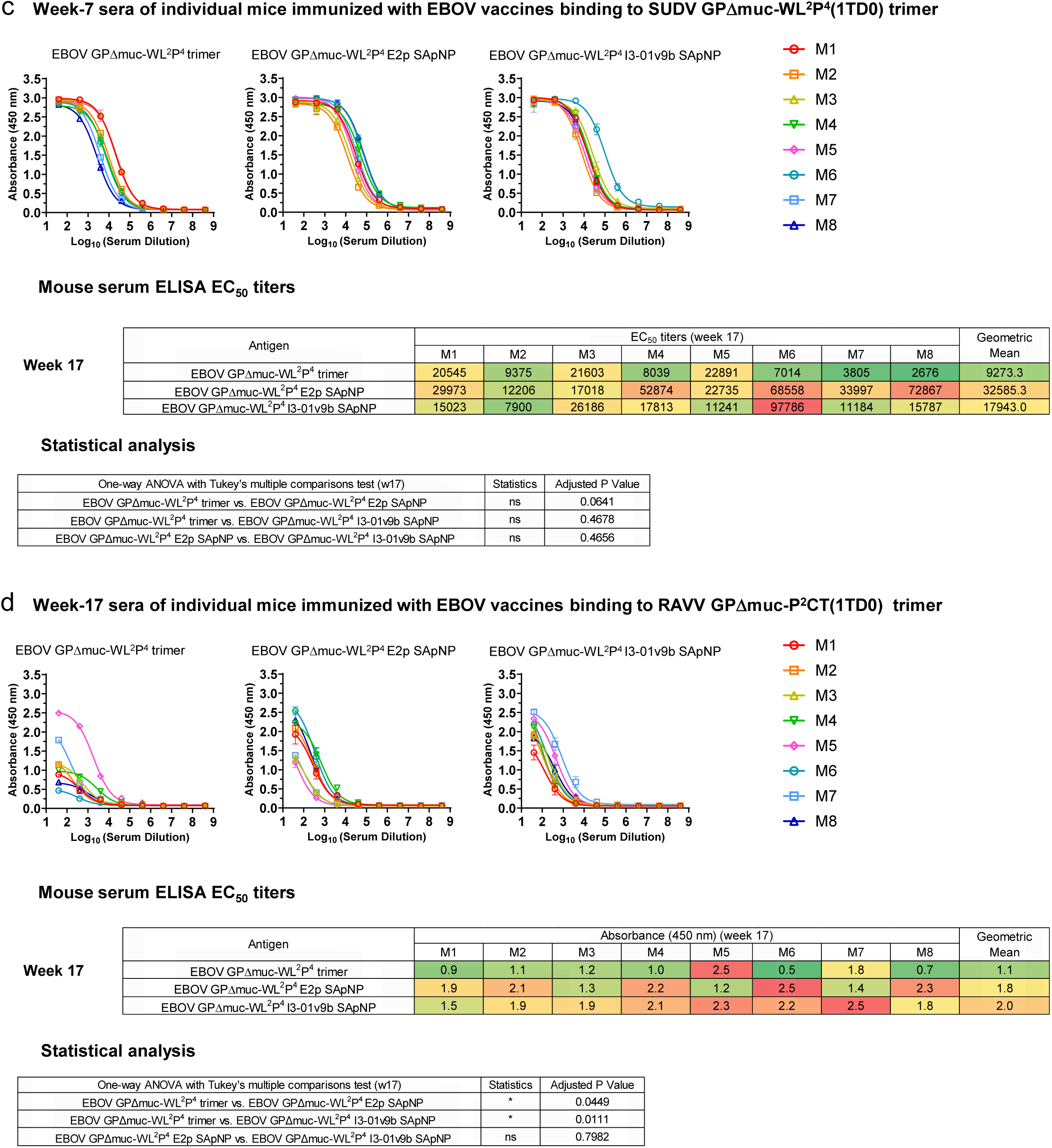
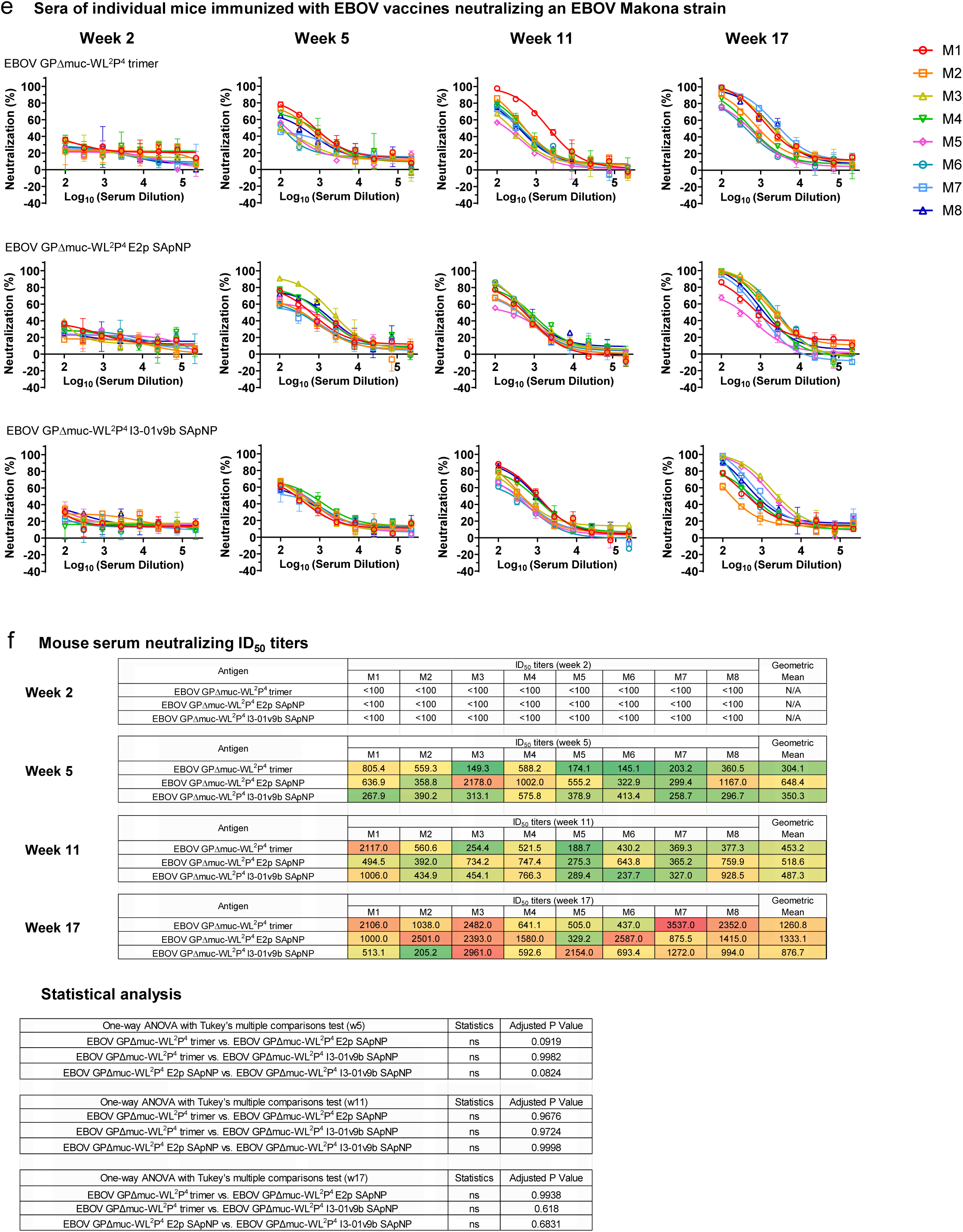
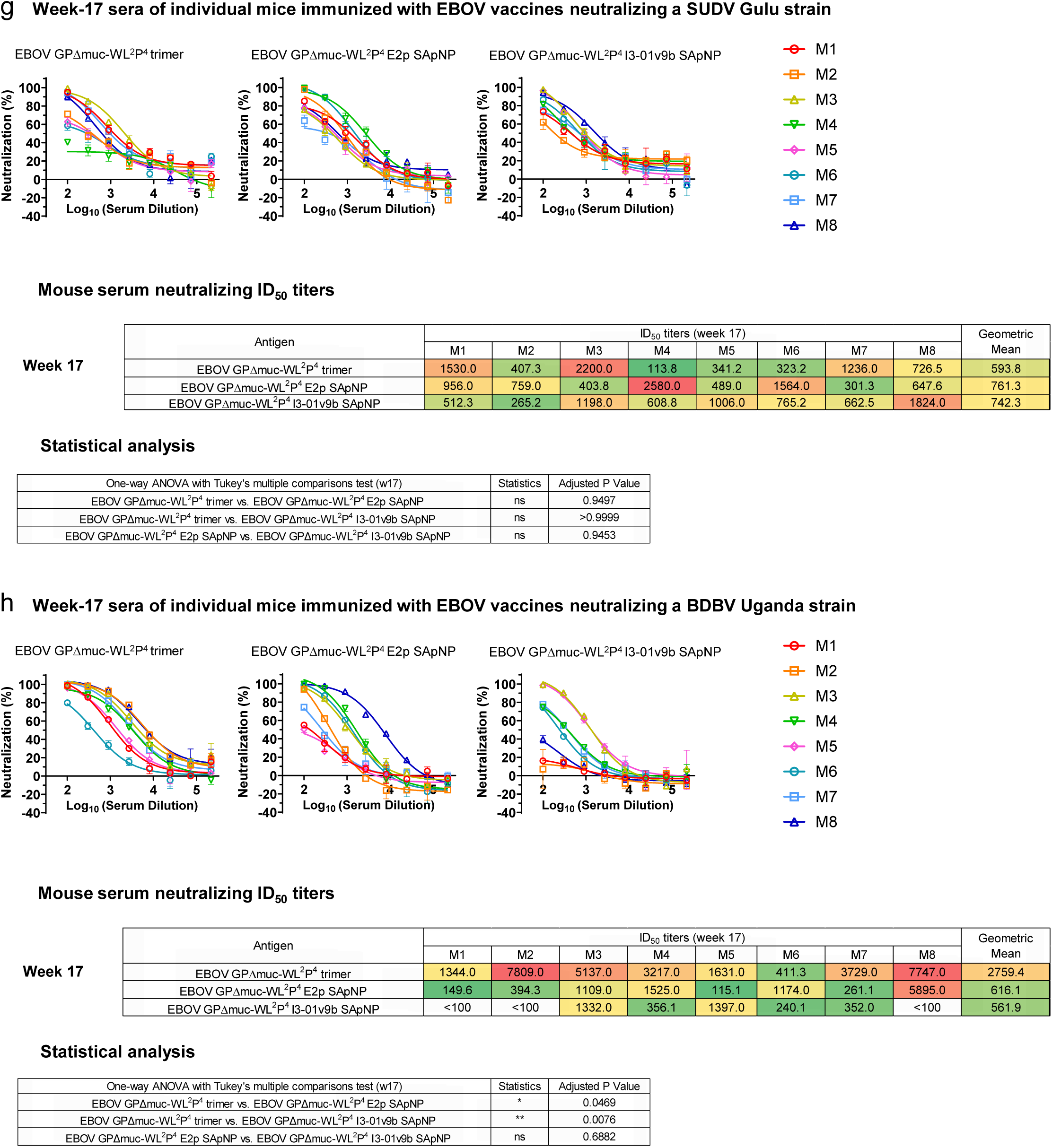
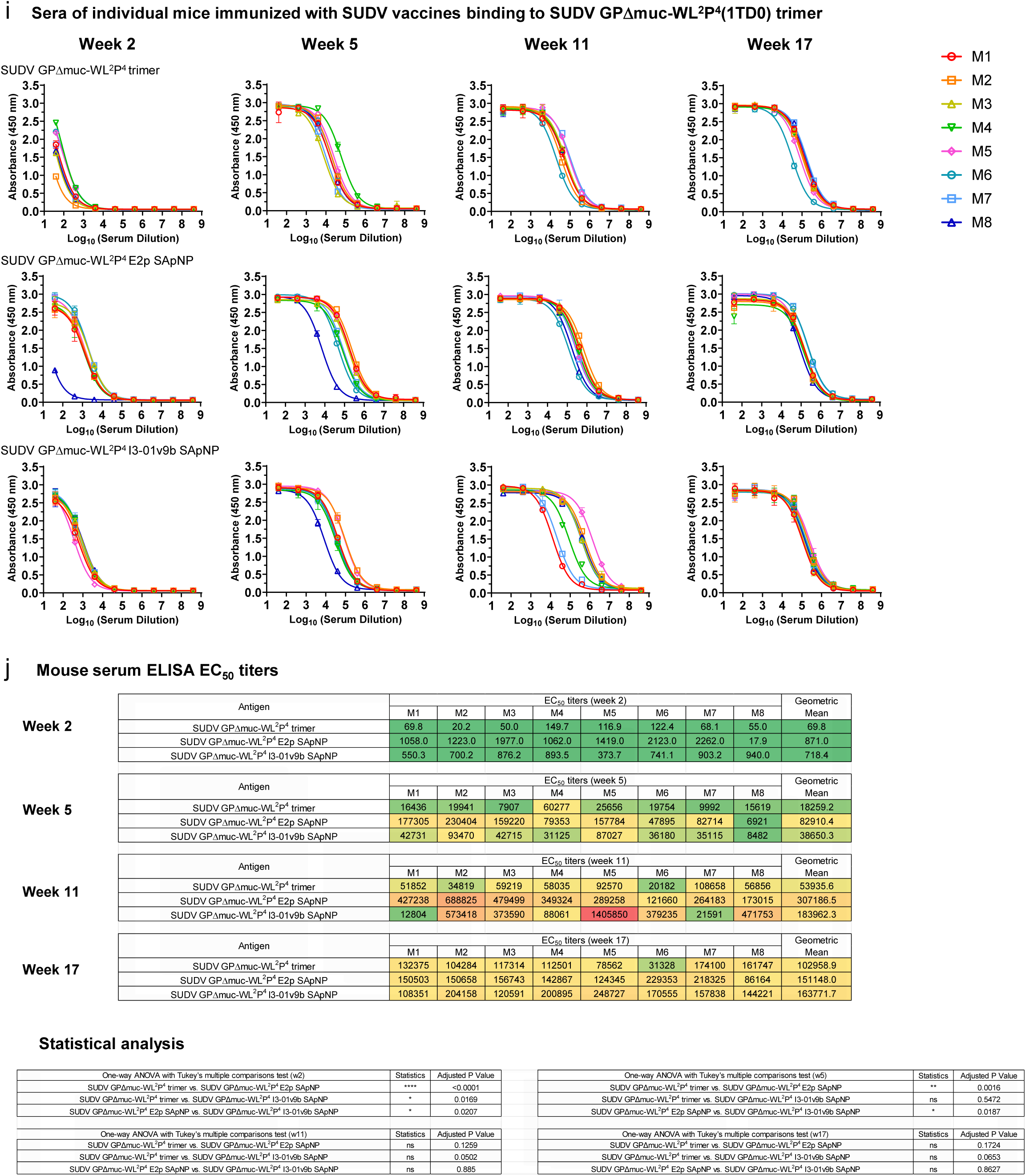
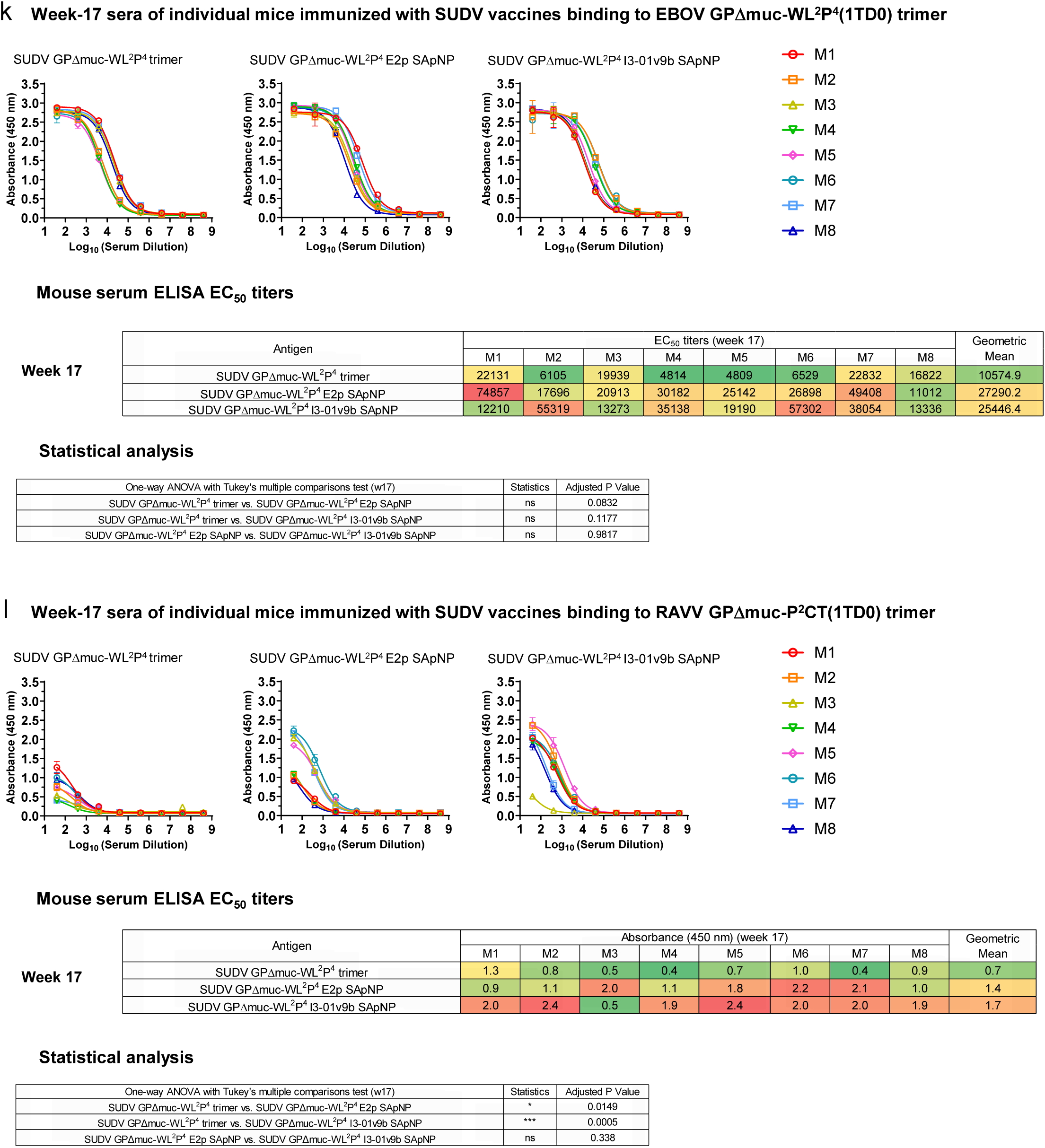
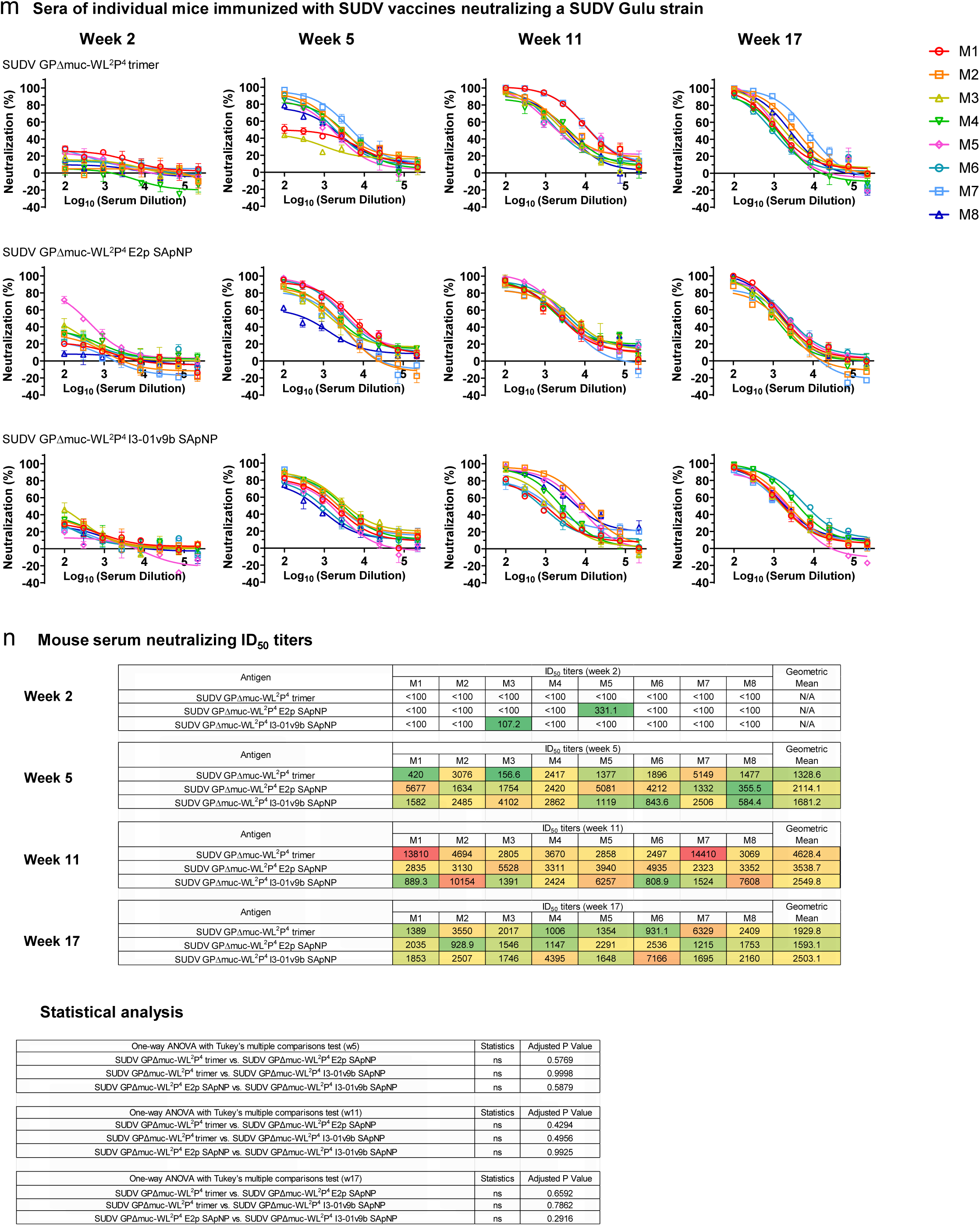
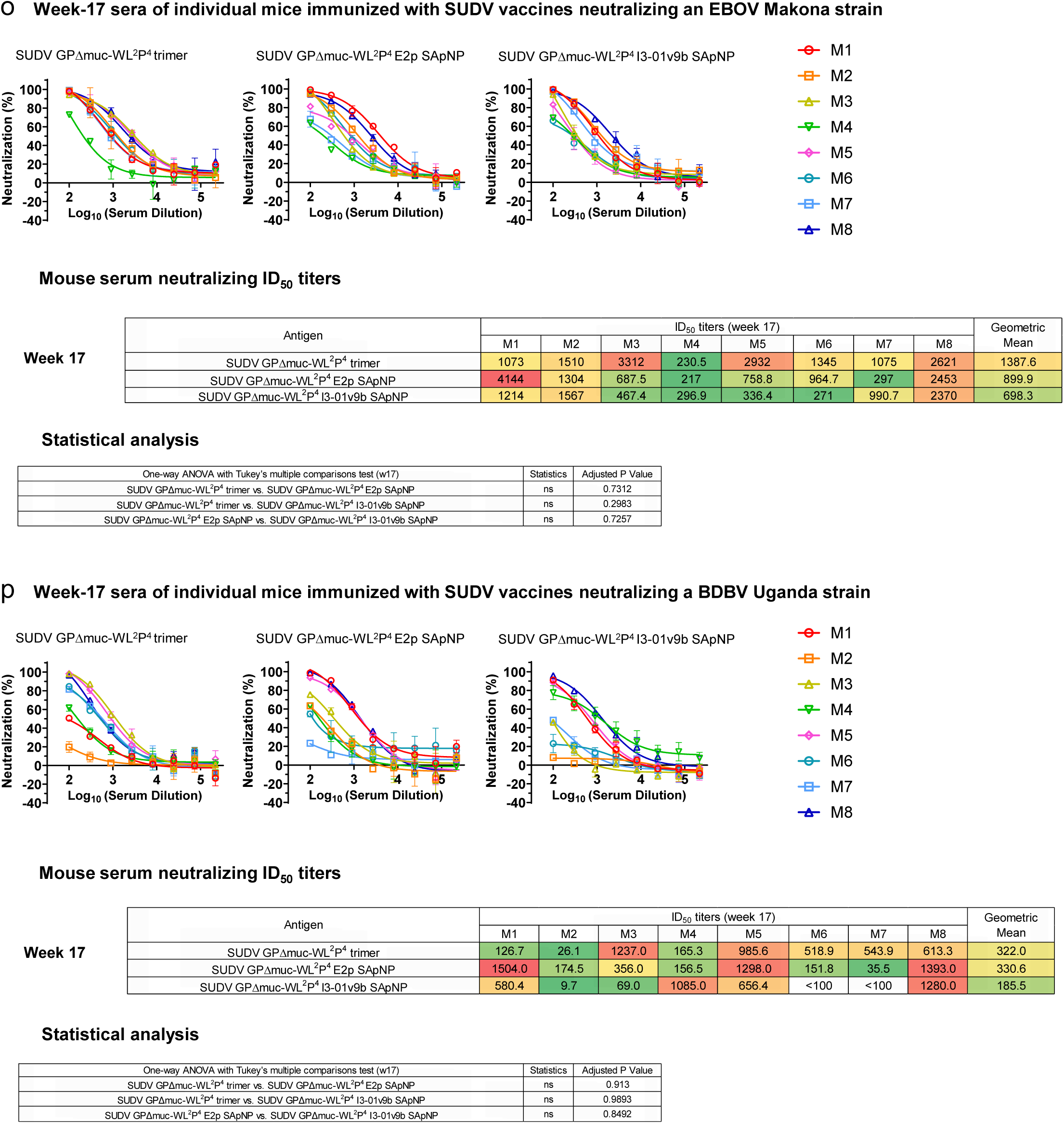
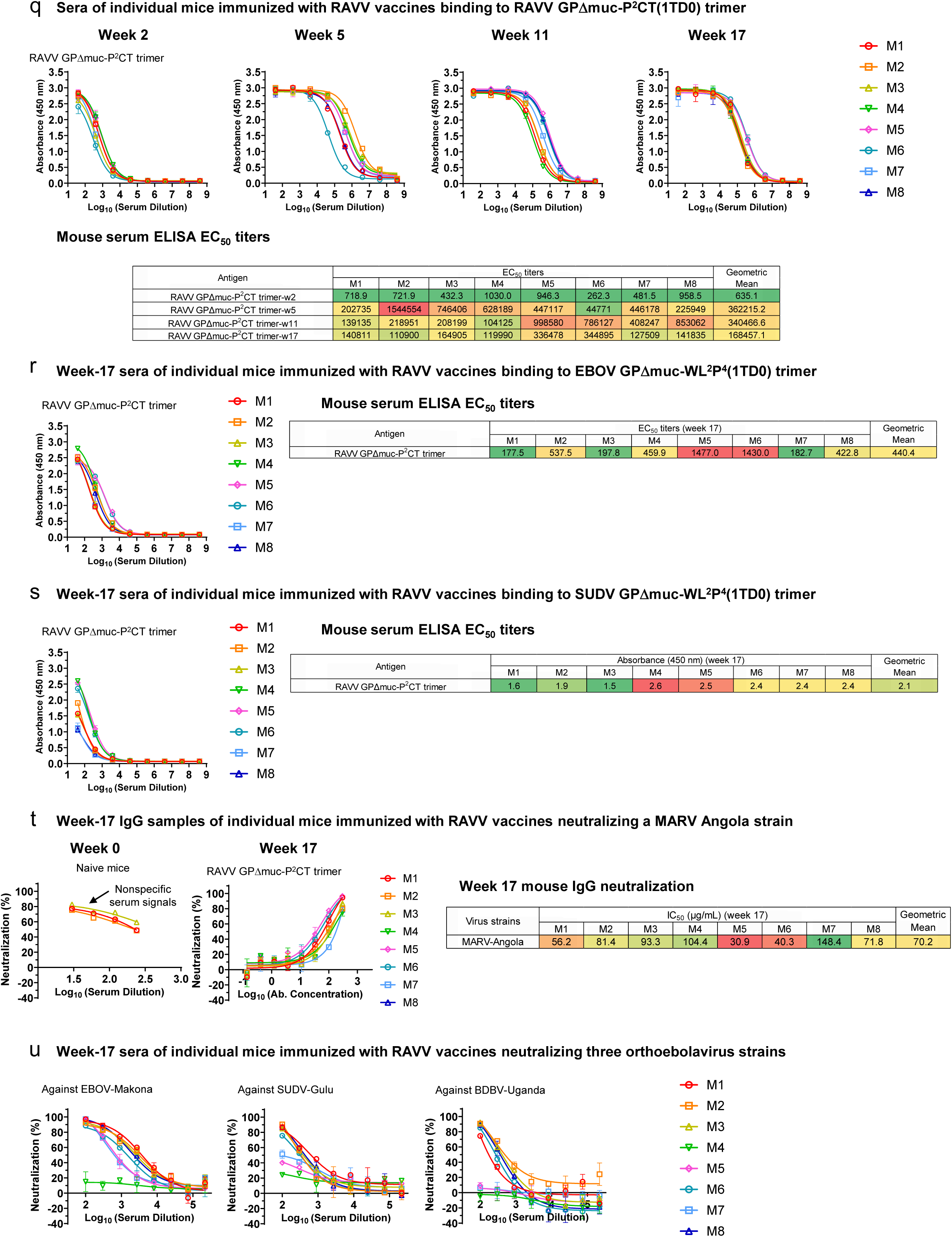
Immunogenicity of EBOV, SUDV, and RAVV GPΔmuc vaccines in mice. (**a**) ELISA curves of mouse sera from EBOV GPΔmuc trimer and SApNP vaccine groups (n = 8 mice/group) binding to the coating antigen EBOV GPΔmuc-WL^2^P^4^(1TD0) trimer (**b**) (Top) Summary of geometric mean EC_50_ titers measured for Ebola GPΔmuc vaccine groups against EBOV GPΔmuc-WL^2^P^4^(1TD0) trimer. Color coding indicates EC_50_ levels (green to red: low to high binding). (Bottom) Summary of statistical analysis performed for each timepoint. Note: EC_50_ values at week 2 were derived by setting the minimum/maximum OD_450_ values to 0.0/2.8. (**c**) ELISA curves of mouse sera from EBOV GPΔmuc vaccine groups at week 17 after four immunizations binding to SUDV GPΔmuc-WL^2^P^4^(1TD0) trimer. Summary of geometric mean EC_50_ titers and statistical analysis. (**d**) ELISA curves of mouse sera from Ebola GPΔmuc vaccine groups at week 17 after four immunizations binding to RAVV GPΔmuc-P^2^CT(1TD0) trimer. Summary of absorbance at 450 nm (A450) values was included. (**e**) Neutralization curves of mouse sera from EBOV GPΔmuc trimer and SApNP vaccine groups against EBOV Makona pseudovirus. (**f**) (Top) Summary of geometric mean ID_50_ titers measured for EBOV GPΔmuc vaccine groups against EBOV Makona pseudovirus. Color coding: white (no neutralization), green to red (low to high neutralization). Note: ID_50_ values were calculated using %neutralization constraints of 0.0 (min) and 100.0 (max). (Bottom) Summary of statistical analysis as in (**b**). (**g**) Neutralization curves of mouse sera from EBOV GPΔmuc vaccine groups at week 17 against SUDV Gulu pseudovirus. Summary of geometric means of ID_50_ values and statistical analysis. (**h**) Neutralization curves of mouse sera from EBOV GPΔmuc vaccine groups at week 17 against BDBV Uganda pseudovirus. Summary of geometric mean ID_50_ values and statistical analysis. (**i**) ELISA curves of mouse sera from SUDV GPΔmuc trimer and SApNP vaccine groups binding to SUDV GPΔmuc-WL^2^P^4^(1TD0) trimer. (**j**) (Top) Summary of geometric means of EC_50_ titers measured for SUDV GPΔmuc vaccine groups against SUDV GPΔmuc-WL^2^P^4^(1TD0) trimer. (Bottom) Summary of statistical analysis as in (**b**). Note: EC_50_ values at week 2 were derived by setting the minimum/maximum OD_450_ values to 0.0/2.9. (**k**) ELISA curves of mouse sera from SUDV GPΔmuc vaccine groups at week 17 after four immunizations binding to EBOV GPΔmuc-WL^2^P^4^(1TD0) trimer. Summary of geometric mean EC_50_ titers and statistical analysis. (**l**) ELISA curves of mouse sera from SUDV GPΔmuc vaccine groups at week 17 after four immunizations binding to RAVV GPΔmuc-P^2^CT(1TD0) trimer. Summary of A450 values was included. (**m**) Neutralization curves of mouse sera from SUDV GPΔmuc trimer and SApNP vaccine groups against SUDV Gulu pseudovirus. (**n**) (Top) Summary of geometric means of ID_50_ titers measured for EBOV GPΔmuc vaccine groups against EBOV Makona pseudovirus. (Bottom) Summary of statistical analysis as in (**b**). (**o**) Neutralization curves of mouse sera from EBOV GPΔmuc vaccine groups at week 17 against EBOV Makona pseudovirus. Summary of geometric mean ID_50_ values and statistical analysis. (**p**) Neutralization curves of mouse sera from EBOV GPΔmuc vaccine groups at week 17 against BDBV Uganda pseudovirus. Summary of geometric means of ID_50_ values and statistical analysis. (**q**) (Top) ELISA curves of mouse sera from RAVV GPΔmuc trimer vaccine groups binding to RAVV GPΔmuc-P^2^CT(1TD0) trimer. (Bottom) Summary of geometric mean EC_50_ values. Note: EC_50_ values at week 2 were derived by setting the minimum/maximum OD_450_ values to 0.0/2.9. (**r**) ELISA curves of mouse sera from RAVV GPΔmuc vaccine groups at week 17 after four immunizations binding to EBOV GPΔmuc-WL^2^P^4^(1TD0) trimer. Summary of geometric mean EC_50_ titers. (**s**) ELISA curves of mouse sera from RAVV GPΔmuc vaccine groups at week 17 after four immunizations binding to SUDV GPΔmuc-WL^2^P^4^(1TD0) trimer. Summary of A450 values was included. (**t**) Neutralization curves of purified IgG from RAVV GPΔmuc vaccine groups at week 17 against MARV Angola pseudovirus. Summary of IC_50_ values. Left panel: sera from three naive mice showed nonspecific background in MARV Angola pseudovirus assays, suggesting that IgG purification is required to eliminate the nonspecific serum reactivity. (**u**) Neutralization curves of mouse sera from RAVV GPΔmuc vaccine groups at week 17 against orthoebolavirus strains. Error bars represent the difference between duplicate measurements at each concentration for each sample. EC_50_, ID_50_, and IC_50_ values were calculated using GraphPad Prism version 10.3.1. Data were analyzed using one-way ANOVA, followed by Tukey’s multiple comparison post hoc test for each timepoint. For significance, ns (not significant), \**p* < 0.05, \*\**p* < 0.01, \*\*\**p* < 0.001, and \*\*\*\**p* < 0.0001.

**Fig. S11.**
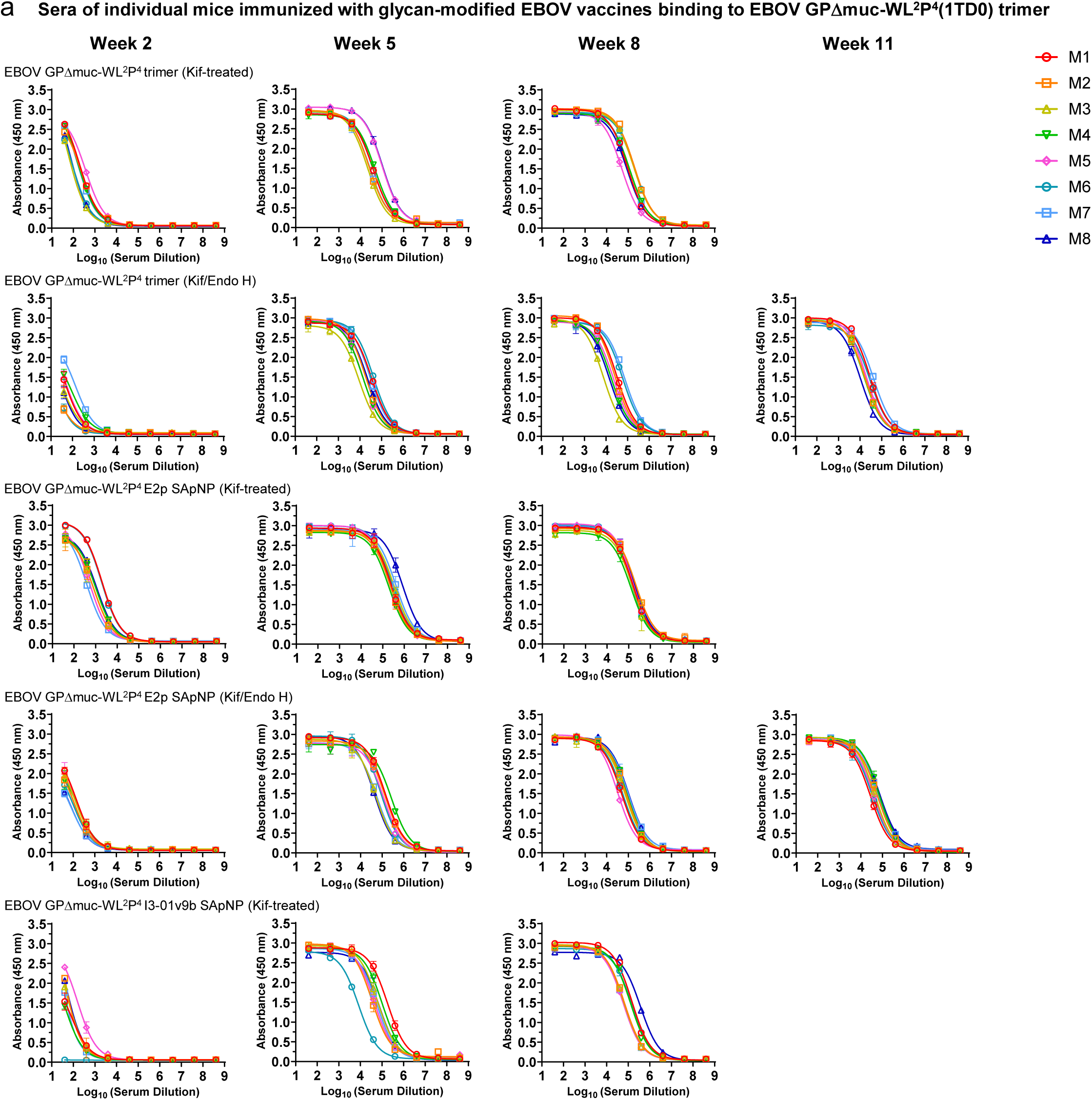
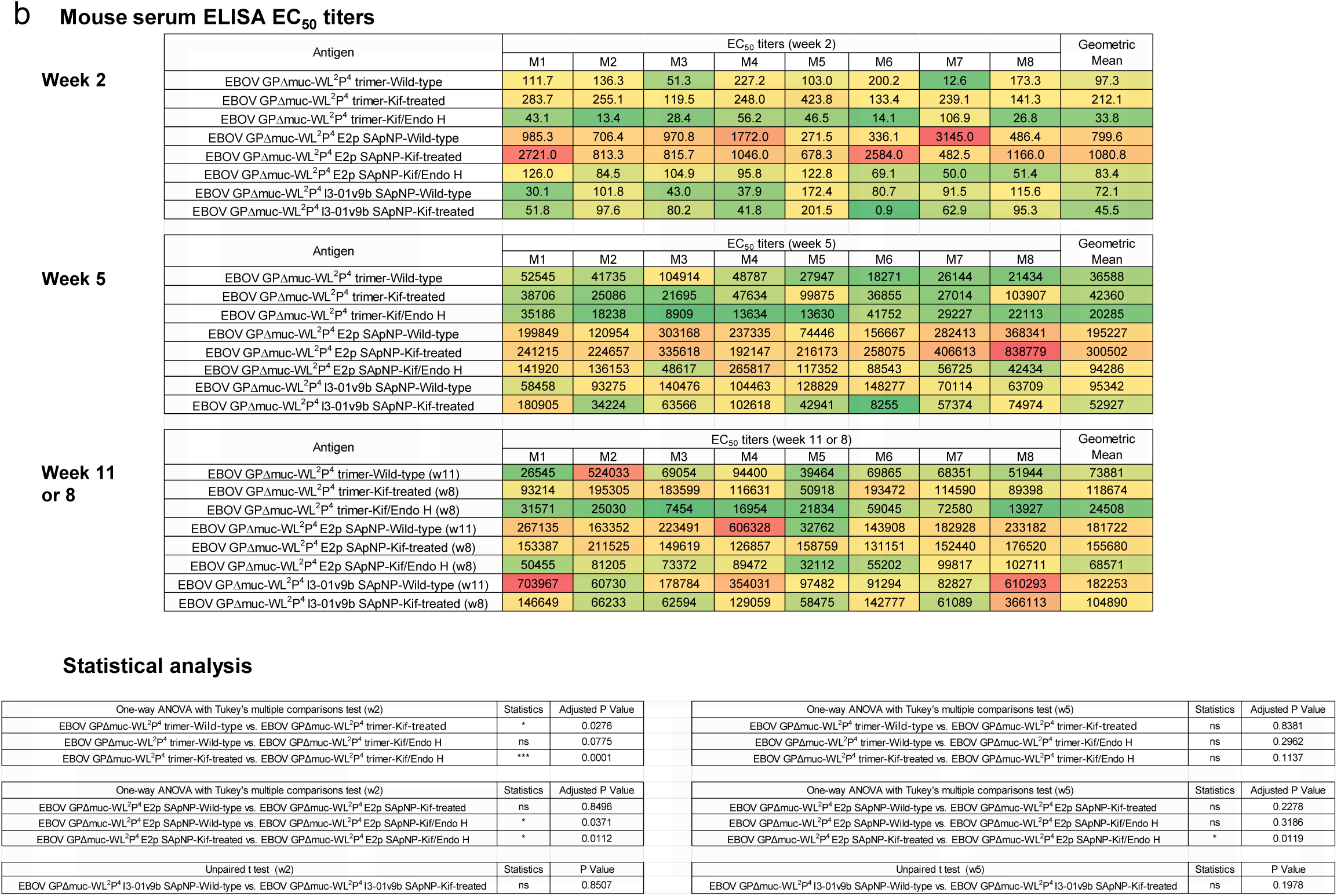
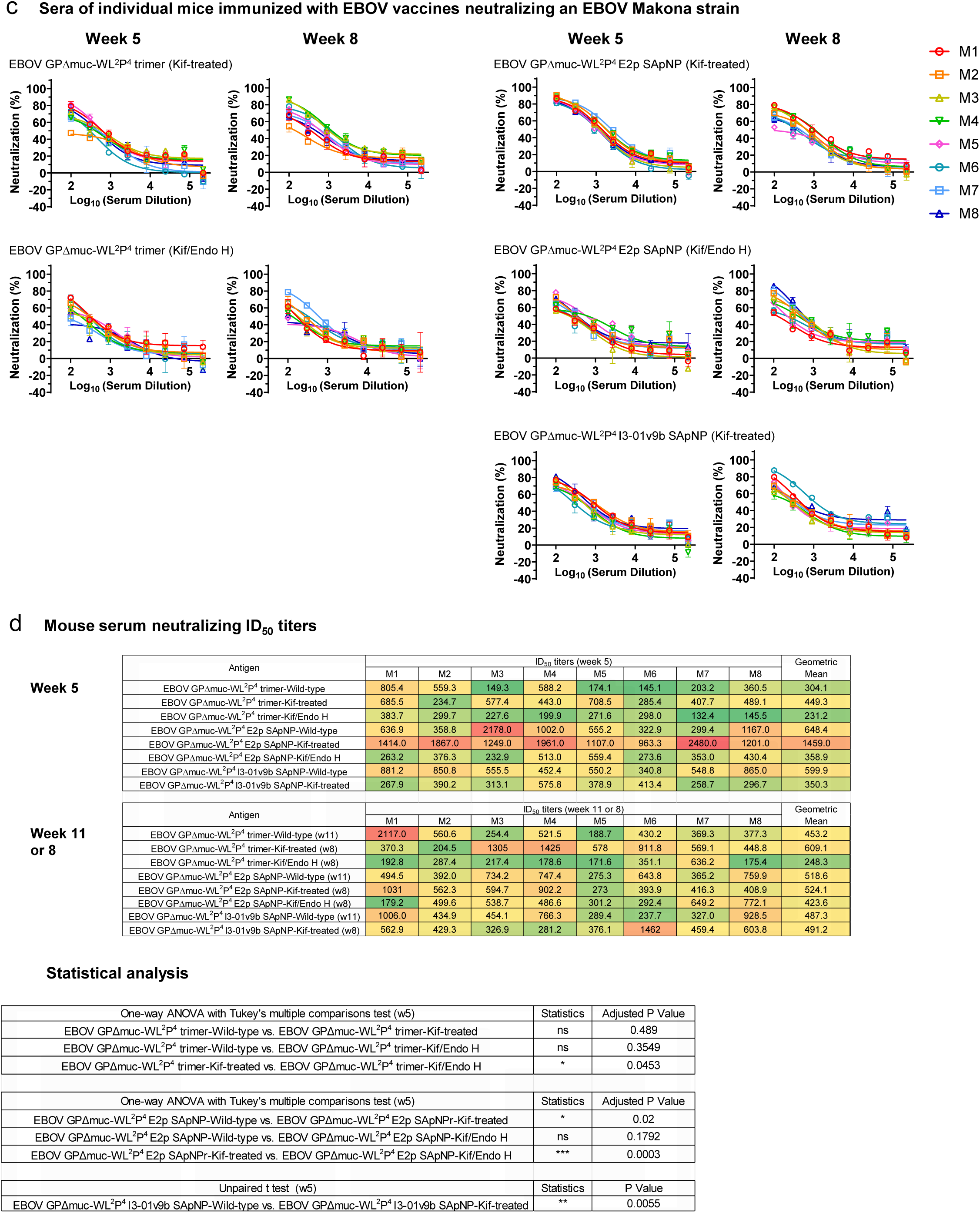
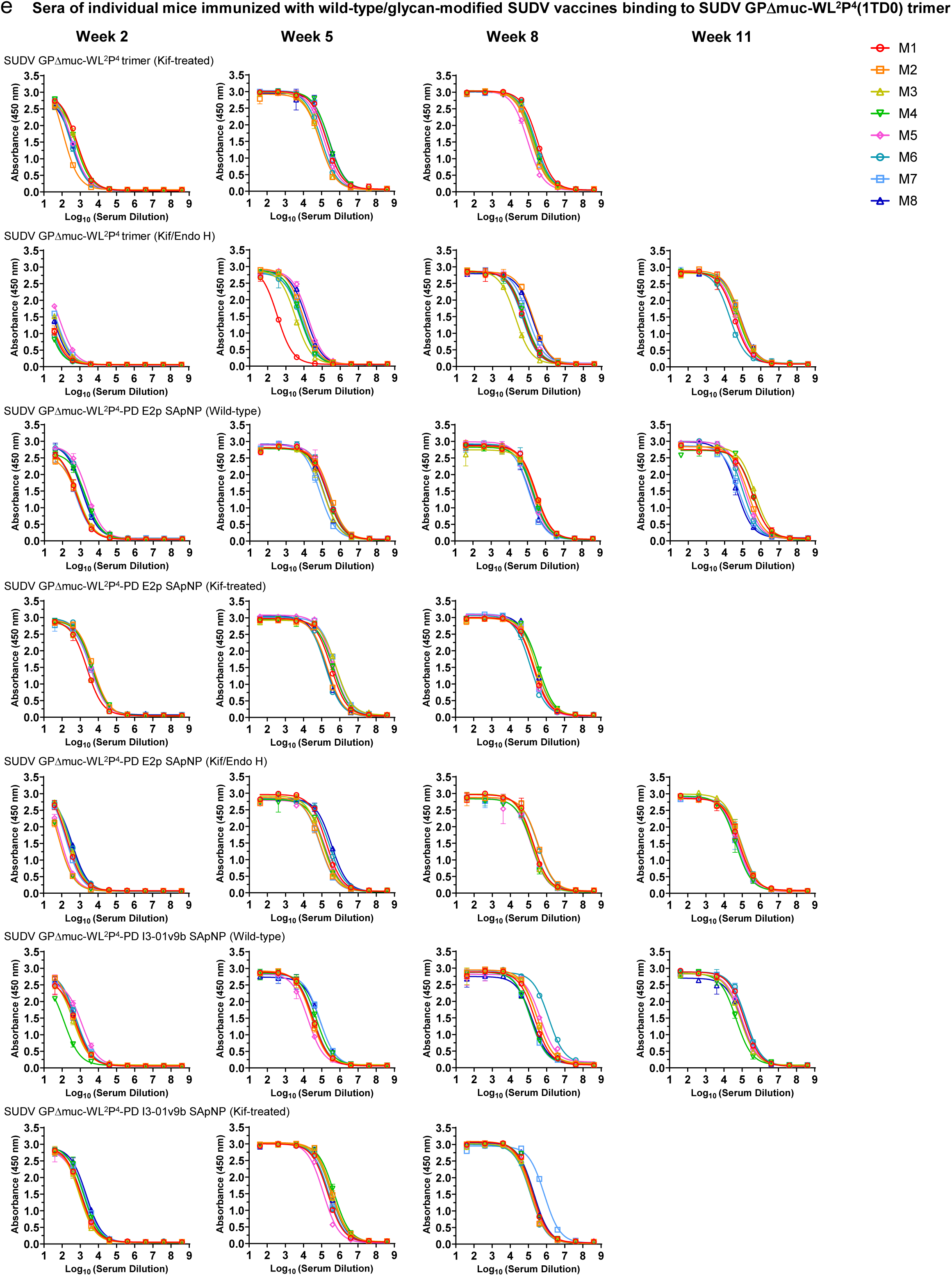
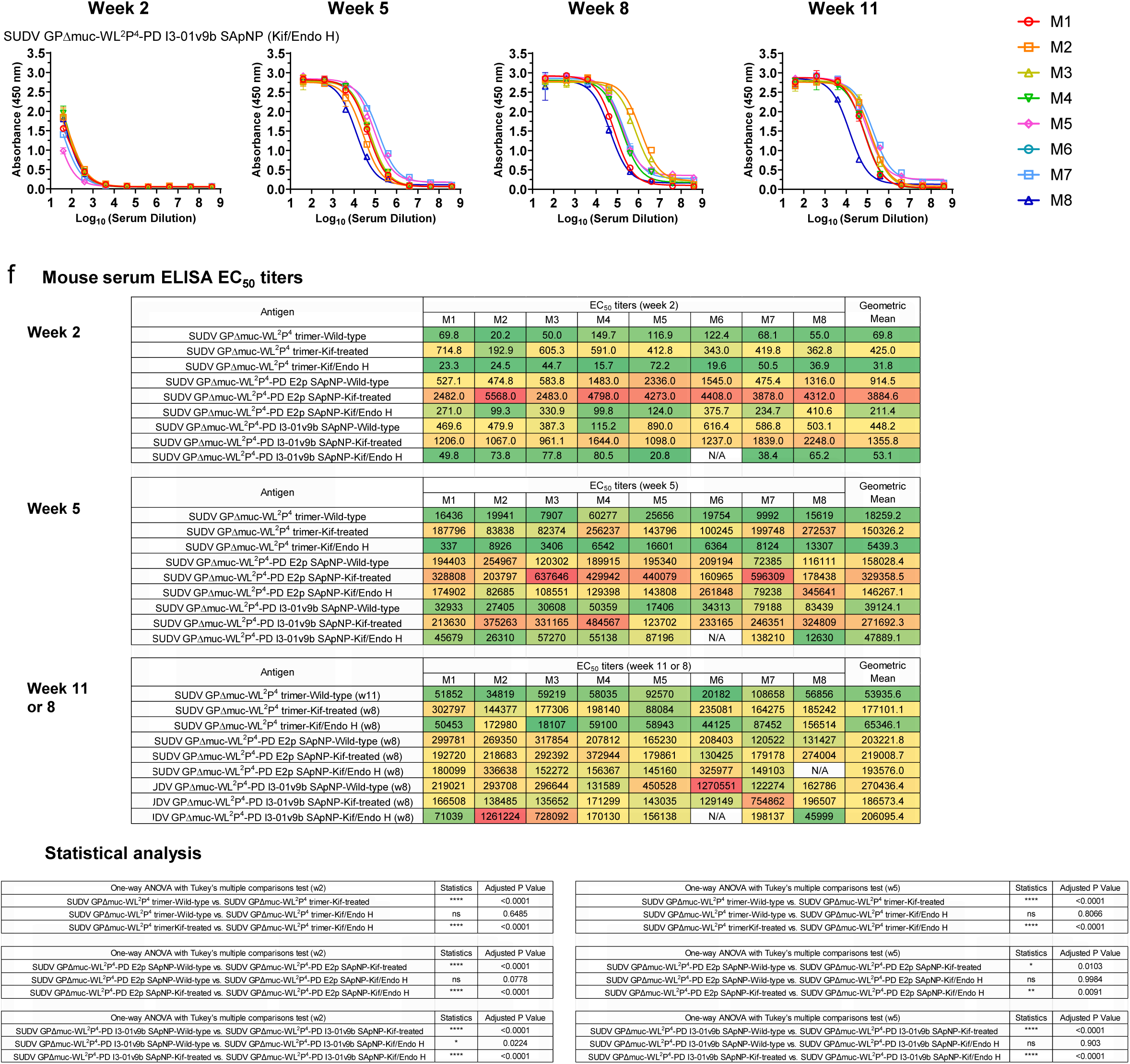
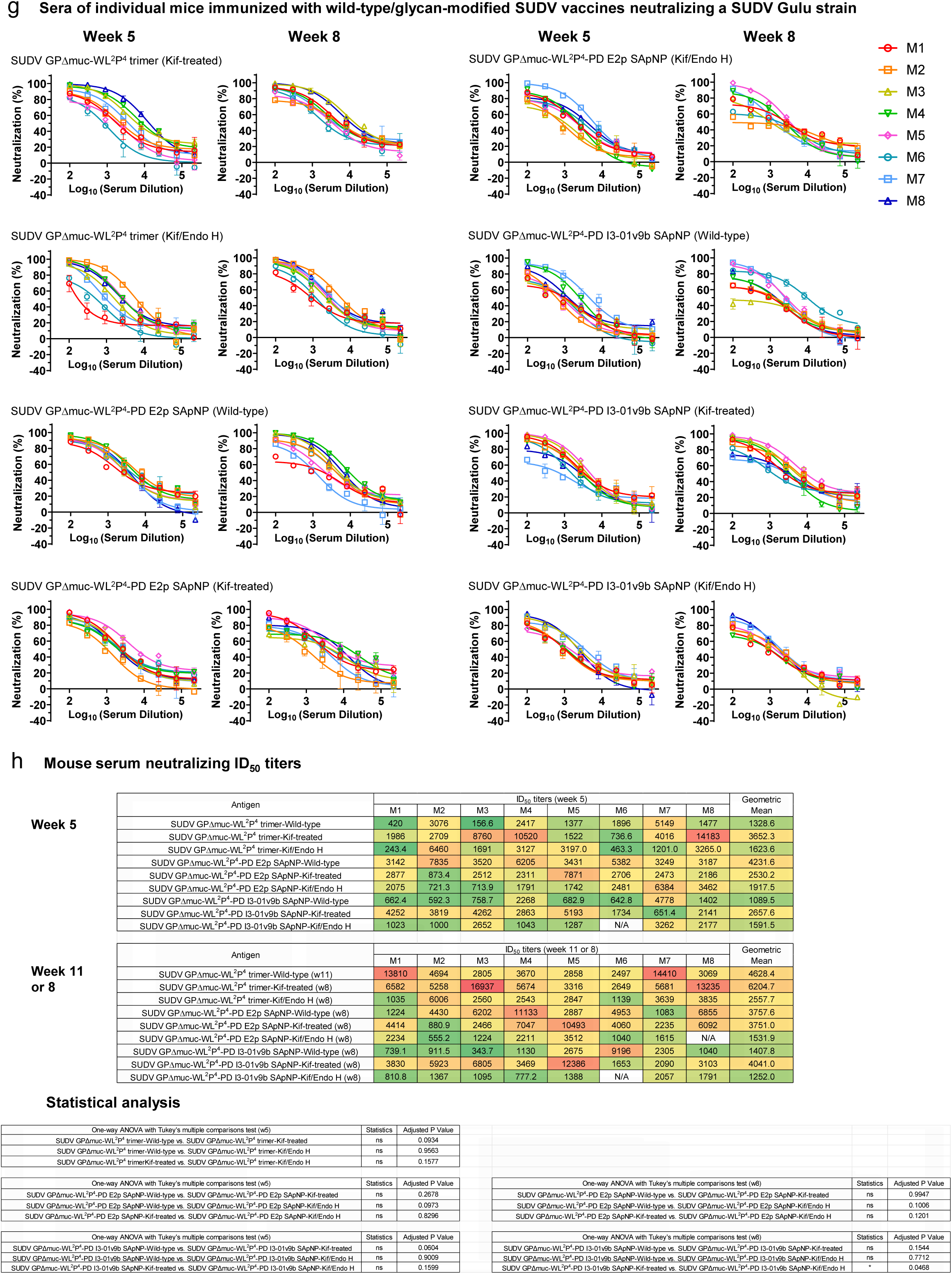
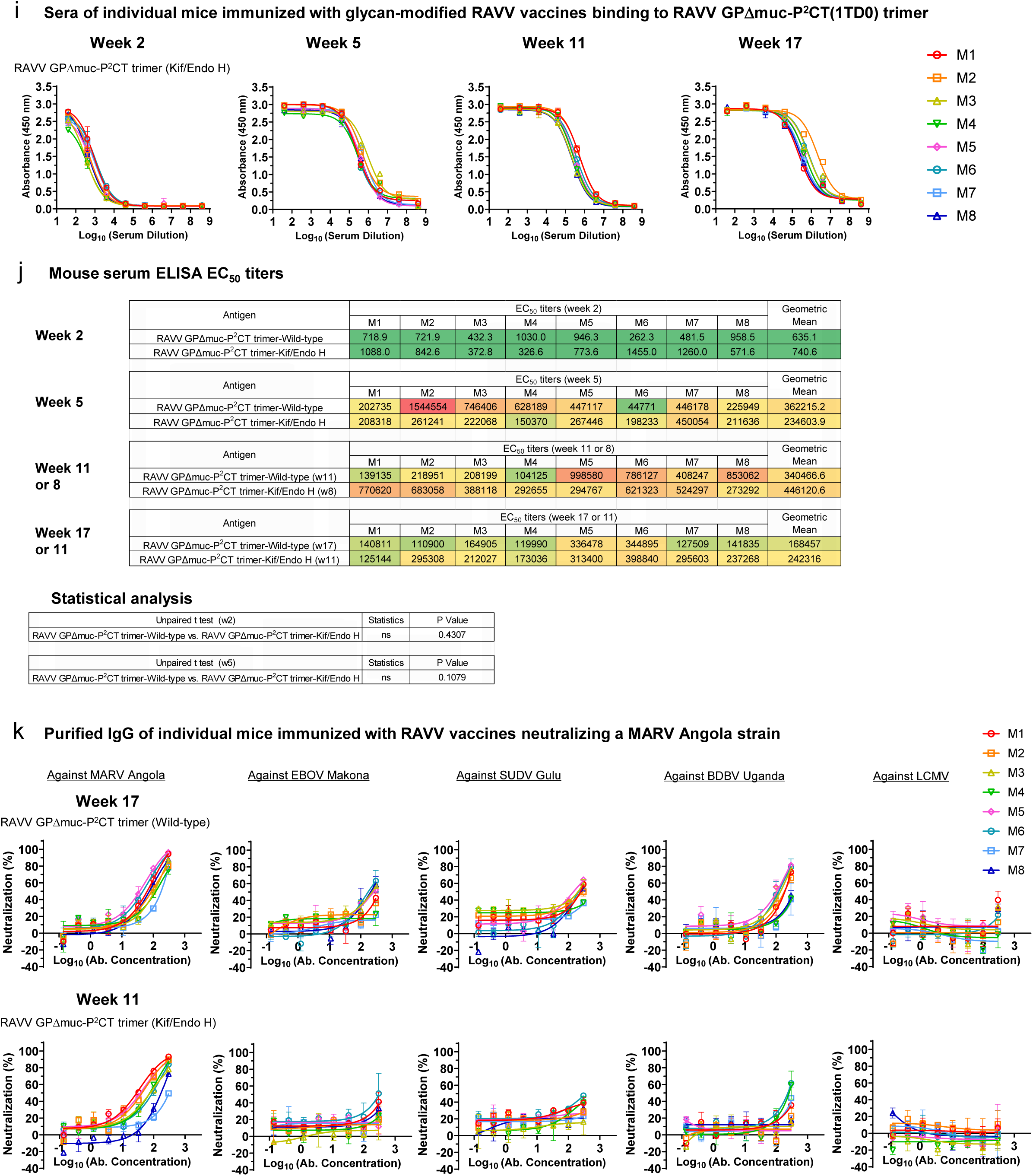
Immunogenicity of glycan-modified EBOV, SUDV, and RAVV GPΔmuc vaccines in mice. (**a**) ELISA curves of mouse sera from glycan-modified EBOV GPΔmuc trimer and SApNP vaccine groups (n = 8 mice/group) binding to EBOV GPΔmuc-WL^2^P^4^(1TD0) trimer. (**b**) (Top) Summary of geometric mean EC_50_ titers measured for glycan-modified EBOV GPΔmuc vaccine groups against EBOV GPΔmuc-WL^2^P^4^(1TD0) trimer. Color coding indicates EC_50_ levels (green to red: low to high binding). (Bottom) Summary of statistical analysis performed for each timepoint. Note: EC_50_ values at week 2 were derived by setting the minimum/maximum OD_450_ values to 0.0/2.8. (**c**) Neutralization curves of mouse sera from glycan-modified EBOV GPΔmuc trimer and SApNP vaccine groups against EBOV Makona pseudovirus. (**d**) (Top) Summary of geometric means of ID_50_ titers measured for EBOV GPΔmuc vaccine groups against EBOV Makona pseudovirus. Color coding: white (no neutralization), green to red (low to high neutralization). Note: ID_50_ values were calculated using %neutralization constraints of 0.0 (min) and 100.0 (max). (Bottom) Summary of statistical analysis. (**e**) ELISA curves of mouse sera from glycan-modified SUDV GPΔmuc trimer and SApNP vaccine groups binding to SUDV GPΔmuc-WL^2^P^4^(1TD0) trimer. (**f**) (Top) Summary of geometric mean EC_50_ titers measured for glycan-modified SUDV GPΔmuc vaccine groups against SUDV GPΔmuc-WL^2^P^4^(1TD0) trimer. (Bottom) Summary of statistical analysis. Note: EC_50_ values at week 2 were derived by setting the minimum/maximum OD_450_ values to 0.0/2.9. (**g**) Neutralization curves of mouse sera from glycan-modified SUDV GPΔmuc trimer and SApNP vaccine groups against SUDV Gulu pseudovirus. (**h**) (Top) Summary of geometric means of ID_50_ titers measured for SUDV GPΔmuc vaccine groups against SUDV Gulu pseudovirus. (Bottom) Summary of statistical analysis. (**i**) ELISA curves of mouse sera from glycan modified RAVV GPΔmuc trimer vaccine groups binding to RAVV GPΔmuc-P^2^CT(1TD0) trimer. (**j**) (Top) Summary of geometric mean EC_50_ titers measured for glycan-modified RAVV GPΔmuc vaccine groups against RAVV GPΔmuc-P^2^CT(1TD0) trimer. (Bottom) Summary of statistical analysis. Note: EC_50_ values at week 2 were derived by setting the minimum/maximum OD_450_ values to 0.0/2.9. (**k**) Neutralization curves of purified mouse IgG from glycan-modified RAVV GPΔmuc vaccine groups after four immunizations at week 17 or week 11 against MARV Angola or orthoebolavirus strains. Lymphocytic choriomeningitis virus (LCMV) was included as a negative control to confirm the cross-NAb responses measured using purified IgG. Error bars represent the difference between duplicate measurements at each concentration for each sample. EC_50_ and ID_50_ values were calculated using GraphPad Prism version 10.3.1. Data were analyzed using one-way ANOVA, followed by Tukey’s multiple comparison post hoc test for each timepoint. Two-tailed unpaired t-tests were used to compare the geometric means between two independent groups. Statistical significance was defined as follows: ns (not significant), *p < 0.05, **p < 0.01, ***p < 0.001, and \*\*\*\**p* < 0.0001.

**Fig. S12.**
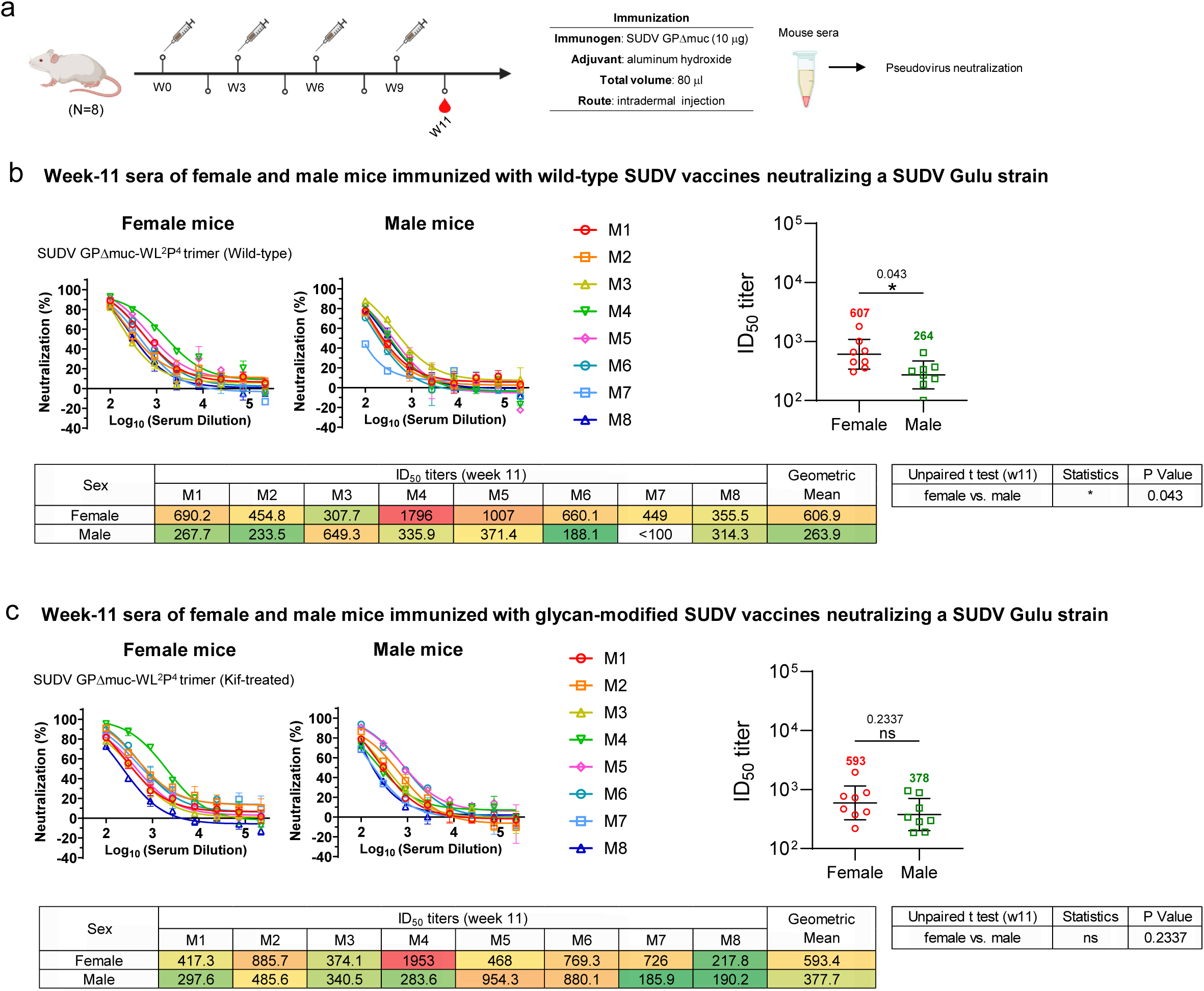
Influence of sex on immunogenicity of SUDV GPΔmuc-induced neutralizing antibody responses in mice. (**a**) Schematic illustration of the mouse immunization regimen for SUDV GPΔmuc trimer vaccines (n = 8 mice per group). Each mouse received 80 μl of a vaccine antigen/aluminum hydroxide (AH) adjuvant mix containing 10 μg of SUDV antigen and 40 μl of AH. Mice were immunized via the intradermal route through four footpad injections at weeks 0, 3, 6, and 9, with 3-week intervals between doses. (**b**) Neutralization curves of female and male mouse sera from the wild-type SUDV GPΔmuc trimer group at week 11 (after four immunizations), tested against SUDV Gulu pseudovirus. (**c**) Neutralization curves of female and male mouse sera from the glycan-modified SUDV GPΔmuc trimer group under the same conditions. Comparison of NAb responses (by ID₅₀ titers) between female and male mice is shown, along with a summary of geometric mean ID₅₀ values for each group against SUDV Gulu pseudovirus. Error bars represent the difference between duplicate measurements at each concentration for each sample. ID_50_ values were calculated using GraphPad Prism version 10.3.1. Two-tailed unpaired t-tests were used to compare the geometric means between two independent groups. Statistical significance was defined as follows: ns (not significant), *p < 0.05.

**Table S1.**
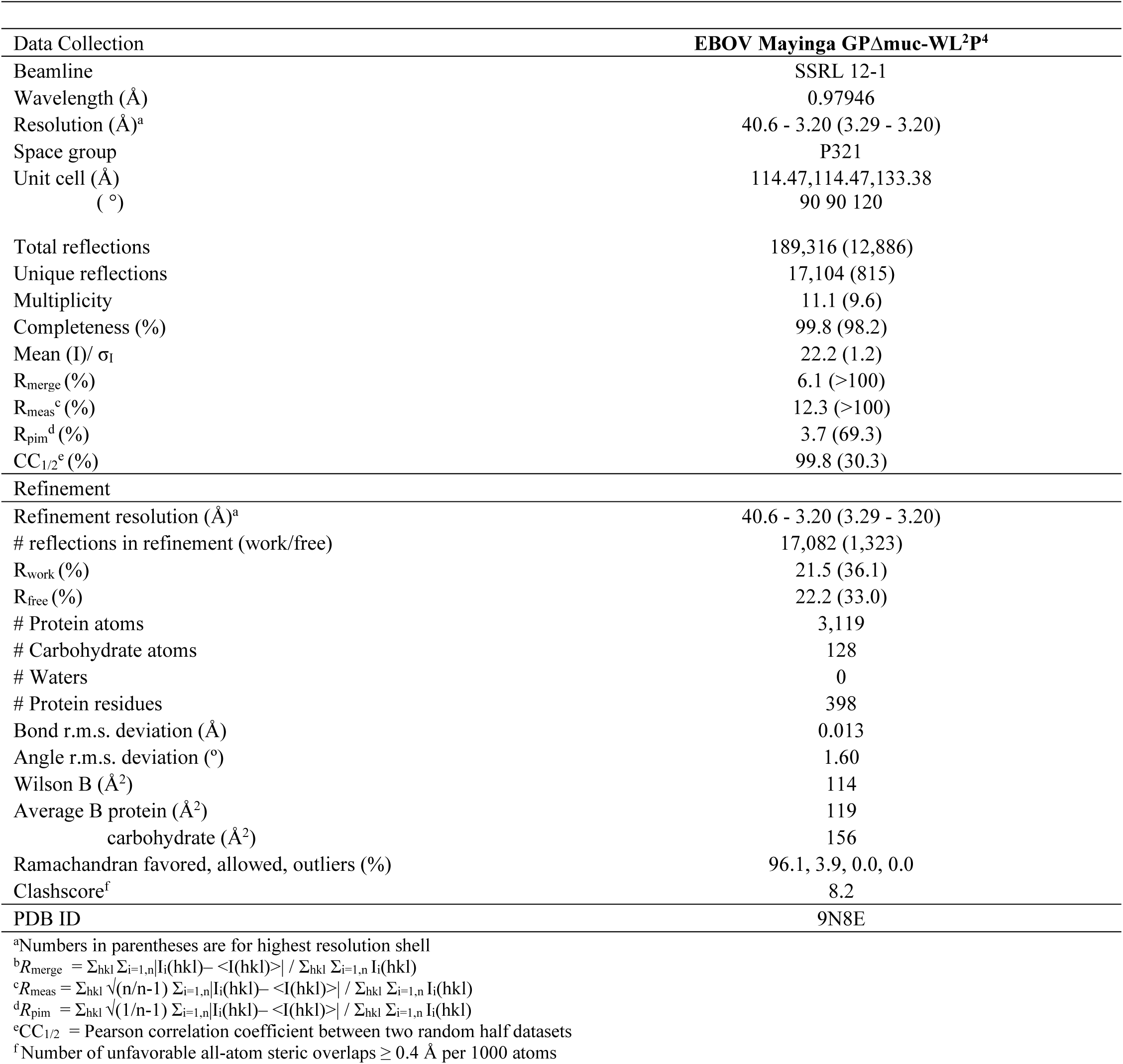
Data collection and refinement statistics for EBOV Mayinga GPΔmuc-WL^2^P^4^.

**Table S2.**
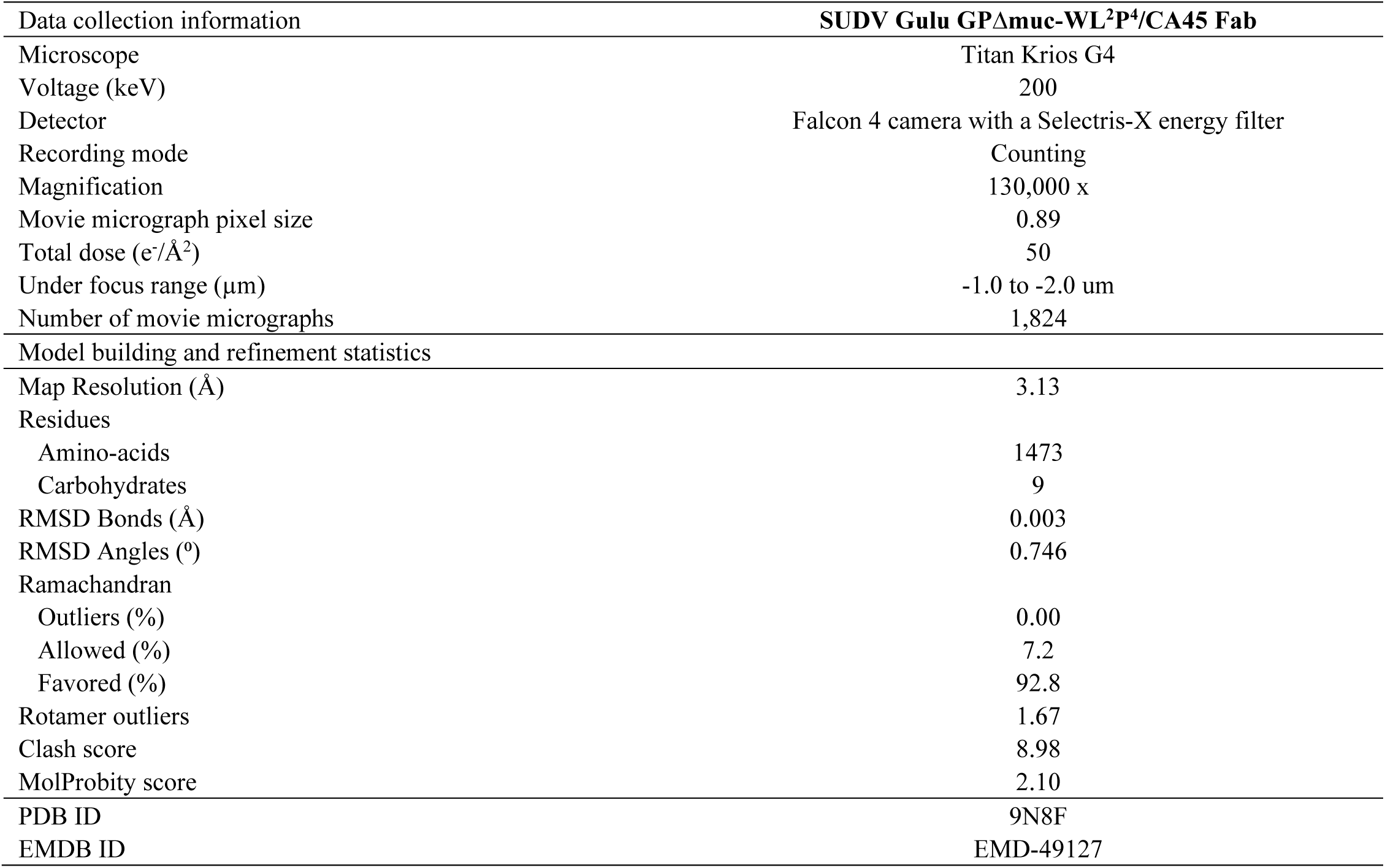
Cryo-EM data collection information, model building, and refinement statistics.

