## Supplementary Information for "Rational design of next-generation filovirus vaccines through glycoprotein stabilization, nanoparticle display, and glycan modification"

**Fig. S1**

a

**>EBOV Mayinga GPΔmuc-WL$^2$P$^2$-foldon-[His$_6$]**

MGVTGILQLPRDRFKRTSFFLWVIILFQRTFSIPLGVIHNSALQVSDVDKLVCRDKLSSTNQLRSVGLNLEGNGVATDVPSATKRWGFRSGVPPKVVNYEAGEWAENC
YNLEIKKPDGSECLPAAPDGIRGFPRCRYVHKVSGTGPCAGDFAFHKEGAFFLYDRLASTVIYRGTTFAEGVVAFLILPQAKKDFFSSHPLREPVNATEDPSSGYYST
TIRYQATGFGTNETEYLFEVDNLTYVQLESRFTPQFLLQLNETIYTSGKRSNTTGKLIWKVNPEIDTTIGEWAFWETKKNLTRKIRSEELSFTVVSTHHQDTGEESAS
SGKLGLITNTIAGVAGLITGGRRTRREAIVNAQPKCNPNLHYWTTQDEGAAIGLAWIPYFGPAAEGIYIEGLMHNQDGLICGLRQLANETTQALQLFLRATPELRTFS
ILNRKAIDFLLQRWGGTCHILGPDCCIEPHDLTKNITDKIDQIIHDFVDKTLPDASGYIPEAPRDGQAYVRKDGEWVLLSTFL[GSHHHHHH]

**>EBOV Mayinga GPΔmuc-WL$^2$P$^4$-foldon-[His$_6$]**

MGVTGILQLPRDRFKRTSFFLWVIILFQRTFSIPLGVIHNSALQVSDVDKLVCRDKLSSTNQLRSVGLNLEGNGVATDVPSATKRWGFRSGVPPKVVNYEAGEWAENC
YNLEIKKPDGSECLPAAPDGIRGFPRCRYVHKVSGTGPCAGDFAFHKEGAFFLYDRLASTVIYRGTTFAEGVVAFLILPQAKKDFFSSHPLREPVNATEDPSSGYYST
TIRYQATGFGTNETEYLFEVDNLTYVQLESRFTPQFLLQLNETIYTSGKRSNTTGKLIWKVNPEIDTTIGEWAFWETKKNLTRKIRSEELSFTVVSTHHQDTGEESAS
SGKLGLITNTIAGVAGLITGGRRTRREAIVNAQPKCNPNLHYWTTQDEGAAIGLAWIPYFGPAAEGIYIEGLMHNQDGLICGLRQLANETTQALQLFLRATTEPRTFS
ILNRKAIDFLLQRWGGTCHILGPDCCIEPHDLTKNITDKIDQIIHDFVDKTLPDASGYIPEAPRDGQAYVRKDGEWVLLSTFL[GSHHHHHH]

**>SUDV Gulu GPΔmuc-WL$^2$P$^2$-foldon-[His$_6$]**

MGILPSPGMPALLSLVSLLSVLLMGCVAEMPLGVVTNSTLEVTEIDQLVCKDHLASTDQLKSVGLNLEGSGVSTDIPSATKRWGFRSGVPPKVVSYEAGEWAENCYNL
EIKKPDGSECLPPPPDGVRGFPRCRYVHKAQGTGPCPGDYAFHKDGAFFLYDRLASTVIYRGVNFAEGVIAFLILAKPKETFLQSPPIREAVNYTENTSSYYATSYLE
YEIENFGAQHSTTLFKIDNNTFVRLDRPHTPQFLFQLNDTIHLHQQLSNTTGRLIWTLDANINADIGEWAFWENKKNLSEQLRGEELSFEALSLNETEDDDAASSSTS
NGLITSTVTGILGSLGLRKRSRRQTNTKATGKCNPNLHYWTAQEQHNAAGIAWIPYFGPGAEGIYTEGLMHNQNALVCGLRQLANETTQALQLFLRATPELRTYTILN
RKAIDFLLRRWGGTCRILGPDCCIEPHDLTKNITDKINQIIHDFIDNPLPNASGYIPEAPRDGQAYVRKDGEWVLLSTFL[GSHHHHHH]

**>SUDV Gulu GPΔmuc-WL$^2$P$^4$-foldon-[His$_6$]**

MGILPSPGMPALLSLVSLLSVLLMGCVAEMPLGVVTNSTLEVTEIDQLVCKDHLASTDQLKSVGLNLEGSGVSTDIPSATKRWGFRSGVPPKVVSYEAGEWAENCYNL
EIKKPDGSECLPPPPDGVRGFPRCRYVHKAQGTGPCPGDYAFHKDGAFFLYDRLASTVIYRGVNFAEGVIAFLILAKPKETFLQSPPIREAVNYTENTSSYYATSYLE
YEIENFGAQHSTTLFKIDNNTFVRLDRPHTPQFLFQLNDTIHLHQQLSNTTGRLIWTLDANINADIGEWAFWENKKNLSEQLRGEELSFEALSLNETEDDDAASSSTS
NGLITSTVTGILGSLGLRKRSRRQTNTKATGKCNPNLHYWTAQEQHNAAGIAWIPYFGPGAEGIYTEGLMHNQNALVCGLRQLANETTQALQLFLRATTEPRTYTILN
RKAIDFLLRRWGGTCRILGPDCCIEPHDLTKNITDKINQIIHDFIDNPLPNASGYIPEAPRDGQAYVRKDGEWVLLSTFL[GSHHHHHH]

**>BDBV R4386L GPΔmuc-WL$^2$P$^2$-foldon-[His$_6$]**

MGILPSPGMPALLSLVSLLSVLLMGCVAEIPLGVVHNNTLQVSDIDKLVCRDKLSSTSQLKSVGLNLEGNGVATDVPTATKRWGFRAGVPPKVVNYEAGEWAENCYNL
DIKKADGSECLPEAPEGVRGFPRCRYVHKVSGTGPCPEGYAFHKEGAFFLYDRLASTIIYRSTTFSEGVVAFLILPETKKDFFQSPPLHEPANMTTDPSSYYHTVTLN
YVADNFGTNMTNFLFQVDHLTYVQLEPRFTPQFLVQLNETIYTNGRRSNTTGTLIWKVNPTVDTGVGEWAFWENKKNFTKTLSSEELSVIFVPRAQDPGSNDISESTE
PGPLTNTTRGAANLLTGSRRTRREITLRTQAKCNPNLHYWTTQDEGAAIGLAWIPYFGPAAEGIYTEGIMHNQNGLICGLRQLANETTQALQLFLRATPELRTFSILN
RKAIDFLLQRWGGTCHILGPDCCIEPHDLTKNITDKIDQIIHDFIDKPLPDASGYIPEAPRDGQAYVRKDGEWVLLSTFL[GSHHHHHH]

**>BDBV R4386L GPΔmuc-WL$^2$P$^4$-foldon-[His$_6$]**

MGILPSPGMPALLSLVSLLSVLLMGCVAEIPLGVVHNNTLQVSDIDKLVCRDKLSSTSQLKSVGLNLEGNGVATDVPTATKRWGFRAGVPPKVVNYEAGEWAENCYNL
DIKKADGSECLPEAPEGVRGFPRCRYVHKVSGTGPCPEGYAFHKEGAFFLYDRLASTIIYRSTTFSEGVVAFLILPETKKDFFQSPPLHEPANMTTDPSSYYHTVTLN
YVADNFGTNMTNFLFQVDHLTYVQLEPRFTPQFLVQLNETIYTNGRRSNTTGTLIWKVNPTVDTGVGEWAFWENKKNFTKTLSSEELSVIFVPRAQDPGSNDISESTE
PGPLTNTTRGAANLLTGSRRTRREITLRTQAKCNPNLHYWTTQDEGAAIGLAWIPYFGPAAEGIYTEGIMHNQNGLICGLRQLANETTQALQLFLRATTEPRTFSILN
RKAIDFLLQRWGGTCHILGPDCCIEPHDLTKNITDKIDQIIHDFIDKPLPDASGYIPEAPRDGQAYVRKDGEWVLLSTFL[GSHHHHHH]

b **SEC profile of 6 orthoebolavirus GPΔmuc constructs expressed in HEK293F cells and purified by nickel and IAC columns**

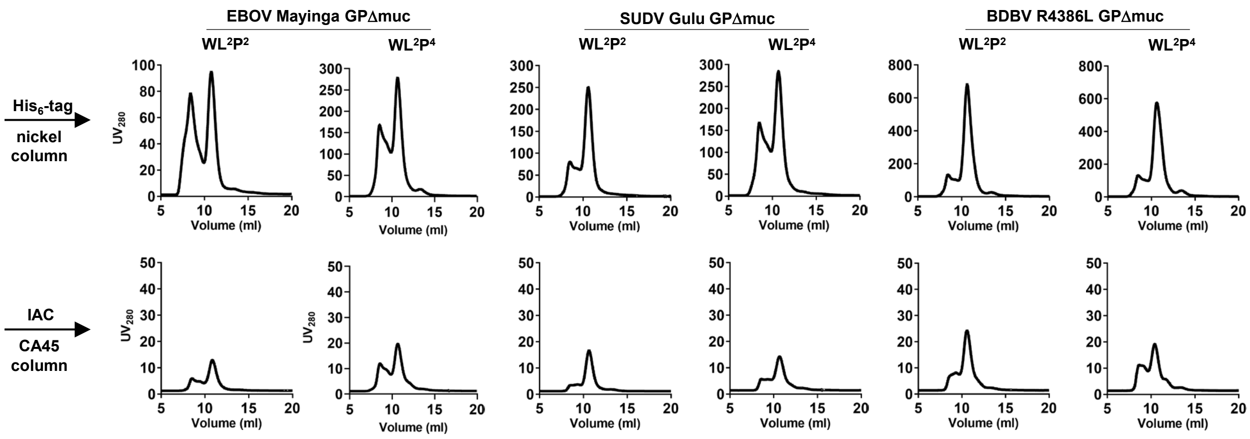

c

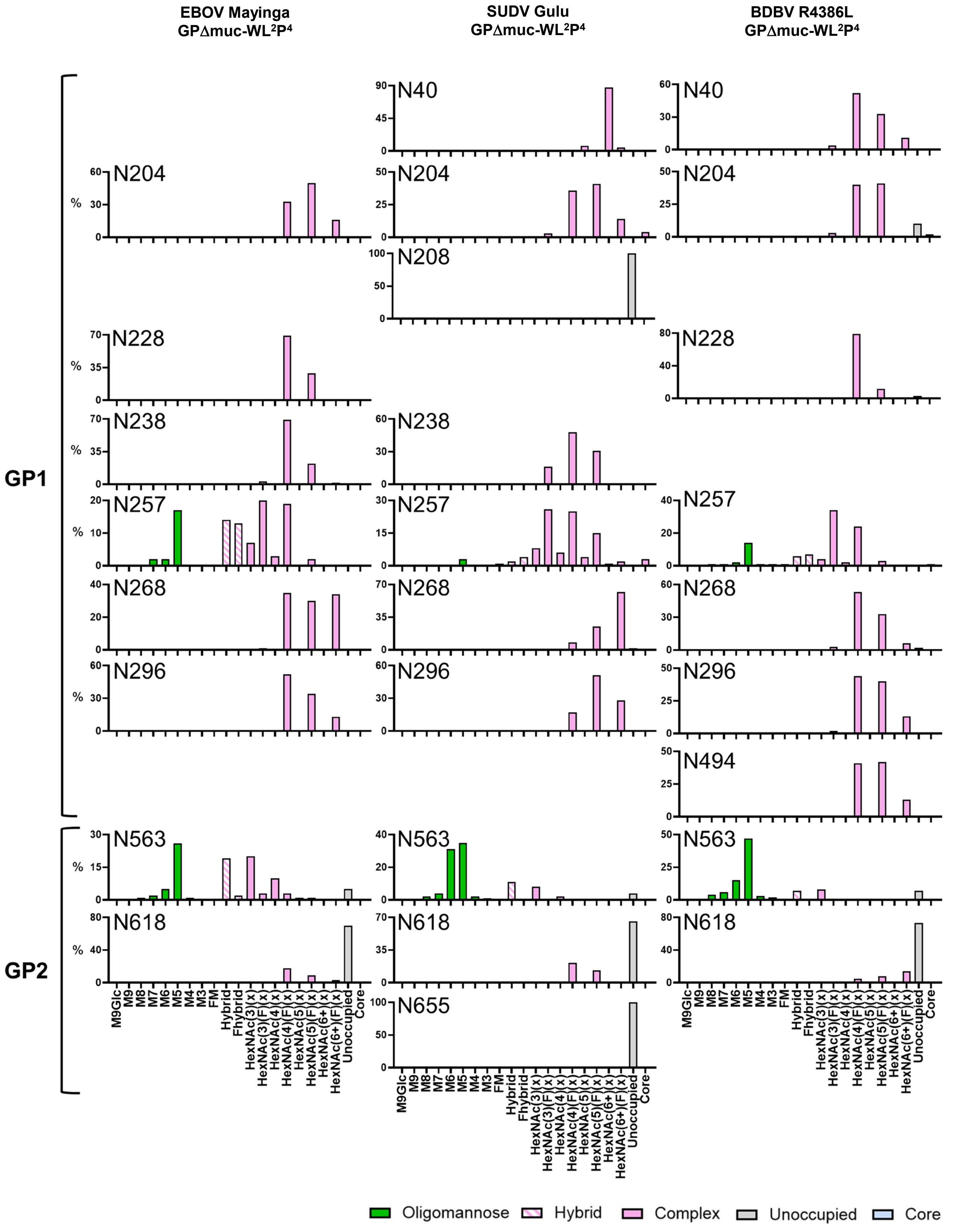
EBOV Mayinga
GPΔmuc-WL²P⁴
SUDV Gulu
GPΔmuc-WL²P⁴
BDBV R4386L
GPΔmuc-WL²P⁴
GP1
GP2
N40
N204
N208
N228
N238
N257
N268
N296
N494
N563
N618
N655
%
M9Glc
M9
M8
M7
M6
M5
M4
M3
FM
Hybrid
Fhybrid
HexNAc(3)(x)
HexNAc(3)(F)(x)
HexNAc(4)(x)
HexNAc(4)(F)(x)
HexNAc(5)(x)
HexNAc(5)(F)(x)
HexNAc(6+)(x)
HexNAc(6+)(F)(x)
Unoccupied
Core
Oligomannose
Hybrid
Complex
Unoccupied
Core

**Fig. S1**

d Flowchart of image processing and model building protocol

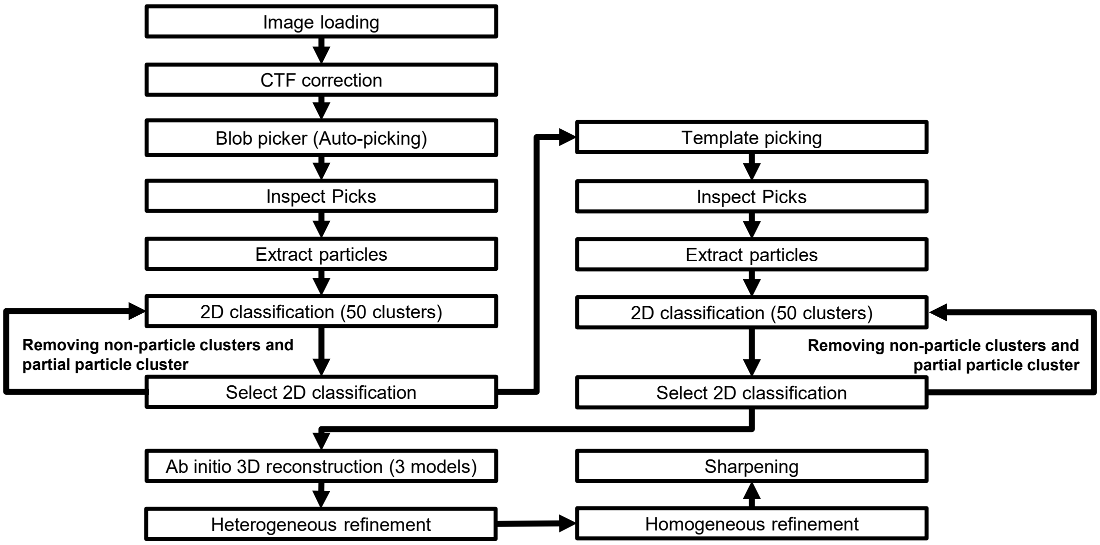


e **Selected 2D classifications for 3D modeling of EBOV Mayinga GPΔmuc**

**GPΔmuc-WL²P⁴ trimer/ADI-15878 Fab complex**

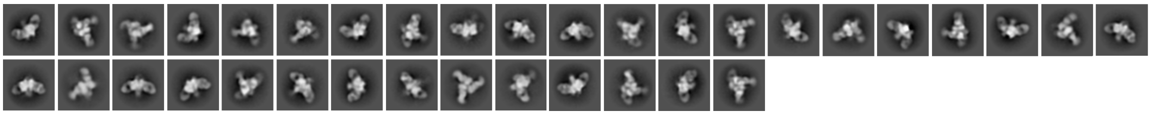

**GPΔmuc-WL²P⁴ trimer**

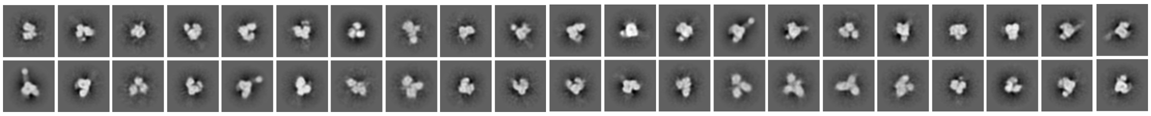

f **Selected 2D classifications for 3D modeling of SUDV Gulu GPΔmuc**

**GPΔmuc-WL²P⁴ trimer/ADI-15878 Fab complex**

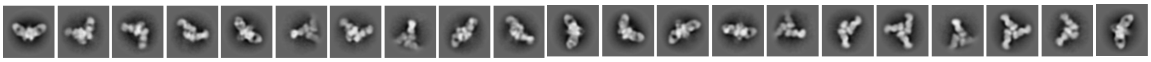

**GPΔmuc-WL²P⁴ trimer**

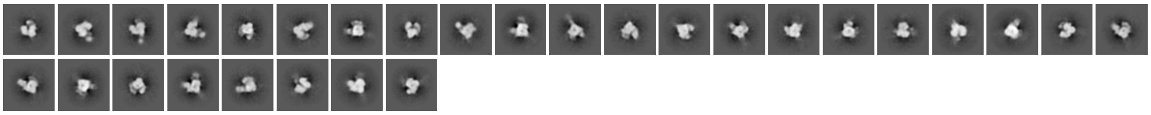

g **Selected 2D classifications for 3D modeling of BDBV R3486L GPΔmuc**

**GPΔmuc-WL²P⁴ trimer/ADI-15878 Fab complex**

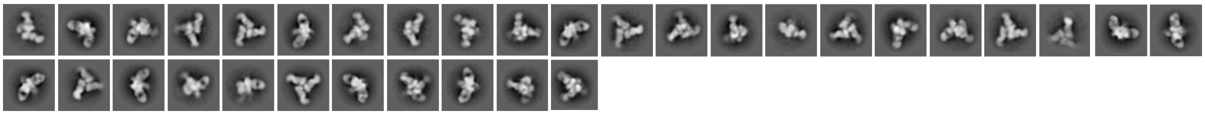

**GPΔmuc-WL²P⁴ trimer**

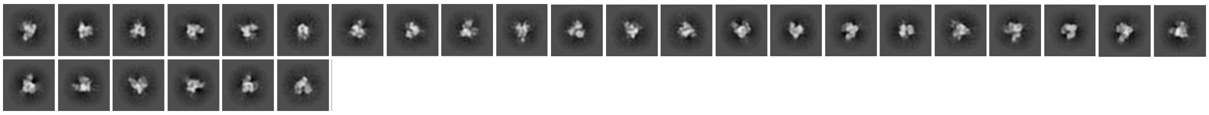

#### h ELISA analysis of 6 rationally designed orthoebolavirus GPΔmuc trimers binding to 5 filovirus NAbs

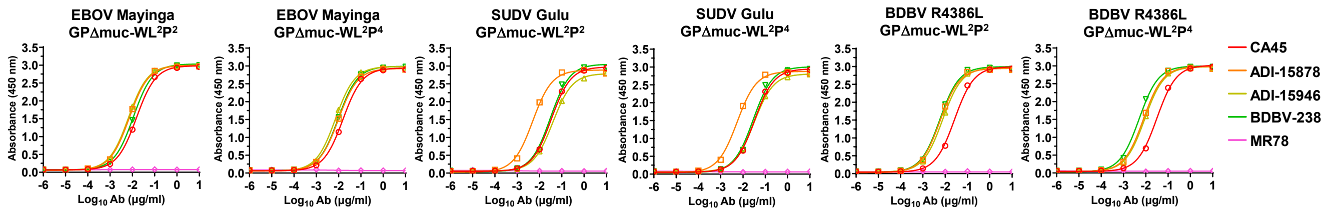


| EBOV Mayinga GPΔmuc-WL²P² | | | | | | | | | |
|---|---|---|---|---|---|---|---|---|---|
| CA45 | | ADI-15878 | | ADI-15946 | | BDBV-289 | | MR78 | |
| EC50 | STD | EC50 | STD | EC50 | STD | EC50 | STD | EC50 | STD |
| 0.01520 | 0.00023 | 0.00727 | 0.00072 | 0.00669 | 0.00017 | 0.01109 | 0.00029 | N.D. | N.D. |

| EBOV Mayinga GPΔmuc-WL²P⁴ | | | | | | | | | |
|---|---|---|---|---|---|---|---|---|---|
| CA45 | | ADI-15878 | | ADI-15946 | | BDBV-289 | | MR78 | |
| EC50 | STD | EC50 | STD | EC50 | STD | EC50 | STD | EC50 | STD |
| 0.015639 | 0.000587 | 0.009683 | 0.001136 | 0.007233 | 0.000747 | 0.009632 | 0.00102 | N.D. | N.D. |

| SUDV Gulu GPΔmuc-WL²P² | | | | | | | | | |
|---|---|---|---|---|---|---|---|---|---|
| CA45 | | ADI-15878 | | ADI-15946 | | BDBV-289 | | MR78 | |
| EC50 | STD | EC50 | STD | EC50 | STD | EC50 | STD | EC50 | STD |
| 0.0338 | 2.83E-05 | 0.004959 | 7.07E-06 | 0.039574 | 0.002213 | 0.030734 | 0.000834 | N.D. | N.D. |

| SUDV Gulu GPΔmuc-WL²P⁴ | | | | | | | | | |
|---|---|---|---|---|---|---|---|---|---|
| CA45 | | ADI-15878 | | ADI-15946 | | BDBV-289 | | MR78 | |
| EC50 | STD | EC50 | STD | EC50 | STD | EC50 | STD | EC50 | STD |
| 0.032764 | 0.001209 | 0.005461 | 0.000156 | 0.031179 | 0.002638 | 0.028805 | 0.000792 | N.D. | N.D. |

| BDBV R4386L GPΔmuc-WL²P² | | | | | | | | | |
|---|---|---|---|---|---|---|---|---|---|
| CA45 | | ADI-15878 | | ADI-15946 | | BDBV-289 | | MR78 | |
| EC50 | STD | EC50 | STD | EC50 | STD | EC50 | STD | EC50 | STD |
| 0.026339 | 0.001499 | 0.005969 | 0.000224 | 0.007626 | 0.000158 | 0.005773 | 3.39E-05 | N.D. | N.D. |

| BDBV R4386L GPΔmuc-WL²P⁴ | | | | | | | | | |
|---|---|---|---|---|---|---|---|---|---|
| CA45 | | ADI-15878 | | ADI-15946 | | BDBV-289 | | MR78 | |
| EC50 | STD | EC50 | STD | EC50 | STD | EC50 | STD | EC50 | STD |
| 0.031360 | 0.000170 | 0.007866 | 0.000318 | 0.008934 | 0.000208 | 0.005026 | 0.000132 | N.D. | N.D. |

#### i BLI analysis of 6 rationally designed orthoebolavirus GPΔmuc trimers binding to 5 filovirus NAbs

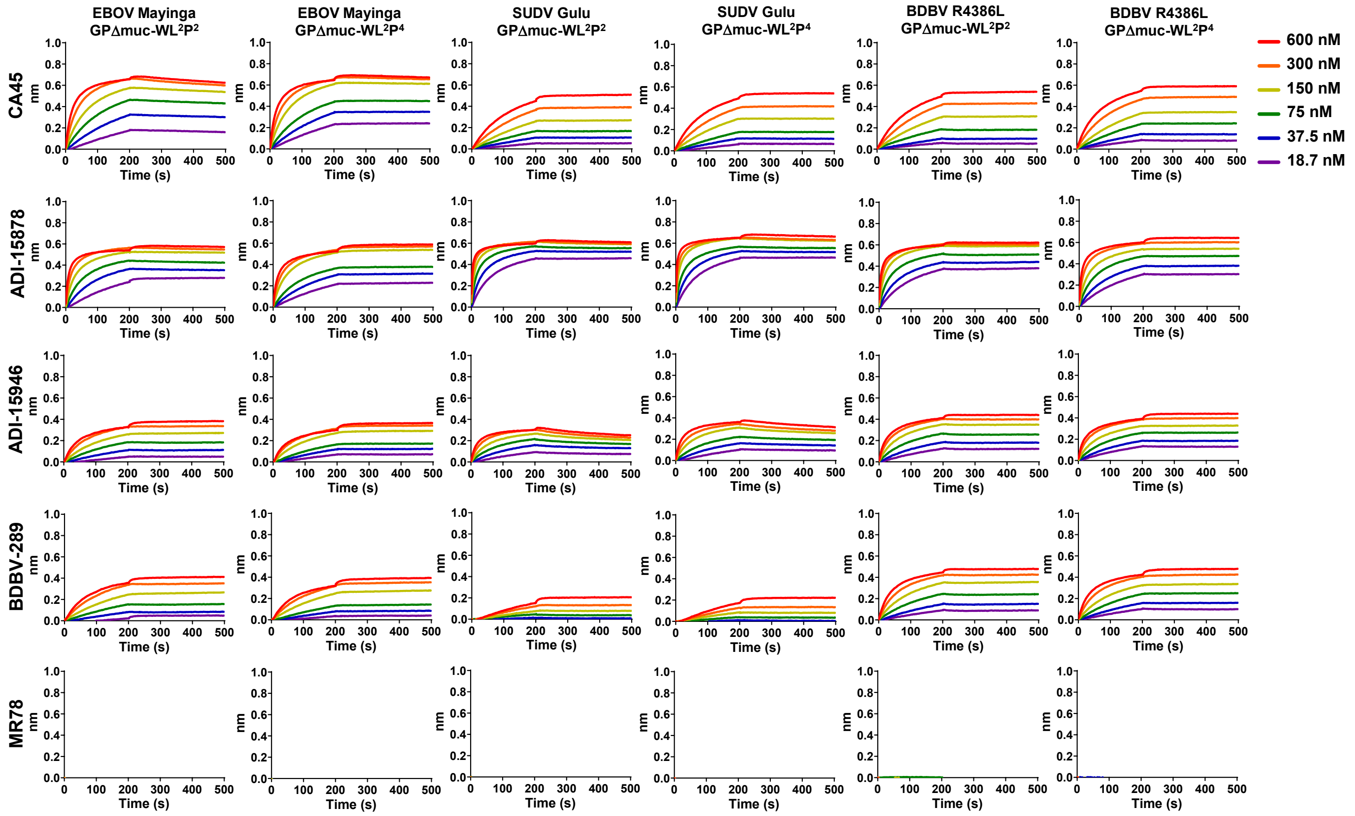


***K**D* **values for 6 rationally designed orthoebolavirus GPΔmuc trimers binding to 5 filovirus NAbs (nM)** [a]

| | CA45 | ADI-15878 | ADI-15946 | BDBV-289 | MR78 |
|---|---|---|---|---|---|
| EBOV Mayinga GPΔmuc-WL²P² | 2.62 | <1.0E-3 | <1.0E-3 | <1.0E-3 | — [b] |
| EBOV Mayinga GPΔmuc-WL²P⁴ | <1.0E-3 | <1.0E-3 | <1.0E-3 | <1.0E-3 | — [b] |
| SUDV Gulu GPΔmuc-WL²P² | <1.0E-3 | <1.0E-3 | 7.22 | <1.0E-3 | — [b] |
| SUDV Gulu GPΔmuc-WL²P⁴ | <1.0E-3 | <1.0E-3 | 4.89 | <1.0E-3 | — [b] |
| BDBV R4386L GPΔmuc-WL²P² | <1.0E-3 | <1.0E-3 | <1.0E-3 | <1.0E-3 | — [b] |
| BDBV R4386L GPΔmuc-WL²P⁴ | <1.0E-3 | <1.0E-3 | <1.0E-3 | <1.0E-3 | — [b] |

[a] $K_D$ values were derived from biolayer interferometry (BLI) using the binding equations describing a 1:1 interaction.
[b] "—" indicates cases where the peak signal value at the highest ebolavirus antigen concentration is 0.2 or lower.

**Fig. S1. Construct design and in vitro characterization of orthoebolavirus GPΔmuc trimers.** (**a**) Amino acid sequences of EBOV Mayinga , SUDV Gulu, and BDBV R3486L GPΔmuc-WL$^2$P$^x$-foldon-[His$_6$] constructs containing either the P$^2$ or P$^4$ mutation. Signal peptide, GP, restriction site (AS), foldon, linker (GS), and His$_6$ tag are highlighted in yellow, grey, green, orange, pink, and teal, respectively. The WL$^2$P$^x$ mutations are shown in red. Note: Foldon is a C-terminal trimerization motif used in all orthoebolavirus GPΔmuc constructs and therefore will not be included in the construct names (except here in the sequence definition) to avoid redundancy. The GS linker and His$_6$ tag are enclosed in [ ] to indicate that they are included only in a subset of constructs used for evaluating nickel-based purification. (**b**) SEC profiles of EBOV Mayinga, SUDV Gulu, and BDBV R3486L GPΔmuc proteins produced in 165 ml HEK293F cultures and purified using either a nickel affinity column or a CA45 immunoaffinity column. (**c**) Compositional site-specific glycan analysis of three orthoebolavirus GPΔmuc-WL$^2$P$^4$ trimers. The graphs summarize quantitative mass spectrometric analysis of the glycan population present at individual N-linked glycosylation sites simplified into categories of glycans. The oligomannose-type glycan series (M9 to M5; $Man_9GlcNAc_2$ to $Man_5GlcNAc_2$) is colored green, afucosylated and fucosylated hybrid-type glycans (hybrid and F hybrid) are dashed pink, and complex glycans are grouped according to the number of antennae and presence of core fucosylation and are colored pink. Unoccupancy of an N-linked glycan site is represented in gray. Glycan sites that could not be determined are denoted as "N.D.". (**d**) Flowchart illustrating image processing, 2D classification, and 3D reconstruction of negative stain EM (nsEM) data for EBOV, SUDV, and BDBV GPΔmuc trimers and their complexes with ADI-15878 Fab, using CryoSPARC. (**e**)-(**f**) Representative 2D classification images of EBOV, SUDV, and BDBV GPΔmuc-WL$^2$P$^4$ trimers and their complexes with ADI-15878 Fab. (**g**) ELISA analysis of EBOV, SUDV, and BDBV GPΔmuc trimers binding to 5 filovirus NAbs in IgG form. Briefly, each well was coated with 0.1 μg of the appropriate antigen, and IgG was diluted in a 10-fold dilution series from a starting concentration of 10 μg/ml for all tested antibodies. Error bars represent the difference between duplicate measurements at each concentration for each sample. (**h**) BLI analysis of EBOV, SUDV, and BDBV GPΔmuc trimers binding to 5 filovirus NAbs in IgG form. Sensorgrams were obtained from an Octet RED96 instrument using AHC biosensors. A two-fold concentration gradient of antigen, starting at 600 nM, was used in a dilution series of six. $K_D$ values derived from a 1:1 fitting model are summarized in a table.

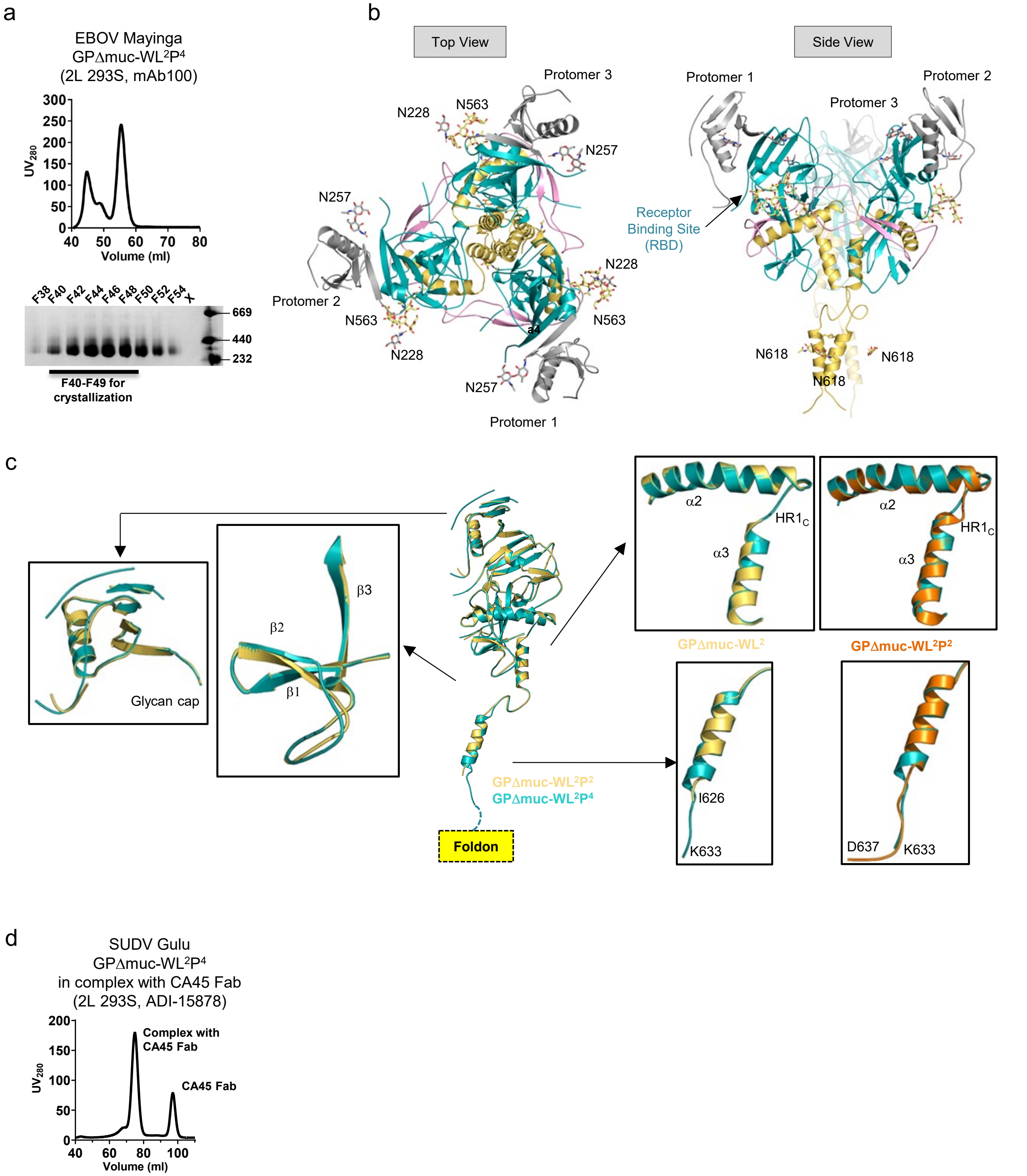
a
EBOV Mayinga
GPΔmuc-WL²P⁴
(2L 293S, mAb100)
UV280
Volume (ml)
F38 F40 F42 F44 F46 F48 F50 F52 F54 X
669
440
232
F40-F49 for crystallization
b
Top View
Side View
Protomer 1
Protomer 2
Protomer 3
N228
N563
N257
N618
Receptor Binding Site (RBD)
c
Glycan cap
β1
β2
β3
α2
α3
HR1C
GPΔmuc-WL²
GPΔmuc-WL²P²
GPΔmuc-WL²P⁴
I626
K633
D637
Foldon
d
SUDV Gulu
GPΔmuc-WL²P⁴
in complex with CA45 Fab
(2L 293S, ADI-15878)
Complex with CA45 Fab
CA45 Fab
UV280
Volume (ml)

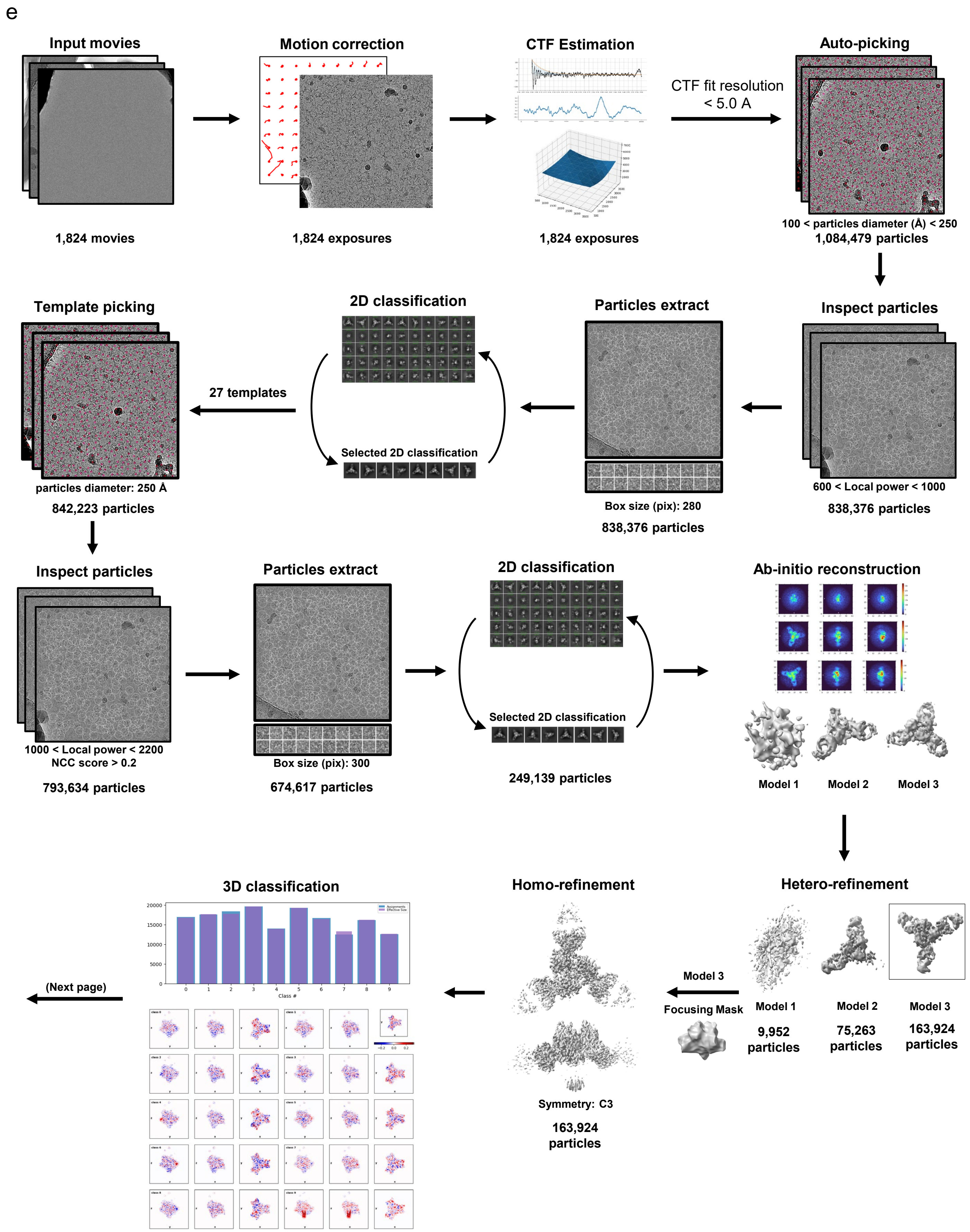
e
Input movies
1,824 movies
Motion correction
1,824 exposures
CTF Estimation
1,824 exposures
CTF fit resolution
< 5.0 A
Auto-picking
100 < particles diameter (Å) < 250
1,084,479 particles
Inspect particles
600 < Local power < 1000
838,376 particles
Particles extract
Box size (pix): 280
838,376 particles
2D classification
Selected 2D classification
27 templates
Template picking
particles diameter: 250 Å
842,223 particles
Inspect particles
1000 < Local power < 2200
NCC score > 0.2
793,634 particles
Particles extract
Box size (pix): 300
674,617 particles
2D classification
Selected 2D classification
249,139 particles
Ab-initio reconstruction
Model 1
Model 2
Model 3
Hetero-refinement
Model 1
9,952 particles
Model 2
75,263 particles
Model 3
163,924 particles
Model 3
Focusing Mask
Homo-refinement
Symmetry: C3
163,924 particles
3D classification
Class #
(Next page)

Fig. S2

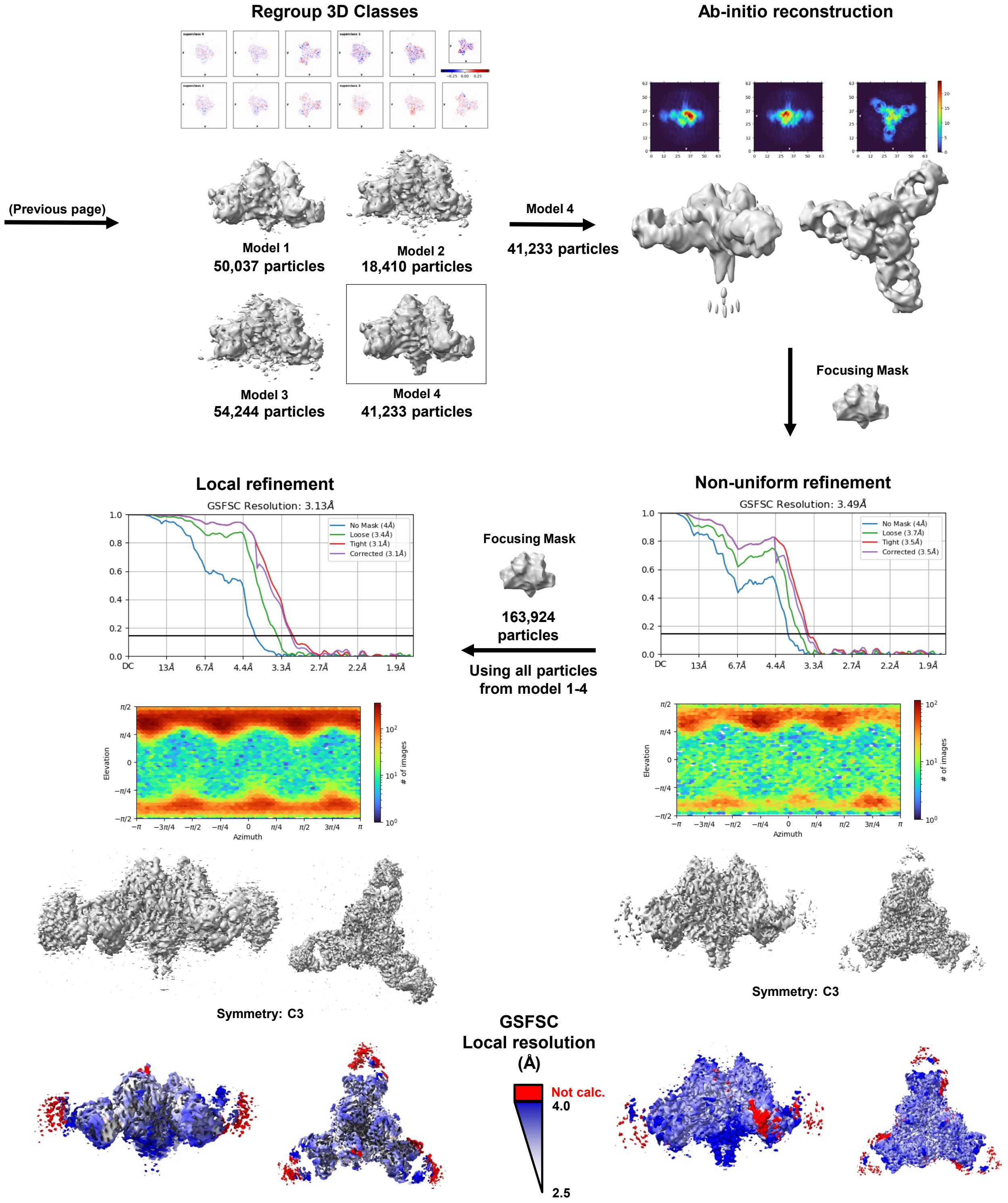
Regroup 3D Classes
Ab-initio reconstruction
(Previous page)
Model 1
50,037 particles
Model 2
18,410 particles
Model 3
54,244 particles
Model 4
41,233 particles
Model 4
41,233 particles
Focusing Mask
Local refinement
GSFSC Resolution: 3.13Å
No Mask (4Å)
Loose (3.4Å)
Tight (3.1Å)
Corrected (3.1Å)
Non-uniform refinement
GSFSC Resolution: 3.49Å
No Mask (4Å)
Loose (3.7Å)
Tight (3.5Å)
Corrected (3.5Å)
Focusing Mask
163,924
particles
Using all particles
from model 1-4
Elevation
Azimuth
### of images
Symmetry: C3
Symmetry: C3
GSFSC
Local resolution
(Å)
Not calc.
4.0
2.5

**Fig. S2**

f

**SUDV GPΔmuc-WL²P⁴ in complex with CA45 Fab**

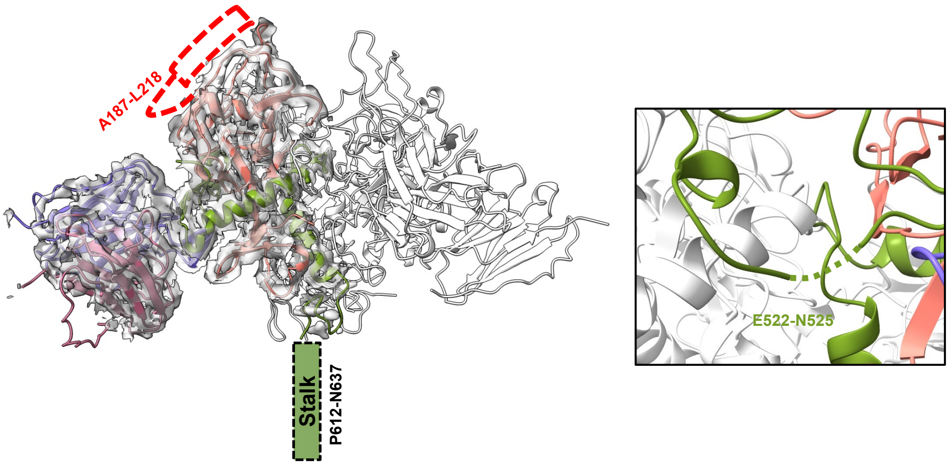


g

**EBOV Mayinga GPΔmuc-WL²P⁴**

**EBOV Mayinga GPΔmuc in complex with CA45 Fab (PDB ID: 6EAY)**

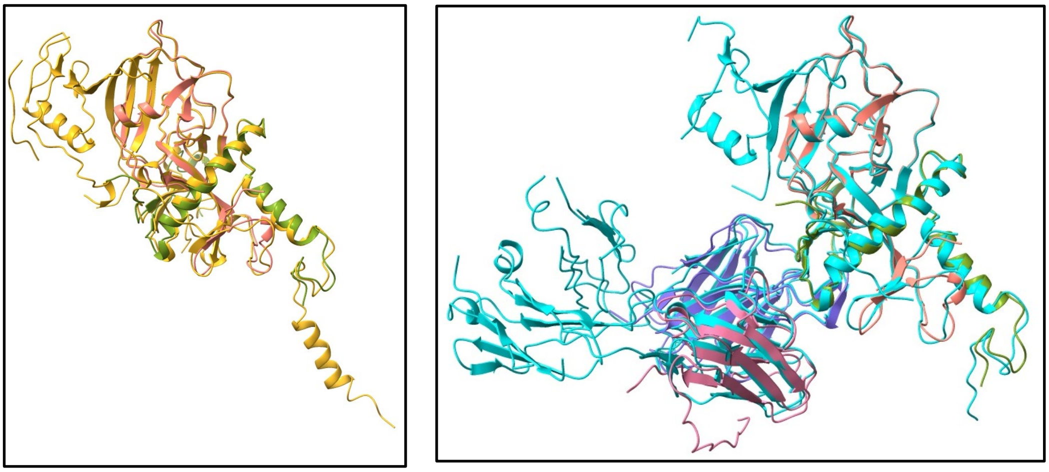

**Cα-RMSD: 0.71 Å**

**Cα-RMSD: 0.69 Å**

h

**Crystal structure of EBOV GPΔmuc in complex with CA45 Fab (PDB ID: 6EAY)**

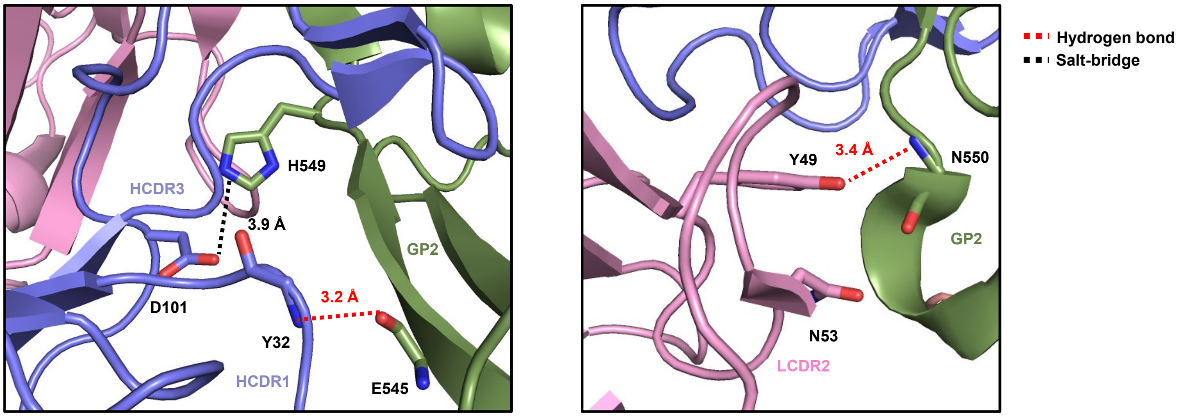


**Heavy chain variable domain of CA45 Fab**

**Light chain variable domain of CA45 Fab**

Fig. S2

i

**Interface between SUDV Gulu GPΔmuc-WL²P⁴ and CA45**

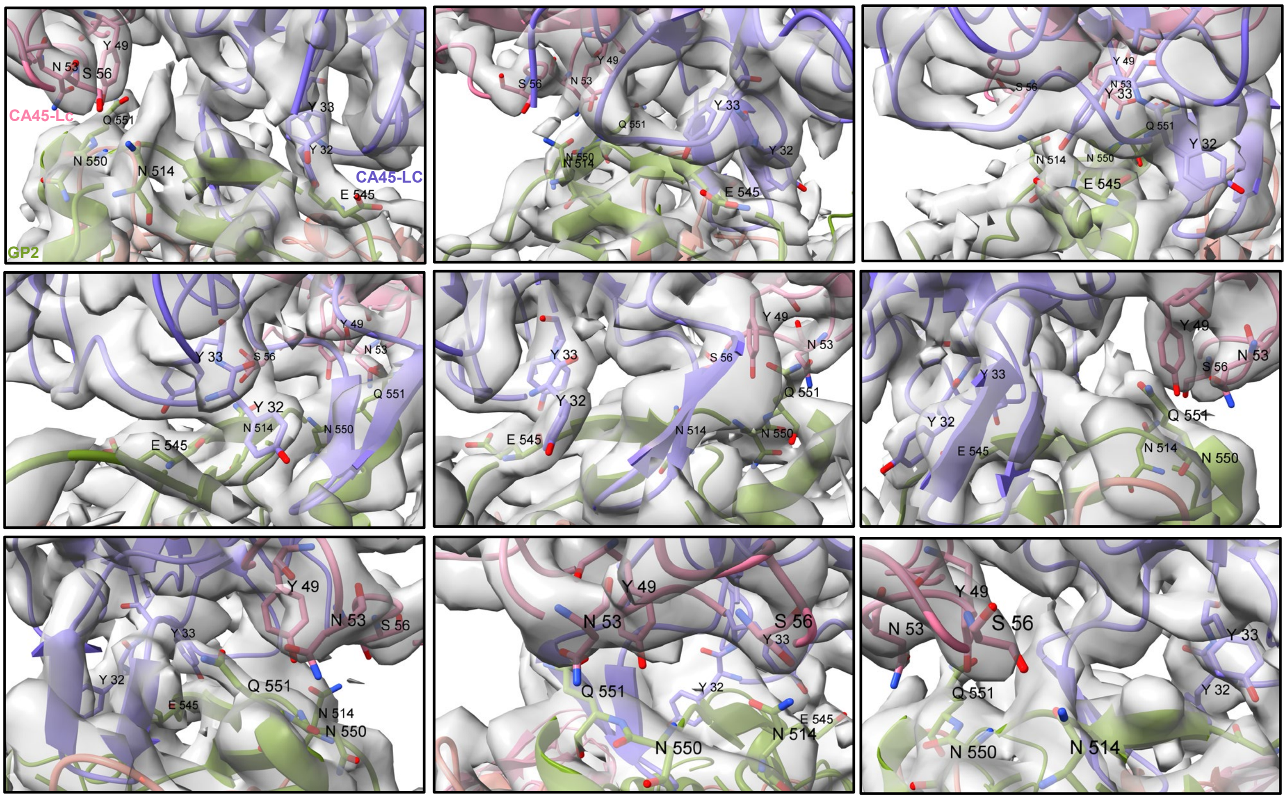


j

**Local resolution at the interface between SUDV Gulu GPΔmuc-WL²P⁴ and CA45**

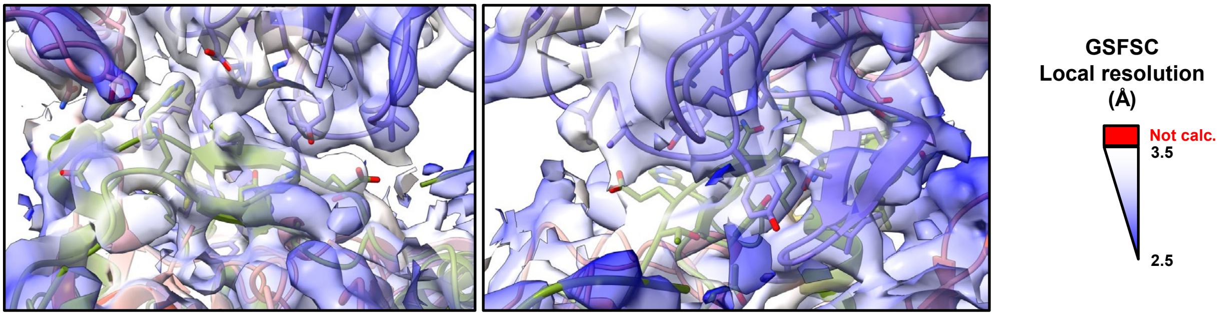


k

**EBOV GPΔmuc in complex with ADI-15878**

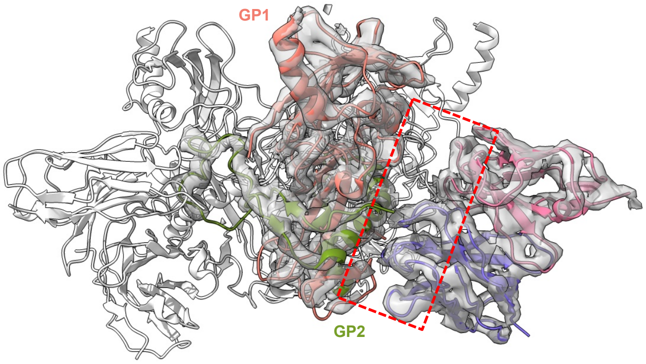


PDB ID: 6DZL (EMDB-8935)

**Interface between EBOV GPΔmuc and ADI-15878**

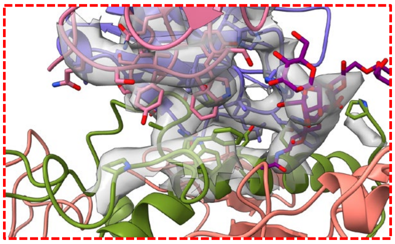

**Fig. S2**

l

**EBOV GPΔmuc in complex with rEBOV-515 and rEBOV-442**

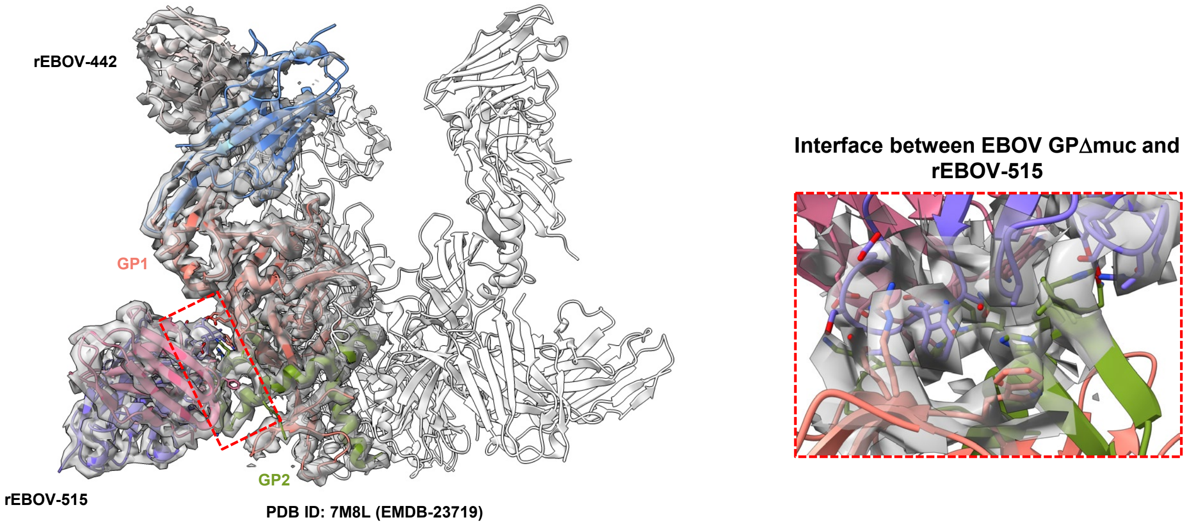


m

**EBOV GPΔmuc in complex with rEBOV-520 and rEBOV-548**

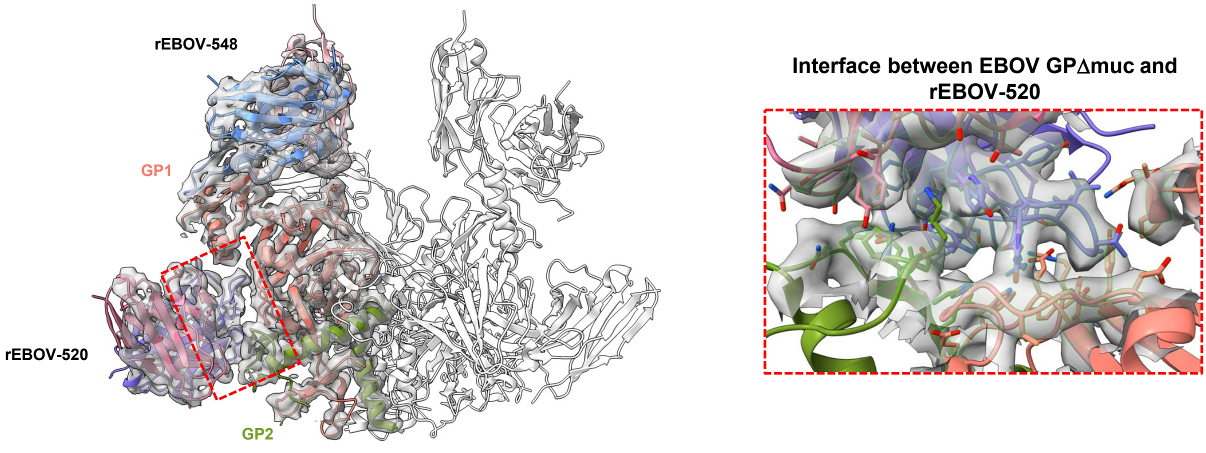


**Fig. S2. Structural characterization of ligand-free EBOV GPΔmuc-WL²P⁴ and CA45 Fab-bound SUDV GPΔmuc-WL²P⁴ trimers.** (**a**) SEC profile of EBOV GPΔmuc-WL²P⁴ trimer expressed in 2L HEK293S cells and BN-PAGE of ADI-15878/SEC-purified EBOV GPΔmuc-WL²P⁴ trimer. Fractions F40 to F49 were used for crystallization screening. The SEC profile was generated from a HiLoad Superdex 200 16/600 PG column (Cytiva). (**b**) Ribbon representation of EBOV GPΔmuc-WL²P⁴ trimer structure (Left: top view; Right: side view). The structure is colored as follows: glycan cap (gray), RBS (cyan), GP2 (yellow), and IFL (pink). *N*-linked glycans at N228, N257, N563, and N618 are shown as color-coded (by atom type) sticks. (**c**) Structural comparison of GPΔmuc-WL²P⁴ and previously reported GPΔmuc-WL² and GPΔmuc-WL²P² trimers. The ribbon models of three GPΔmuc trimers containing WL²P⁴, WL², and WL²P² mutations are shown in cyan, gold, and orange, respectively. Regions that are critical to the GP structure and exhibit appreciable differences in structure are superposed and shown as enlarged insects. (**d**) SEC profile of ADI-15878/SEC-purified SUDV GPΔmuc-WL²P⁴ trimer expressed in 2L HEK293S cells and complexed with ADI-15878 Fab. The SEC profile was generated from a Superose 6 10/300 increase column. (**e**) Schematic representation of image processing, 2D classification, and 3D reconstruction of cryo-EM data obtained for SUDV GPΔmuc-WL²P⁴ trimer in complex with ADI-15878 Fab using CryoSPARC. (**f**) Regions not modeled in the GP–CA45 complex structure due to limited density resolution. (**g**) Left: cryo-EM structure of SUDV GPΔmuc-WL²P⁴ superposed with the crystal structure of EBOV GPΔmuc-WL²P⁴ for comparison at the monomer level. The Cα root mean square deviation (Cα-RMSD) is 0.66 Å for 154 matching Cα atoms. Right: cryo-EM structure of SUDV GPΔmuc-WL²P⁴ in complex with ADI-15878 Fab superposed with the previously reported crystal structure of EBOV GPΔmuc in complex with CA45 Fab (PDB ID: 6EAY). Cα-RMSD is 0.63 Å for 149 matching Cα atoms. (**h**) Structural details of the GP-CA45 interface as illustrated in the crystal structure of EBOV GPΔmuc in complex with CA45 Fab (PDB ID: 6EAY). Hydrogen bonds are shown as red dashed lines, and the salt bridge is indicated with a black dashed line. Distances are labeled. (**i**) Snapshots of the GP-CA45 interface from different views with the density map shown. Interacting residues are displayed in ribbon representation. (**j**) Local resolution map overlaid with the built model at the GP-CA45 interface. (**k**) Published cryo-EM structure of EBOV GPΔmuc in complex with ADI-15878 (PDB ID: 6DZL; EMDB-8935), highlighting structural details at the interface. (**l**) Published cryo-EM structure of EBOV GPΔmuc in complex with rEBOV-515 and rEBOV-442 (PDB ID: 7M8L; EMDB-23719), with interface details shown. (**m**) Published cryo-EM structure of EBOV GPΔmuc in complex with rEBOV-520 and rEBOV-548 (PDB ID: 6PCI; EMDB-20301), highlighting interface details.

### Fig. S3

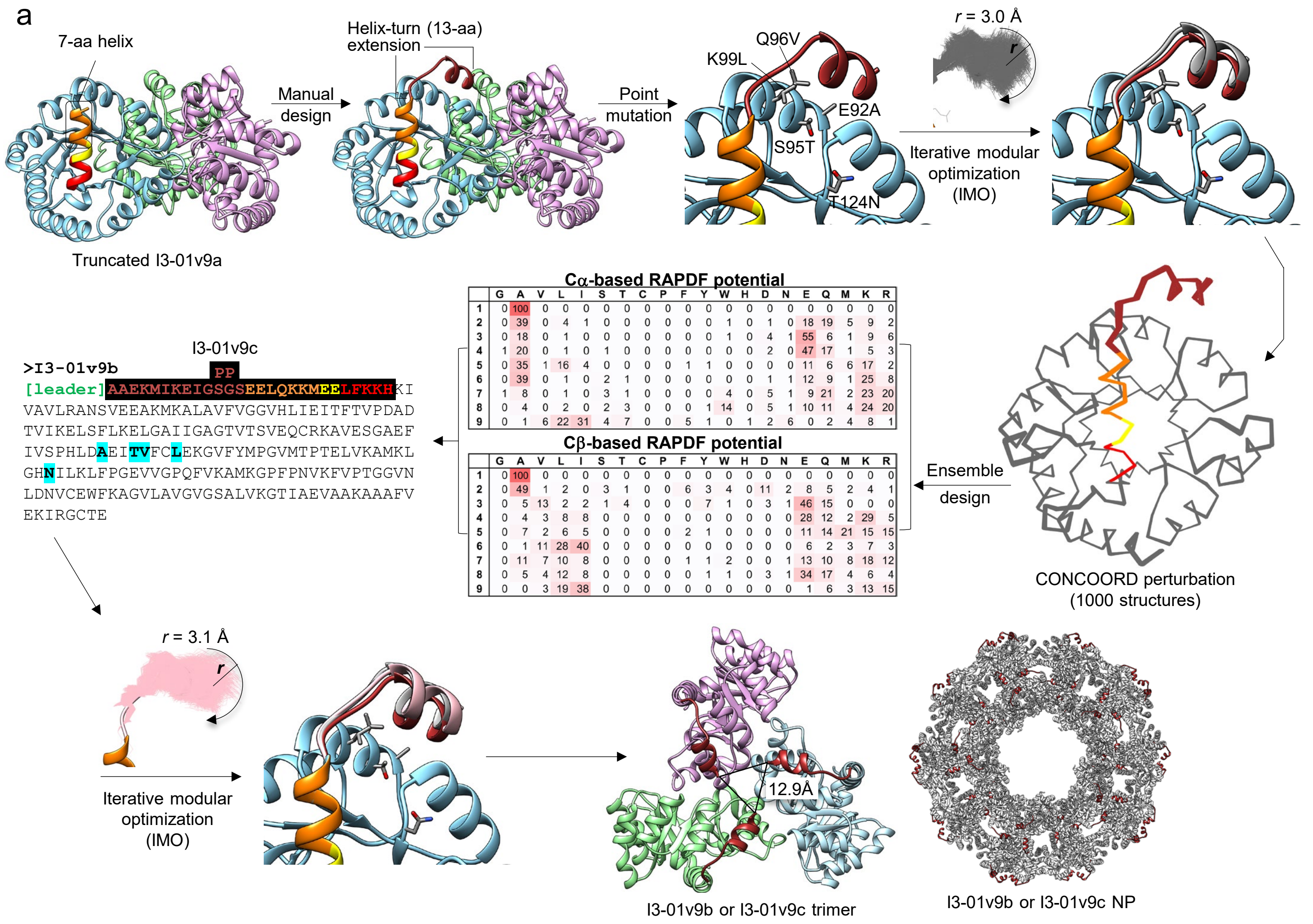


**Cα-based RAPDF potential**

| | G | A | V | L | I | S | T | C | P | F | Y | W | H | D | N | E | Q | M | K | R |
|---|---|---|---|---|---|---|---|---|---|---|---|---|---|---|---|---|---|---|---|---|
| **1** | 0 | 100 | 0 | 0 | 0 | 0 | 0 | 0 | 0 | 0 | 0 | 0 | 0 | 0 | 0 | 0 | 0 | 0 | 0 | 0 |
| **2** | 0 | 39 | 0 | 4 | 1 | 0 | 0 | 0 | 0 | 0 | 0 | 1 | 0 | 1 | 0 | 18 | 19 | 5 | 9 | 2 |
| **3** | 0 | 18 | 0 | 1 | 0 | 0 | 0 | 0 | 0 | 0 | 0 | 1 | 0 | 4 | 1 | 55 | 6 | 1 | 9 | 6 |
| **4** | 1 | 20 | 0 | 1 | 0 | 1 | 0 | 0 | 0 | 0 | 0 | 0 | 0 | 2 | 0 | 47 | 17 | 1 | 5 | 3 |
| **5** | 0 | 35 | 1 | 16 | 4 | 0 | 0 | 0 | 0 | 1 | 1 | 0 | 0 | 0 | 0 | 11 | 6 | 6 | 17 | 2 |
| **6** | 0 | 39 | 0 | 1 | 0 | 2 | 1 | 0 | 0 | 0 | 0 | 0 | 0 | 1 | 1 | 12 | 9 | 1 | 25 | 8 |
| **7** | 1 | 8 | 0 | 1 | 0 | 3 | 1 | 0 | 0 | 0 | 0 | 4 | 0 | 5 | 1 | 9 | 21 | 2 | 23 | 20 |
| **8** | 0 | 4 | 0 | 2 | 0 | 2 | 3 | 0 | 0 | 0 | 1 | 14 | 0 | 5 | 1 | 10 | 11 | 4 | 24 | 20 |
| **9** | 0 | 1 | 6 | 22 | 31 | 4 | 7 | 0 | 0 | 5 | 1 | 0 | 1 | 2 | 6 | 0 | 2 | 4 | 8 | 1 |

**Cβ-based RAPDF potential**

| | G | A | V | L | I | S | T | C | P | F | Y | W | H | D | N | E | Q | M | K | R |
|---|---|---|---|---|---|---|---|---|---|---|---|---|---|---|---|---|---|---|---|---|
| **1** | 0 | 100 | 0 | 0 | 0 | 0 | 0 | 0 | 0 | 0 | 0 | 0 | 0 | 0 | 0 | 0 | 0 | 0 | 0 | 0 |
| **2** | 0 | 49 | 1 | 2 | 0 | 3 | 1 | 0 | 0 | 6 | 3 | 4 | 0 | 11 | 2 | 8 | 5 | 2 | 4 | 1 |
| **3** | 0 | 5 | 13 | 2 | 2 | 1 | 4 | 0 | 0 | 0 | 7 | 1 | 0 | 3 | 1 | 46 | 15 | 0 | 0 | 0 |
| **4** | 0 | 4 | 3 | 8 | 8 | 0 | 0 | 0 | 0 | 0 | 0 | 0 | 0 | 0 | 0 | 28 | 12 | 2 | 29 | 5 |
| **5** | 0 | 7 | 2 | 6 | 5 | 0 | 0 | 0 | 0 | 2 | 1 | 0 | 0 | 0 | 0 | 11 | 14 | 21 | 15 | 15 |
| **6** | 0 | 1 | 11 | 28 | 40 | 0 | 0 | 0 | 0 | 0 | 0 | 0 | 0 | 0 | 0 | 6 | 2 | 3 | 7 | 3 |
| **7** | 0 | 11 | 7 | 10 | 8 | 0 | 0 | 0 | 0 | 1 | 1 | 2 | 0 | 0 | 1 | 13 | 10 | 8 | 18 | 12 |
| **8** | 0 | 5 | 4 | 12 | 8 | 0 | 0 | 0 | 0 | 0 | 1 | 1 | 0 | 3 | 1 | 34 | 17 | 4 | 6 | 4 |
| **9** | 0 | 0 | 3 | 19 | 38 | 0 | 0 | 0 | 0 | 0 | 0 | 0 | 0 | 0 | 0 | 1 | 6 | 3 | 13 | 15 |

**Fig. S3**

b

**>EBOV Mayinga GP$\Delta$muc-WL$^2$P$^4$-I3-01v9b-T**

MGVTGILQLPRDRFKRTSFFLWVIILFQRTFSIPLGVIHNSALQVSDVDKLVCRDKLSSTNQLRSVGLNLEGNGVATDVPSATKRWGFRSGVPPKVVNYEAGEWAENCYNLEI
KKPDGSECLPAAPDGIRGFPRCRYVHKVSGTGPCAGDFAFHKEGAFFLYDRLASTVIYRGTTFAEGVVAFLILPQAKKDFFSSHPLREPVNATEDPSSGYYSTTIRYQATGFG
TNETEYLFEVDNLTYVQLESRFTPQFLLQLNETIYTSGKRSNTTGKLIWKVNPEIDTTIGEWAFWETKKNLTRKIRSEELSFTVVSTHHQDTGEESASSGKLGLITNTIAGVA
GLITGGRRTRREAIVNAQPKCNPNLHYWTTQDEGAAIGLAWIPYFGPAAEGIYIEGLMHNQDGLICGLRQLANETTQALQLFLRATTEPRTFSILNRKAIDFLLQRWGGTCHI
LGPDCCIEPHDLTKNITDKIDQIIHDFVDKTLPDASGAEKMIKEIG**SG**SEELQKKMEELFKKHKIVAVLRANSVEEAKEKALAVFEGGVHLIEITFTVPDADTVIKELSFLKE
KGAIIGAGTVTSVEQCRKAVESGAEFIVSPHLDAEITVFCLEKGVFYMPGVMTPTELVKAMKLGHNILKLFPGEVVGPQFVKAMKGPFPNVKFVPTGGVNLDNVCEWFKAGVL
AVGVGSALVKGTPDEVREKAKAFVEKIRGCTE

**>EBOV Mayinga GP$\Delta$muc-WL$^2$P$^4$-I3-01v9c-T**

MGVTGILQLPRDRFKRTSFFLWVIILFQRTFSIPLGVIHNSALQVSDVDKLVCRDKLSSTNQLRSVGLNLEGNGVATDVPSATKRWGFRSGVPPKVVNYEAGEWAENCYNLEI
KKPDGSECLPAAPDGIRGFPRCRYVHKVSGTGPCAGDFAFHKEGAFFLYDRLASTVIYRGTTFAEGVVAFLILPQAKKDFFSSHPLREPVNATEDPSSGYYSTTIRYQATGFG
TNETEYLFEVDNLTYVQLESRFTPQFLLQLNETIYTSGKRSNTTGKLIWKVNPEIDTTIGEWAFWETKKNLTRKIRSEELSFTVVSTHHQDTGEESASSGKLGLITNTIAGVA
GLITGGRRTRREAIVNAQPKCNPNLHYWTTQDEGAAIGLAWIPYFGPAAEGIYIEGLMHNQDGLICGLRQLANETTQALQLFLRATTEPRTFSILNRKAIDFLLQRWGGTCHI
LGPDCCIEPHDLTKNITDKIDQIIHDFVDKTLPDASGAEKMIKEIG**PP**SEELQKKMEELFKKHKIVAVLRANSVEEAKEKALAVFEGGVHLIEITFTVPDADTVIKELSFLKE
KGAIIGAGTVTSVEQCRKAVESGAEFIVSPHLDAEITVFCLEKGVFYMPGVMTPTELVKAMKLGHNILKLFPGEVVGPQFVKAMKGPFPNVKFVPTGGVNLDNVCEWFKAGVL
AVGVGSALVKGTPDEVREKAKAFVEKIRGCTE

c

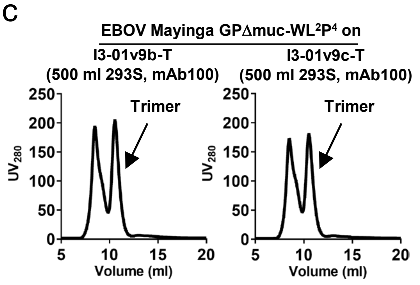


d

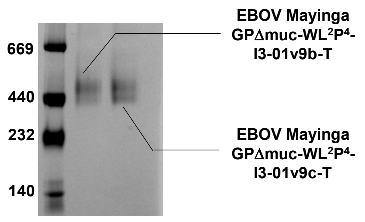


e **2D classes of EBOV Mayinga GP$\Delta$muc-WL$^2$P$^4$-I3-01v9b-T**

**Selected 2D classification for modeling**

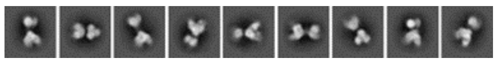

f **2D classes of EBOV Mayinga GP$\Delta$muc-WL$^2$P$^4$-I3-01v9c-T**

**Selected 2D classification for modeling**

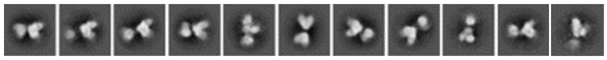

g

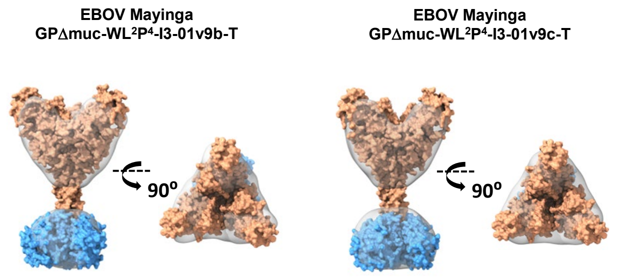


h

**>EBOV Mayinga GP$\Delta$muc-WL$^2$P$^4$-E2p-LD4-PADRE**

MGVTGILQLPRDRFKRTSFFLWVIILFQRTFSIPLGVIHNSALQVSDVDKLVCRDKLSSTNQLRSVGLNLEGNGVATDVPSATKRWGFRSGVPPKVVNYEAGEWAENCYNLEI
KKPDGSECLPAAPDGIRGFPRCRYVHKVSGTGPCAGDFAFHKEGAFFLYDRLASTVIYRGTTFAEGVVAFLILPQAKKDFFSSHPLREPVNATEDPSSGYYSTTIRYQATGFG
TNETEYLFEVDNLTYVQLESRFTPQFLLQLNETIYTSGKRSNTTGKLIWKVNPEIDTTIGEWAFWETKKNLTRKIRSEELSFTVVSTHHQDTGEESASSGKLGLITNTIAGVA
GLITGGRRTRREAIVNAQPKCNPNLHYWTTQDEGAAIGLAWIPYFGPAAEGIYIEGLMHNQDGLICGLRQLANETTQALQLFLRATTEPRTFSILNRKAIDFLLQRWGGTCHI
LGPDCCIEPHDLTKNITDKIDQIIHDFVDKTLPDASGAAAKPATTEGEFPETREKMSGIRRAIAKAMVHSKHTAPHVTLMDEADVKLVAHRKKFKAIAAEKGIKLTFLPYVVK
ALVSALREYPVLNTAIDDETEEIIQKHYYNIGIAADTDRGLLVPVIKHADRKPIFALAQEINELAEKARDGKLTPGEMKGASCTITNIGSAGGQWFTPVINHPEVAILGIGRI
AEKPIVRDGEIVAAPMLALSLSFDHRMIDGATAQKALNHIKRLLSDPELLLMGGGGSFSEEQKKALDLAFYFDRRLTPEWRRYLSQRLGLNEEQIERWFRRKEQQIGWSHPQF
EKGSAKFVAAWTLKAAA

**>EBOV Mayinga GP$\Delta$muc-WL$^2$P$^4$-I3-01v9b-LD7-PADRE**

MGVTGILQLPRDRFKRTSFFLWVIILFQRTFSIPLGVIHNSALQVSDVDKLVCRDKLSSTNQLRSVGLNLEGNGVATDVPSATKRWGFRSGVPPKVVNYEAGEWAENCYNLEI
KKPDGSECLPAAPDGIRGFPRCRYVHKVSGTGPCAGDFAFHKEGAFFLYDRLASTVIYRGTTFAEGVVAFLILPQAKKDFFSSHPLREPVNATEDPSSGYYSTTIRYQATGFG
TNETEYLFEVDNLTYVQLESRFTPQFLLQLNETIYTSGKRSNTTGKLIWKVNPEIDTTIGEWAFWETKKNLTRKIRSEELSFTVVSTHHQDTGEESASSGKLGLITNTIAGVA
GLITGGRRTRREAIVNAQPKCNPNLHYWTTQDEGAAIGLAWIPYFGPAAEGIYIEGLMHNQDGLICGLRQLANETTQALQLFLRATTEPRTFSILNRKAIDFLLQRWGGTCHI
LGPDCCIEPHDLTKNITDKIDQIIHDFVDKTLPDASGAEKMIKEIGSGSEELQKKMEELFKKHKIVAVLRANSVEEAKMKALAVFVGGVHLIEITFTVPDADTVIKELSFLKE
LGAIIGAGTVTSVEQCRKAVESGAEFIVSPHLDAEITVFCLEKGVFYMPGVMTPTELVKAMKLGHNILKLFPGEVVGPQFVKAMKGPFPNVKFVPTGGVNLDNVCEWFKAGVL
AVGVGSALVKGTIAEVAAKAAAFVEKIRGCTEGGGGSSPAVDIGDRLDELEKALEALSAEDGHDDVGQRLESLLRRWNSRRADGSAKFVAAWTLKAAA

**>SUDV Gulu GP$\Delta$muc-WL$^2$P$^4$-E2p-LD4-PADRE**

MGILPSPGMPALLSLVSLLSVLLMGCVAEMPLGVVTNSTLEVTEIDQLVCKDHLASTDQLKSVGLNLEGSGVSTDIPSATKRWGFRSGVPPKVVSYEAGEWAENCYNLEIKKP
DGSECLPPPPDGVRGFPRCRYVHKAQGTGPCPGDYAFHKDGAFFLYDRLASTVIYRGVNFAEGVIAFLILAKPKETFLQSPPIREAVNYTENTSSYYATSYLEYEIENFGAQH
STTLFKIDNNTFVRLDRPHTPQFLFQLNDTIHLHQQLSNTTGRLIWTLDANINADIGEWAFWENKKNLSEQLRGEELSFEALSLNETEDDDAASSSTSNGLITSTVTGILGSL
GLRKRSRRQTNTKATGKCNPNLHYWTAQEQHNAAGIAWIPYFGPGAEGIYTEGLMHNQNALVCGLRQLANETTQALQLFLRATTEPRTYTILNRKAIDFLLRRWGGTCRILGP
DCCIEPHDLTKNITDKINQIIHDFIDNPLPNASGAAAKPATTEGEFPETREKMSGIRRAIAKAMVHSKHTAPHVTLMDEADVTKLVAHRKKFKAIAAEKGIKLTFLPYVVKAL
VSALREYPVLNTAIDDETEEIIQKHYYNIGIAADTDRGLLVPVIKHADRKPIFALAQEINELAEKARDGKLTPGEMKGASCTITNIGSAGGQWFTPVINHPEVAILGIGRIAE
KPIVRDGEIVAAPMLALSLSFDHRMIDGATAQKALNHIKRLLSDPELLLMGGGGSFSEEQKKALDLAFYFDRRLTPEWRRYLSQRLGLNEEQIERWFRRKEQQIGWSHPQFEK
GSAKFVAAWTLKAAA

**>SUDV Gulu GP$\Delta$muc-WL$^2$P$^4$-I3-01v9b-LD7-PADRE**

MGILPSPGMPALLSLVSLLSVLLMGCVAEMPLGVVTNSTLEVTEIDQLVCKDHLASTDQLKSVGLNLEGSGVSTDIPSATKRWGFRSGVPPKVVSYEAGEWAENCYNLEIKKP
DGSECLPPPPDGVRGFPRCRYVHKAQGTGPCPGDYAFHKDGAFFLYDRLASTVIYRGVNFAEGVIAFLILAKPKETFLQSPPIREAVNYTENTSSYYATSYLEYEIENFGAQH
STTLFKIDNNTFVRLDRPHTPQFLFQLNDTIHLHQQLSNTTGRLIWTLDANINADIGEWAFWENKKNLSEQLRGEELSFEALSLNETEDDDAASSSTSNGLITSTVTGILGSL
GLRKRSRRQTNTKATGKCNPNLHYWTAQEQHNAAGIAWIPYFGPGAEGIYTEGLMHNQNALVCGLRQLANETTQALQLFLRATTEPRTYTILNRKAIDFLLRRWGGTCRILGP
DCCIEPHDLTKNITDKINQIIHDFIDNPLPNASGAEKMIKEIGSGSEELQKKMEELFKKHKIVAVLRANSVEEAKMKALAVFVGGVHLIEITFTVPDADTVIKELSFLKELGA
IIGAGTVTSVEQCRKAVESGAEFIVSPHLDAEITVFCLEKGVFYMPGVMTPTELVKAMKLGHNILKLFPGEVVGPQFVKAMKGPFPNVKFVPTGGVNLDNVCEWFKAGVLAVG
VGSALVKGTIAEVAAKAAAFVEKIRGCTEGGGGSSPAVDIGDRLDELEKALEALSAEDGHDDVGQRLESLLRRWNSRRADGSAKFVAAWTLKAAA

### Fig. S3

**>BDBV R4386L GPΔmuc-WL²P⁴-E2p-LD4-PADRE**

MGILPSPGMPALLSLVSLLSVLLMGCVAEIPLGVVHNNTLQVSDIDKLVCRDKLSSTSQLKSVGLNLEGNGVATDVPTATKRWGFRAGVPPKVVNYEAGEWAENCYNLDIKKA
DGSECLPEAPEGVRGFPRCRYVHKVSGTGPCPEGYAFHKEGAFFLYDRLASTIIYRSTTFSEGVVAFLILPETKKDFFQSPPLHEPANMTTDPSSYYHTVTLNYVADNFGTNM
TNFLFQVDHLTYVQLEPRFTPQFLVQLNETIYTNGRRSNTTGTLIWKVNPTVDTGVGEWAFWENKKNFTKTLSSEELSVIFVPRAQDPGSNDISESTEPGPLTNTTRGAANLL
TGSRRTRREITLRTQAKCNPNLHYWTTQDEGAAIGLAWIPYFGPAAEGIYTEGIMHNQNGLICGLRQLANETTQALQLFLRATTEPRTFSILNRKAIDFLLQRWGGTCHILGP
DCCIEPHDLTKNITDKIDQIIHDFIDKPLPDASGAAAKPATTEGEFPETREKMSGIRRAIAKAMVHSKHTAPHVTLMDEADVTKLVAHRKKFKAIAAEKGIKLTFLPYVVKAL
VSALREYPVLNTAIDDETEEIIQKHYYNIGIAADTDRGLLVPVIKHADRKPIFALAQEINELAEKARDGKLTPGEMKGASCTITNIGSAGGQWFTPVINHPEVAILGIGRIAE
KPIVRDGEIVAAPMLALSLSFDHRMIDGATAQKALNHIKRLLSDPELLLMGGGGSFSEEQKKALDLAFYFDRRLTPEWRRYLSQRLGLNEEQIERWFRRKEQQIGWSHPQFEK
GSAKFVAAWTLKAAA

**>BDBV R4386L GPΔmuc-WL²P⁴-I3-01v9b-LD7-PADRE**

MGILPSPGMPALLSLVSLLSVLLMGCVAEIPLGVVHNNTLQVSDIDKLVCRDKLSSTSQLKSVGLNLEGNGVATDVPTATKRWGFRAGVPPKVVNYEAGEWAENCYNLDIKKA
DGSECLPEAPEGVRGFPRCRYVHKVSGTGPCPEGYAFHKEGAFFLYDRLASTIIYRSTTFSEGVVAFLILPETKKDFFQSPPLHEPANMTTDPSSYYHTVTLNYVADNFGTNM
TNFLFQVDHLTYVQLEPRFTPQFLVQLNETIYTNGRRSNTTGTLIWKVNPTVDTGVGEWAFWENKKNFTKTLSSEELSVIFVPRAQDPGSNDISESTEPGPLTNTTRGAANLL
TGSRRTRREITLRTQAKCNPNLHYWTTQDEGAAIGLAWIPYFGPAAEGIYTEGIMHNQNGLICGLRQLANETTQALQLFLRATTEPRTFSILNRKAIDFLLQRWGGTCHILGP
DCCIEPHDLTKNITDKIDQIIHDFIDKPLPDASGAEKMIKEIGSGSEELQKKMEELFKKHKIVAVLRANSVEEAKMKALAVFVGGVHLIEITFTVPDADTVIKELSFLKELGA
IIGAGTVTSVEQCRKAVESGAEFIVSPHLDAEITVFCLEKGVFYMPGVMTPTELVKAMKLGHNILKLFPGEVVGPQFVKAMKGPFPNVKFVPTGGVNLDNVCEWFKAGVLAVG
VGSALVKGTIAEVAAKAAAFVEKIRGCTEGGGGSSPAVDIGDRLDELEKALEALSAEDGHDDVGQRLESLLRRWNSRRADGSAKFVAAWTLKAAA

i

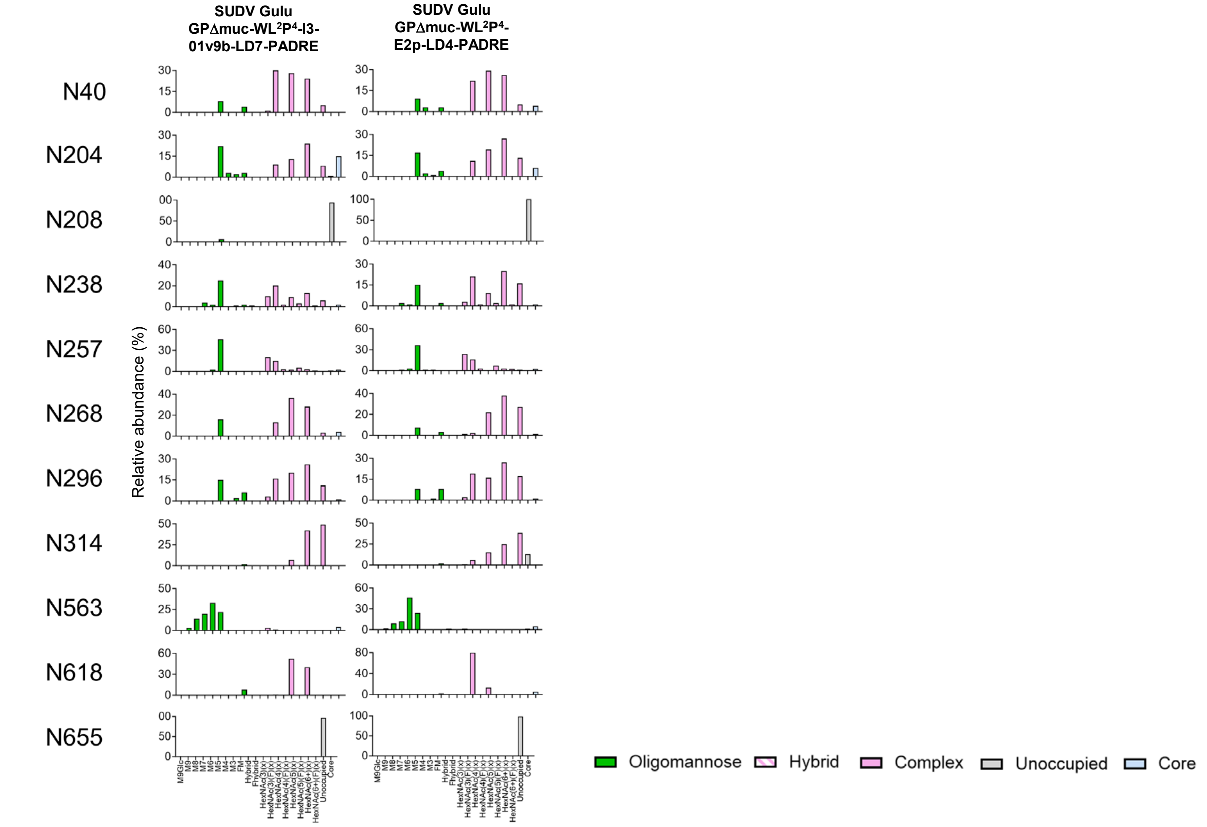


j

**ELISA analysis of 6 rationally designed ebolavirus GPΔmuc SApNPs binding to 5 filovirus NAbs**

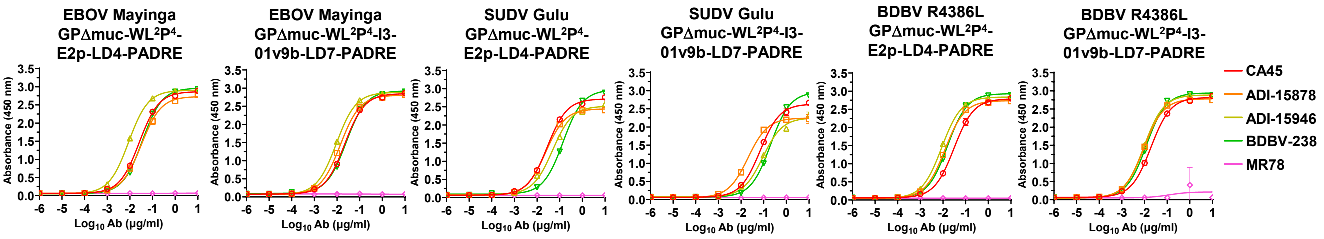


| EBOV Mayinga GPΔmuc-WL²P⁴E2p-LD4-PADRE | | | | | | | | | |
|---|---|---|---|---|---|---|---|---|---|
| CA45 | | ADI-15878 | | ADI-15946 | | BDBV-289 | | MR78 | |
| EC50 | STD | EC50 | STD | EC50 | STD | EC50 | STD | EC50 | STD |
| 0.02593 | 0.00030 | 0.03197 | 0.00696 | 0.00879 | 0.00075 | 0.03550 | 0.00212 | N.D. | N.D. |

| EBOV Mayinga GPΔmuc-WL²P⁴I3-01v9b-LD7-PADRE | | | | | | | | | |
|---|---|---|---|---|---|---|---|---|---|
| CA45 | | ADI-15878 | | ADI-15946 | | BDBV-289 | | MR78 | |
| EC50 | STD | EC50 | STD | EC50 | STD | EC50 | STD | EC50 | STD |
| 0.022141 | 0.000587 | 0.014095 | 0.000148 | 0.009384 | 0.000164 | 0.025555 | 0.00123 | N.D. | N.D. |

| SUDV Gulu GPΔmuc-WL²P⁴E2p-LD4-PADRE | | | | | | | | | |
|---|---|---|---|---|---|---|---|---|---|
| CA45 | | ADI-15878 | | ADI-15946 | | BDBV-289 | | MR78 | |
| EC50 | STD | EC50 | STD | EC50 | STD | EC50 | STD | EC50 | STD |
| 0.02819 | 0.00070 | 0.02407 | 0.00284 | 0.04811 | 0.00204 | 0.14057 | 0.00849 | N.D. | N.D. |

| SUDV Gulu GPΔmuc-WL²P⁴I3-01v9b-LD7-PADRE | | | | | | | | | |
|---|---|---|---|---|---|---|---|---|---|
| CA45 | | ADI-15878 | | ADI-15946 | | BDBV-289 | | MR78 | |
| EC50 | STD | EC50 | STD | EC50 | STD | EC50 | STD | EC50 | STD |
| 0.0699 | 0.0024 | 0.0204 | 0.0024 | 0.0910 | 0.0007 | 0.2017 | 0.0139 | N.D. | N.D. |

| BDBV R4386L GPΔmuc-WL²P⁴E2p-LD4-PADRE | | | | | | | | | |
|---|---|---|---|---|---|---|---|---|---|
| CA45 | | ADI-15878 | | ADI-15946 | | BDBV-289 | | MR78 | |
| EC50 | STD | EC50 | STD | EC50 | STD | EC50 | STD | EC50 | STD |
| 0.0307 | 0.0035 | 0.0123 | 0.0002 | 0.0087 | 0.0004 | 0.0159 | 0.0002 | N.D. | N.D. |

| BDBV R4386L GPΔmuc-WL²P⁴I3-01v9b-LD7-PADRE | | | | | | | | | |
|---|---|---|---|---|---|---|---|---|---|
| CA45 | | ADI-15878 | | ADI-15946 | | BDBV-289 | | MR78 | |
| EC50 | STD | EC50 | STD | EC50 | STD | EC50 | STD | EC50 | STD |
| 0.0191 | 0.0014 | 0.0093 | 0.0005 | 0.0095 | 0.0005 | 0.0120 | 0.0003 | N.D. | N.D. |

k **BLI analysis of 6 rationally designed orthoebolavirus GPΔmuc SApNPs binding to 5 filovirus NAbs**

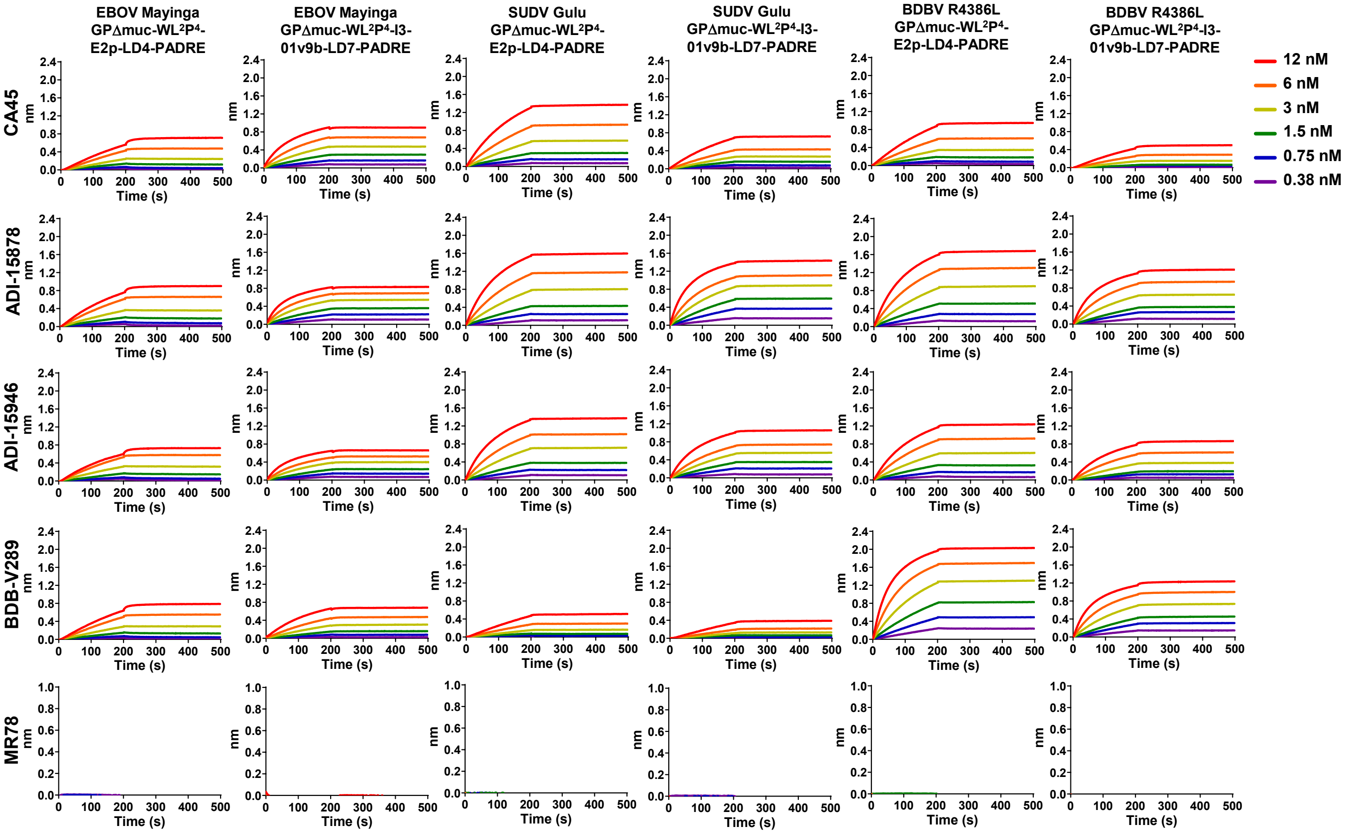


**$K_D$ values for 6 redesigned Filo-virus GP NP binding to 5 representative antibodies (nM)** [a]

| | CA45 | ADI-15878 | ADI-15946 | BDBV-289 | MR78 |
|---|---|---|---|---|---|
| EBOV Mayinga GPΔmuc-WL²P⁴-E2p-LD4-PADRE | <1.0E-3 | <1.0E-3 | <1.0E-3 | <1.0E-3 | — [b] |
| EBOV Mayinga GPΔmuc-WL²P⁴-I3-01v9b-LD7-PADRE | <1.0E-3 | <1.0E-3 | <1.0E-3 | <1.0E-3 | — [b] |
| SUDV Gulu GPΔmuc-WL²P⁴-E2p-LD4-PADRE | <1.0E-3 | <1.0E-3 | <1.0E-3 | <1.0E-3 | — [b] |
| SUDV Gulu GPΔmuc-WL²P⁴-I3-01v9b-LD7-PADRE | <1.0E-3 | <1.0E-3 | <1.0E-3 | <1.0E-3 | — [b] |
| BDBV R4386L GPΔmuc-WL²P⁴-E2p-LD4-PADRE | <1.0E-3 | <1.0E-3 | <1.0E-3 | <1.0E-3 | — [b] |
| BDBV R4386L GPΔmuc-WL²P⁴-I3-01v9b-LD7-PADRE | <1.0E-3 | <1.0E-3 | <1.0E-3 | <1.0E-3 | — [b] |

[a] $K_D$ values were derived from biolayer interferometry (BLI) using the binding equations describing a 1:1 interaction.
[b] "—" indicates cases where the peak signal value at the highest orthoebolavirus antigen concentration is 0.2 or lower.

**Fig. S3. Rational design of I3-01v9b/c and in vitro characterization of orthoebolavirus GPΔmuc-presenting SApNPs.** (**a**) Schematic representation of computational design of I3-01v9b/c. Top left: structural model of I3-01v9a with the 11-aa N-terminal helix truncated to 7 aa; Top middle left: a 13-aa helix-turn fragment (9-aa helix + 4-aa turn, all alanine) was fused to the 7-aa N-terminal helix of the truncated I3-01v9a in such a way the new 9-aa N-terminal helix from the fused fragment would pack within the groove of two helices that are part of the I3-01 core; Top middle right: four mutations, E92A, S95T, Q96V, and K99L, were introduced to the I3-01 core helices to remove their steric clashes with the new N-terminal helix; Top right: iterative modular optimization (IMO) of the helix-turn backbone, with a radius of 3.0 Å measured for the first amino acid at the N-terminus; Middle right: 1000 slightly perturbed backbone conformations generated by CONCOORD, a torsion-space sampling program; Middle middle: results from an ensemble-based protein design program used to predict the optimal sequence for the 9-aa N-terminal helix within the fragment using C$\alpha$ and C$\beta$-based RAPDF potentials. The 4-aa turn was set to "GSGS"; Middle left: sequence of I3-01v9b designed by combining data from both RAPDF potentials, with the difference between I3-01v9b and I3-01v9c (the "PP" mutation) labeled on the sequence; Bottom left: further backbone relaxation using IMO with a minimum perturbation angle; Bottom right: a structural model of the trimeric I3-01v9b/c, termed I3-01v9b/c-T, in which the N-termini form a triangle of 12.9 Å, and a structural model of fully assembled I3-01v9b/c NP. (**b**) Amino acids sequences of EBOV GPΔmuc-WL2P4-I3-01v9b/c-T, which are EBOV Mayinga GPΔmuc-WL$^2$P$^4$ anchored to the trimeric I3-01v9b/c-T scaffolds. Signal peptide, GP, restriction site (AS), and I3-01v9b/c-T are highlighted in yellow, grey, green and pink, respectively. The double proline mutation in the N-terminus of I3-01v9c-T that differs from I3-01v9b-T is highlighted in red. (**c**) SEC profiles of EBOV GPΔmuc-WL2P4-I3-01v9b/c-T expressed in 500 ml HEK293S cells and purified using an ADI-15878 column. (**d**) BN-PAGE of ADI-15878/SEC-purified EBOV GPΔmuc-WL2P4-I3-01v9b/c-T. (**e**) and (**f**) Representative 2D classification images of ADI-15878/SEC-purified EBOV GPΔmuc-WL2P4-I3-01v9b/c-T trimers. (**g**) 3D reconstructions of ADI-15878/SEC-purified EBOV GPΔmuc-WL2P4-I3-01v9b/c-T trimers. Crystal structures of EBOV GPΔmuc-WL$^2$P$^2$ (PDB ID:7JPH) and the bacterial enzyme from which I3-01 was derived (PDB ID:1VLW) were fitted into the model and shown in orange and blue, respectively. (**h**) Amino acids sequences of EBOV, SUDV, and BDBV GPΔmuc-WL$^2$P$^4$-E2P-LD4-PADRE and -I3-01v9b-LD7-PADRE SApNP constructs. LD stands for locking domain (LD), which is fused to the C-terminus of the NP backbone and forms an inner layer to stabilize the NP shell from the inside. PADRE is a helper T-cell epitope and forms a hydrophobic core at the center of the assembled NP to further stabilize the NP structure and to induce a strong T-help response upon vaccination. (**i**) Compositional site-specific glycan analysis of SUDV GPΔmuc-WL$^2$P$^4$-E2P-LD4-PADRE and I3-01v9b-LD7-PADRE SApNPs, as in **Fig. S1c**. (**j**) ELISA analysis of EBOV, SUDV, and BDBV GPΔmuc-WL$^2$P$^4$-E2P-LD4-PADRE and I3-01v9b-LD7-PADRE SApNPs binding to 5 filovirus NAbs, as in **Fig. S1h**. (**k**) BLI analysis of EBOV, SUDV, and BDBV GPΔmuc-WL$^2$P$^4$-E2P-LD4-PADRE and I3-01v9b-LD7-PADRE SApNPs binding to 5 filovirus NAbs. A two-fold concentration gradient of antigen, starting at 12 nM, was used in a dilution series of six. $K_D$ values derived from a 1:1 fitting model are summarized in a table.

#### Fig. S4

a

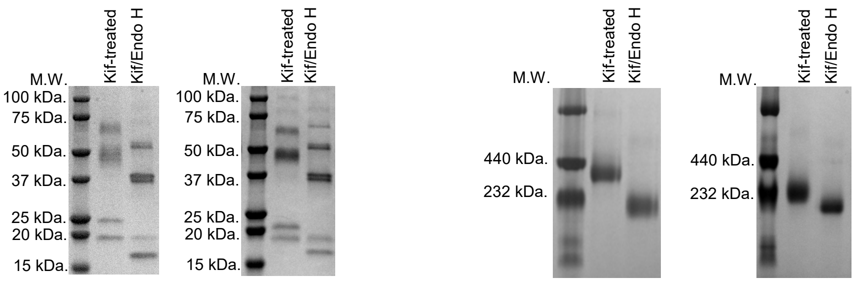


b ELISA analysis of Kif-treated and glycan-trimmed EBOV and SUDV GPΔmuc-WL²P⁴ trimers binding to 5 filovirus NAbs

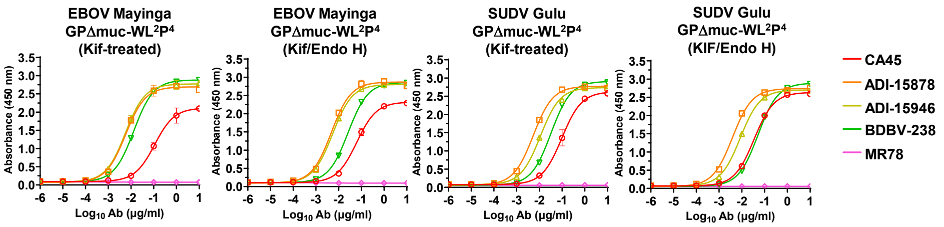


| EBOV Mayinga GPΔmuc-WL²P⁴ (Kif-treated) | | | | | | | | | |
|---|---|---|---|---|---|---|---|---|---|
| CA45 | | ADI-15878 | | ADI-15946 | | BDBV-289 | | MR78 | |
| EC50 | STD | EC50 | STD | EC50 | STD | EC50 | STD | EC50 | STD |
| 0.1110 | 0.0252 | 0.0058 | 0.0006 | 0.0057 | 0.0003 | 0.0133 | 0.0004 | N.D. | N.D. |

| EBOV Mayinga GPΔmuc-WL²P⁴ (Kif/Endo H) | | | | | | | | | |
|---|---|---|---|---|---|---|---|---|---|
| CA45 | | ADI-15878 | | ADI-15946 | | BDBV-289 | | MR78 | |
| EC50 | STD | EC50 | STD | EC50 | STD | EC50 | STD | EC50 | STD |
| 0.0624 | 0.0093 | 0.0047 | 0.0001 | 0.0055 | 0.0006 | 0.0261 | 0.0021 | N.D. | N.D. |

| SUDV Gulu GPΔmuc-WL²P⁴ (Kif-treated) | | | | | | | | | |
|---|---|---|---|---|---|---|---|---|---|
| CA45 | | ADI-15878 | | ADI-15946 | | BDBV-289 | | MR78 | |
| EC50 | STD | EC50 | STD | EC50 | STD | EC50 | STD | EC50 | STD |
| 0.0973 | 0.0354 | 0.0054 | 0.0003 | 0.0111 | 0.0003 | 0.0325 | 0.0013 | N.D. | N.D. |

| SUDV Gulu GPΔmuc-WL²P⁴ (Kif/Endo H) | | | | | | | | | |
|---|---|---|---|---|---|---|---|---|---|
| CA45 | | ADI-15878 | | ADI-15946 | | BDBV-289 | | MR78 | |
| EC50 | STD | EC50 | STD | EC50 | STD | EC50 | STD | EC50 | STD |
| 0.0347 | 0.0012 | 0.0039 | 0.0002 | 0.0086 | 0.0002 | 0.0518 | 0.0010 | N.D. | N.D. |

c BLI analysis of Kif-treated and glycan-trimmed EBOV and SUDV GPΔmuc-WL²P⁴ trimers binding to 5 filovirus NAbs

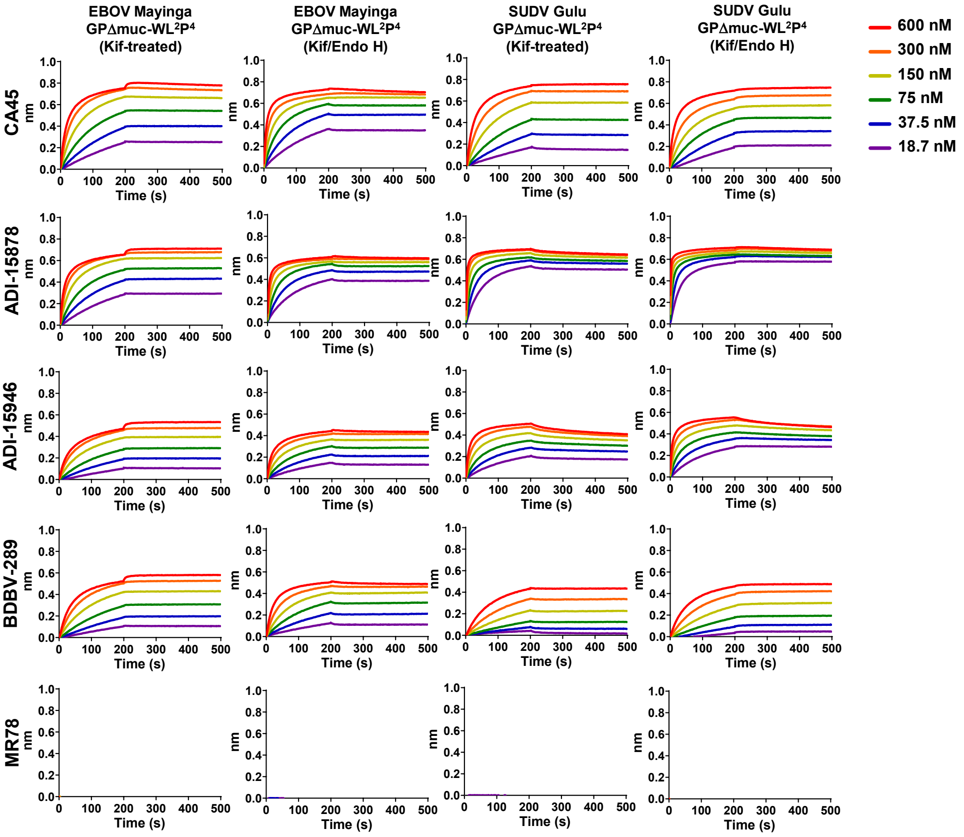

### Fig. S4

**$K_D$ values for 4 glycan-modified orthoebolavirus GPΔmuc trimers binding to 5 filovirus NAbs (nM) [a]**

| | CA45 | ADI-15878 | ADI-15946 | BDBV-289 | MR78 |
|---|---|---|---|---|---|
| EBOV Mayinga GPΔmuc-WL²P⁴ (Kif-treated) | <1.0E-3 | <1.0E-3 | <1.0E-3 | <1.0E-3 | — [b] |
| EBOV Mayinga GPΔmuc-WL²P⁴ (Kif/Endo H) | <1.0E-3 | <1.0E-3 | <1.0E-3 | <1.0E-3 | — [b] |
| SUDV Gulu GPΔmuc-WL²P⁴ (Kif-treated) | <1.0E-3 | 2.99E-2 | 2.03 | <1.0E-3 | — [b] |
| SUDV Gulu GPΔmuc-WL²P⁴ (Kif/Endo H) | <1.0E-3 | <1.0E-3 | 9.52E-1 | <1.0E-3 | — [b] |

[a] $K_D$ values were derived from bio-layer interferometry (BLI) using the binding equations describing a 1:1 interaction.
[b] "—" indicates cases where the peak signal value at the highest orthoebolavirus antigen concentration is 0.2 or lower.

d

**>SUDV Gulu GPΔmuc-WL²P⁴-PD-E2p-LD4-PADRE**
MGILPSPGMPALLSLVSLLSVLLMGCVAEMPLGVVTNSTLEVTEIDQLVCKDHLASTDQLKSVGLNLEGSGVSTDIPSATKRWGFRSGVPPKVVSYEAGEWAENCYNLEIKKP
DGSECLPPPPDGVRGFPRCRYVHKAQGTGPCPGDYAFHKDGAFFLYDRLASTVIYRGVNFAEGVIAFLILAKPKETFLQSPPIREAVNYTENTSSYYATSYLEYEIENFGAQH
STTLFKIDNNTFVRLDRPHTPQFLFQLNDTIHLHQQLSNTTGRLIWTLDANINADIGEWAFWENKKNLSEQLRGEELSFEALSLNETEDDDAASSSTSNGLITSTVTGILGSL
GLRKRSRRQTNTKATGKCNPNLHYWTAQEQHNAAGIAWIPYFGPGAEGIYTEGLMHNQNALVCGLRQLANETTQALQLFLRATTEPRTYTILNRKAIDFLLRRWGGTCRILGP
DCCIEPHDLTKNITDKINQIIHDFIDNPLPDASGAAAKPATTEGEFPETREKMSGIRRAIAKAMVHSKHTAPHVTLMDEADVT
KLVAHRKKFKAIAAEKGIKLTFLPYVVKALVSALREYPVLNTAIDDETEEIIQKHYYNIGIAADTDRGLLVPVIKHADRKPIFALAQEINELAEKARDGKLTPGEMKGASCTI
TNIGSAGGQWFTPVINHPEVAILGIGRIAEKPIVRDGEIVAAPMLALSLSFDHRMIDGATAQKALNHIKRLLSDPELLLMGGGGSFSEEQKKALDLAFYFDRRLTPEWRRYLS
QRLGLNEEQIERWFRRKEQQIGWSHPQFEKGSAKFVAAWTLKAAA

**>SUDV Gulu GPΔmuc-WL²P⁴-PD-I3-01v9b-LD7-PADRE**
MGILPSPGMPALLSLVSLLSVLLMGCVAEMPLGVVTNSTLEVTEIDQLVCKDHLASTDQLKSVGLNLEGSGVSTDIPSATKRWGFRSGVPPKVVSYEAGEWAENCYNLEIKKP
DGSECLPPPPDGVRGFPRCRYVHKAQGTGPCPGDYAFHKDGAFFLYDRLASTVIYRGVNFAEGVIAFLILAKPKETFLQSPPIREAVNYTENTSSYYATSYLEYEIENFGAQH
STTLFKIDNNTFVRLDRPHTPQFLFQLNDTIHLHQQLSNTTGRLIWTLDANINADIGEWAFWENKKNLSEQLRGEELSFEALSLNETEDDDAASSSTSNGLITSTVTGILGSL
GLRKRSRRQTNTKATGKCNPNLHYWTAQEQHNAAGIAWIPYFGPGAEGIYTEGLMHNQNALVCGLRQLANETTQALQLFLRATTEPRTYTILNRKAIDFLLRRWGGTCRILGP
DCCIEPHDLTKNITDKINQIIHDFIDNPLPDASGAEKMIKEIGSGSEELQKKMEELFKKHKIVAVLRANSVEEAKEKALAVFEGGVHLIEITFTVPDADTVIKELSFLKEKGA
IIGAGTVTSVEQCRKAVESGAEFIVSPHLDAEITVFCLEKGVFYMPGVMTPTELVKAMKLGHNILKLFPGEVVGPQFVKAMKGPFPNVKFVPTGGVNLDNVCEWFKAGVLAVG
VGSALVKGTPDEVREKAKAFVEKIRGCTEGGGGSSPAVDIGDRLDELEKALEALSAEDGHDDVGQRLESLLRRWNSRRADGSAKFVAAWTLKAAA

e

**SUDV Gulu**
**GPΔmuc-WL²P⁴-PD**
**E2p-LD4-PADRE**

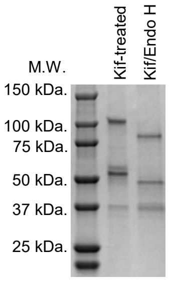


**SUDV Gulu**
**GPΔmuc-WL²P⁴-PD**
**I3-01v9b-LD7-PADRE**

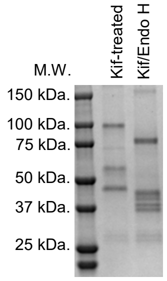

f Site-specific glycan profiles of wild-type and Kif-treated SUDV GPΔmuc-WL²P⁴-PD-presenting SApNPs

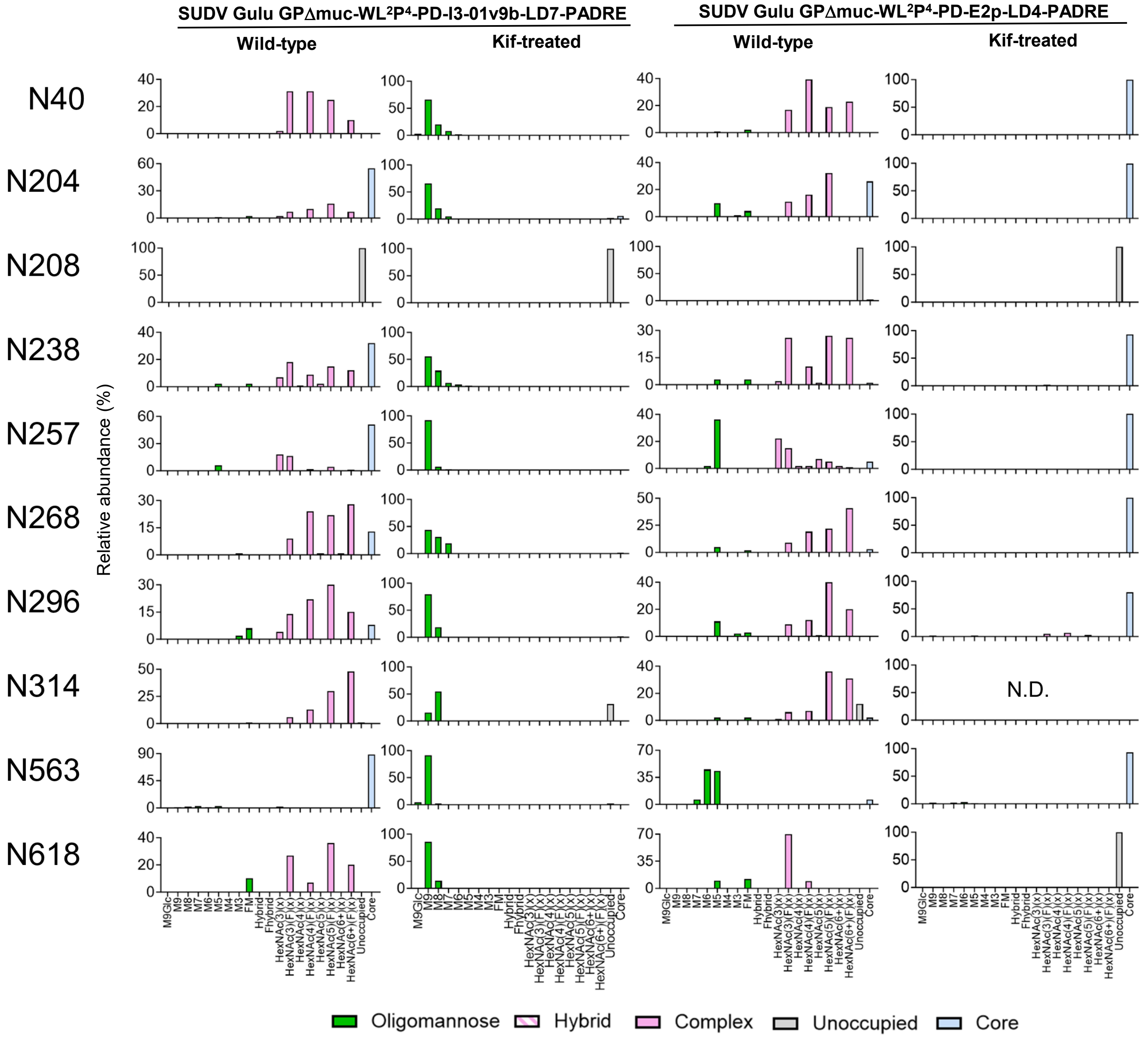

### Fig. S4

g

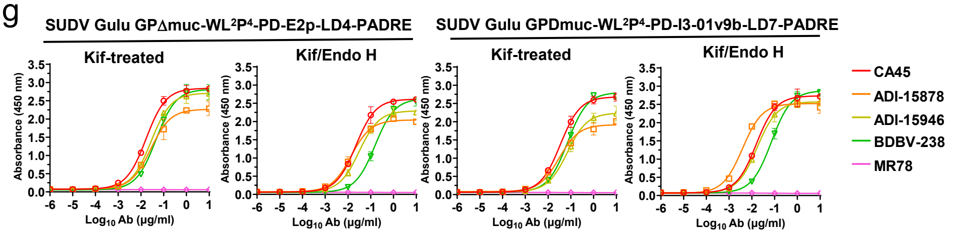


| SUDV Gulu GPΔmuc-WL2P4-PD-E2p-LD4-PADRE (Kif-treated) | | | | | | | | | |
|---|---|---|---|---|---|---|---|---|---|
| CA45 | | ADI-15878 | | ADI-15946 | | BDBV-829 | | MR78 | |
| EC50 | STD | EC50 | STD | EC50 | STD | EC50 | STD | EC50 | STD |
| 0.0168 | 0.0014 | 0.0277 | 0.0049 | 0.0283 | 0.0021 | 0.0462 | 0.0006 | N.D. | N.D. |

| SUDV Gulu GPΔmuc-WL2P4-PD-E2p-LD4-PADRE (Kif/Endo H) | | | | | | | | | |
|---|---|---|---|---|---|---|---|---|---|
| CA45 | | ADI-15878 | | ADI-15946 | | BDBV-829 | | MR78 | |
| EC50 | STD | EC50 | STD | EC50 | STD | EC50 | STD | EC50 | STD |
| 0.0226 | 0.0048 | 0.0308 | 0.002 | 0.0312 | 0.0043 | 0.1550 | 0.0179 | N.D. | N.D. |

| SUDV Gulu GPΔmuc-WL2P4-PD-I3-01v9b-LD7-PADRE (Kif-treated) | | | | | | | | | |
|---|---|---|---|---|---|---|---|---|---|
| CA45 | | ADI-15878 | | ADI-15946 | | BDBV-829 | | MR78 | |
| EC50 | STD | EC50 | STD | EC50 | STD | EC50 | STD | EC50 | STD |
| 0.0348 | 0.0024 | 0.0254 | 0.0163 | 0.0708 | 0.0049 | 0.0718 | 0.0181 | N.D. | N.D. |

| SUDV Gulu GPΔmuc-WL2P4-PD-I3-01v9b-LD7-PADRE (Kif/Endo H) | | | | | | | | | |
|---|---|---|---|---|---|---|---|---|---|
| CA45 | | ADI-15878 | | ADI-15946 | | BDBV-829 | | MR78 | |
| EC50 | STD | EC50 | STD | EC50 | STD | EC50 | STD | EC50 | STD |
| 0.0155 | 0.0023 | 0.0039 | 0.0004 | 0.0174 | 0.0011 | 0.0726 | 0.0083 | N.D. | N.D. |

h

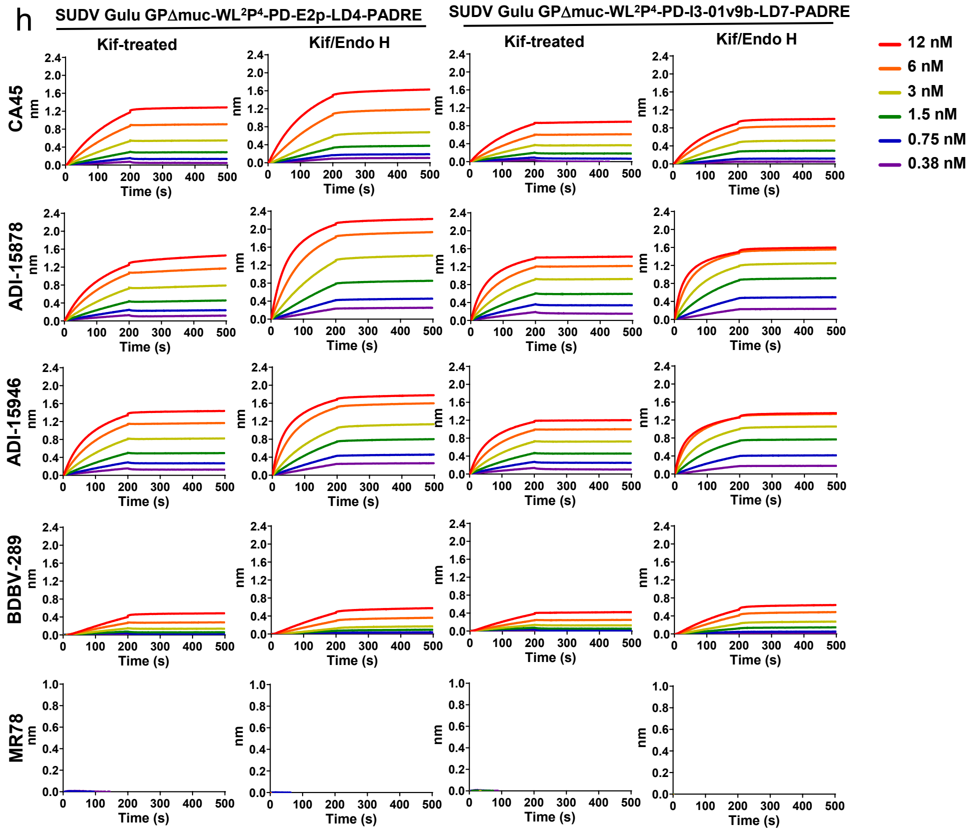


**$K_D$ values for 4 glycan-modified SUDVGPΔmuc-WL$^2$P$^4$-PD-presenting SApNPs binding to 5 filovirus NAbs (nM) [a]**

| | CA45 | ADI-15878 | ADI-15946 | BDBV-289 | MR78 |
|---|---|---|---|---|---|
| SUDV Gulu GPΔmuc-WL$^2$P$^4$-PD-E2p-LD4-PADRE (Kif-treated) | <1.0E-3 | <1.0E-3 | <1.0E-3 | <1.0E-3 | — [b] |
| SUDV Gulu GPΔmuc-WL$^2$P$^4$-PD-E2p-LD4-PADRE (Kif/Endo H) | <1.0E-3 | <1.0E-3 | <1.0E-3 | <1.0E-3 | — [b] |
| SUDV Gulu GPΔmuc-WL$^2$P$^4$-PD-I3-01v9b-LD7-PADRE (Kif-treated) | <1.0E-3 | <1.0E-3 | <1.0E-3 | <1.0E-3 | — [b] |
| SUDV Gulu GPΔmuc-WL$^2$P$^4$-PD-I3-01v9b-LD7-PADRE (Kif/Endo H) | <1.0E-3 | <1.0E-3 | <1.0E-3 | <1.0E-3 | — [b] |

[a] $K_D$ values were derived from biolayer interferometry (BLI) using the binding equations describing a 1:1 interaction.
[b] "—" indicates cases where the peak signal value at the highest SUDV antigen concentration is 0.2 or lower.

**Fig. S4. In vitro characterization for glycan-modified orthoebolavirus GPΔmuc trimers and SApNPs.** (**a**) SDS-PAGE and BN-PAGE of glycan-modified EBOV Mayinga and SUDV Gulu GPΔmuc-WL$^2$P$^4$ trimers. They were either expressed in the presence of kifunensine (Kif-treated) to obtain oligomannose-only glycans or further trimmed using endo H (Kif/Endo H) to obtain a mono-layer of GlcNAc stumps. (**b**) ELISA analysis of glycan-modified EBOV Mayinga and SUDV Gulu GPΔmuc-WL$^2$P$^4$ trimers binding to 5 filovirus NAbs in the IgG form, as in **Fig S1h.** (**c**) BLI analysis of glycan-modified EBOV Mayinga and SUDV Gulu GPΔmuc-WL$^2$P$^4$ trimers binding to 5 filovirus NAbs in the IgG form, as in **Fig S1i**. (**d**) Construct sequences for "multilayered" E2p (E2p-LD4-PADRE) and I3-01v9b (I3-0v9b-LD7-PADRE) SApNPs presenting 20 copies of a SUDV Gulu GPΔmuc-WL$^2$P$^4$-PD trimer, which contains a mutation (N637D) at the C-terminus to remove a potential glycosylation site between GP and NP. (**e**) SDS-PAGE of 3 glycan-modified SApNPs corresponding to EBOV Mayinga GPΔmuc-WL$^2$P$^4$ on the E2p-LD4-PADRE scaffold and SUDV Gulu GPΔmuc-WL$^2$P$^4$-PD on the E2p-LD4-PADRE and I3-01v9b-LD7-PADRE scaffolds. (**f**) Compositional site-specific glycan analysis of SUDV GPΔmuc-WL$^2$P$^4$-PD-presenting E2p and I3-01v9b SApNPs, as in **Fig. S1c**. (**g**) ELISA analysis of glycan-modified SUDV Gulu GPΔmuc-WL2P4-PD-presenting E2P and I3-01v9b SApNPs binding to 5 filovirus NAbs in the IgG form, as in **Fig S1h**. (**h**) BLI analysis of glycan-modified SUDV Gulu GPΔmuc-WL$^2$P$^4$-PD-presenting E2P and I3-01v9b SApNPs binding to 5 filovirus NAbs in the IgG form, as in **Fig S3k**.

### Fig. S5

a

**>RAVV GPΔTM-foldon-His$_6$**

MGVTGILQLPRDRFKRTSFFLWVIILFQRTFSIKTLPVLEIASNSQPQDVDSVCSGTLQKTEDVHLMGFTLSGQKVADSPLEASKRWAFRTGVPPKNVEYTEGEEAKTCYNIS
VTDPSGKSLLLDPPSNIRDYPKCKTVHHIQGQNPHAQGIALHLWGAFFLYDRVASTTMYRGKVFTEGNIAAMIVNKTVHRMIFSRQGQGYRHMNLTSTNKYWTSSNETQRNDT
GCFGILQEYNSTNNQTCPPSLKPPSLPTVTPSIHSTNTQINTAK**SGT**MNPSSDDEDLMISGSGSGEQGPHTTLNVVTEQKQSSTILSTPSLHPSTSQHEQNSTNPSRHAVTEH
NGTDPTTQPATLLNNTNTTPTYNTLKYNLSTPSPPTRNITNNDTQRELAESEQTNAQLNTTLDPTENPTTGQDTNSTTNIIMTTSDITSKHPTNSSPDSSPTT**RPPI**YFRKKR
SIFWKEGDIFPFLDGLINTEIDFDPIPNTETIFDESPSFNTSTNEEQHTPPNISLTFSYFPDKNGDTAYSGENENDCDAELRIWSVQEDDLAAGLSWIPFFGPGIEGLYTAGL
IKNQNNLVCRLRRLANQTAKSLELLLRVTTEERTFSLINRHAIDFLLTRWGGTCKVLGPDCCIGIEDLSKNISEQIDKIRKDEQKEETGASGYIPEAPRDGQAYVRKDGEWVL
LSTFLGSHHHHHH

**>RAVV GPΔmuc-foldon-His$_6$**

MGVTGILQLPRDRFKRTSFFLWVIILFQRTFSIKTLPVLEIASNSQPQDVDSVCSGTLQKTEDVHLMGFTLSGQKVADSPLEASKRWAFRTGVPPKNVEYTEGEEAKTCYNIS
VTDPSGKSLLLDPPSNIRDYPKCKTVHHIQGQNPHAQGIALHLWGAFFLYDRVASTTMYRGKVFTEGNIAAMIVNKTVHRMIFSRQGQGYRHMNLTSTNKYWTSSNETQRNDT
GCFGILQEYNSTNNQTCPPSLKPPSLPTVTPSIHSTNTQINTAK**SGTRPP**IYFRKKRSIFWKEGDIFPFLDGLINTEIDFDPIPNTETIFDESPSFNTSTNEEQHTPPNISLT
FSYFPDKNGDTAYSGENENDCDAELRIWSVQEDDLAAGLSWIPFFGPGIEGLYTAGLIKNQNNLVCRLRRLANQTAKSLELLLRVTTEERTFSLINRHAIDFLLTRWGGTCKV
LGPDCCIGIEDLSKNISEQIDKIRKDEQKEETGASGYIPEAPRDGQAYVRKDGEWVLLSTFLGSHHHHHH

**>RAVV GPΔmuc-P$^2$-foldon-His$_6$**

MGVTGILQLPRDRFKRTSFFLWVIILFQRTFSIKTLPVLEIASNSQPQDVDSVCSGTLQKTEDVHLMGFTLSGQKVADSPLEASKRWAFRTGVPPKNVEYTEGEEAKTCYNIS
VTDPSGKSLLLDPPSNIRDYPKCKTVHHIQGQNPHAQGIALHLWGAFFLYDRVASTTMYRGKVFTEGNIAAMIVNKTVHRMIFSRQGQGYRHMNLTSTNKYWTSSNETQRNDT
GCFGILQEYNSTNNQTCPPSLKPPSLPTVTPSIHSTNTQINTAK**SGTRPP**IYFRKKRSIFWKEGDIFPFLDGLINTEIDFDPIPNTETIFDESPSFNTSTNEEQHTPPNISLT
FSYFPDKNGDTAYSGENENDCDAELRIWSVQEDDLAAGLSWIPFFGPGIEGLYTAGLIKNQNNLVCRLRRLANQTAKSLELLLRVT**P**EERTFSLINRHAIDFLLTRWGGTCKV
LGPDCCIGIEDLSKNISEQIDKIRKDEQKEETGASGYIPEAPRDGQAYVRKDGEWVLLSTFLGSHHHHHH

**>RAVV GPΔmuc-P$^4$-foldon-His$_6$**

MGVTGILQLPRDRFKRTSFFLWVIILFQRTFSIKTLPVLEIASNSQPQDVDSVCSGTLQKTEDVHLMGFTLSGQKVADSPLEASKRWAFRTGVPPKNVEYTEGEEAKTCYNIS
VTDPSGKSLLLDPPSNIRDYPKCKTVHHIQGQNPHAQGIALHLWGAFFLYDRVASTTMYRGKVFTEGNIAAMIVNKTVHRMIFSRQGQGYRHMNLTSTNKYWTSSNETQRNDT
GCFGILQEYNSTNNQTCPPSLKPPSLPTVTPSIHSTNTQINTAK**SGTRPP**IYFRKKRSIFWKEGDIFPFLDGLINTEIDFDPIPNTETIFDESPSFNTSTNEEQHTPPNISLT
FSYFPDKNGDTAYSGENENDCDAELRIWSVQEDDLAAGLSWIPFFGPGIEGLYTAGLIKNQNNLVCRLRRLANQTAKSLELLLRVTTE**P**RTFSLINRHAIDFLLTRWGGTCKV
LGPDCCIGIEDLSKNISEQIDKIRKDEQKEETGASGYIPEAPRDGQAYVRKDGEWVLLSTFLGSHHHHHH

**>RAVV GPΔmuc-P$^2$CT-foldon-His$_6$**

MGVTGILQLPRDRFKRTSFFLWVIILFQRTFSIKTLPVLEIASNSQPQDVDSVCSGTLQKTEDVHLMGFTLSGQKVADSPLEASKRWAFRTGVPPKNVEYTEGEEAKTCYNIS
VTDPSGKSLLLDPPSNIRDYPKCKTVHHIQGQNPHAQGIALHLWGAFFLYDRVASTTMYRGKVFTEGNIAAMIVNKTVHRMIFSRQGQGYRHMNLTSTNKYWTSSNETQRNDT
GCFGILQEYNSTNNQTCPPSLKPPSLPTVTPSIHSTNTQINTAK**SGTRPP**IYFRKKRSIFWKEGDIFPFLDGLINTEIDFDPIPNTETIFDESPSFNTSTNEEQHTPPNISLT
FSYFPDKNGDTAYSGENENDCDAELRIWSVQEDDLAAGLSWIPFFGPGIEGLYTAGLIKNQNNLVCRLRRLANQTAKSLELLLRVT**P**EERTFSLINRHAIDFLLTRWGGTCKV
LGPDCCIGIEDLSKNISEQIDKIRKD**FV**KEETGASGYIPEAPRDGQAYVRKDGEWVLLSTFLGSHHHHHH

b

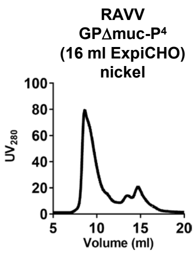


c

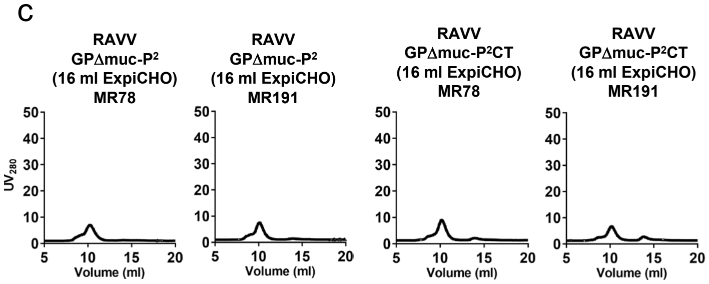


d

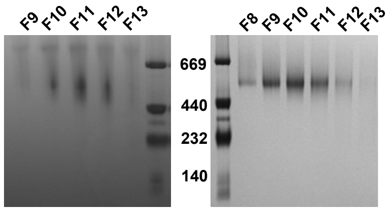


e

2D classes of RAVV GPΔmuc-P$^2$CT / MR78 Fab complex

Selected 2D classifications for 3D modeling

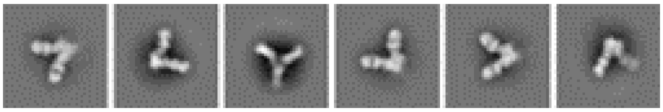

f

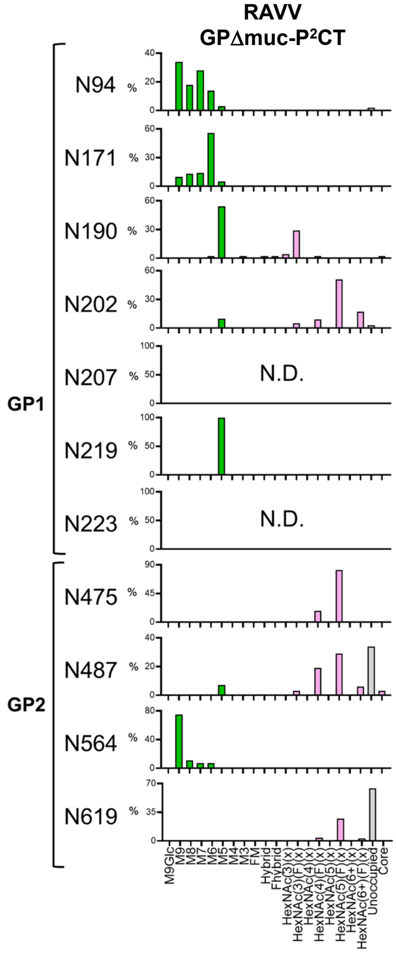


g

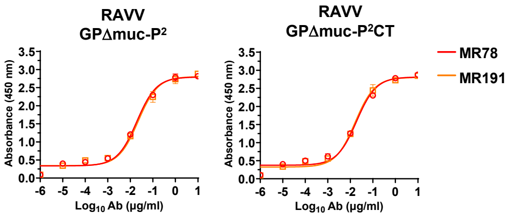


| RAVV GPΔmuc-P² | | | |
|---|---|---|---|
| MR78 | | MR191 | |
| EC50 | STD | EC50 | STD |
| 0.0191 | 0.0007 | 0.0166 | 0.0007 |

| RAVV GPΔmuc-P²CT | | | |
|---|---|---|---|
| MR78 | | MR191 | |
| EC50 | STD | EC50 | STD |
| 0.0196 | 0.0015 | 0.0230 | 0.0046 |

h

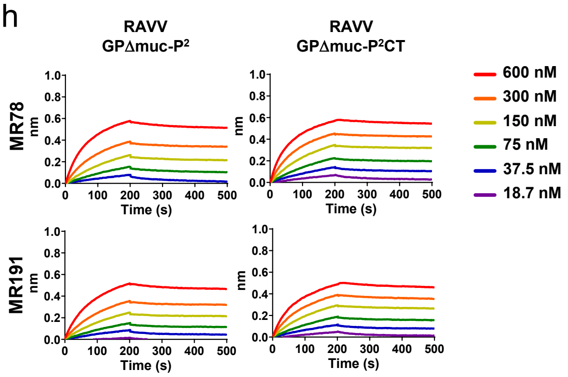


**$K_D$ values for two RAVV GPΔmuc trimers binding to 2 marburgvirus NAbs in the IgG form (nM)** [a]

| | MR78 | MR191 |
|---|---|---|
| RAVV GPΔmuc-P² | 6.61E-1 | 1.55E+1 |
| RAVV GPΔmuc-P²CT | 8.06 | 1.30E+1 |

[a] $K_D$ values were derived from bio-layer interferometry (BLI) using the binding equations describing a 1:1 interaction.
[b] "—" indicates cases where the peak signal value at the highest RSV-F concentration is 0.2 or lower.

i

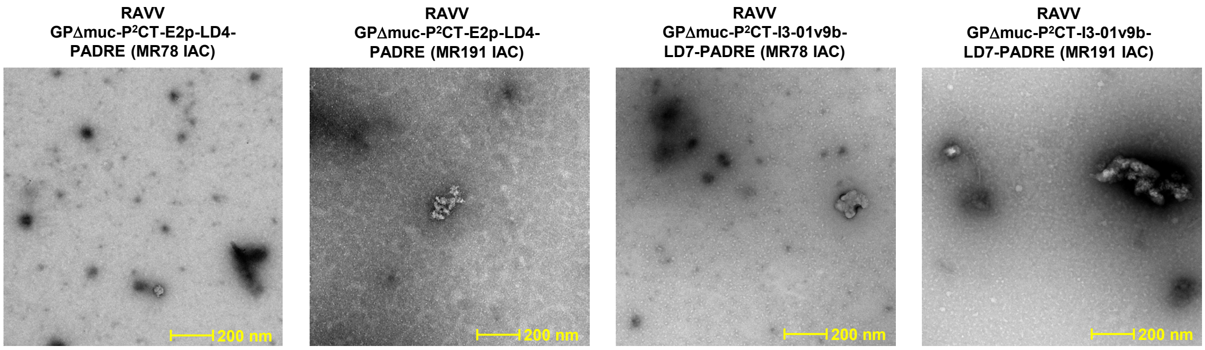


**Fig. S5. Construct design and in vitro characterization of RAVV GP trimers.** (**a**) Constructs sequences of RAVV GP designs. Signal peptide, mucin-like domain (MLD), HR1c, restriction site (AS), foldon, linker (GS), and $His_6$-tag are highlighted in yellow, gray, cyan, green, orange, light pink, and light sea green, respectively. The proline mutation on HR1c and CT are shown in red. Notably, the foldon motif, GS linker, and $His_6$ tag are present in all RAVV GP trimers tested in this study and will not be included in the construct names to avoid redundancy. (**b**) SEC profile of nickel-purified RAVV GPΔmuc-$P^4$ expressed in 16 ml ExpiCHO cells. (**c**) SECs profile of NAb MR78/MR191-purified RAVV GPΔmuc-$P^2$ and GPΔmuc-$P^2$CT each expressed in 16 ml ExpiCHO cells. (**d**) BN-PAGE of the trimer fractions from the SEC analysis of nickel-purified RAVV GPΔmuc-$P^2$ and GPΔmuc-$P^2$CT. (**e**) Representative 2D classification images of the RAVV GPΔmuc-$P^2$CT trimer in complex with MR78 Fab. (**f**) Compositional site-specific glycan analysis for RAVV GPΔmuc-$P^2$CT trimer, as in **Fig. S1c**. (**g**) ELISA analysis of RAVV GPΔmuc-$P^2$ and GPΔmuc-$P^2$CT trimers binding to NAbs MR78 and MR191 in the IgG form. Briefly, each well was coated with 0.1 μg of the appropriate antigen and IgG antibodies were diluted in a 10-fold dilution series from a starting concentration of 10 μg/ml for all antibodies. Error bars represent the difference between duplicate measurements at each concentration for each sample. (**h**) BLI analysis of RAVV GPΔmuc-$P^2$ and GPΔmuc-$P^2$CT trimers binding to NAbs MR78 and MR191 in the IgG form, as in **Fig. S1i**. (i) nsEM analysis of RAVV GPΔmuc-$P^2$CT E2p and I3-01v9b constructs following ExpiCHO expression and MR78- or MR191-based IAC purification. Scale bar is labeled in yellow on the EM micrographs. No discernible protein NPs are shown in any of the EM micrographs.

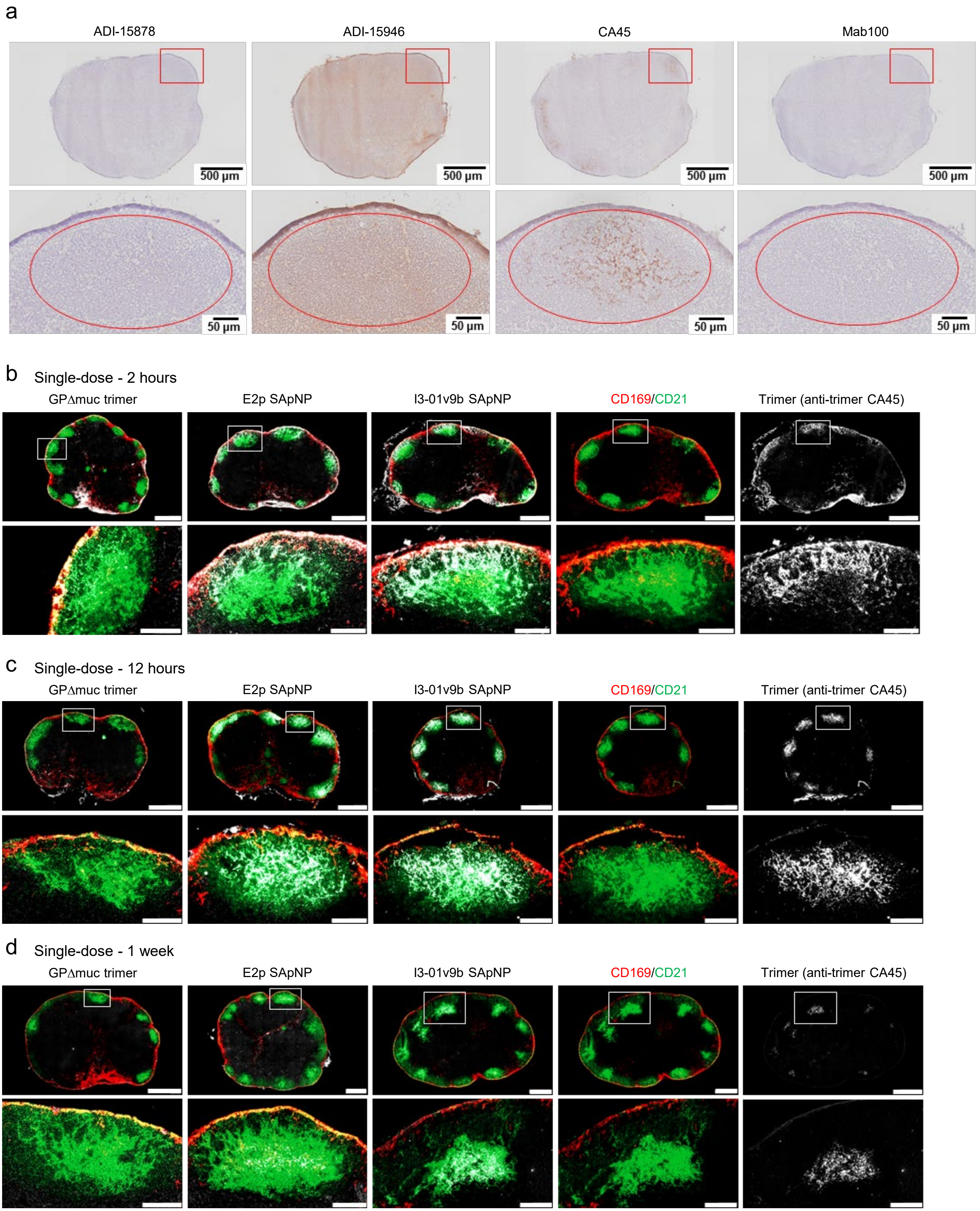
a
ADI-15878
ADI-15946
CA45
Mab100
500 µm
50 µm
b Single-dose - 2 hours
GPΔmuc trimer
E2p SApNP
I3-01v9b SApNP
CD169/CD21
Trimer (anti-trimer CA45)
c Single-dose - 12 hours
GPΔmuc trimer
E2p SApNP
I3-01v9b SApNP
CD169/CD21
Trimer (anti-trimer CA45)
d Single-dose - 1 week
GPΔmuc trimer
E2p SApNP
I3-01v9b SApNP
CD169/CD21
Trimer (anti-trimer CA45)

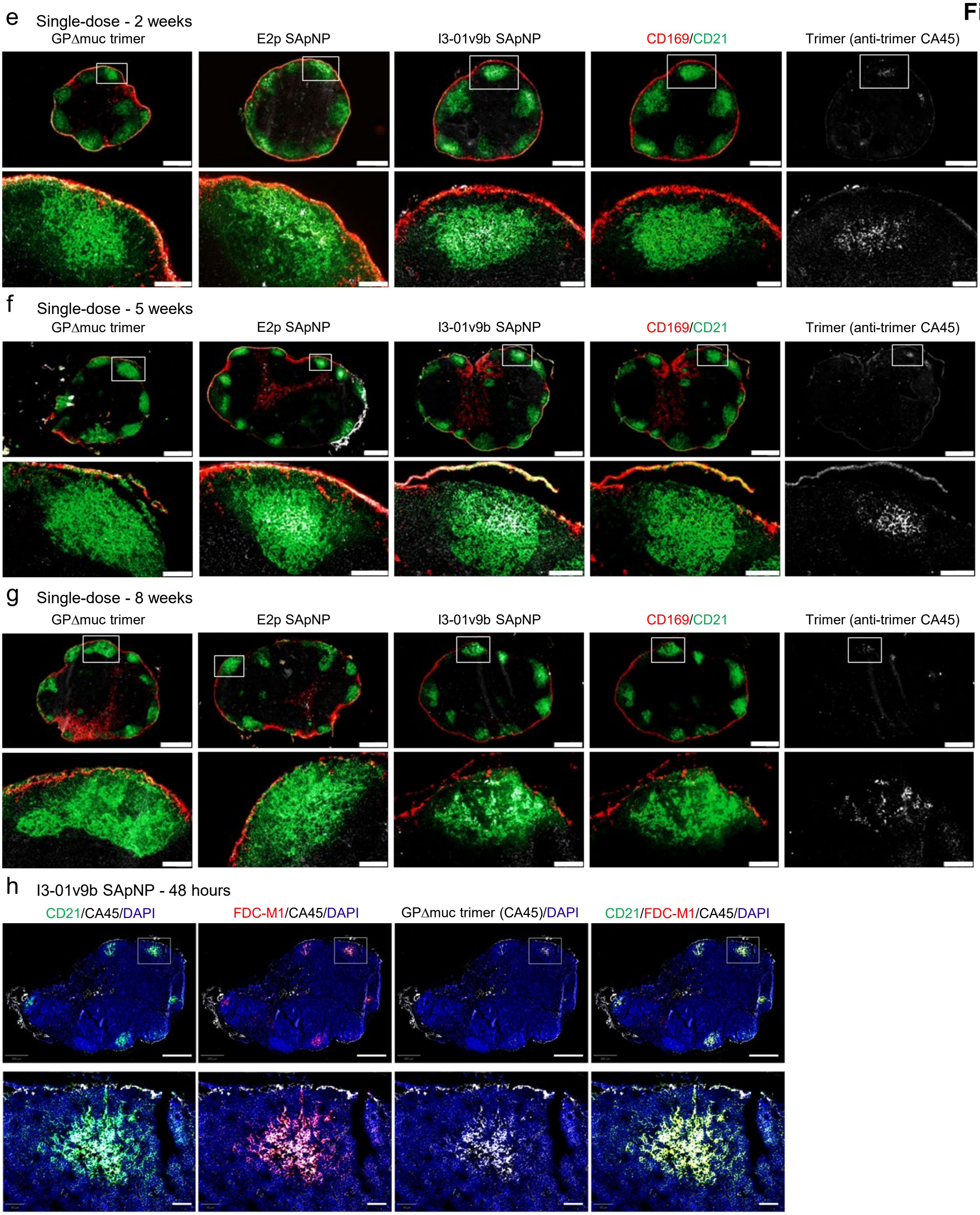


**Fig. S6. Immunohistological images of SUDV GPΔmuc trimer and SApNPs in lymph nodes.** (**a**) Immunostaining images of lymph node tissues from mice injected with SUDV GPΔmuc-presenting SApNPs, stained using antibodies ADI-15878, ADI-15946, CA45, and mAb100. **(b–g)** Colocalization of SUDV GPΔmuc trimer, E2p SApNP, and I3-01v9b SApNP with FDC networks in lymph node follicles at various time points following a single-dose injection (10 µg per injection, 40 µg total per mouse): (**b**) 2 hours, (**c**) 12 hours, (**d**) 1 week, (**e**) 2 weeks, (**f**) 5 weeks, and (**g**) 8 weeks. Immunofluorescent images are pseudo-color-coded as follows: CD21$^{+}$ (green), CD169$^{+}$ (red), and CA45 (white). (**h**) Colocalization of SUDV GPΔmuc-presenting I3-01v9b SApNPs (labeled by CA45, white) with FDC networks labeled by CD21$^{+}$ (green) and FDC-M1$^{+}$ (red) at 48 hours post-injection. Scale bars: 500 µm (entire lymph node) and 100 µm (enlarged follicle image).

**Fig. S7**

**a** E2p SApNPs (yellow arrow) are aligned on FDC dendrites at 2 hours after injection

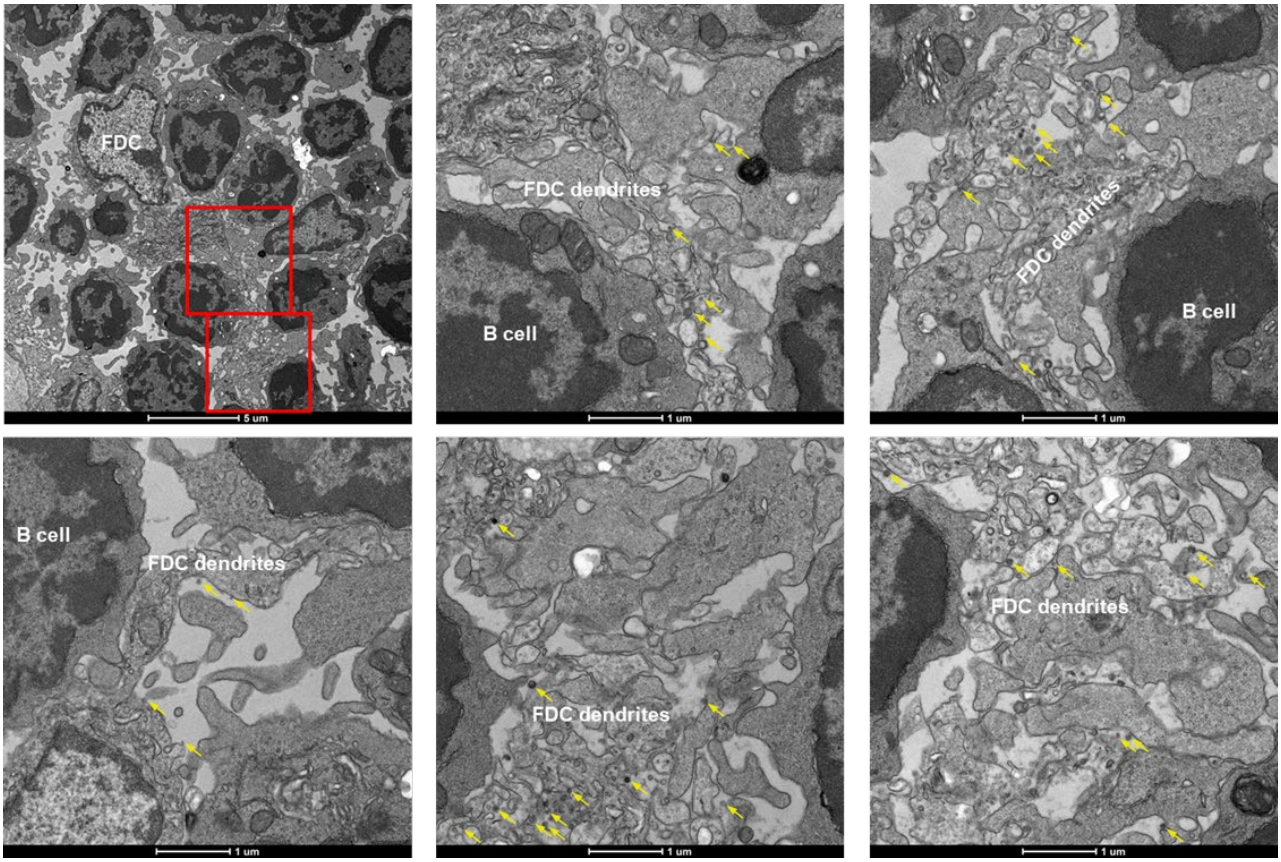


**b** E2p SApNPs (yellow arrow) are aligned on FDC dendrites at 12 hours after injection

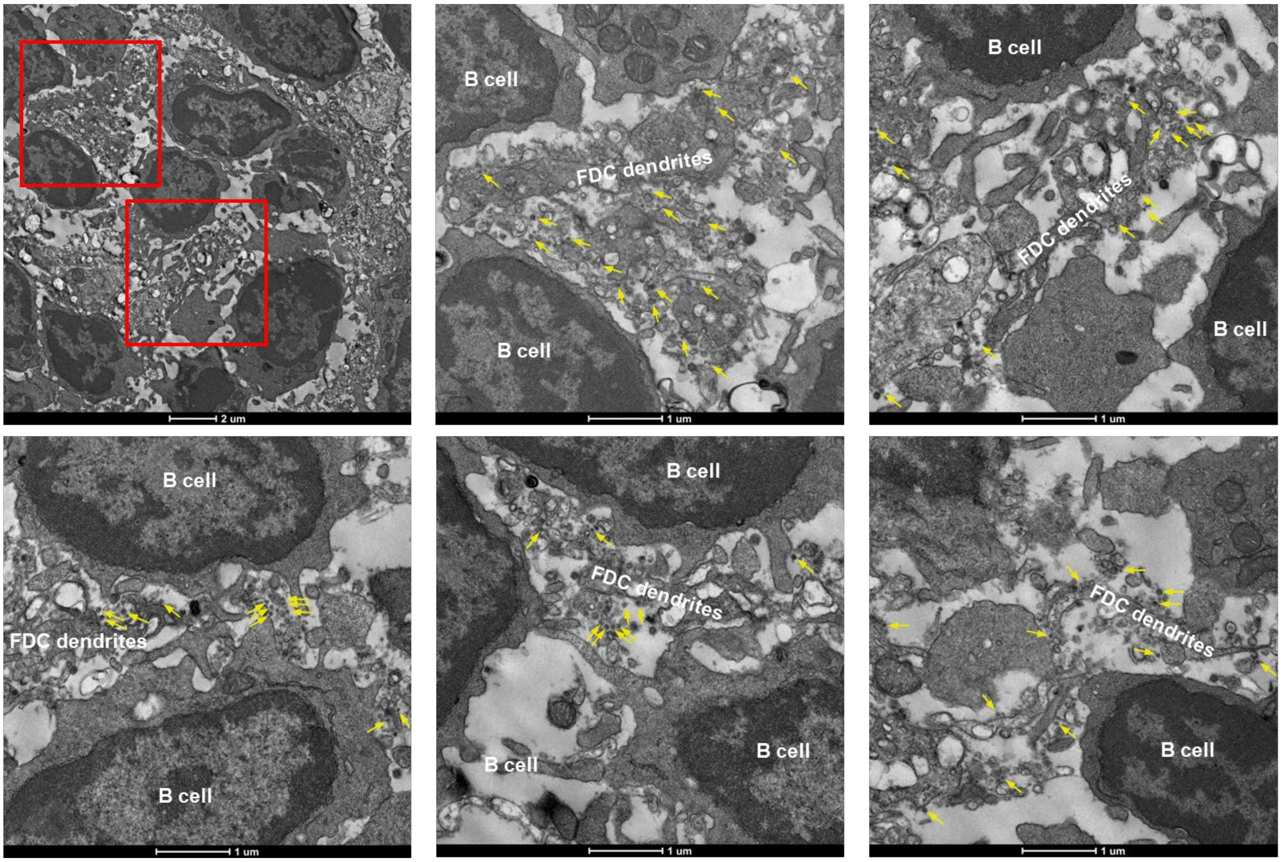

c E2p SApNPs (yellow arrow) are aligned on FDC dendrites or taken up by B cells at 48 hours after injection

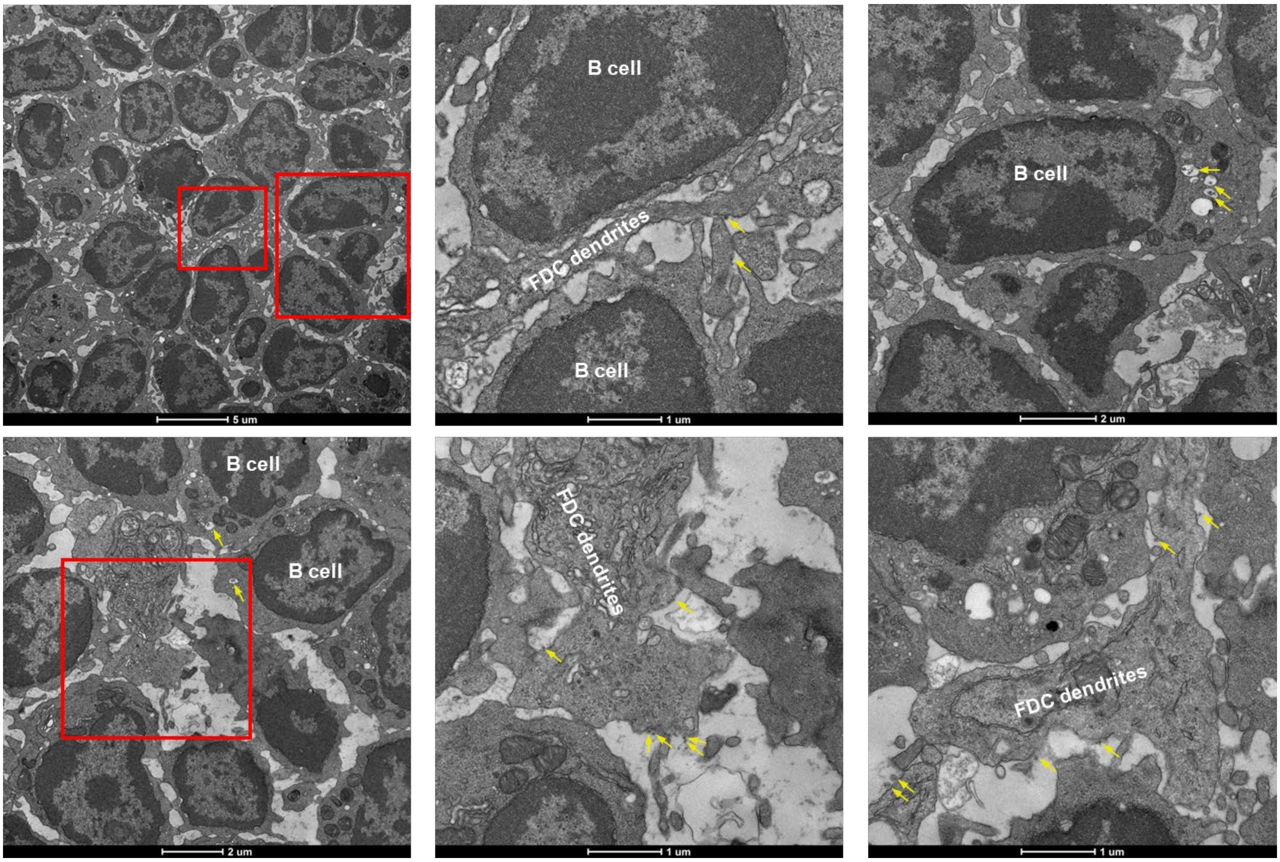


d I3-01v9b SApNPs (yellow arrow) are aligned on FDC dendrites or inside endolysosomes of FDC dendrites at 2 hours after injection

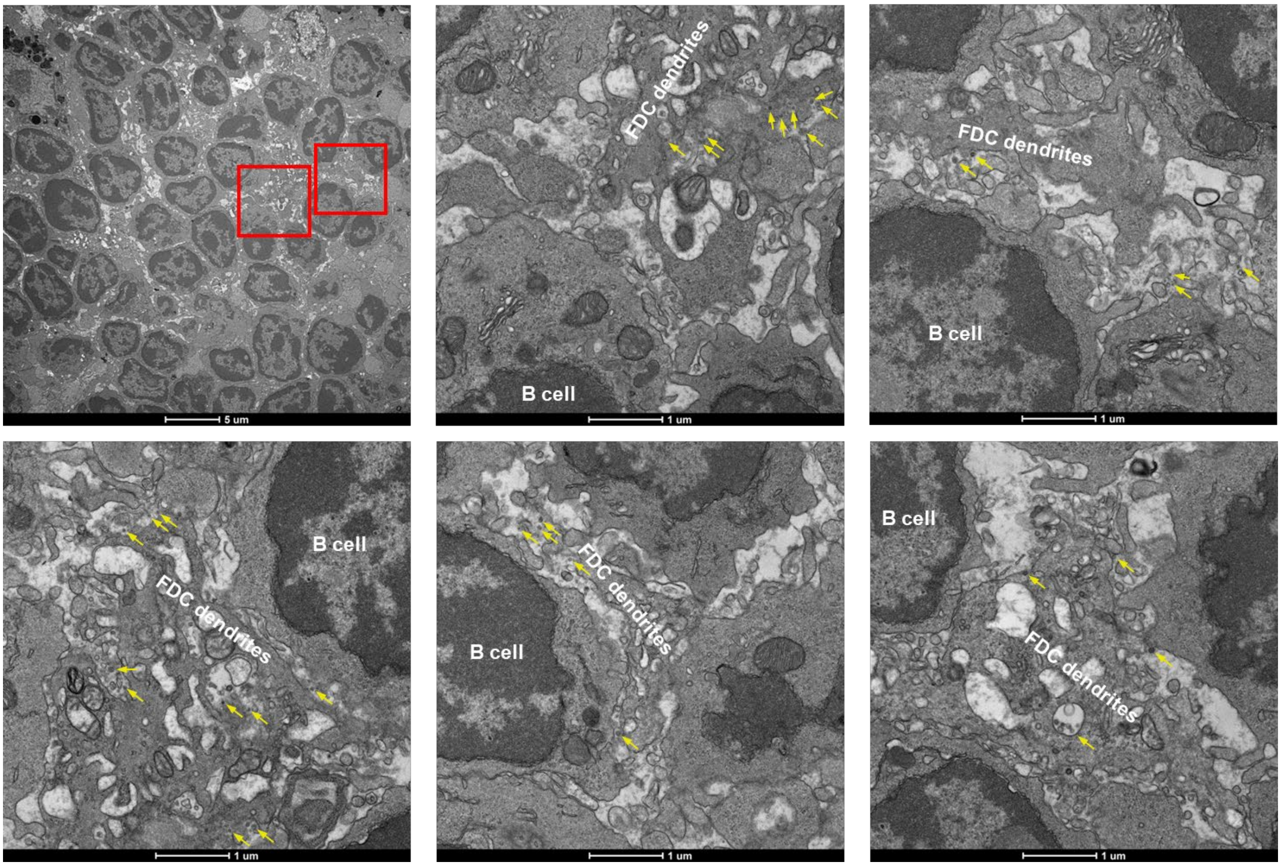

**Fig. S7**

e I3-01v9b SApNPs (yellow arrow) are aligned on FDC dendrites or taken up by B cells at 12 hours after injection

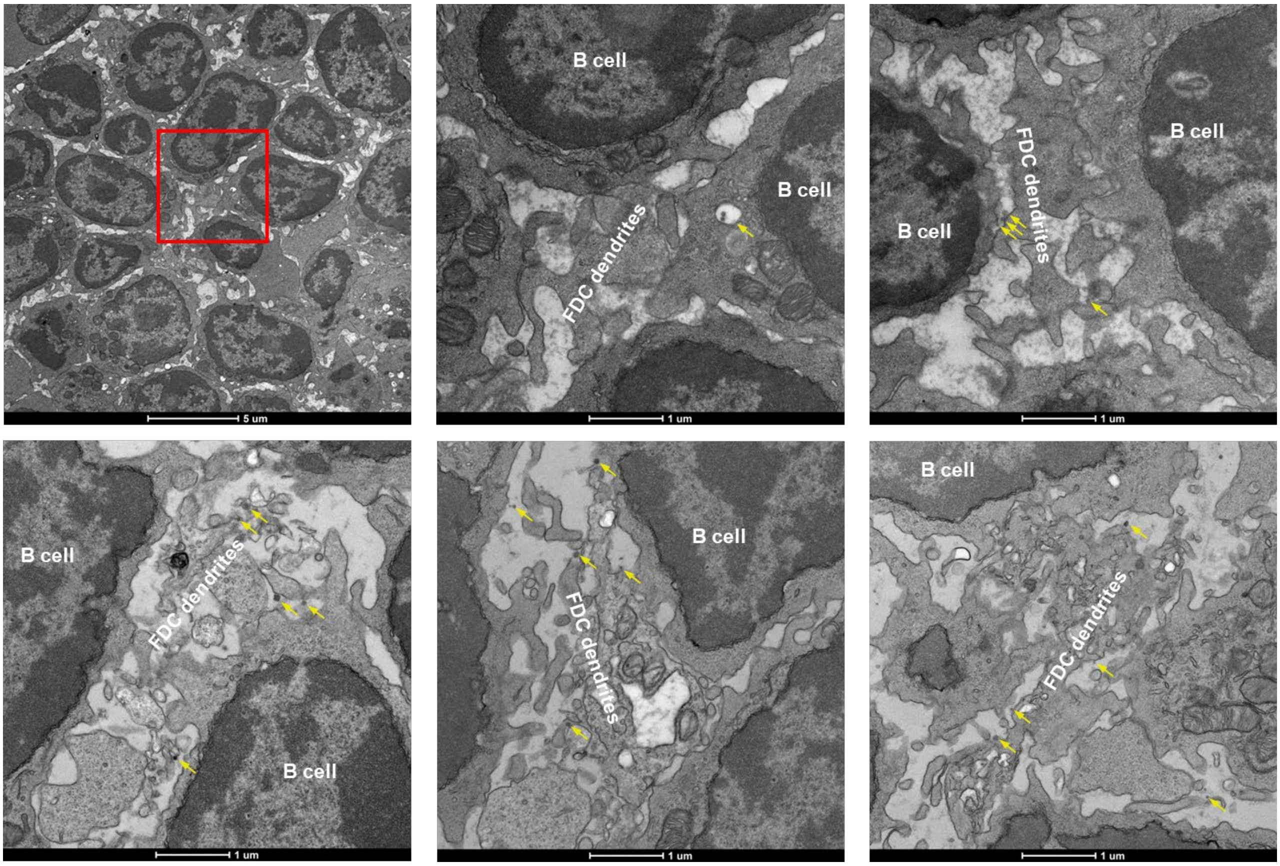


f I3-01v9b SApNPs (yellow arrow) are aligned on FDC dendrites at 48 hours after injection

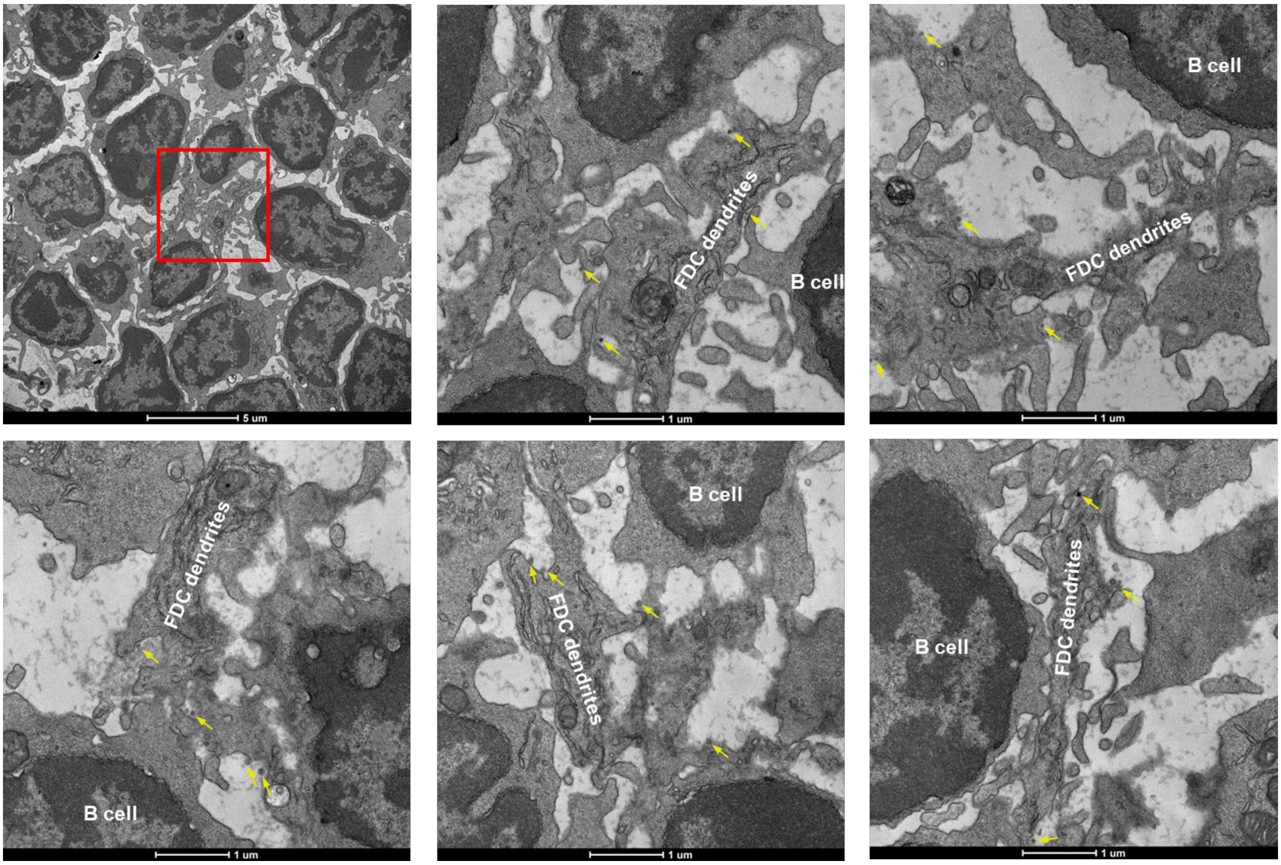

g E2p SApNPs (yellow arrow) are associated with phagocytic cells at 2 hours after injection

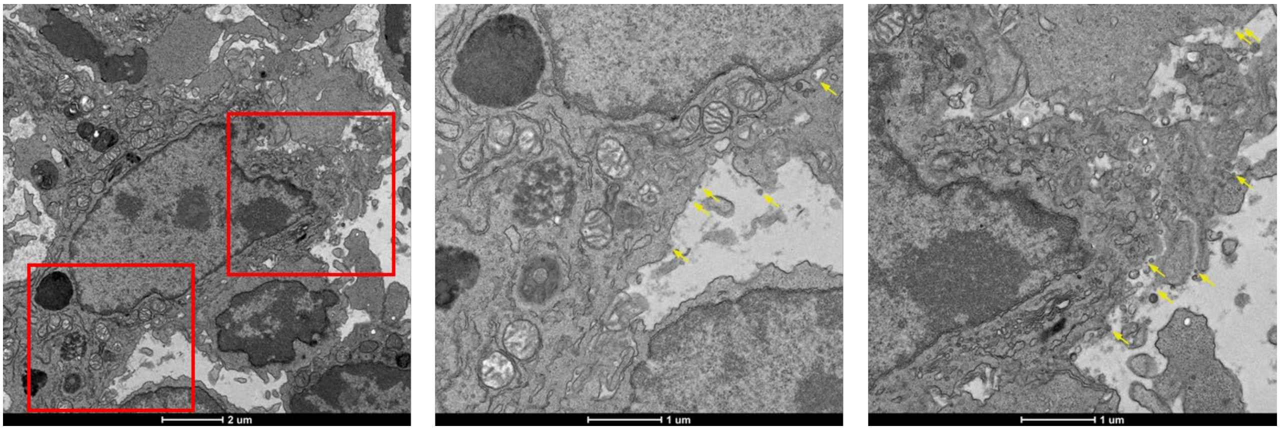


h E2p SApNPs (yellow arrow) are associated with phagocytic cells at 12 hours after injection

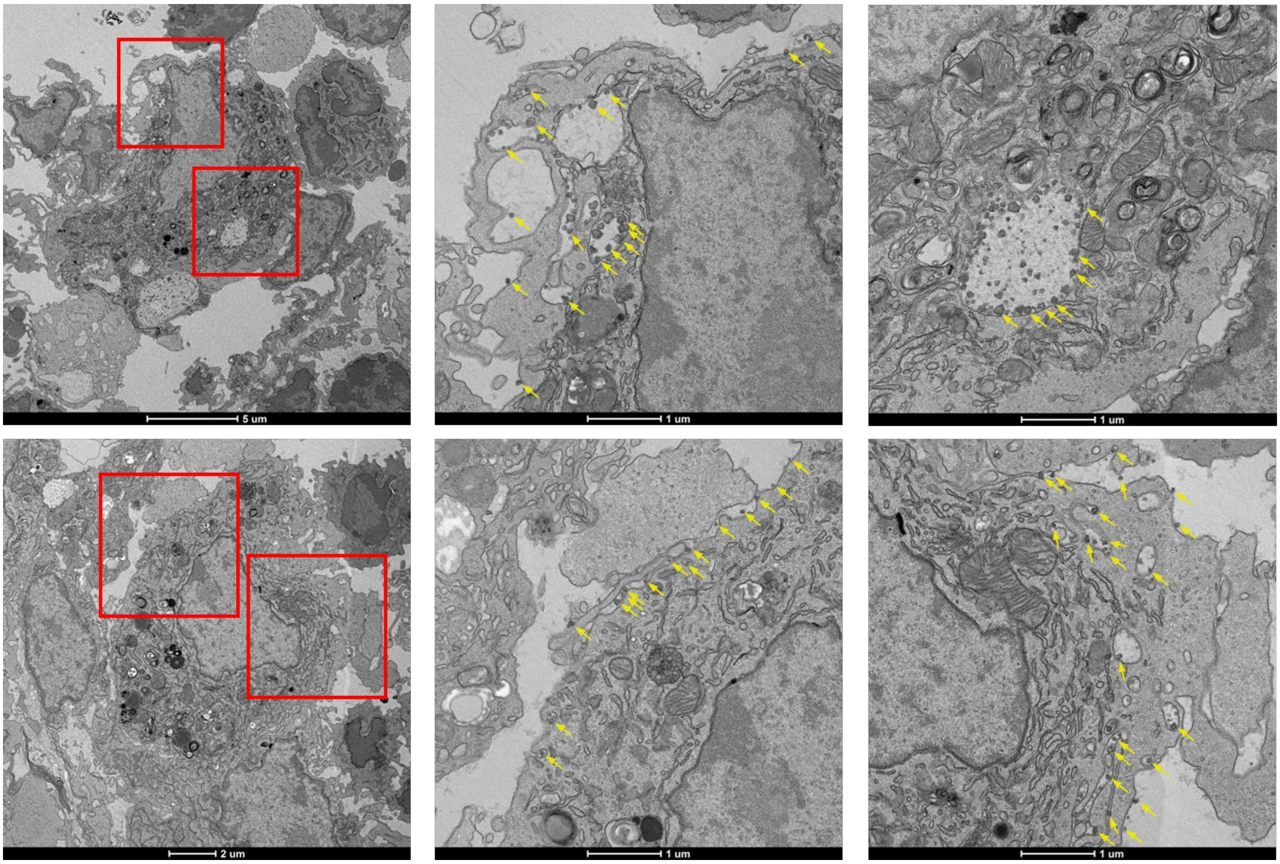

i E2p SApNPs (yellow arrow) are associated with phagocytic cells at 48 hours after injection

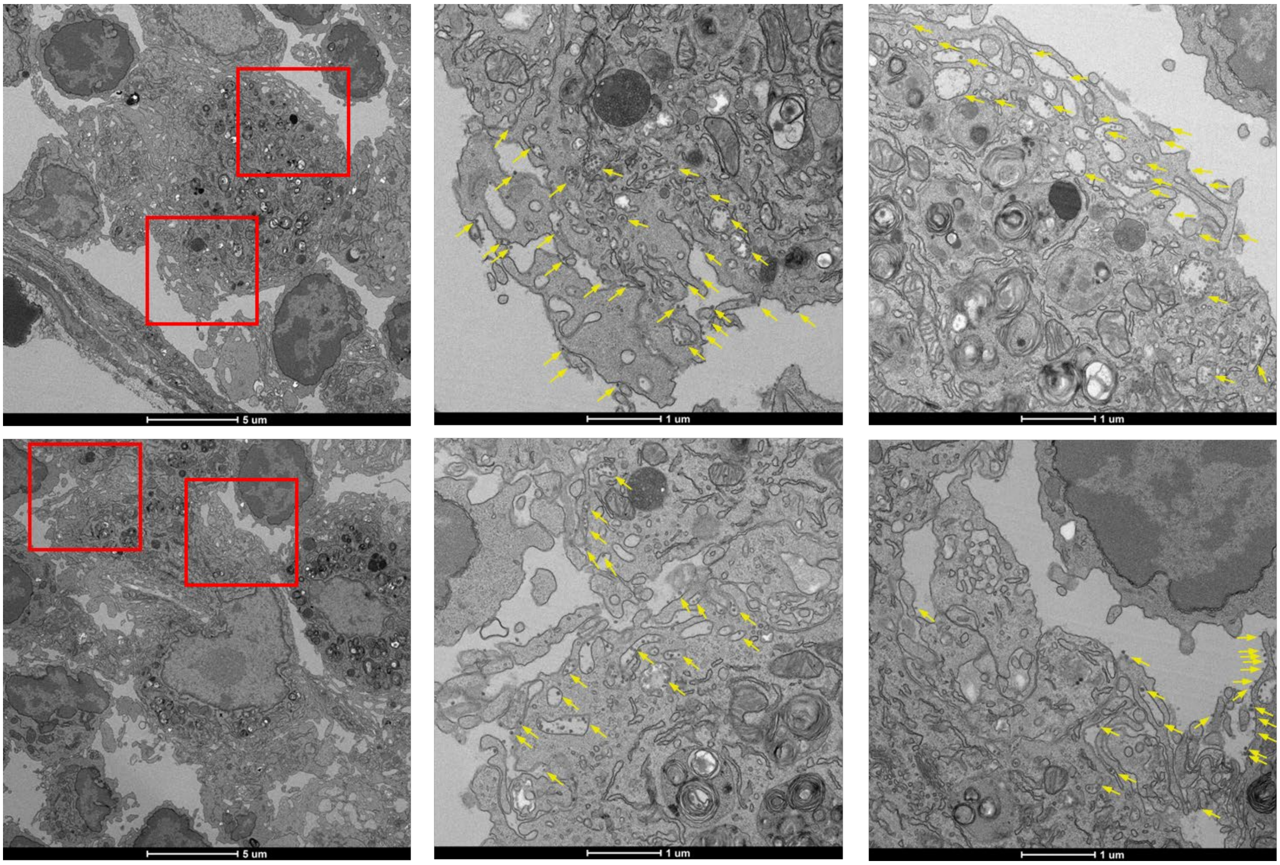


j I3-01v9b SApNPs (yellow arrow) are associated with phagocytic cells at 2 hours after injection

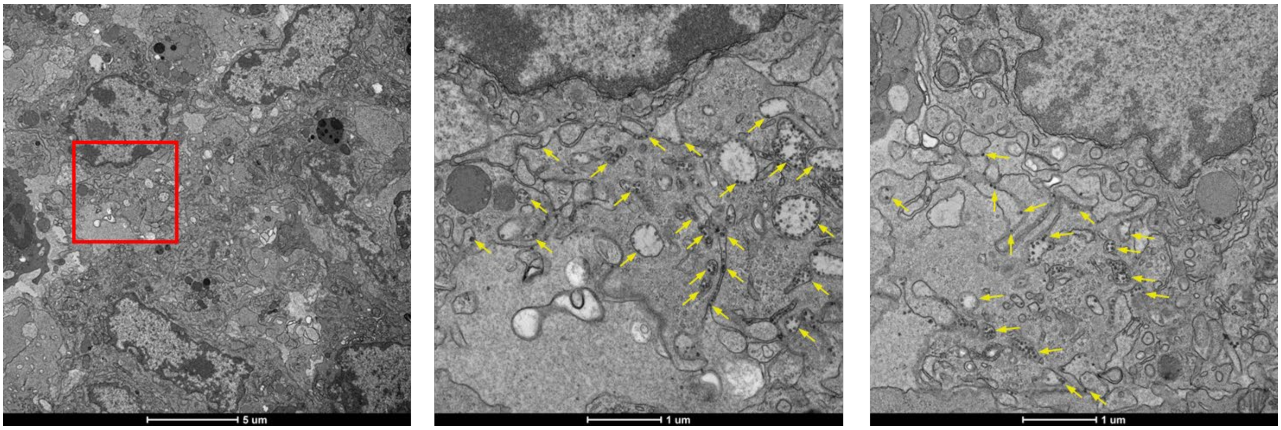

**Fig. S7**

k I3-01v9b SApNPs (yellow arrow) are associated with phagocytic cells at 12 hours after injection

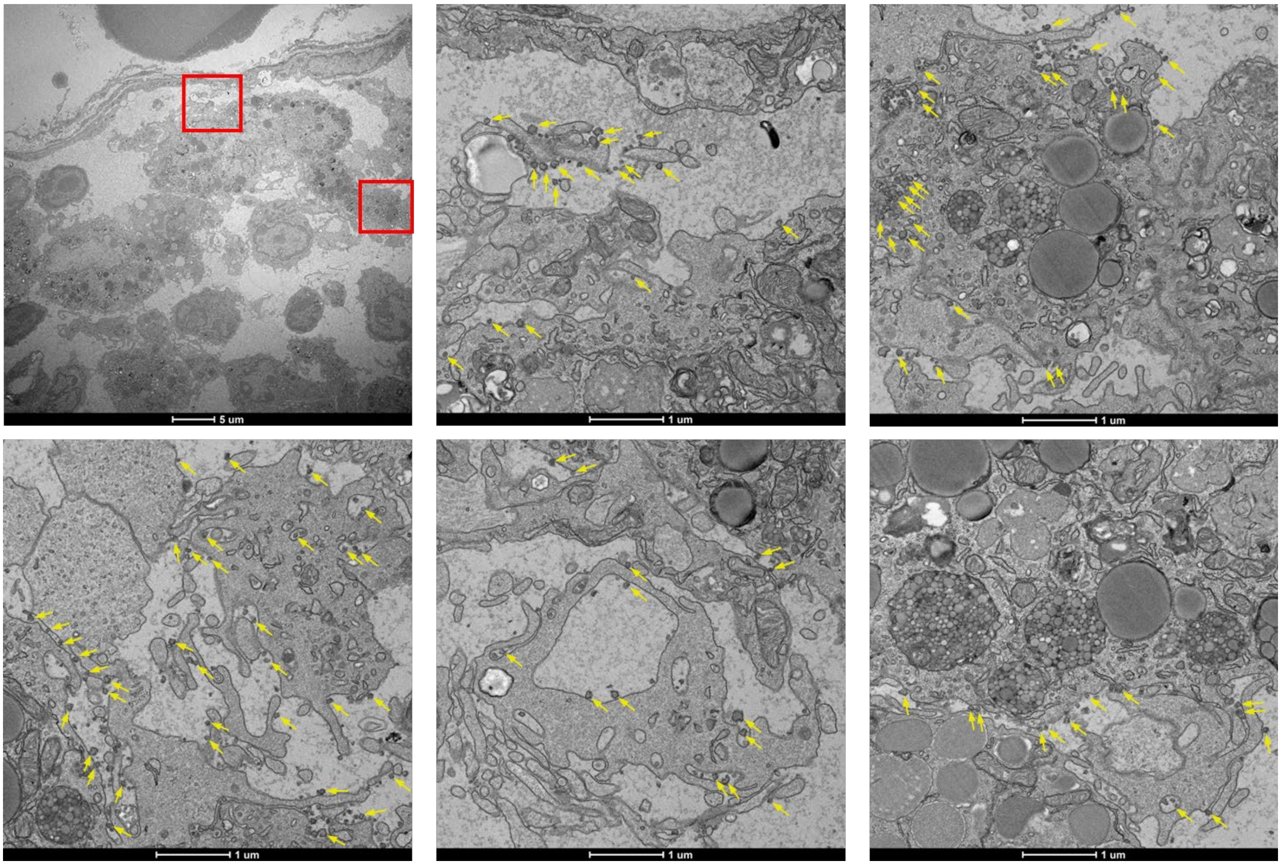


l I3-01v9b SApNPs (yellow arrow) are associated with phagocytic cells at 48 hours after injection

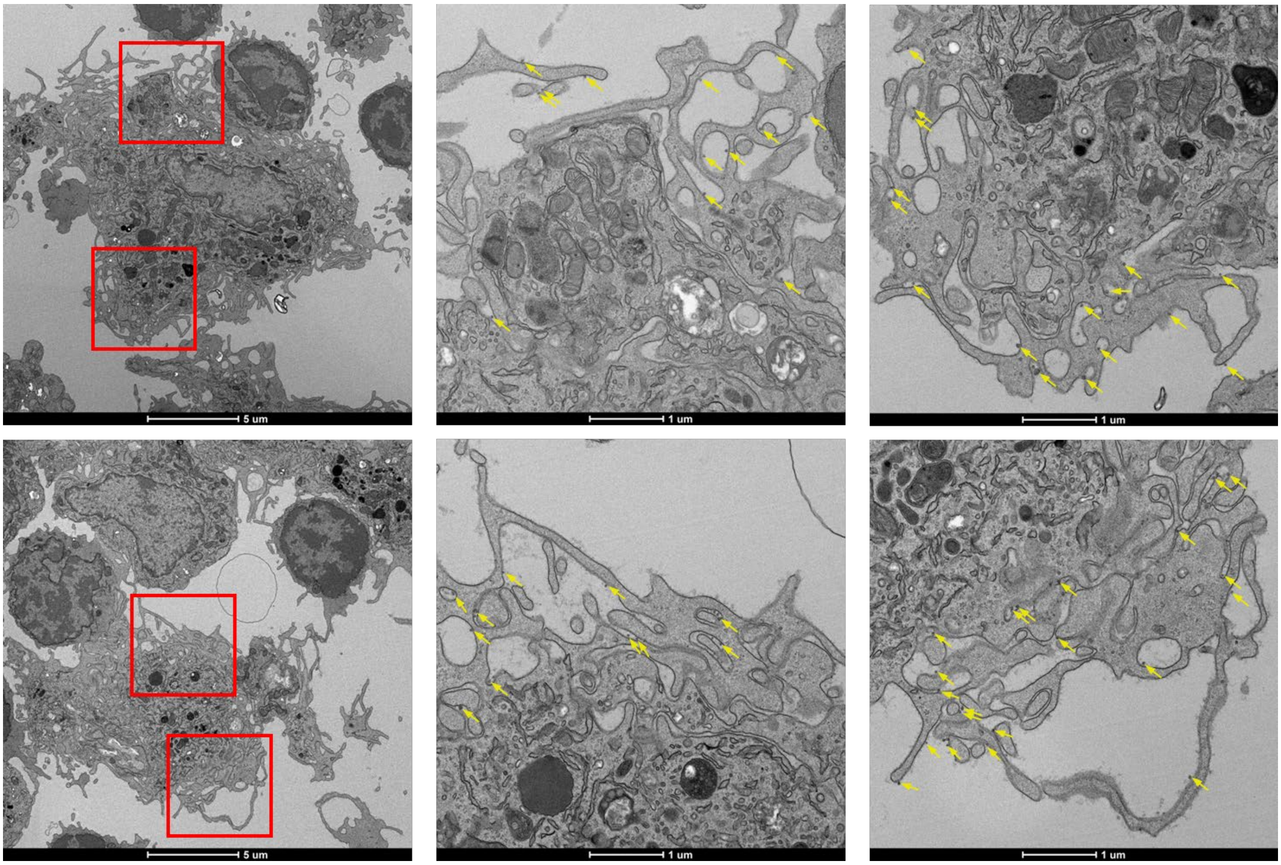

**Fig. S7**

m Aluminum phosphate (green arrow) tends to aggregate in the ECM at 2 hours after injection

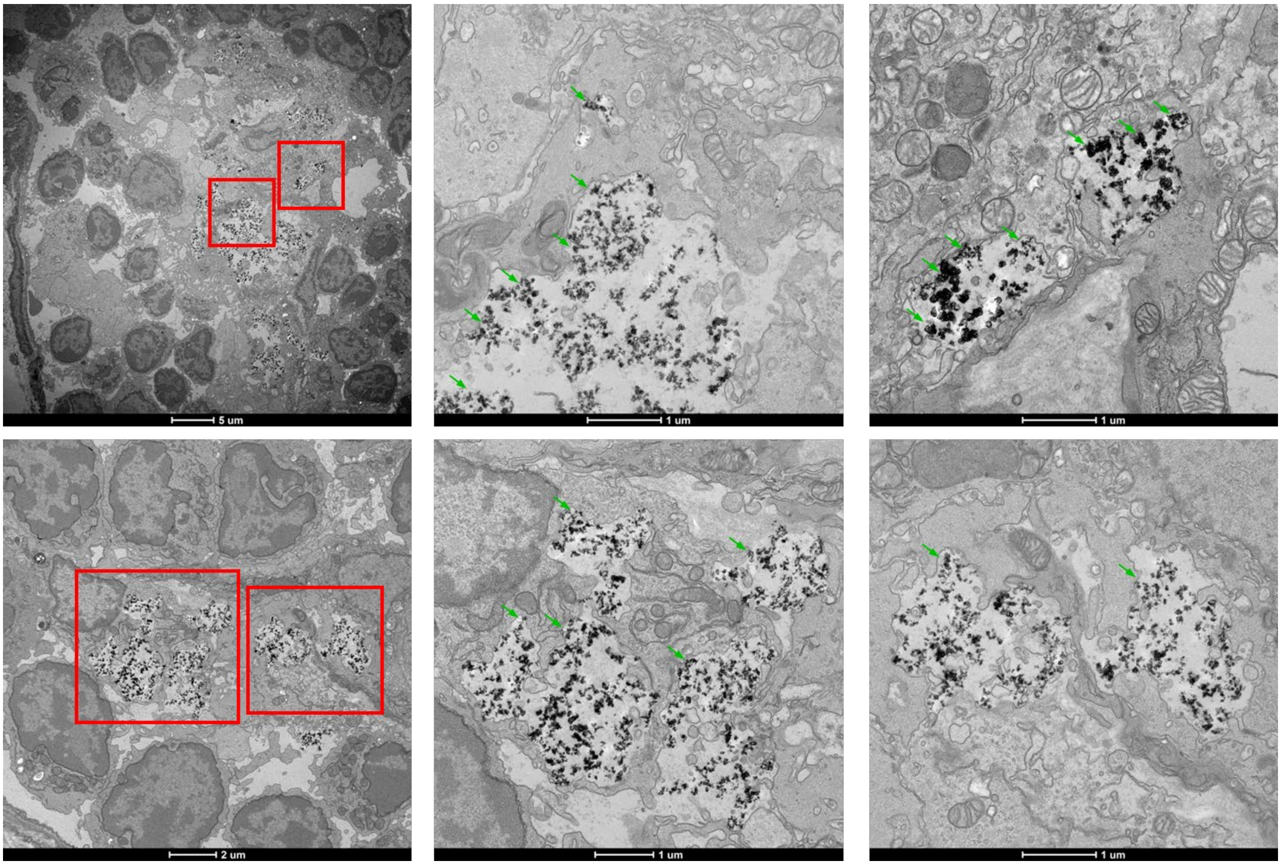


n I3-01v9b SApNPs (yellow arrow) and aluminum phosphate (green arrow) with phagocytic cells at 2 hours after injection

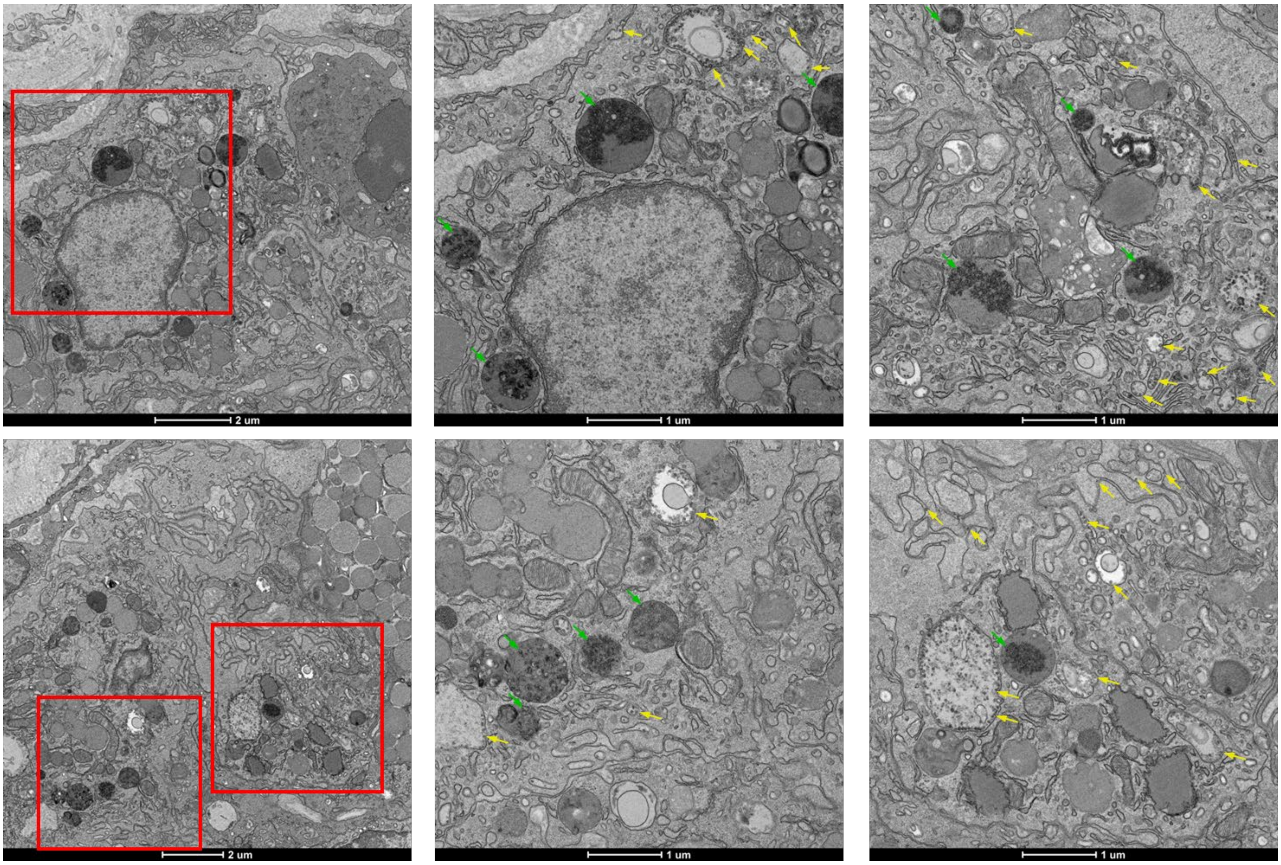

**Fig. S7**

o I3-01v9b SApNPs (yellow arrow) and aluminum phosphate (green arrow) with phagocytic cells at 12 hours after injection

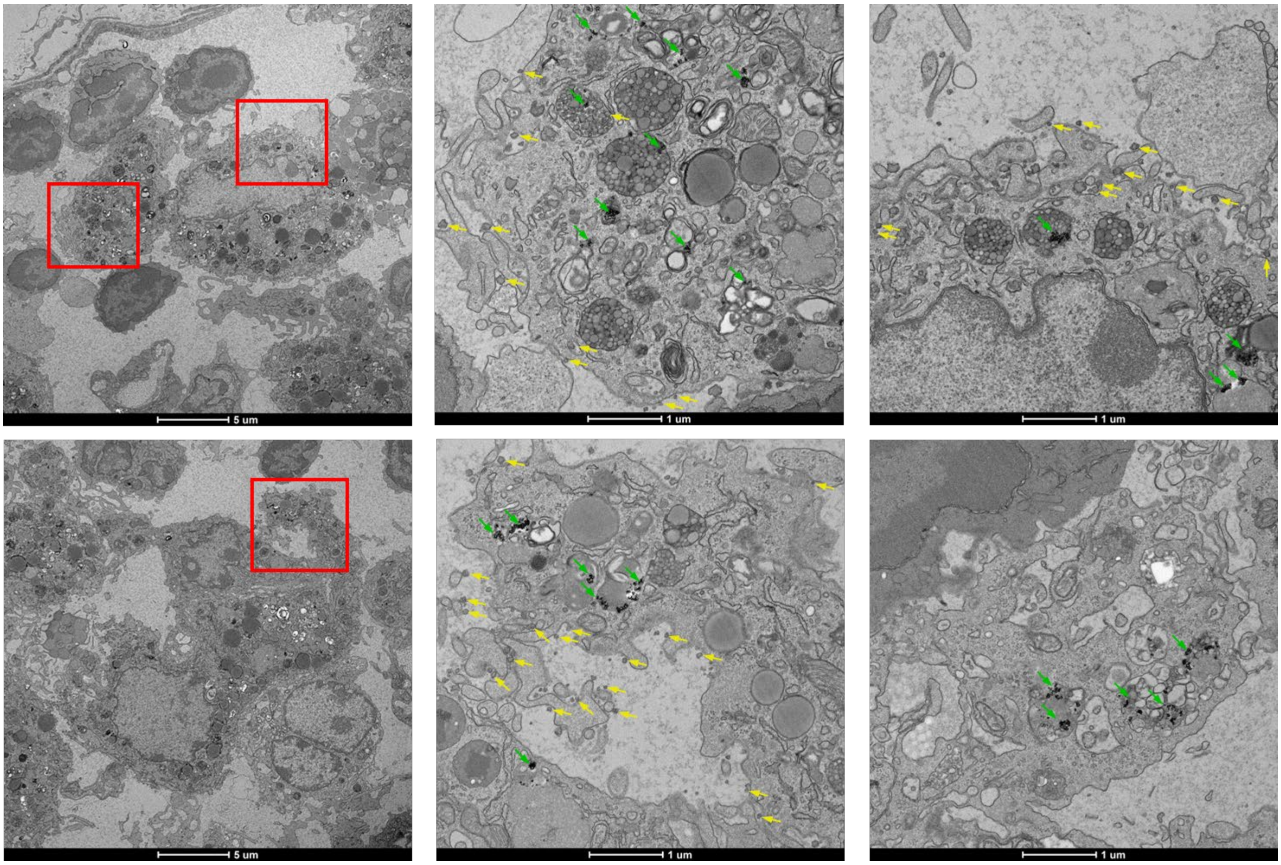


p I3-01v9b SApNPs (yellow arrow) and aluminum phosphate (green arrow) with phagocytic cells at 48 hours after injection

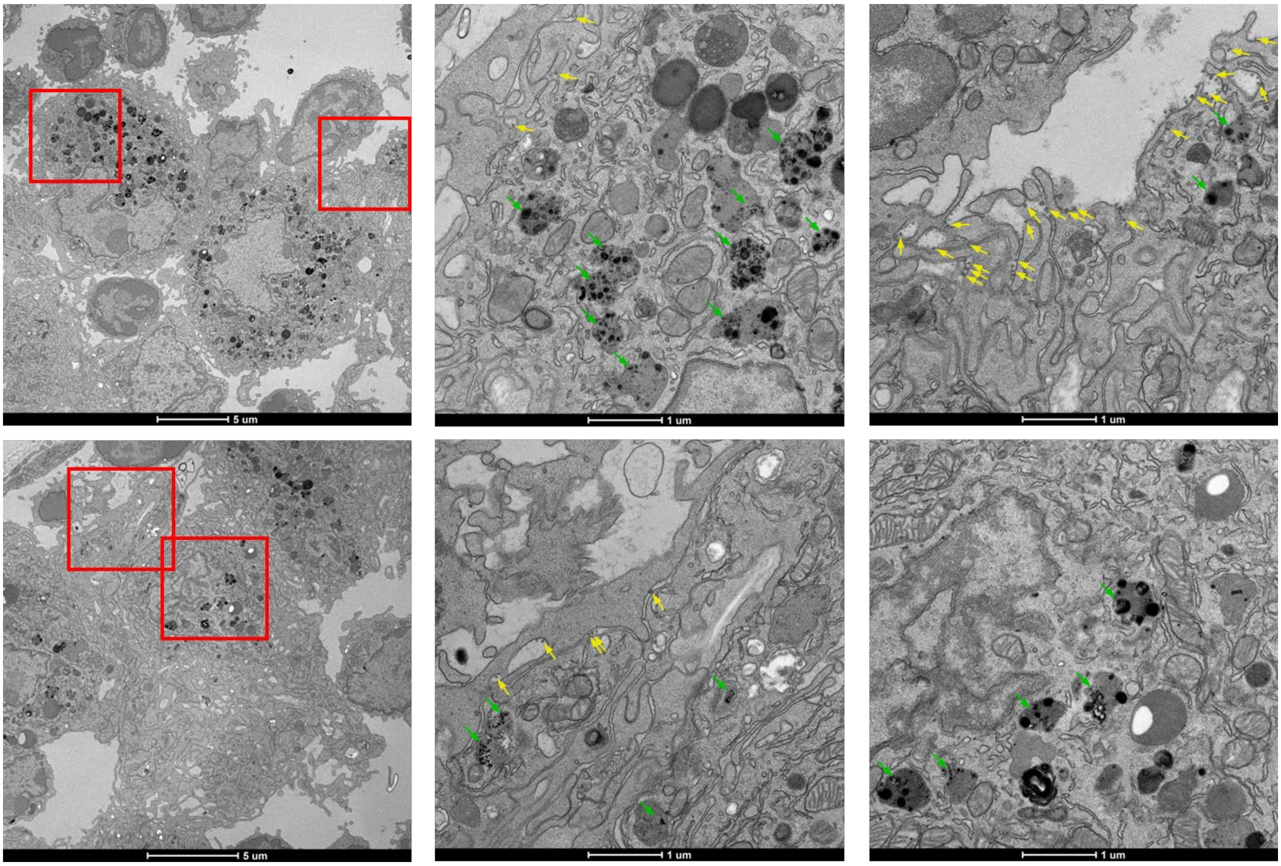

**Fig. S7. TEM analysis of SUDV GPΔmuc-presenting SApNPs interacting with FDCs and phagocytic cells in lymph nodes.** TEM images show the distribution and cellular interactions of SUDV GPΔmuc-presenting SApNPs following a single-dose injection (2 footpads, 50 µg/footpad). (**a–c**) E2p SApNPs aligned along FDC dendrites at 2 hours, 12 hours, and 48 hours post-injection, respectively. (**d–f**) I3-01v9b SApNPs aligned on FDC dendrites at the same time points. Aluminum phosphate (AP) adjuvant particles are not observed in association with SApNPs on FDC dendrites. (**g–i**) E2p SApNPs located on the surface or within endolysosomes of phagocytic cells at 2 hours, 12 hours, and 48 hours post-injection, respectively. (**j–l**) I3-01v9b SApNPs associated with phagocytic cells at the same time points. (**m**) AP adjuvant particles aggregating in the extracellular matrix 2 hours after injection. (**n–p**) I3-01v9b SApNPs formulated with AP observed within phagocytic cells at 2 hours, 12 hours, and 48 hours post-injection. TEM imaging was performed on two popliteal lymph nodes per SApNP construct. E2p and I3-01v9b SApNPs are indicated by yellow arrows; AP adjuvant particles are indicated by green arrows.

Fig. S8

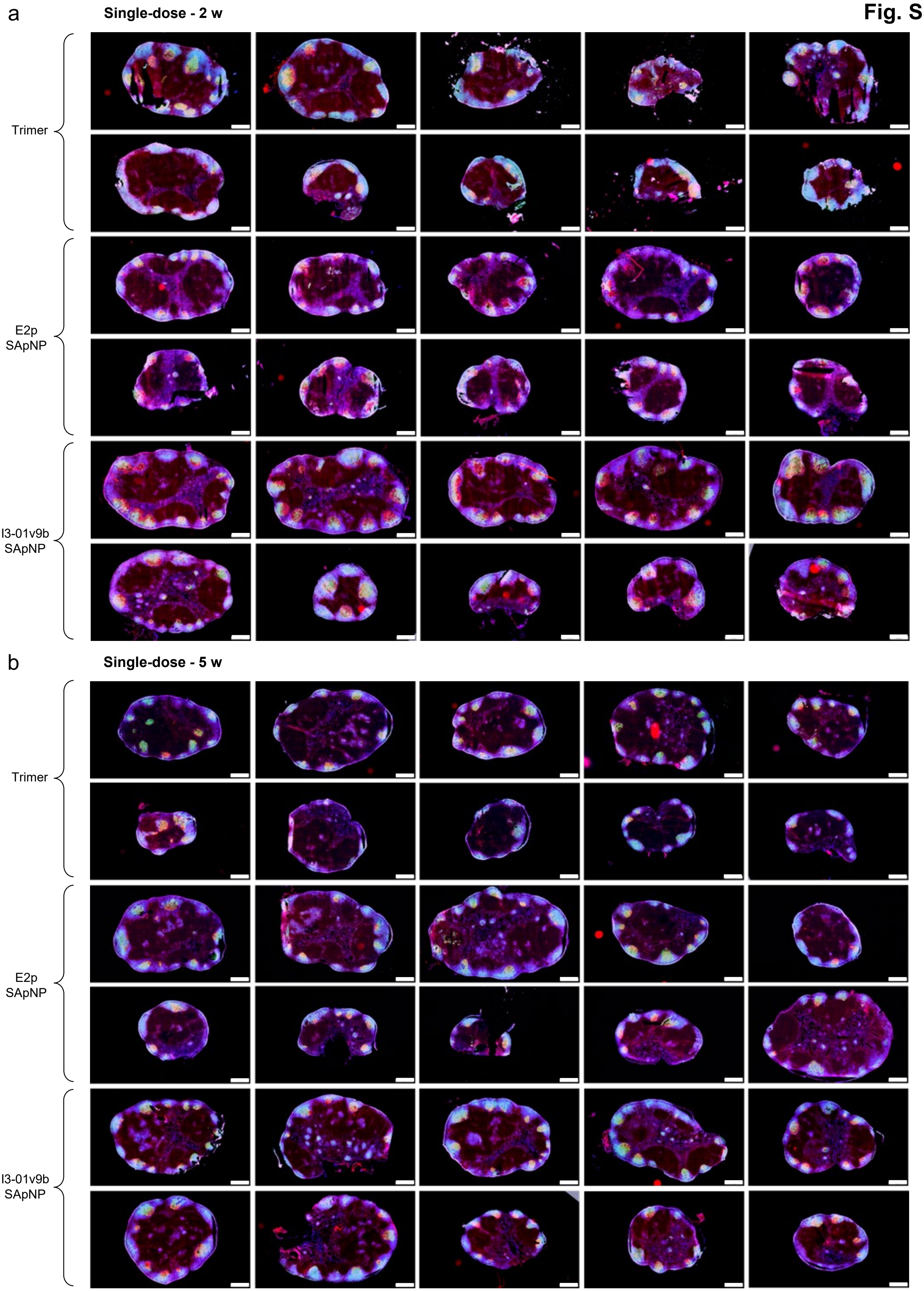
a
Single-dose - 2 w
Trimer
E2p
SApNP
I3-01v9b
SApNP
b
Single-dose - 5 w
Trimer
E2p
SApNP
I3-01v9b
SApNP

Fig. S8

c Single-dose - 8 w

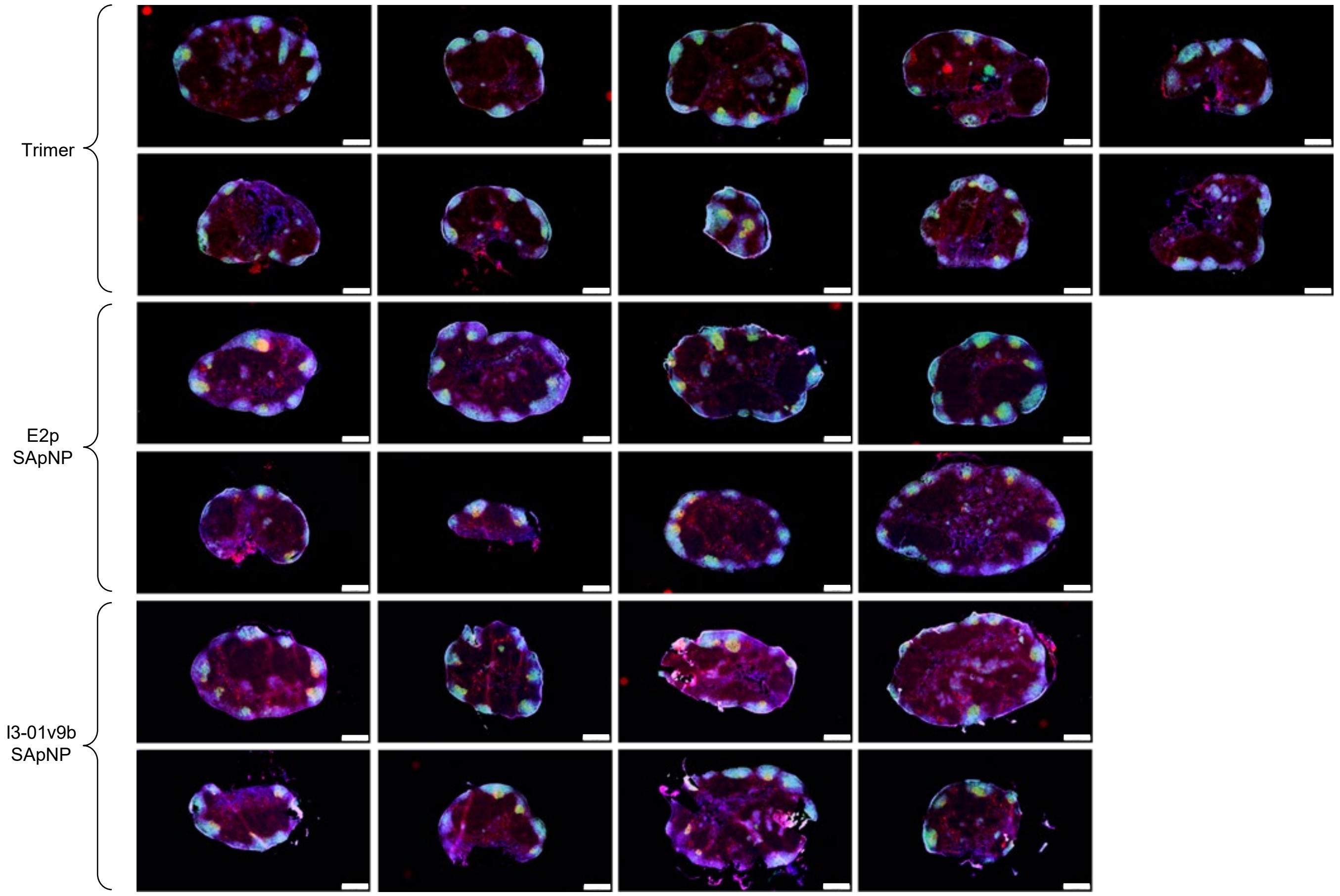


d Prime-boost - 3 w + 2 w

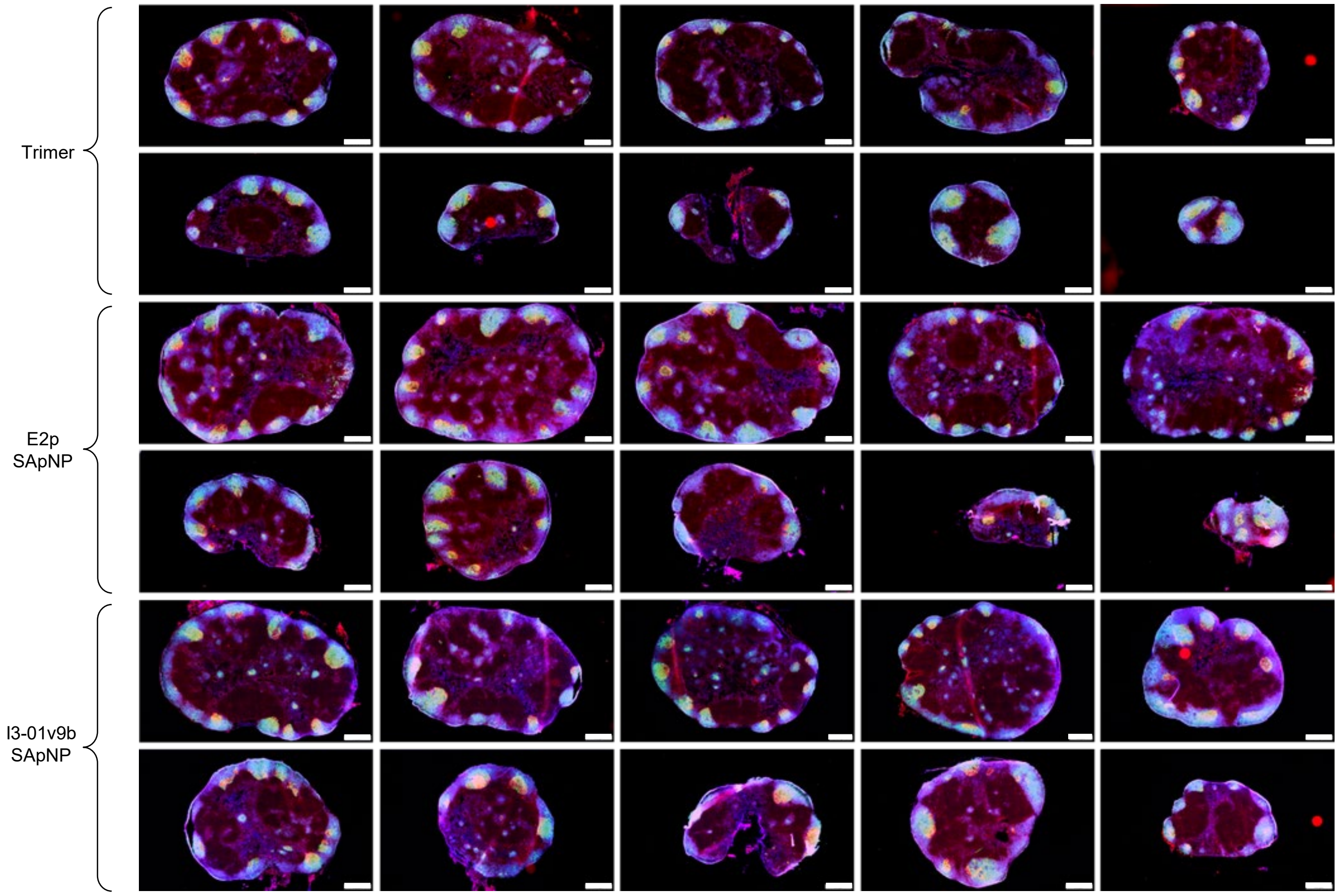

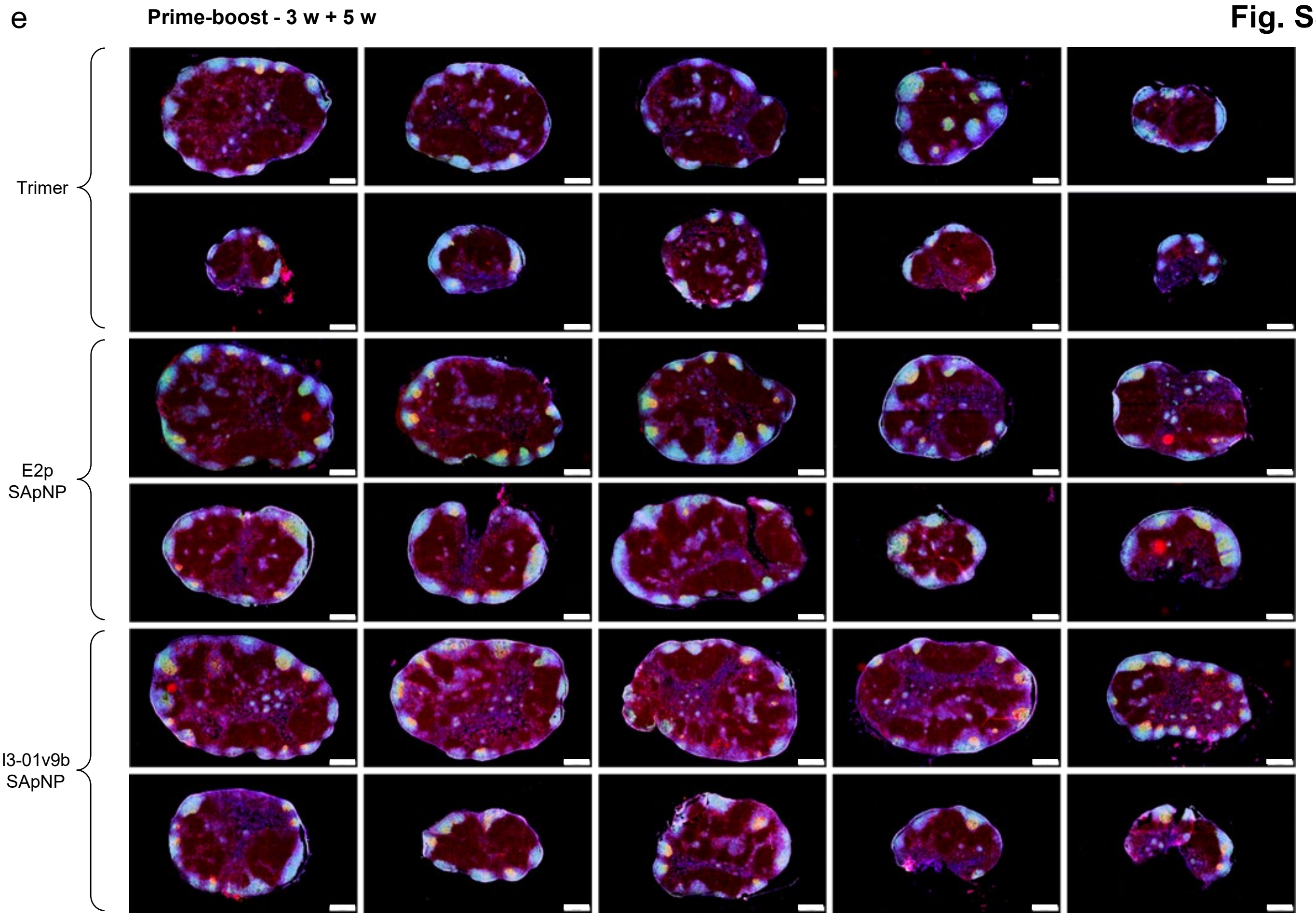


**Fig. S8. Immunohistological analysis of SUDV GPΔmuc trimer and SApNP vaccine-induced germinal centers (GCs).** Immunohistological images of GCs at (**a**) 2, (**b**) 5, and (**c**) 8 weeks following a single-dose injection of GPΔmuc trimer or GPΔmuc-presenting E2p and I3-01v9b SApNPs (10 μg per injection; total of 40 μg per mouse). GC images at (**d**) 2 and (**e**) 5 weeks after a booster dose, administered 3 weeks after the initial immunization (n = 5 mice/group). Immunofluorescent images are pseudo-color coded ($GL7^{+}$, red; $CD21^{+}$, green; B220, blue). Scale bars = 500 μm for each lymph node image.

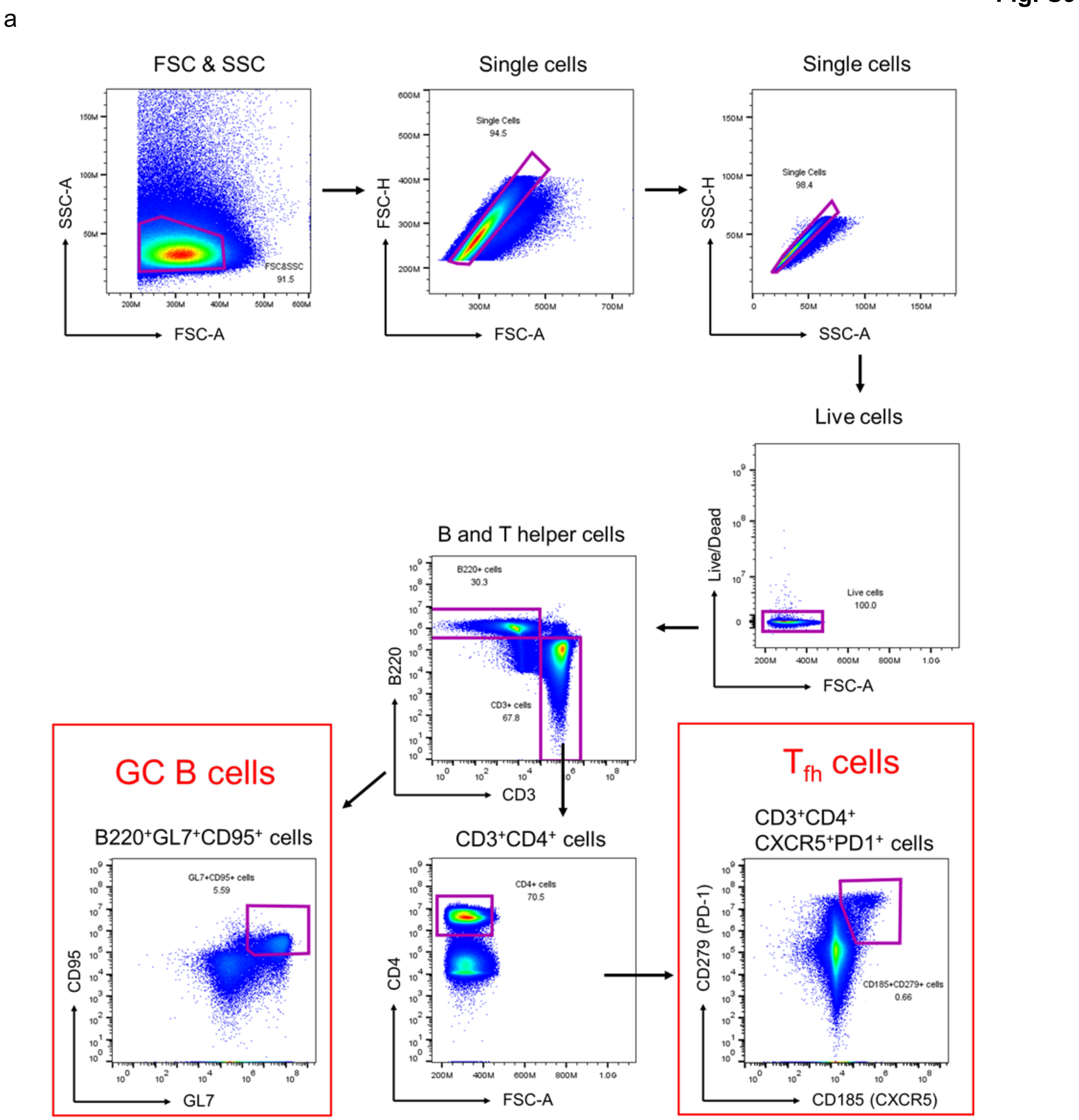


**Fig. S9. Flow cytometry analysis of germinal centers (GCs) induced by SUDV GPΔmuc trimer and SApNPs vaccines.** Gating strategy for identifying GC B cells and T follicular helper (Tfh) cells by flow cytometry (n = 5 mice per group).

**Fig. S10**

**a Sera of individual mice immunized with EBOV vaccines binding to EBOV GPΔmuc-WL$^2$P$^4$(1TD0) trimer**

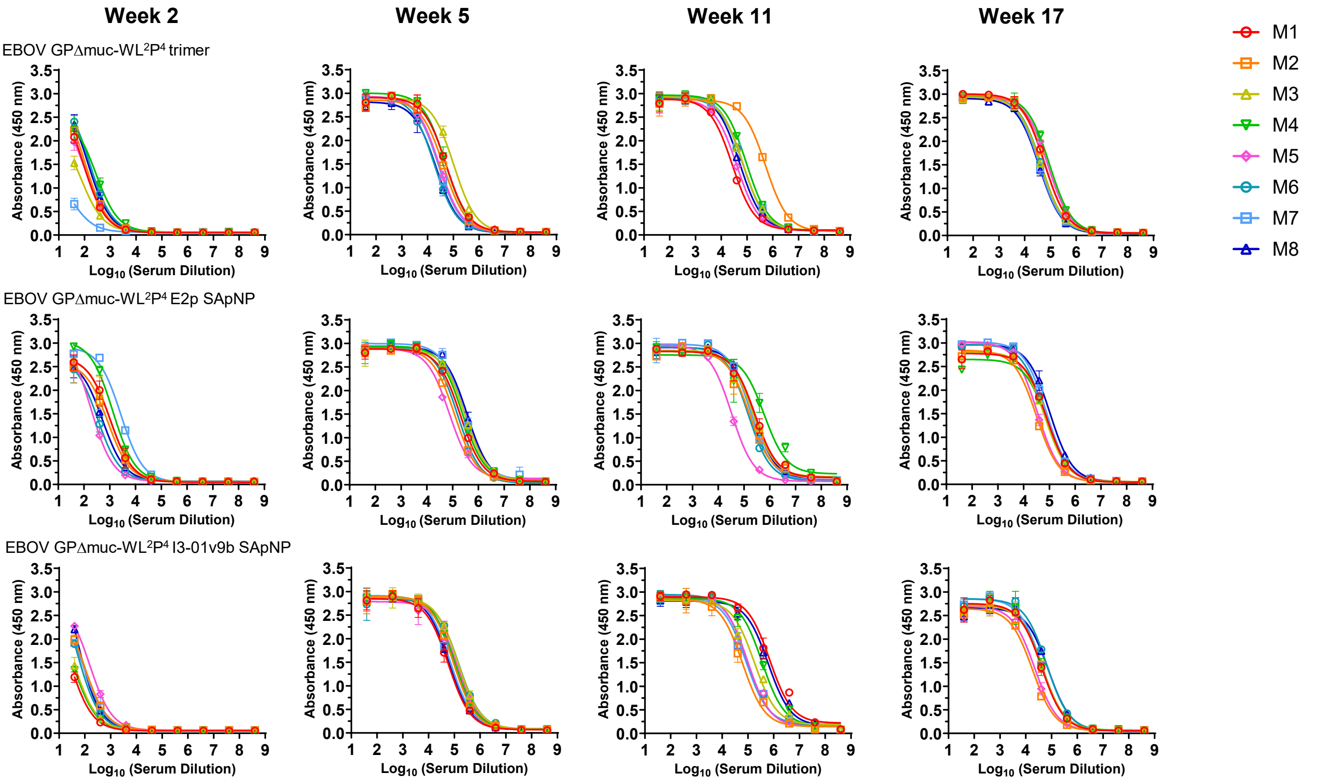


**b Mouse serum ELISA EC$_{50}$ titers**

**Week 2**

| Antigen | EC$_{50}$ titers (week 2) | | | | | | | | Geometric Mean |
|---|---|---|---|---|---|---|---|---|---|
| | M1 | M2 | M3 | M4 | M5 | M6 | M7 | M8 | |
| EBOV GPΔmuc-WL$^2$P$^4$ trimer | 111.7 | 136.3 | 51.3 | 227.2 | 103.0 | 200.2 | 12.6 | 173.3 | 97.3 |
| EBOV GPΔmuc-WL$^2$P$^4$ E2p SApNP | 985.3 | 706.4 | 970.8 | 1772.0 | 271.5 | 336.1 | 3145.0 | 486.4 | 799.6 |
| EBOV GPΔmuc-WL$^2$P$^4$ I3-01v9b SApNP | 30.1 | 101.8 | 43.0 | 37.9 | 172.4 | 80.7 | 91.5 | 115.6 | 72.1 |

**Week 5**

| Antigen | EC$_{50}$ titers (week 5) | | | | | | | | Geometric Mean |
|---|---|---|---|---|---|---|---|---|---|
| | M1 | M2 | M3 | M4 | M5 | M6 | M7 | M8 | |
| EBOV GPΔmuc-WL$^2$P$^4$ trimer | 52545 | 41735 | 104914 | 48787 | 27947 | 18271 | 26144 | 21434 | 36588.3 |
| EBOV GPΔmuc-WL$^2$P$^4$ E2p SApNP | 199849 | 120954 | 303168 | 237335 | 74446 | 156667 | 282413 | 368341 | 195227.1 |
| EBOV GPΔmuc-WL$^2$P$^4$ I3-01v9b SApNP | 58458 | 93275 | 140476 | 104463 | 128829 | 148277 | 70114 | 63709 | 95341.8 |

**Week 11**

| Antigen | EC$_{50}$ titers (week 11) | | | | | | | | Geometric Mean |
|---|---|---|---|---|---|---|---|---|---|
| | M1 | M2 | M3 | M4 | M5 | M6 | M7 | M8 | |
| EBOV GPΔmuc-WL$^2$P$^4$ trimer | 26545 | 524033 | 69054 | 94400 | 39464 | 69865 | 68351 | 51944 | 73880.6 |
| EBOV GPΔmuc-WL$^2$P$^4$ E2p SApNP | 267135 | 163352 | 223491 | 606328 | 32762 | 143908 | 182928 | 233182 | 181721.7 |
| EBOV GPΔmuc-WL$^2$P$^4$ I3-01v9b SApNP | 703967 | 60730 | 178784 | 354031 | 97482 | 91294 | 82827 | 610293 | 182253.3 |

**Week 17**

| Antigen | EC$_{50}$ titers (week 17) | | | | | | | | Geometric Mean |
|---|---|---|---|---|---|---|---|---|---|
| | M1 | M2 | M3 | M4 | M5 | M6 | M7 | M8 | |
| EBOV GPΔmuc-WL$^2$P$^4$-FOLD | 59703 | 59841 | 42414 | 92526 | 80033 | 44111 | 36894 | 36448 | 53444.3 |
| EBOV GPΔmuc-WL$^2$P$^4$ E2p SApNP | 78072 | 31500 | 66659 | 79702 | 33830 | 63116 | 75359 | 105584 | 62128.2 |
| EBOV GPΔmuc-WL$^2$P$^4$ I3-01v9b SApNP | 40965 | 18206 | 43559 | 42711 | 22529 | 65820 | 41237 | 74760 | 39837.1 |

**Statistical analysis**

| One-way ANOVA with Tukey's multiple comparisons test (w2) | Statistics | Adjusted P Value |
|---|---|---|
| EBOV GPΔmuc-WL$^2$P$^4$ trimer vs. EBOV GPΔmuc-WL$^2$P$^4$ E2p SApNP | ** | 0.0067 |
| EBOV GPΔmuc-WL$^2$P$^4$ trimer vs. EBOV GPΔmuc-WL$^2$P$^4$ I3-01v9b SApNP | ns | 0.9871 |
| EBOV GPΔmuc-WL$^2$P$^4$ E2p SApNP vs. EBOV GPΔmuc-WL$^2$P$^4$ I3-01v9b SApNP | ** | 0.0047 |

| One-way ANOVA with Tukey's multiple comparisons test (w5) | Statistics | Adjusted P Value |
|---|---|---|
| EBOV GPΔmuc-WL$^2$P$^4$ trimer vs. EBOV GPΔmuc-WL$^2$P$^4$ E2p SApNP | **** | <0.0001 |
| EBOV GPΔmuc-WL$^2$P$^4$ trimer vs. EBOV GPΔmuc-WL$^2$P$^4$ I3-01v9b SApNP | ns | 0.1769 |
| EBOV GPΔmuc-WL$^2$P$^4$ E2p SApNP vs. EBOV GPΔmuc-WL$^2$P$^4$ I3-01v9b SApNP | ** | 0.0034 |

| One-way ANOVA with Tukey's multiple comparisons test (w11) | Statistics | Adjusted P Value |
|---|---|---|
| EBOV GPΔmuc-WL$^2$P$^4$ trimer vs. EBOV GPΔmuc-WL$^2$P$^4$ E2p SApNP | ns | 0.5058 |
| EBOV GPΔmuc-WL$^2$P$^4$ trimer vs. EBOV GPΔmuc-WL$^2$P$^4$ I3-01v9b SApNP | ns | 0.2942 |
| EBOV GPΔmuc-WL$^2$P$^4$ E2p SApNP vs. EBOV GPΔmuc-WL$^2$P$^4$ I3-01v9b SApNP | ns | 0.9135 |

| One-way ANOVA with Tukey's multiple comparisons test (w17) | Statistics | Adjusted P Value |
|---|---|---|
| EBOV GPΔmuc-WL$^2$P$^4$ trimer vs. EBOV GPΔmuc-WL$^2$P$^4$ E2p SApNP | ns | 0.6171 |
| EBOV GPΔmuc-WL$^2$P$^4$ trimer vs. EBOV GPΔmuc-WL$^2$P$^4$ I3-01v9b SApNP | ns | 0.4761 |
| EBOV GPΔmuc-WL$^2$P$^4$ E2p SApNP vs. EBOV GPΔmuc-WL$^2$P$^4$ I3-01v9b SApNP | ns | 0.1078 |

#### C Week-7 sera of individual mice immunized with EBOV vaccines binding to SUDV GPΔmuc-WL²P⁴(1TD0) trimer

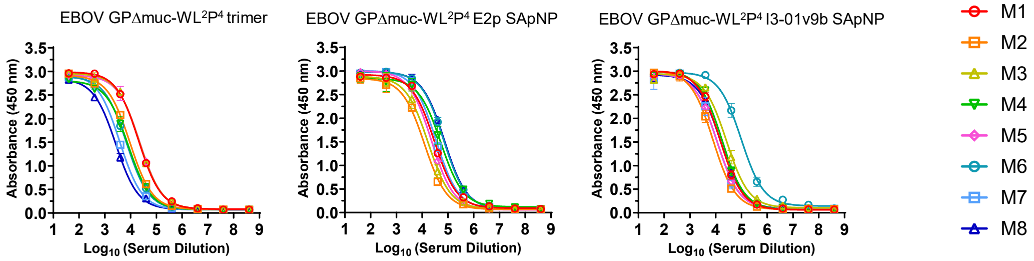


##### Mouse serum ELISA $EC_{50}$ titers

**Week 17**

| Antigen | M1 | M2 | M3 | M4 | M5 | M6 | M7 | M8 | Geometric Mean |
|---|---|---|---|---|---|---|---|---|---|
| | $EC_{50}$ titers (week 17) | | | | | | | | |
| EBOV GPΔmuc-WL²P⁴ trimer | 20545 | 9375 | 21603 | 8039 | 22891 | 7014 | 3805 | 2676 | 9273.3 |
| EBOV GPΔmuc-WL²P⁴ E2p SApNP | 29973 | 12206 | 17018 | 52874 | 22735 | 68558 | 33997 | 72867 | 32585.3 |
| EBOV GPΔmuc-WL²P⁴ I3-01v9b SApNP | 15023 | 7900 | 26186 | 17813 | 11241 | 97786 | 11184 | 15787 | 17943.0 |

##### Statistical analysis

| One-way ANOVA with Tukey's multiple comparisons test (w17) | Statistics | Adjusted P Value |
|---|---|---|
| EBOV GPΔmuc-WL²P⁴ trimer vs. EBOV GPΔmuc-WL²P⁴ E2p SApNP | ns | 0.0641 |
| EBOV GPΔmuc-WL²P⁴ trimer vs. EBOV GPΔmuc-WL²P⁴ I3-01v9b SApNP | ns | 0.4678 |
| EBOV GPΔmuc-WL²P⁴ E2p SApNP vs. EBOV GPΔmuc-WL²P⁴ I3-01v9b SApNP | ns | 0.4656 |

#### d Week-17 sera of individual mice immunized with EBOV vaccines binding to RAVV GPΔmuc-P²CT(1TD0) trimer

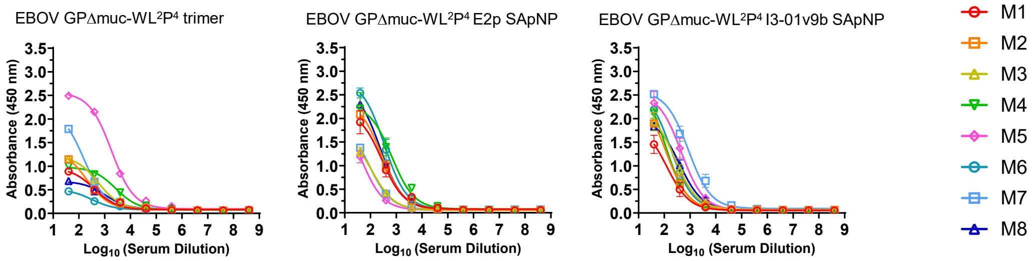


##### Mouse serum ELISA $EC_{50}$ titers

**Week 17**

| Antigen | M1 | M2 | M3 | M4 | M5 | M6 | M7 | M8 | Geometric Mean |
|---|---|---|---|---|---|---|---|---|---|
| | Absorbance (450 nm) (week 17) | | | | | | | | |
| EBOV GPΔmuc-WL²P⁴ trimer | 0.9 | 1.1 | 1.2 | 1.0 | 2.5 | 0.5 | 1.8 | 0.7 | 1.1 |
| EBOV GPΔmuc-WL²P⁴ E2p SApNP | 1.9 | 2.1 | 1.3 | 2.2 | 1.2 | 2.5 | 1.4 | 2.3 | 1.8 |
| EBOV GPΔmuc-WL²P⁴ I3-01v9b SApNP | 1.5 | 1.9 | 1.9 | 2.1 | 2.3 | 2.2 | 2.5 | 1.8 | 2.0 |

##### Statistical analysis

| One-way ANOVA with Tukey's multiple comparisons test (w17) | Statistics | Adjusted P Value |
|---|---|---|
| EBOV GPΔmuc-WL²P⁴ trimer vs. EBOV GPΔmuc-WL²P⁴ E2p SApNP | * | 0.0449 |
| EBOV GPΔmuc-WL²P⁴ trimer vs. EBOV GPΔmuc-WL²P⁴ I3-01v9b SApNP | * | 0.0111 |
| EBOV GPΔmuc-WL²P⁴ E2p SApNP vs. EBOV GPΔmuc-WL²P⁴ I3-01v9b SApNP | ns | 0.7982 |

Fig. S10

**e** **Sera of individual mice immunized with EBOV vaccines neutralizing an EBOV Makona strain**

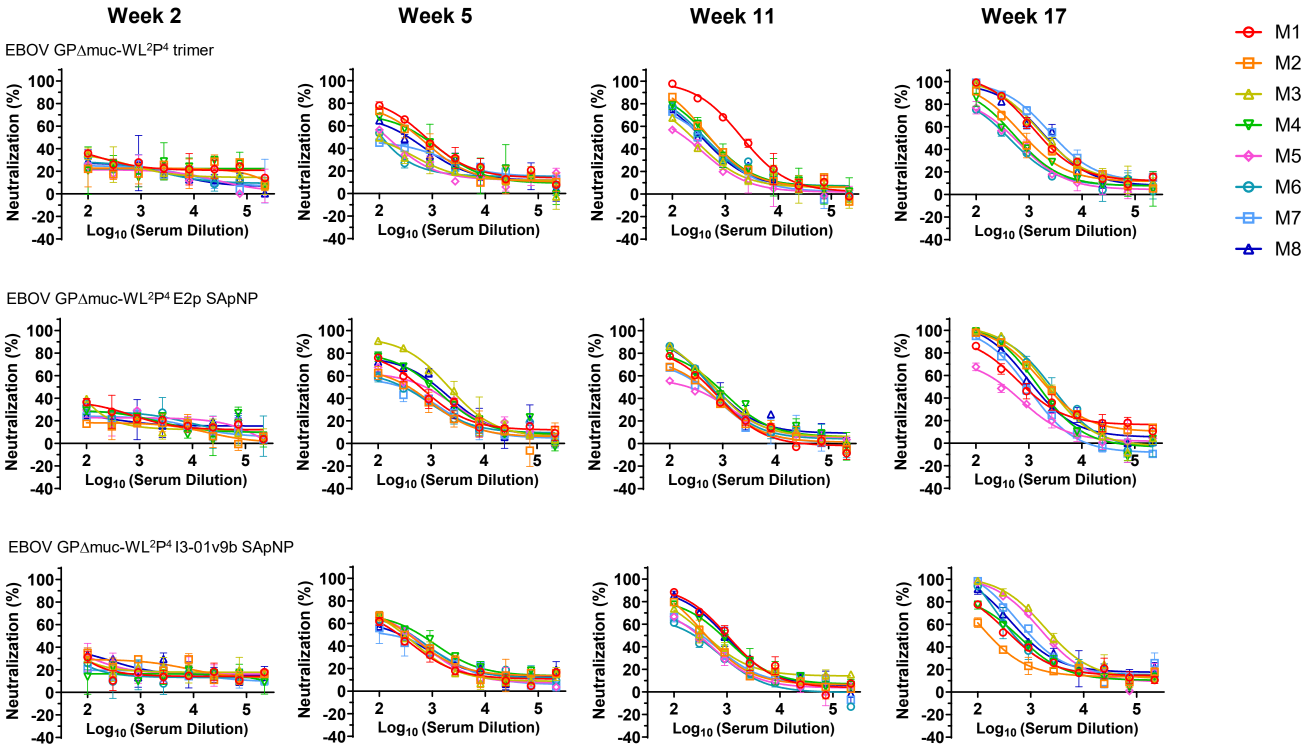


**f** **Mouse serum neutralizing $ID_{50}$ titers**

**Week 2**

| Antigen | $ID_{50}$ titers (week 2) | | | | | | | | Geometric Mean |
|---|---|---|---|---|---|---|---|---|---|
| | M1 | M2 | M3 | M4 | M5 | M6 | M7 | M8 | |
| EBOV GPΔmuc-WL$^2$P$^4$ trimer | <100 | <100 | <100 | <100 | <100 | <100 | <100 | <100 | N/A |
| EBOV GPΔmuc-WL$^2$P$^4$ E2p SApNP | <100 | <100 | <100 | <100 | <100 | <100 | <100 | <100 | N/A |
| EBOV GPΔmuc-WL$^2$P$^4$ I3-01v9b SApNP | <100 | <100 | <100 | <100 | <100 | <100 | <100 | <100 | N/A |

**Week 5**

| Antigen | $ID_{50}$ titers (week 5) | | | | | | | | Geometric Mean |
|---|---|---|---|---|---|---|---|---|---|
| | M1 | M2 | M3 | M4 | M5 | M6 | M7 | M8 | |
| EBOV GPΔmuc-WL$^2$P$^4$ trimer | 805.4 | 559.3 | 149.3 | 588.2 | 174.1 | 145.1 | 203.2 | 360.5 | 304.1 |
| EBOV GPΔmuc-WL$^2$P$^4$ E2p SApNP | 636.9 | 358.8 | 2178.0 | 1002.0 | 555.2 | 322.9 | 299.4 | 1167.0 | 648.4 |
| EBOV GPΔmuc-WL$^2$P$^4$ I3-01v9b SApNP | 267.9 | 390.2 | 313.1 | 575.8 | 378.9 | 413.4 | 258.7 | 296.7 | 350.3 |

**Week 11**

| Antigen | $ID_{50}$ titers (week 11) | | | | | | | | Geometric Mean |
|---|---|---|---|---|---|---|---|---|---|
| | M1 | M2 | M3 | M4 | M5 | M6 | M7 | M8 | |
| EBOV GPΔmuc-WL$^2$P$^4$ trimer | 2117.0 | 560.6 | 254.4 | 521.5 | 188.7 | 430.2 | 369.3 | 377.3 | 453.2 |
| EBOV GPΔmuc-WL$^2$P$^4$ E2p SApNP | 494.5 | 392.0 | 734.2 | 747.4 | 275.3 | 643.8 | 365.2 | 759.9 | 518.6 |
| EBOV GPΔmuc-WL$^2$P$^4$ I3-01v9b SApNP | 1006.0 | 434.9 | 454.1 | 766.3 | 289.4 | 237.7 | 327.0 | 928.5 | 487.3 |

**Week 17**

| Antigen | $ID_{50}$ titers (week 17) | | | | | | | | Geometric Mean |
|---|---|---|---|---|---|---|---|---|---|
| | M1 | M2 | M3 | M4 | M5 | M6 | M7 | M8 | |
| EBOV GPΔmuc-WL$^2$P$^4$ trimer | 2106.0 | 1038.0 | 2482.0 | 641.1 | 505.0 | 437.0 | 3537.0 | 2352.0 | 1260.8 |
| EBOV GPΔmuc-WL$^2$P$^4$ E2p SApNP | 1000.0 | 2501.0 | 2393.0 | 1580.0 | 329.2 | 2587.0 | 875.5 | 1415.0 | 1333.1 |
| EBOV GPΔmuc-WL$^2$P$^4$ I3-01v9b SApNP | 513.1 | 205.2 | 2961.0 | 592.6 | 2154.0 | 693.4 | 1272.0 | 994.0 | 876.7 |

#### Statistical analysis

| One-way ANOVA with Tukey's multiple comparisons test (w5) | Statistics | Adjusted P Value |
|---|---|---|
| EBOV GPΔmuc-WL$^2$P$^4$ trimer vs. EBOV GPΔmuc-WL$^2$P$^4$ E2p SApNP | ns | 0.0919 |
| EBOV GPΔmuc-WL$^2$P$^4$ trimer vs. EBOV GPΔmuc-WL$^2$P$^4$ I3-01v9b SApNP | ns | 0.9982 |
| EBOV GPΔmuc-WL$^2$P$^4$ E2p SApNP vs. EBOV GPΔmuc-WL$^2$P$^4$ I3-01v9b SApNP | ns | 0.0824 |

| One-way ANOVA with Tukey's multiple comparisons test (w11) | Statistics | Adjusted P Value |
|---|---|---|
| EBOV GPΔmuc-WL$^2$P$^4$ trimer vs. EBOV GPΔmuc-WL$^2$P$^4$ E2p SApNP | ns | 0.9676 |
| EBOV GPΔmuc-WL$^2$P$^4$ trimer vs. EBOV GPΔmuc-WL$^2$P$^4$ I3-01v9b SApNP | ns | 0.9724 |
| EBOV GPΔmuc-WL$^2$P$^4$ E2p SApNP vs. EBOV GPΔmuc-WL$^2$P$^4$ I3-01v9b SApNP | ns | 0.9998 |

| One-way ANOVA with Tukey's multiple comparisons test (w17) | Statistics | Adjusted P Value |
|---|---|---|
| EBOV GPΔmuc-WL$^2$P$^4$ trimer vs. EBOV GPΔmuc-WL$^2$P$^4$ E2p SApNP | ns | 0.9938 |
| EBOV GPΔmuc-WL$^2$P$^4$ trimer vs. EBOV GPΔmuc-WL$^2$P$^4$ I3-01v9b SApNP | ns | 0.618 |
| EBOV GPΔmuc-WL$^2$P$^4$ E2p SApNP vs. EBOV GPΔmuc-WL$^2$P$^4$ I3-01v9b SApNP | ns | 0.6831 |

### Fig. S10

#### g Week-17 sera of individual mice immunized with EBOV vaccines neutralizing a SUDV Gulu strain

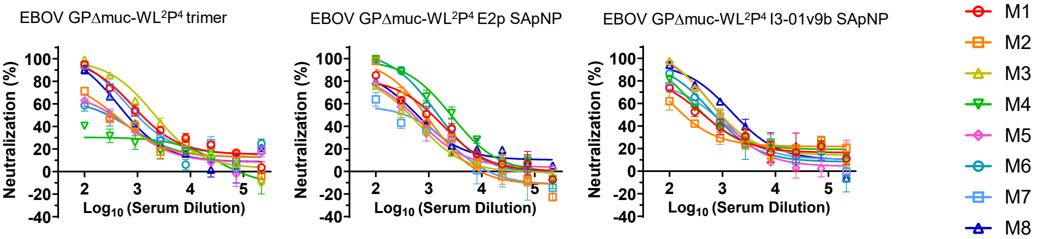


##### Mouse serum neutralizing ID$_{50}$ titers

**Week 17**

| Antigen | ID$_{50}$ titers (week 17) | | | | | | | | Geometric Mean |
|---|---|---|---|---|---|---|---|---|---|
| | M1 | M2 | M3 | M4 | M5 | M6 | M7 | M8 | |
| EBOV GPΔmuc-WL$^2$P$^4$ trimer | 1530.0 | 407.3 | 2200.0 | 113.8 | 341.2 | 323.2 | 1236.0 | 726.5 | 593.8 |
| EBOV GPΔmuc-WL$^2$P$^4$ E2p SApNP | 956.0 | 759.0 | 403.8 | 2580.0 | 489.0 | 1564.0 | 301.3 | 647.6 | 761.3 |
| EBOV GPΔmuc-WL$^2$P$^4$ I3-01v9b SApNP | 512.3 | 265.2 | 1198.0 | 608.8 | 1006.0 | 765.2 | 662.5 | 1824.0 | 742.3 |

##### Statistical analysis

| One-way ANOVA with Tukey's multiple comparisons test (w17) | Statistics | Adjusted P Value |
|---|---|---|
| EBOV GPΔmuc-WL$^2$P$^4$ trimer vs. EBOV GPΔmuc-WL$^2$P$^4$ E2p SApNP | ns | 0.9497 |
| EBOV GPΔmuc-WL$^2$P$^4$ trimer vs. EBOV GPΔmuc-WL$^2$P$^4$ I3-01v9b SApNP | ns | >0.9999 |
| EBOV GPΔmuc-WL$^2$P$^4$ E2p SApNP vs. EBOV GPΔmuc-WL$^2$P$^4$ I3-01v9b SApNP | ns | 0.9453 |

#### h Week-17 sera of individual mice immunized with EBOV vaccines neutralizing a BDBV Uganda strain

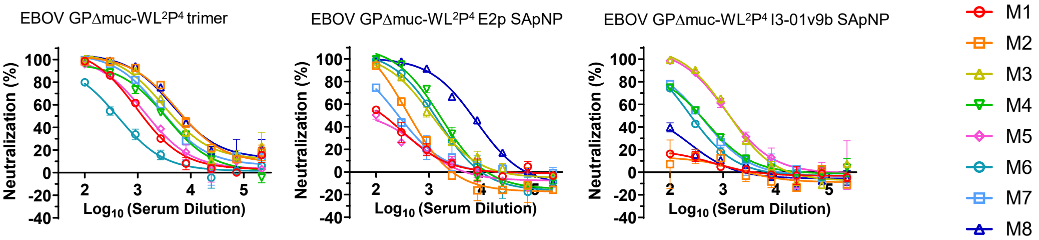


##### Mouse serum neutralizing ID$_{50}$ titers

**Week 17**

| Antigen | ID$_{50}$ titers (week 17) | | | | | | | | Geometric Mean |
|---|---|---|---|---|---|---|---|---|---|
| | M1 | M2 | M3 | M4 | M5 | M6 | M7 | M8 | |
| EBOV GPΔmuc-WL$^2$P$^4$ trimer | 1344.0 | 7809.0 | 5137.0 | 3217.0 | 1631.0 | 411.3 | 3729.0 | 7747.0 | 2759.4 |
| EBOV GPΔmuc-WL$^2$P$^4$ E2p SApNP | 149.6 | 394.3 | 1109.0 | 1525.0 | 115.1 | 1174.0 | 261.1 | 5895.0 | 616.1 |
| EBOV GPΔmuc-WL$^2$P$^4$ I3-01v9b SApNP | <100 | <100 | 1332.0 | 356.1 | 1397.0 | 240.1 | 352.0 | <100 | 561.9 |

##### Statistical analysis

| One-way ANOVA with Tukey's multiple comparisons test (w17) | Statistics | Adjusted P Value |
|---|---|---|
| EBOV GPΔmuc-WL$^2$P$^4$ trimer vs. EBOV GPΔmuc-WL$^2$P$^4$ E2p SApNP | * | 0.0469 |
| EBOV GPΔmuc-WL$^2$P$^4$ trimer vs. EBOV GPΔmuc-WL$^2$P$^4$ I3-01v9b SApNP | ** | 0.0076 |
| EBOV GPΔmuc-WL$^2$P$^4$ E2p SApNP vs. EBOV GPΔmuc-WL$^2$P$^4$ I3-01v9b SApNP | ns | 0.6882 |

#### i Sera of individual mice immunized with SUDV vaccines binding to SUDV GPΔmuc-WL²P⁴(1TD0) trimer

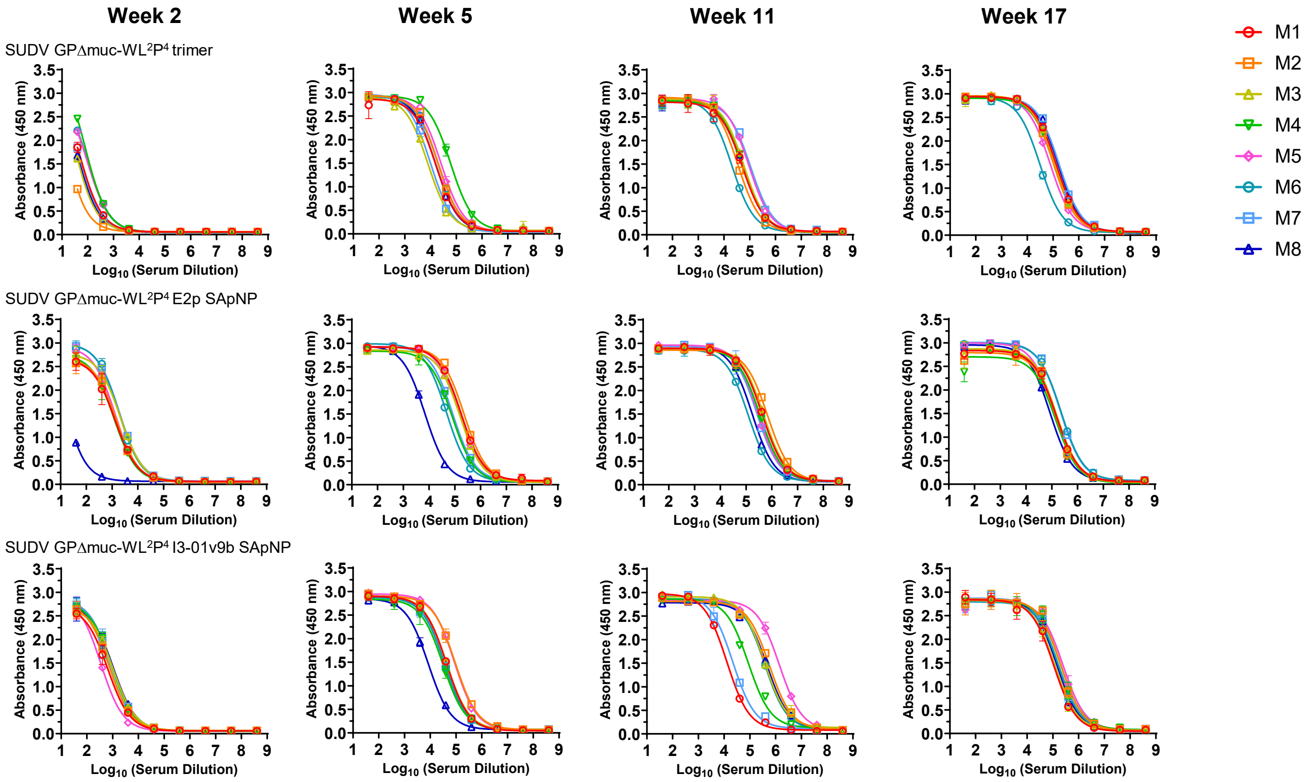


#### j Mouse serum ELISA $EC_{50}$ titers

**Week 2**

| Antigen | $EC_{50}$ titers (week 2) | | | | | | | | Geometric Mean |
|---|---|---|---|---|---|---|---|---|---|
| | M1 | M2 | M3 | M4 | M5 | M6 | M7 | M8 | |
| SUDV GPΔmuc-WL$^2$P$^4$ trimer | 69.8 | 20.2 | 50.0 | 149.7 | 116.9 | 122.4 | 68.1 | 55.0 | 69.8 |
| SUDV GPΔmuc-WL$^2$P$^4$ E2p SApNP | 1058.0 | 1223.0 | 1977.0 | 1062.0 | 1419.0 | 2123.0 | 2262.0 | 17.9 | 871.0 |
| SUDV GPΔmuc-WL$^2$P$^4$ I3-01v9b SApNP | 550.3 | 700.2 | 876.2 | 893.5 | 373.7 | 741.1 | 903.2 | 940.0 | 718.4 |

**Week 5**

| Antigen | $EC_{50}$ titers (week 5) | | | | | | | | Geometric Mean |
|---|---|---|---|---|---|---|---|---|---|
| | M1 | M2 | M3 | M4 | M5 | M6 | M7 | M8 | |
| SUDV GPΔmuc-WL$^2$P$^4$ trimer | 16436 | 19941 | 7907 | 60277 | 25656 | 19754 | 9992 | 15619 | 18259.2 |
| SUDV GPΔmuc-WL$^2$P$^4$ E2p SApNP | 177305 | 230404 | 159220 | 79353 | 157784 | 47895 | 82714 | 6921 | 82910.4 |
| SUDV GPΔmuc-WL$^2$P$^4$ I3-01v9b SApNP | 42731 | 93470 | 42715 | 31125 | 87027 | 36180 | 35115 | 8482 | 38650.3 |

**Week 11**

| Antigen | $EC_{50}$ titers (week 11) | | | | | | | | Geometric Mean |
|---|---|---|---|---|---|---|---|---|---|
| | M1 | M2 | M3 | M4 | M5 | M6 | M7 | M8 | |
| SUDV GPΔmuc-WL$^2$P$^4$ trimer | 51852 | 34819 | 59219 | 58035 | 92570 | 20182 | 108658 | 56856 | 53935.6 |
| SUDV GPΔmuc-WL$^2$P$^4$ E2p SApNP | 427238 | 688825 | 479499 | 349324 | 289258 | 121660 | 264183 | 173015 | 307186.5 |
| SUDV GPΔmuc-WL$^2$P$^4$ I3-01v9b SApNP | 12804 | 573418 | 373590 | 88061 | 1405850 | 379235 | 21591 | 471753 | 183962.3 |

**Week 17**

| Antigen | $EC_{50}$ titers (week 17) | | | | | | | | Geometric Mean |
|---|---|---|---|---|---|---|---|---|---|
| | M1 | M2 | M3 | M4 | M5 | M6 | M7 | M8 | |
| SUDV GPΔmuc-WL$^2$P$^4$ trimer | 132375 | 104284 | 117314 | 112501 | 78562 | 31328 | 174100 | 161747 | 102958.9 |
| SUDV GPΔmuc-WL$^2$P$^4$ E2p SApNP | 150503 | 150658 | 156743 | 142867 | 124345 | 229353 | 218325 | 86164 | 151148.0 |
| SUDV GPΔmuc-WL$^2$P$^4$ I3-01v9b SApNP | 108351 | 204158 | 120591 | 200895 | 248727 | 170555 | 157838 | 144221 | 163771.7 |

##### Statistical analysis

| One-way ANOVA with Tukey's multiple comparisons test (w2) | Statistics | Adjusted P Value |
|---|---|---|
| SUDV GPΔmuc-WL$^2$P$^4$ trimer vs. SUDV GPΔmuc-WL$^2$P$^4$ E2p SApNP | **** | <0.0001 |
| SUDV GPΔmuc-WL$^2$P$^4$ trimer vs. SUDV GPΔmuc-WL$^2$P$^4$ I3-01v9b SApNP | * | 0.0169 |
| SUDV GPΔmuc-WL$^2$P$^4$ E2p SApNP vs. SUDV GPΔmuc-WL$^2$P$^4$ I3-01v9b SApNP | * | 0.0207 |

| One-way ANOVA with Tukey's multiple comparisons test (w5) | Statistics | Adjusted P Value |
|---|---|---|
| SUDV GPΔmuc-WL$^2$P$^4$ trimer vs. SUDV GPΔmuc-WL$^2$P$^4$ E2p SApNP | ** | 0.0016 |
| SUDV GPΔmuc-WL$^2$P$^4$ trimer vs. SUDV GPΔmuc-WL$^2$P$^4$ I3-01v9b SApNP | ns | 0.5472 |
| SUDV GPΔmuc-WL$^2$P$^4$ E2p SApNP vs. SUDV GPΔmuc-WL$^2$P$^4$ I3-01v9b SApNP | * | 0.0187 |

| One-way ANOVA with Tukey's multiple comparisons test (w11) | Statistics | Adjusted P Value |
|---|---|---|
| SUDV GPΔmuc-WL$^2$P$^4$ trimer vs. SUDV GPΔmuc-WL$^2$P$^4$ E2p SApNP | ns | 0.1259 |
| SUDV GPΔmuc-WL$^2$P$^4$ trimer vs. SUDV GPΔmuc-WL$^2$P$^4$ I3-01v9b SApNP | ns | 0.0502 |
| SUDV GPΔmuc-WL$^2$P$^4$ E2p SApNP vs. SUDV GPΔmuc-WL$^2$P$^4$ I3-01v9b SApNP | ns | 0.885 |

| One-way ANOVA with Tukey's multiple comparisons test (w17) | Statistics | Adjusted P Value |
|---|---|---|
| SUDV GPΔmuc-WL$^2$P$^4$ trimer vs. SUDV GPΔmuc-WL$^2$P$^4$ E2p SApNP | ns | 0.1724 |
| SUDV GPΔmuc-WL$^2$P$^4$ trimer vs. SUDV GPΔmuc-WL$^2$P$^4$ I3-01v9b SApNP | ns | 0.0653 |
| SUDV GPΔmuc-WL$^2$P$^4$ E2p SApNP vs. SUDV GPΔmuc-WL$^2$P$^4$ I3-01v9b SApNP | ns | 0.8627 |

#### k Week-17 sera of individual mice immunized with SUDV vaccines binding to EBOV GPΔmuc-WL$^2$P$^4$(1TD0) trimer

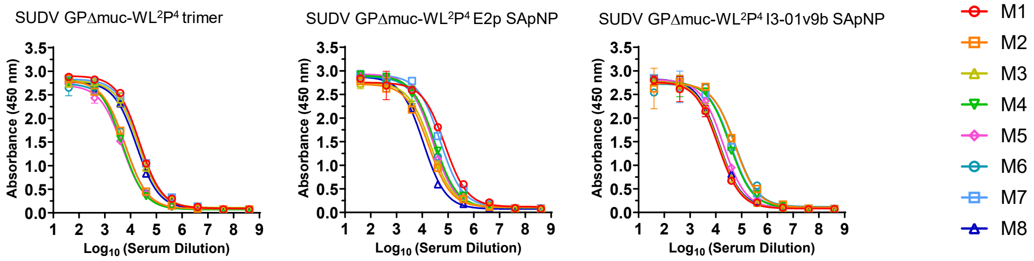


##### Mouse serum ELISA EC$_{50}$ titers

**Week 17**

| Antigen | EC$_{50}$ titers (week 17) | | | | | | | | Geometric Mean |
|---|---|---|---|---|---|---|---|---|---|
| | M1 | M2 | M3 | M4 | M5 | M6 | M7 | M8 | |
| SUDV GPΔmuc-WL$^2$P$^4$ trimer | 22131 | 6105 | 19939 | 4814 | 4809 | 6529 | 22832 | 16822 | 10574.9 |
| SUDV GPΔmuc-WL$^2$P$^4$ E2p SApNP | 74857 | 17696 | 20913 | 30182 | 25142 | 26898 | 49408 | 11012 | 27290.2 |
| SUDV GPΔmuc-WL$^2$P$^4$ I3-01v9b SApNP | 12210 | 55319 | 13273 | 35138 | 19190 | 57302 | 38054 | 13336 | 25446.4 |

##### Statistical analysis

| One-way ANOVA with Tukey's multiple comparisons test (w17) | Statistics | Adjusted P Value |
|---|---|---|
| SUDV GPΔmuc-WL$^2$P$^4$ trimer vs. SUDV GPΔmuc-WL$^2$P$^4$ E2p SApNP | ns | 0.0832 |
| SUDV GPΔmuc-WL$^2$P$^4$ trimer vs. SUDV GPΔmuc-WL$^2$P$^4$ I3-01v9b SApNP | ns | 0.1177 |
| SUDV GPΔmuc-WL$^2$P$^4$ E2p SApNP vs. SUDV GPΔmuc-WL$^2$P$^4$ I3-01v9b SApNP | ns | 0.9817 |

#### l Week-17 sera of individual mice immunized with SUDV vaccines binding to RAVV GPΔmuc-P$^2$CT(1TD0) trimer

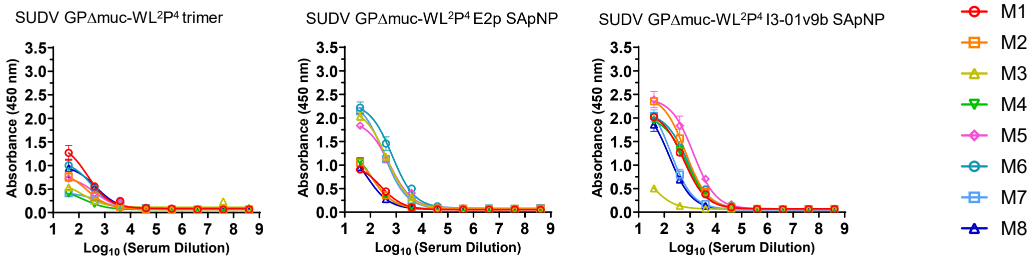


##### Mouse serum ELISA EC$_{50}$ titers

**Week 17**

| Antigen | Absorbance (450 nm) (week 17) | | | | | | | | Geometric Mean |
|---|---|---|---|---|---|---|---|---|---|
| | M1 | M2 | M3 | M4 | M5 | M6 | M7 | M8 | |
| SUDV GPΔmuc-WL$^2$P$^4$ trimer | 1.3 | 0.8 | 0.5 | 0.4 | 0.7 | 1.0 | 0.4 | 0.9 | 0.7 |
| SUDV GPΔmuc-WL$^2$P$^4$ E2p SApNP | 0.9 | 1.1 | 2.0 | 1.1 | 1.8 | 2.2 | 2.1 | 1.0 | 1.4 |
| SUDV GPΔmuc-WL$^2$P$^4$ I3-01v9b SApNP | 2.0 | 2.4 | 0.5 | 1.9 | 2.4 | 2.0 | 2.0 | 1.9 | 1.7 |

##### Statistical analysis

| One-way ANOVA with Tukey's multiple comparisons test (w17) | Statistics | Adjusted P Value |
|---|---|---|
| SUDV GPΔmuc-WL$^2$P$^4$ trimer vs. SUDV GPΔmuc-WL$^2$P$^4$ E2p SApNP | * | 0.0149 |
| SUDV GPΔmuc-WL$^2$P$^4$ trimer vs. SUDV GPΔmuc-WL$^2$P$^4$ I3-01v9b SApNP | *** | 0.0005 |
| SUDV GPΔmuc-WL$^2$P$^4$ E2p SApNP vs. SUDV GPΔmuc-WL$^2$P$^4$ I3-01v9b SApNP | ns | 0.338 |

**Fig. S10**

**m** **Sera of individual mice immunized with SUDV vaccines neutralizing a SUDV Gulu strain**

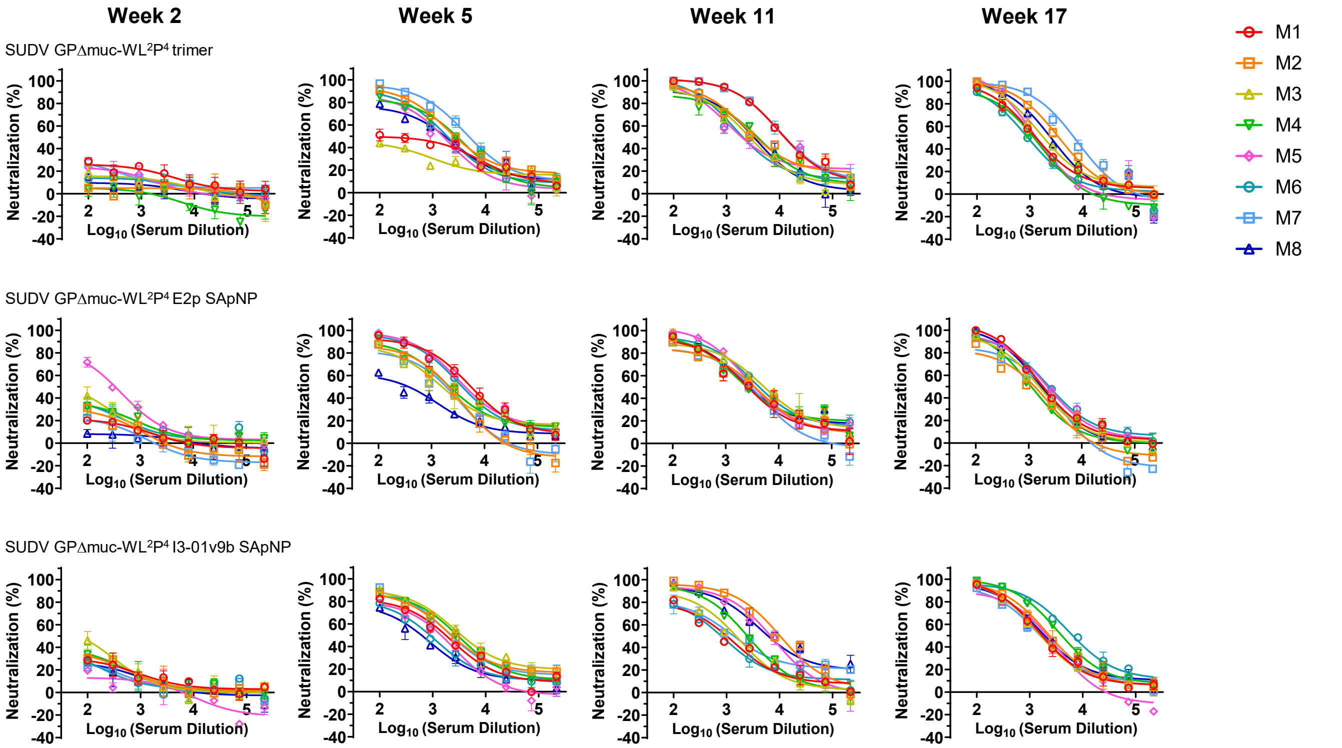


**n** **Mouse serum neutralizing $ID_{50}$ titers**

**Week 2**

| Antigen | $ID_{50}$ titers (week 2) M1 | M2 | M3 | M4 | M5 | M6 | M7 | M8 | Geometric Mean |
|---|---|---|---|---|---|---|---|---|---|
| SUDV GPΔmuc-WL$^2$P$^4$ trimer | <100 | <100 | <100 | <100 | <100 | <100 | <100 | <100 | N/A |
| SUDV GPΔmuc-WL$^2$P$^4$ E2p SApNP | <100 | <100 | <100 | <100 | 331.1 | <100 | <100 | <100 | N/A |
| SUDV GPΔmuc-WL$^2$P$^4$ I3-01v9b SApNP | <100 | <100 | 107.2 | <100 | <100 | <100 | <100 | <100 | N/A |

**Week 5**

| Antigen | $ID_{50}$ titers (week 5) M1 | M2 | M3 | M4 | M5 | M6 | M7 | M8 | Geometric Mean |
|---|---|---|---|---|---|---|---|---|---|
| SUDV GPΔmuc-WL$^2$P$^4$ trimer | 420 | 3076 | 156.6 | 2417 | 1377 | 1896 | 5149 | 1477 | 1328.6 |
| SUDV GPΔmuc-WL$^2$P$^4$ E2p SApNP | 5677 | 1634 | 1754 | 2420 | 5081 | 4212 | 1332 | 355.5 | 2114.1 |
| SUDV GPΔmuc-WL$^2$P$^4$ I3-01v9b SApNP | 1582 | 2485 | 4102 | 2862 | 1119 | 843.6 | 2506 | 584.4 | 1681.2 |

**Week 11**

| Antigen | $ID_{50}$ titers (week 11) M1 | M2 | M3 | M4 | M5 | M6 | M7 | M8 | Geometric Mean |
|---|---|---|---|---|---|---|---|---|---|
| SUDV GPΔmuc-WL$^2$P$^4$ trimer | 13810 | 4694 | 2805 | 3670 | 2858 | 2497 | 14410 | 3069 | 4628.4 |
| SUDV GPΔmuc-WL$^2$P$^4$ E2p SApNP | 2835 | 3130 | 5528 | 3311 | 3940 | 4935 | 2323 | 3352 | 3538.7 |
| SUDV GPΔmuc-WL$^2$P$^4$ I3-01v9b SApNP | 889.3 | 10154 | 1391 | 2424 | 6257 | 808.9 | 1524 | 7608 | 2549.8 |

**Week 17**

| Antigen | $ID_{50}$ titers (week 17) M1 | M2 | M3 | M4 | M5 | M6 | M7 | M8 | Geometric Mean |
|---|---|---|---|---|---|---|---|---|---|
| SUDV GPΔmuc-WL$^2$P$^4$ trimer | 1389 | 3550 | 2017 | 1006 | 1354 | 931.1 | 6329 | 2409 | 1929.8 |
| SUDV GPΔmuc-WL$^2$P$^4$ E2p SApNP | 2035 | 928.9 | 1546 | 1147 | 2291 | 2536 | 1215 | 1753 | 1593.1 |
| SUDV GPΔmuc-WL$^2$P$^4$ I3-01v9b SApNP | 1853 | 2507 | 1746 | 4395 | 1648 | 7166 | 1695 | 2160 | 2503.1 |

**Statistical analysis**

| One-way ANOVA with Tukey's multiple comparisons test (w5) | Statistics | Adjusted P Value |
|---|---|---|
| SUDV GPΔmuc-WL$^2$P$^4$ trimer vs. SUDV GPΔmuc-WL$^2$P$^4$ E2p SApNP | ns | 0.5769 |
| SUDV GPΔmuc-WL$^2$P$^4$ trimer vs. SUDV GPΔmuc-WL$^2$P$^4$ I3-01v9b SApNP | ns | 0.9998 |
| SUDV GPΔmuc-WL$^2$P$^4$ E2p SApNP vs. SUDV GPΔmuc-WL$^2$P$^4$ I3-01v9b SApNP | ns | 0.5879 |

| One-way ANOVA with Tukey's multiple comparisons test (w11) | Statistics | Adjusted P Value |
|---|---|---|
| SUDV GPΔmuc-WL$^2$P$^4$ trimer vs. SUDV GPΔmuc-WL$^2$P$^4$ E2p SApNP | ns | 0.4294 |
| SUDV GPΔmuc-WL$^2$P$^4$ trimer vs. SUDV GPΔmuc-WL$^2$P$^4$ I3-01v9b SApNP | ns | 0.4956 |
| SUDV GPΔmuc-WL$^2$P$^4$ E2p SApNP vs. SUDV GPΔmuc-WL$^2$P$^4$ I3-01v9b SApNP | ns | 0.9925 |

| One-way ANOVA with Tukey's multiple comparisons test (w17) | Statistics | Adjusted P Value |
|---|---|---|
| SUDV GPΔmuc-WL$^2$P$^4$ trimer vs. SUDV GPΔmuc-WL$^2$P$^4$ E2p SApNP | ns | 0.6592 |
| SUDV GPΔmuc-WL$^2$P$^4$ trimer vs. SUDV GPΔmuc-WL$^2$P$^4$ I3-01v9b SApNP | ns | 0.7862 |
| SUDV GPΔmuc-WL$^2$P$^4$ E2p SApNP vs. SUDV GPΔmuc-WL$^2$P$^4$ I3-01v9b SApNP | ns | 0.2916 |

Fig. S10

#### o Week-17 sera of individual mice immunized with SUDV vaccines neutralizing an EBOV Makona strain

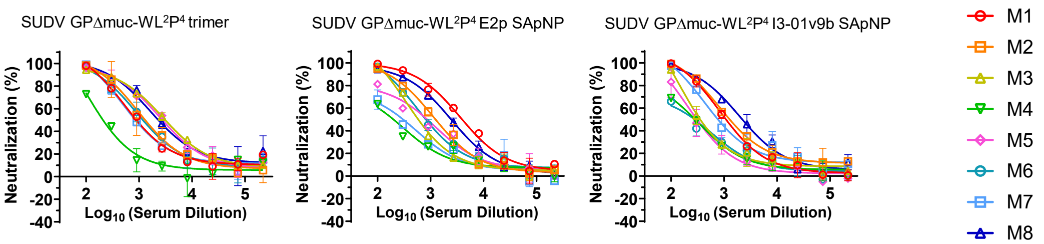


##### Mouse serum neutralizing ID$_{50}$ titers

**Week 17**

| Antigen | ID$_{50}$ titers (week 17) | | | | | | | | Geometric Mean |
|---|---|---|---|---|---|---|---|---|---|
| | M1 | M2 | M3 | M4 | M5 | M6 | M7 | M8 | |
| SUDV GPΔmuc-WL$^2$P$^4$ trimer | 1073 | 1510 | 3312 | 230.5 | 2932 | 1345 | 1075 | 2621 | 1387.6 |
| SUDV GPΔmuc-WL$^2$P$^4$ E2p SApNP | 4144 | 1304 | 687.5 | 217 | 758.8 | 964.7 | 297 | 2453 | 899.9 |
| SUDV GPΔmuc-WL$^2$P$^4$ I3-01v9b SApNP | 1214 | 1567 | 467.4 | 296.9 | 336.4 | 271 | 990.7 | 2370 | 698.3 |

##### Statistical analysis

| One-way ANOVA with Tukey's multiple comparisons test (w17) | Statistics | Adjusted P Value |
|---|---|---|
| SUDV GPΔmuc-WL$^2$P$^4$ trimer vs. SUDV GPΔmuc-WL$^2$P$^4$ E2p SApNP | ns | 0.7312 |
| SUDV GPΔmuc-WL$^2$P$^4$ trimer vs. SUDV GPΔmuc-WL$^2$P$^4$ I3-01v9b SApNP | ns | 0.2983 |
| SUDV GPΔmuc-WL$^2$P$^4$ E2p SApNP vs. SUDV GPΔmuc-WL$^2$P$^4$ I3-01v9b SApNP | ns | 0.7257 |

#### p Week-17 sera of individual mice immunized with SUDV vaccines neutralizing a BDBV Uganda strain

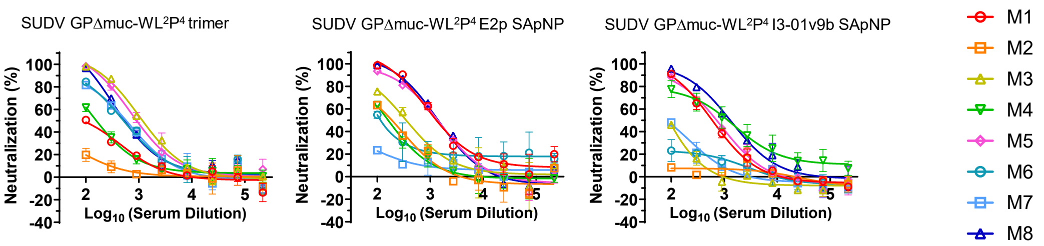


##### Mouse serum neutralizing ID$_{50}$ titers

**Week 17**

| Antigen | ID$_{50}$ titers (week 17) | | | | | | | | Geometric Mean |
|---|---|---|---|---|---|---|---|---|---|
| | M1 | M2 | M3 | M4 | M5 | M6 | M7 | M8 | |
| SUDV GPΔmuc-WL$^2$P$^4$ trimer | 126.7 | 26.1 | 1237.0 | 165.3 | 985.6 | 518.9 | 543.9 | 613.3 | 322.0 |
| SUDV GPΔmuc-WL$^2$P$^4$ E2p SApNP | 1504.0 | 174.5 | 356.0 | 156.5 | 1298.0 | 151.8 | 35.5 | 1393.0 | 330.6 |
| SUDV GPΔmuc-WL$^2$P$^4$ I3-01v9b SApNP | 580.4 | 9.7 | 69.0 | 1085.0 | 656.4 | <100 | <100 | 1280.0 | 185.5 |

##### Statistical analysis

| One-way ANOVA with Tukey's multiple comparisons test (w17) | Statistics | Adjusted P Value |
|---|---|---|
| SUDV GPΔmuc-WL$^2$P$^4$ trimer vs. SUDV GPΔmuc-WL$^2$P$^4$ E2p SApNP | ns | 0.913 |
| SUDV GPΔmuc-WL$^2$P$^4$ trimer vs. SUDV GPΔmuc-WL$^2$P$^4$ I3-01v9b SApNP | ns | 0.9893 |
| SUDV GPΔmuc-WL$^2$P$^4$ E2p SApNP vs. SUDV GPΔmuc-WL$^2$P$^4$ I3-01v9b SApNP | ns | 0.8492 |

### Fig. S10

**q  Sera of individual mice immunized with RAVV vaccines binding to RAVV GPΔmuc-P²CT(1TD0) trimer**

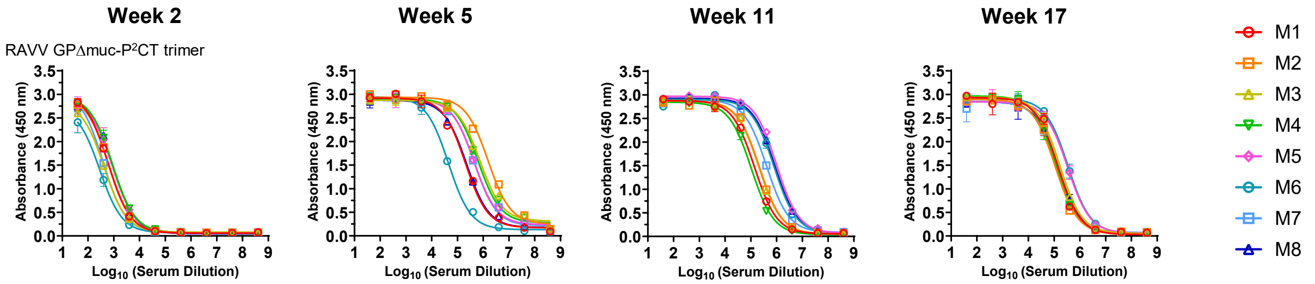


**Mouse serum ELISA $EC_{50}$ titers**

| Antigen | $EC_{50}$ titers | | | | | | | | Geometric Mean |
|---|---|---|---|---|---|---|---|---|---|
| | M1 | M2 | M3 | M4 | M5 | M6 | M7 | M8 | |
| RAVV GPΔmuc-P²CT trimer-w2 | 718.9 | 721.9 | 432.3 | 1030.0 | 946.3 | 262.3 | 481.5 | 958.5 | 635.1 |
| RAVV GPΔmuc-P²CT trimer-w5 | 202735 | 1544554 | 746406 | 628189 | 447117 | 44771 | 446178 | 225949 | 362215.2 |
| RAVV GPΔmuc-P²CT trimer-w11 | 139135 | 218951 | 208199 | 104125 | 998580 | 786127 | 408247 | 853062 | 340466.6 |
| RAVV GPΔmuc-P²CT trimer-w17 | 140811 | 110900 | 164905 | 119990 | 336478 | 344895 | 127509 | 141835 | 168457.1 |

**r  Week-17 sera of individual mice immunized with RAVV vaccines binding to EBOV GPΔmuc-WL²P⁴(1TD0) trimer**

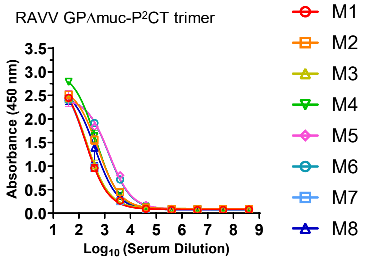


**Mouse serum ELISA $EC_{50}$ titers**

| Antigen | $EC_{50}$ titers (week 17) | | | | | | | | Geometric Mean |
|---|---|---|---|---|---|---|---|---|---|
| | M1 | M2 | M3 | M4 | M5 | M6 | M7 | M8 | |
| RAVV GPΔmuc-P²CT trimer | 177.5 | 537.5 | 197.8 | 459.9 | 1477.0 | 1430.0 | 182.7 | 422.8 | 440.4 |

**s  Week-17 sera of individual mice immunized with RAVV vaccines binding to SUDV GPΔmuc-WL²P⁴(1TD0) trimer**

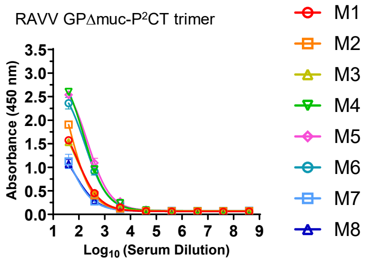


**Mouse serum ELISA $EC_{50}$ titers**

| Antigen | Absorbance (450 nm) (week 17) | | | | | | | | Geometric Mean |
|---|---|---|---|---|---|---|---|---|---|
| | M1 | M2 | M3 | M4 | M5 | M6 | M7 | M8 | |
| RAVV GPΔmuc-P²CT trimer | 1.6 | 1.9 | 1.5 | 2.6 | 2.5 | 2.4 | 2.4 | 2.4 | 2.1 |

**t  Week-17 IgG samples of individual mice immunized with RAVV vaccines neutralizing a MARV Angola strain**

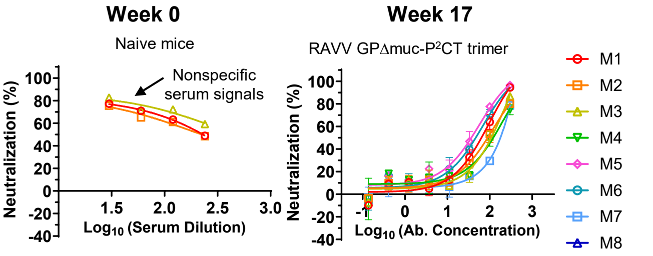


**Week 17 mouse IgG neutralization**

| Virus strains | $IC_{50}$ (μg/mL) (week 17) | | | | | | | | Geometric Mean |
|---|---|---|---|---|---|---|---|---|---|
| | M1 | M2 | M3 | M4 | M5 | M6 | M7 | M8 | |
| MARV-Angola | 56.2 | 81.4 | 93.3 | 104.4 | 30.9 | 40.3 | 148.4 | 71.8 | 70.2 |

**u  Week-17 sera of individual mice immunized with RAVV vaccines neutralizing three orthoebolavirus strains**

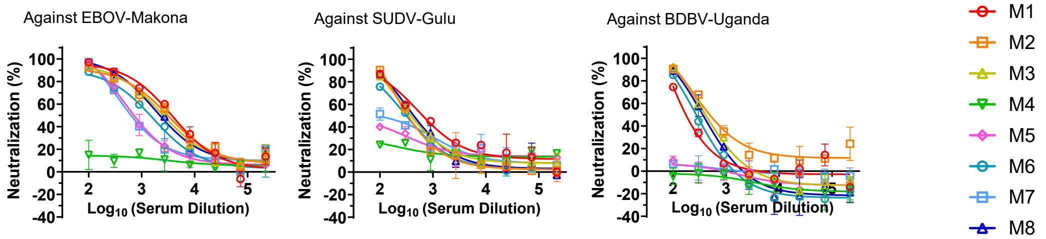

**Fig. S10. Immunogenicity of EBOV, SUDV, and RAVV GPΔmuc vaccines in mice.** (**a**) ELISA curves of mouse sera from EBOV GPΔmuc trimer and SApNP vaccine groups (n = 8 mice/group) binding to the coating antigen EBOV GPΔmuc-WL$^2$P$^4$(1TD0) trimer (**b**) (Top) Summary of geometric mean $EC_{50}$ titers measured for Ebola GPΔmuc vaccine groups against EBOV GPΔmuc-WL$^2$P$^4$(1TD0) trimer. Color coding indicates $EC_{50}$ levels (green to red: low to high binding). (Bottom) Summary of statistical analysis performed for each timepoint. Note: $EC_{50}$ values at week 2 were derived by setting the minimum/maximum $OD_{450}$ values to 0.0/2.8. (**c**) ELISA curves of mouse sera from EBOV GPΔmuc vaccine groups at week 17 after four immunizations binding to SUDV GPΔmuc-WL$^2$P$^4$(1TD0) trimer. Summary of geometric mean $EC_{50}$ titers and statistical analysis. (**d**) ELISA curves of mouse sera from Ebola GPΔmuc vaccine groups at week 17 after four immunizations binding to RAVV GPΔmuc-P$^2$CT(1TD0) trimer. Summary of absorbance at 450 nm (A450) values was included. (**e**) Neutralization curves of mouse sera from EBOV GPΔmuc trimer and SApNP vaccine groups against EBOV Makona pseudovirus. (**f**) (Top) Summary of geometric mean $ID_{50}$ titers measured for EBOV GPΔmuc vaccine groups against EBOV Makona pseudovirus. Color coding: white (no neutralization), green to red (low to high neutralization). Note: $ID_{50}$ values were calculated using %neutralization constraints of 0.0 (min) and 100.0 (max). (Bottom) Summary of statistical analysis as in (**b**). (**g**) Neutralization curves of mouse sera from EBOV GPΔmuc vaccine groups at week 17 against SUDV Gulu pseudovirus. Summary of geometric means of $ID_{50}$ values and statistical analysis. (**h**) Neutralization curves of mouse sera from EBOV GPΔmuc vaccine groups at week 17 against BDBV Uganda pseudovirus. Summary of geometric mean $ID_{50}$ values and statistical analysis. (**i**) ELISA curves of mouse sera from SUDV GPΔmuc trimer and SApNP vaccine groups binding to SUDV GPΔmuc-WL$^2$P$^4$(1TD0) trimer. (**j**) (Top) Summary of geometric means of $EC_{50}$ titers measured for SUDV GPΔmuc vaccine groups against SUDV GPΔmuc-WL$^2$P$^4$(1TD0) trimer. (Bottom) Summary of statistical analysis as in (**b**). Note: $EC_{50}$ values at week 2 were derived by setting the minimum/maximum $OD_{450}$ values to 0.0/2.9. (**k**) ELISA curves of mouse sera from SUDV GPΔmuc vaccine groups at week 17 after four immunizations binding to EBOV GPΔmuc-WL$^2$P$^4$(1TD0) trimer. Summary of geometric mean $EC_{50}$ titers and statistical analysis. (**l**) ELISA curves of mouse sera from SUDV GPΔmuc vaccine groups at week 17 after four immunizations binding to RAVV GPΔmuc-P$^2$CT(1TD0) trimer. Summary of A450 values was included. (**m**) Neutralization curves of mouse sera from SUDV GPΔmuc trimer and SApNP vaccine groups against SUDV Gulu pseudovirus. (**n**) (Top) Summary of geometric means of $ID_{50}$ titers measured for EBOV GPΔmuc vaccine groups against EBOV Makona pseudovirus. (Bottom) Summary of statistical analysis as in (**b**). (**o**) Neutralization curves of mouse sera from EBOV GPΔmuc vaccine groups at week 17 against EBOV Makona pseudovirus. Summary of geometric mean $ID_{50}$ values and statistical analysis. (**p**) Neutralization curves of mouse sera from EBOV GPΔmuc vaccine groups at week 17 against BDBV Uganda pseudovirus. Summary of geometric means of $ID_{50}$ values and statistical analysis. (**q**) (Top) ELISA curves of mouse sera from RAVV GPΔmuc trimer vaccine groups binding to RAVV GPΔmuc-P$^2$CT(1TD0) trimer. (Bottom) Summary of geometric mean $EC_{50}$ values. Note: $EC_{50}$ values at week 2 were derived by setting the minimum/maximum $OD_{450}$ values to 0.0/2.9. (**r**) ELISA curves of mouse sera from RAVV GPΔmuc vaccine groups at week 17 after four immunizations binding to EBOV GPΔmuc-WL$^2$P$^4$(1TD0) trimer. Summary of geometric mean $EC_{50}$ titers. (**s**) ELISA curves of mouse sera from RAVV GPΔmuc vaccine groups at week 17 after four immunizations binding to SUDV GPΔmuc-WL$^2$P$^4$(1TD0) trimer. Summary of A450 values was included. (**t**) Neutralization curves of purified IgG from RAVV GPΔmuc vaccine groups at week 17 against MARV Angola pseudovirus. Summary of $IC_{50}$ values. Left panel: sera from three naive mice showed nonspecific background in MARV Angola pseudovirus assays, suggesting that IgG purification is required to eliminate the nonspecific serum reactivity. (**u**) Neutralization curves of mouse sera from RAVV GPΔmuc vaccine groups at week 17 against orthoebolavirus strains. Error bars represent the difference between duplicate measurements at each concentration for each sample. $EC_{50}$, $ID_{50}$, and $IC_{50}$ values were calculated using GraphPad Prism version 10.3.1. Data were analyzed using one-way ANOVA, followed by Tukey's multiple comparison post hoc test for each timepoint. For significance, ns (not significant), *$p < 0.05$, **$p < 0.01$, ***$p < 0.001$, and ****$p < 0.0001$.

### Figure S11

**a Sera of individual mice immunized with glycan-modified EBOV vaccines binding to EBOV GPΔmuc-WL²P⁴(1TD0) trimer**

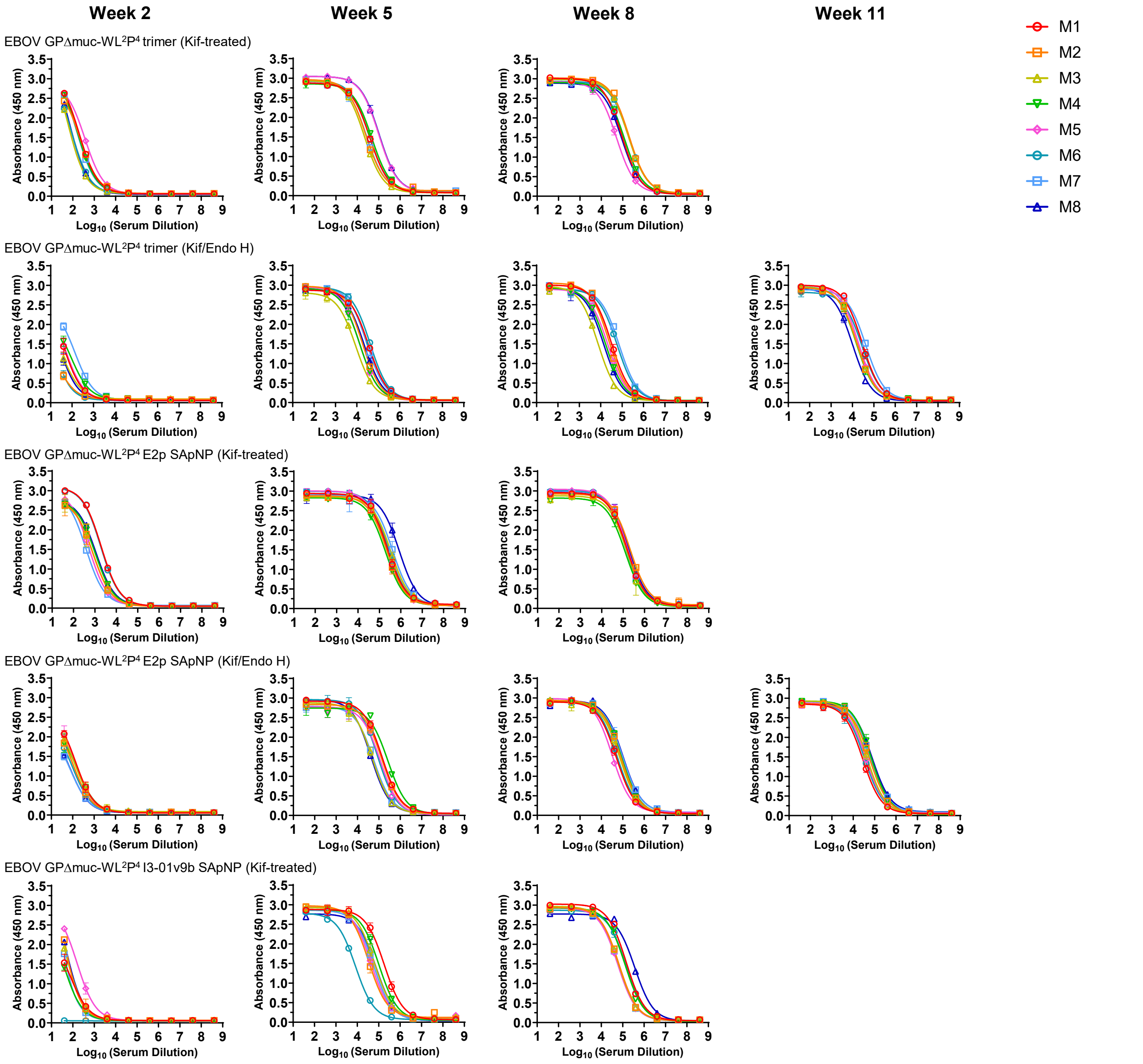

### Figure S11

#### b Mouse serum ELISA $EC_{50}$ titers

**Week 2**

| Antigen | $EC_{50}$ titers (week 2) | | | | | | | | Geometric Mean |
|---|---|---|---|---|---|---|---|---|---|
| | M1 | M2 | M3 | M4 | M5 | M6 | M7 | M8 | |
| EBOV GPΔmuc-WL$^2$P$^4$ trimer-Wild-type | 111.7 | 136.3 | 51.3 | 227.2 | 103.0 | 200.2 | 12.6 | 173.3 | 97.3 |
| EBOV GPΔmuc-WL$^2$P$^4$ trimer-Kif-treated | 283.7 | 255.1 | 119.5 | 248.0 | 423.8 | 133.4 | 239.1 | 141.3 | 212.1 |
| EBOV GPΔmuc-WL$^2$P$^4$ trimer-Kif/Endo H | 43.1 | 13.4 | 28.4 | 56.2 | 46.5 | 14.1 | 106.9 | 26.8 | 33.8 |
| EBOV GPΔmuc-WL$^2$P$^4$ E2p SApNP-Wild-type | 985.3 | 706.4 | 970.8 | 1772.0 | 271.5 | 336.1 | 3145.0 | 486.4 | 799.6 |
| EBOV GPΔmuc-WL$^2$P$^4$ E2p SApNP-Kif-treated | 2721.0 | 813.3 | 815.7 | 1046.0 | 678.3 | 2584.0 | 482.5 | 1166.0 | 1080.8 |
| EBOV GPΔmuc-WL$^2$P$^4$ E2p SApNP-Kif/Endo H | 126.0 | 84.5 | 104.9 | 95.8 | 122.8 | 69.1 | 50.0 | 51.4 | 83.4 |
| EBOV GPΔmuc-WL$^2$P$^4$ I3-01v9b SApNP-Wild-type | 30.1 | 101.8 | 43.0 | 37.9 | 172.4 | 80.7 | 91.5 | 115.6 | 72.1 |
| EBOV GPΔmuc-WL$^2$P$^4$ I3-01v9b SApNP-Kif-treated | 51.8 | 97.6 | 80.2 | 41.8 | 201.5 | 0.9 | 62.9 | 95.3 | 45.5 |

**Week 5**

| Antigen | $EC_{50}$ titers (week 5) | | | | | | | | Geometric Mean |
|---|---|---|---|---|---|---|---|---|---|
| | M1 | M2 | M3 | M4 | M5 | M6 | M7 | M8 | |
| EBOV GPΔmuc-WL$^2$P$^4$ trimer-Wild-type | 52545 | 41735 | 104914 | 48787 | 27947 | 18271 | 26144 | 21434 | 36588 |
| EBOV GPΔmuc-WL$^2$P$^4$ trimer-Kif-treated | 38706 | 25086 | 21695 | 47634 | 99875 | 36855 | 27014 | 103907 | 42360 |
| EBOV GPΔmuc-WL$^2$P$^4$ trimer-Kif/Endo H | 35186 | 18238 | 8909 | 13634 | 13630 | 41752 | 29227 | 22113 | 20285 |
| EBOV GPΔmuc-WL$^2$P$^4$ E2p SApNP-Wild-type | 199849 | 120954 | 303168 | 237335 | 74446 | 156667 | 282413 | 368341 | 195227 |
| EBOV GPΔmuc-WL$^2$P$^4$ E2p SApNP-Kif-treated | 241215 | 224657 | 335618 | 192147 | 216173 | 258075 | 406613 | 838779 | 300502 |
| EBOV GPΔmuc-WL$^2$P$^4$ E2p SApNP-Kif/Endo H | 141920 | 136153 | 48617 | 265817 | 117352 | 88543 | 56725 | 42434 | 94286 |
| EBOV GPΔmuc-WL$^2$P$^4$ I3-01v9b SApNP-Wild-type | 58458 | 93275 | 140476 | 104463 | 128829 | 148277 | 70114 | 63709 | 95342 |
| EBOV GPΔmuc-WL$^2$P$^4$ I3-01v9b SApNP-Kif-treated | 180905 | 34224 | 63566 | 102618 | 42941 | 8255 | 57374 | 74974 | 52927 |

**Week 11 or 8**

| Antigen | $EC_{50}$ titers (week 11 or 8) | | | | | | | | Geometric Mean |
|---|---|---|---|---|---|---|---|---|---|
| | M1 | M2 | M3 | M4 | M5 | M6 | M7 | M8 | |
| EBOV GPΔmuc-WL$^2$P$^4$ trimer-Wild-type (w11) | 26545 | 524033 | 69054 | 94400 | 39464 | 69865 | 68351 | 51944 | 73881 |
| EBOV GPΔmuc-WL$^2$P$^4$ trimer-Kif-treated (w8) | 93214 | 195305 | 183599 | 116631 | 50918 | 193472 | 114590 | 89398 | 118674 |
| EBOV GPΔmuc-WL$^2$P$^4$ trimer-Kif/Endo H (w8) | 31571 | 25030 | 7454 | 16954 | 21834 | 59045 | 72580 | 13927 | 24508 |
| EBOV GPΔmuc-WL$^2$P$^4$ E2p SApNP-Wild-type (w11) | 267135 | 163352 | 223491 | 606328 | 32762 | 143908 | 182928 | 233182 | 181722 |
| EBOV GPΔmuc-WL$^2$P$^4$ E2p SApNP-Kif-treated (w8) | 153387 | 211525 | 149619 | 126857 | 158759 | 131151 | 152440 | 176520 | 155680 |
| EBOV GPΔmuc-WL$^2$P$^4$ E2p SApNP-Kif/Endo H (w8) | 50455 | 81205 | 73372 | 89472 | 32112 | 55202 | 99817 | 102711 | 68571 |
| EBOV GPΔmuc-WL$^2$P$^4$ I3-01v9b SApNP-Wild-type (w11) | 703967 | 60730 | 178784 | 354031 | 97482 | 91294 | 82827 | 610293 | 182253 |
| EBOV GPΔmuc-WL$^2$P$^4$ I3-01v9b SApNP-Kif-treated (w8) | 146649 | 66233 | 62594 | 129059 | 58475 | 142777 | 61089 | 366113 | 104890 |

#### Statistical analysis

| One-way ANOVA with Tukey's multiple comparisons test (w2) | Statistics | Adjusted P Value |
|---|---|---|
| EBOV GPΔmuc-WL$^2$P$^4$ trimer-Wild-type vs. EBOV GPΔmuc-WL$^2$P$^4$ trimer-Kif-treated | * | 0.0276 |
| EBOV GPΔmuc-WL$^2$P$^4$ trimer-Wild-type vs. EBOV GPΔmuc-WL$^2$P$^4$ trimer-Kif/Endo H | ns | 0.0775 |
| EBOV GPΔmuc-WL$^2$P$^4$ trimer-Kif-treated vs. EBOV GPΔmuc-WL$^2$P$^4$ trimer-Kif/Endo H | *** | 0.0001 |

| One-way ANOVA with Tukey's multiple comparisons test (w5) | Statistics | Adjusted P Value |
|---|---|---|
| EBOV GPΔmuc-WL$^2$P$^4$ trimer-Wild-type vs. EBOV GPΔmuc-WL$^2$P$^4$ trimer-Kif-treated | ns | 0.8381 |
| EBOV GPΔmuc-WL$^2$P$^4$ trimer-Wild-type vs. EBOV GPΔmuc-WL$^2$P$^4$ trimer-Kif/Endo H | ns | 0.2962 |
| EBOV GPΔmuc-WL$^2$P$^4$ trimer-Kif-treated vs. EBOV GPΔmuc-WL$^2$P$^4$ trimer-Kif/Endo H | ns | 0.1137 |

| One-way ANOVA with Tukey's multiple comparisons test (w2) | Statistics | Adjusted P Value |
|---|---|---|
| EBOV GPΔmuc-WL$^2$P$^4$ E2p SApNP-Wild-type vs. EBOV GPΔmuc-WL$^2$P$^4$ E2p SApNP-Kif-treated | ns | 0.8496 |
| EBOV GPΔmuc-WL$^2$P$^4$ E2p SApNP-Wild-type vs. EBOV GPΔmuc-WL$^2$P$^4$ E2p SApNP-Kif/Endo H | * | 0.0371 |
| EBOV GPΔmuc-WL$^2$P$^4$ E2p SApNP-Kif-treated vs. EBOV GPΔmuc-WL$^2$P$^4$ E2p SApNP-Kif/Endo H | * | 0.0112 |

| One-way ANOVA with Tukey's multiple comparisons test (w5) | Statistics | Adjusted P Value |
|---|---|---|
| EBOV GPΔmuc-WL$^2$P$^4$ E2p SApNP-Wild-type vs. EBOV GPΔmuc-WL$^2$P$^4$ E2p SApNP-Kif-treated | ns | 0.2278 |
| EBOV GPΔmuc-WL$^2$P$^4$ E2p SApNP-Wild-type vs. EBOV GPΔmuc-WL$^2$P$^4$ E2p SApNP-Kif/Endo H | ns | 0.3186 |
| EBOV GPΔmuc-WL$^2$P$^4$ E2p SApNP-Kif-treated vs. EBOV GPΔmuc-WL$^2$P$^4$ E2p SApNP-Kif/Endo H | * | 0.0119 |

| Unpaired t test (w2) | Statistics | P Value |
|---|---|---|
| EBOV GPΔmuc-WL$^2$P$^4$ I3-01v9b SApNP-Wild-type vs. EBOV GPΔmuc-WL$^2$P$^4$ I3-01v9b SApNP-Kif-treated | ns | 0.8507 |

| Unpaired t test (w5) | Statistics | P Value |
|---|---|---|
| EBOV GPΔmuc-WL$^2$P$^4$ I3-01v9b SApNP-Wild-type vs. EBOV GPΔmuc-WL$^2$P$^4$ I3-01v9b SApNP-Kif-treated | ns | 0.1978 |

### Figure S11

#### C Sera of individual mice immunized with EBOV vaccines neutralizing an EBOV Makona strain

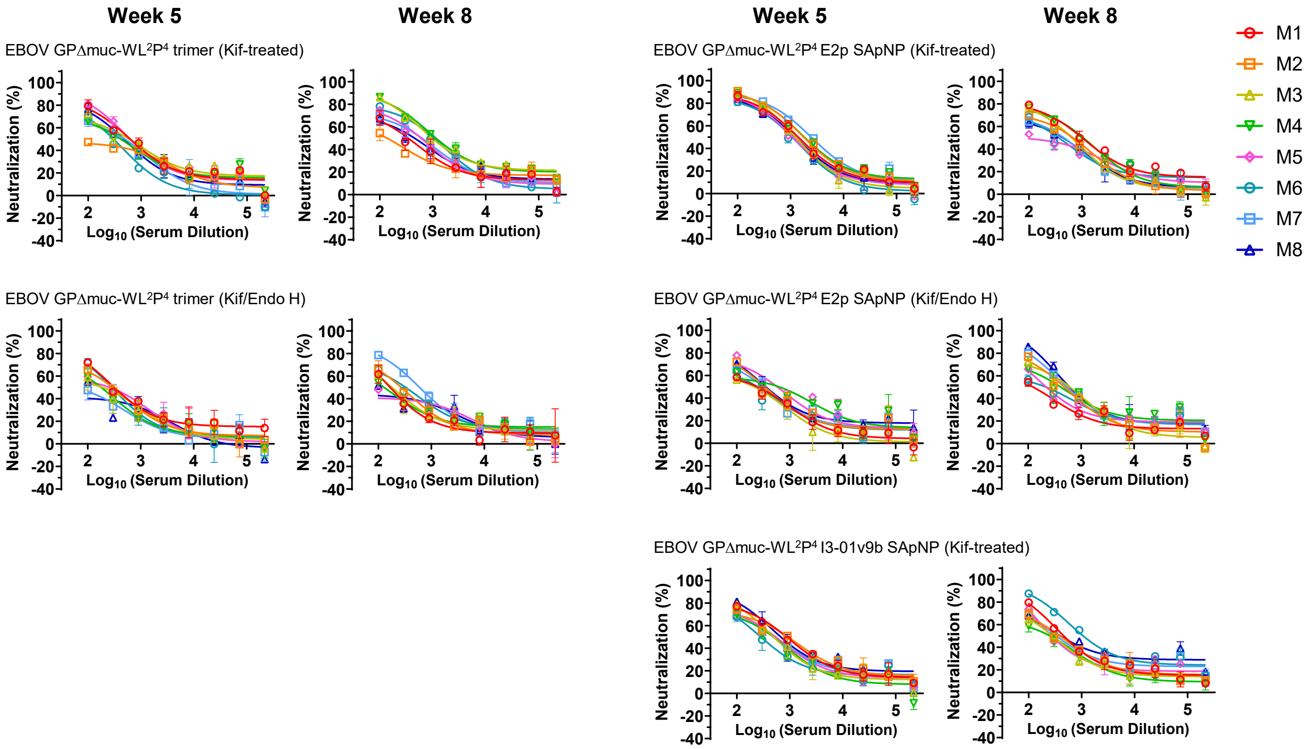


#### d Mouse serum neutralizing $ID_{50}$ titers

**Week 5**

| Antigen | $ID_{50}$ titers (week 5) | | | | | | | | Geometric Mean |
|---|---|---|---|---|---|---|---|---|---|
| | M1 | M2 | M3 | M4 | M5 | M6 | M7 | M8 | |
| EBOV GPΔmuc-WL$^2$P$^4$ trimer-Wild-type | 805.4 | 559.3 | 149.3 | 588.2 | 174.1 | 145.1 | 203.2 | 360.5 | 304.1 |
| EBOV GPΔmuc-WL$^2$P$^4$ trimer-Kif-treated | 685.5 | 234.7 | 577.4 | 443.0 | 708.5 | 285.4 | 407.7 | 489.1 | 449.3 |
| EBOV GPΔmuc-WL$^2$P$^4$ trimer-Kif/Endo H | 383.7 | 299.7 | 227.6 | 199.9 | 271.6 | 298.0 | 132.4 | 145.5 | 231.2 |
| EBOV GPΔmuc-WL$^2$P$^4$ E2p SApNP-Wild-type | 636.9 | 358.8 | 2178.0 | 1002.0 | 555.2 | 322.9 | 299.4 | 1167.0 | 648.4 |
| EBOV GPΔmuc-WL$^2$P$^4$ E2p SApNP-Kif-treated | 1414.0 | 1867.0 | 1249.0 | 1961.0 | 1107.0 | 963.3 | 2480.0 | 1201.0 | 1459.0 |
| EBOV GPΔmuc-WL$^2$P$^4$ E2p SApNP-Kif/Endo H | 263.2 | 376.3 | 232.9 | 513.0 | 559.4 | 273.6 | 353.0 | 430.4 | 358.9 |
| EBOV GPΔmuc-WL$^2$P$^4$ I3-01v9b SApNP-Wild-type | 881.2 | 850.8 | 555.5 | 452.4 | 550.2 | 340.8 | 548.8 | 865.0 | 599.9 |
| EBOV GPΔmuc-WL$^2$P$^4$ I3-01v9b SApNP-Kif-treated | 267.9 | 390.2 | 313.1 | 575.8 | 378.9 | 413.4 | 258.7 | 296.7 | 350.3 |

**Week 11 or 8**

| Antigen | $ID_{50}$ titers (week 11 or 8) | | | | | | | | Geometric Mean |
|---|---|---|---|---|---|---|---|---|---|
| | M1 | M2 | M3 | M4 | M5 | M6 | M7 | M8 | |
| EBOV GPΔmuc-WL$^2$P$^4$ trimer-Wild-type (w11) | 2117.0 | 560.6 | 254.4 | 521.5 | 188.7 | 430.2 | 369.3 | 377.3 | 453.2 |
| EBOV GPΔmuc-WL$^2$P$^4$ trimer-Kif-treated (w8) | 370.3 | 204.5 | 1305 | 1425 | 578 | 911.8 | 569.1 | 448.8 | 609.1 |
| EBOV GPΔmuc-WL$^2$P$^4$ trimer-Kif/Endo H (w8) | 192.8 | 287.4 | 217.4 | 178.6 | 171.6 | 351.1 | 636.2 | 175.4 | 248.3 |
| EBOV GPΔmuc-WL$^2$P$^4$ E2p SApNP-Wild-type (w11) | 494.5 | 392.0 | 734.2 | 747.4 | 275.3 | 643.8 | 365.2 | 759.9 | 518.6 |
| EBOV GPΔmuc-WL$^2$P$^4$ E2p SApNP-Kif-treated (w8) | 1031 | 562.3 | 594.7 | 902.2 | 273 | 393.9 | 416.3 | 408.9 | 524.1 |
| EBOV GPΔmuc-WL$^2$P$^4$ E2p SApNP-Kif/Endo H (w8) | 179.2 | 499.6 | 538.7 | 486.6 | 301.2 | 292.4 | 649.2 | 772.1 | 423.6 |
| EBOV GPΔmuc-WL$^2$P$^4$ I3-01v9b SApNP-Wild-type (w11) | 1006.0 | 434.9 | 454.1 | 766.3 | 289.4 | 237.7 | 327.0 | 928.5 | 487.3 |
| EBOV GPΔmuc-WL$^2$P$^4$ I3-01v9b SApNP-Kif-treated (w8) | 562.9 | 429.3 | 326.9 | 281.2 | 376.1 | 1462 | 459.4 | 603.8 | 491.2 |

##### Statistical analysis

| One-way ANOVA with Tukey's multiple comparisons test (w5) | Statistics | Adjusted P Value |
|---|---|---|
| EBOV GPΔmuc-WL$^2$P$^4$ trimer-Wild-type vs. EBOV GPΔmuc-WL$^2$P$^4$ trimer-Kif-treated | ns | 0.489 |
| EBOV GPΔmuc-WL$^2$P$^4$ trimer-Wild-type vs. EBOV GPΔmuc-WL$^2$P$^4$ trimer-Kif/Endo H | ns | 0.3549 |
| EBOV GPΔmuc-WL$^2$P$^4$ trimer-Kif-treated vs. EBOV GPΔmuc-WL$^2$P$^4$ trimer-Kif/Endo H | * | 0.0453 |

| One-way ANOVA with Tukey's multiple comparisons test (w5) | Statistics | Adjusted P Value |
|---|---|---|
| EBOV GPΔmuc-WL$^2$P$^4$ E2p SApNP-Wild-type vs. EBOV GPΔmuc-WL$^2$P$^4$ E2p SApNPr-Kif-treated | * | 0.02 |
| EBOV GPΔmuc-WL$^2$P$^4$ E2p SApNP-Wild-type vs. EBOV GPΔmuc-WL$^2$P$^4$ E2p SApNP-Kif/Endo H | ns | 0.1792 |
| EBOV GPΔmuc-WL$^2$P$^4$ E2p SApNPr-Kif-treated vs. EBOV GPΔmuc-WL$^2$P$^4$ E2p SApNP-Kif/Endo H | *** | 0.0003 |

| Unpaired t test (w5) | Statistics | P Value |
|---|---|---|
| EBOV GPΔmuc-WL$^2$P$^4$ I3-01v9b SApNP-Wild-type vs. EBOV GPΔmuc-WL$^2$P$^4$ I3-01v9b SApNP-Kif-treated | ** | 0.0055 |

#### Figure S11

**e Sera of individual mice immunized with wild-type/glycan-modified SUDV vaccines binding to SUDV GPΔmuc-WL²P⁴(1TD0) trimer**

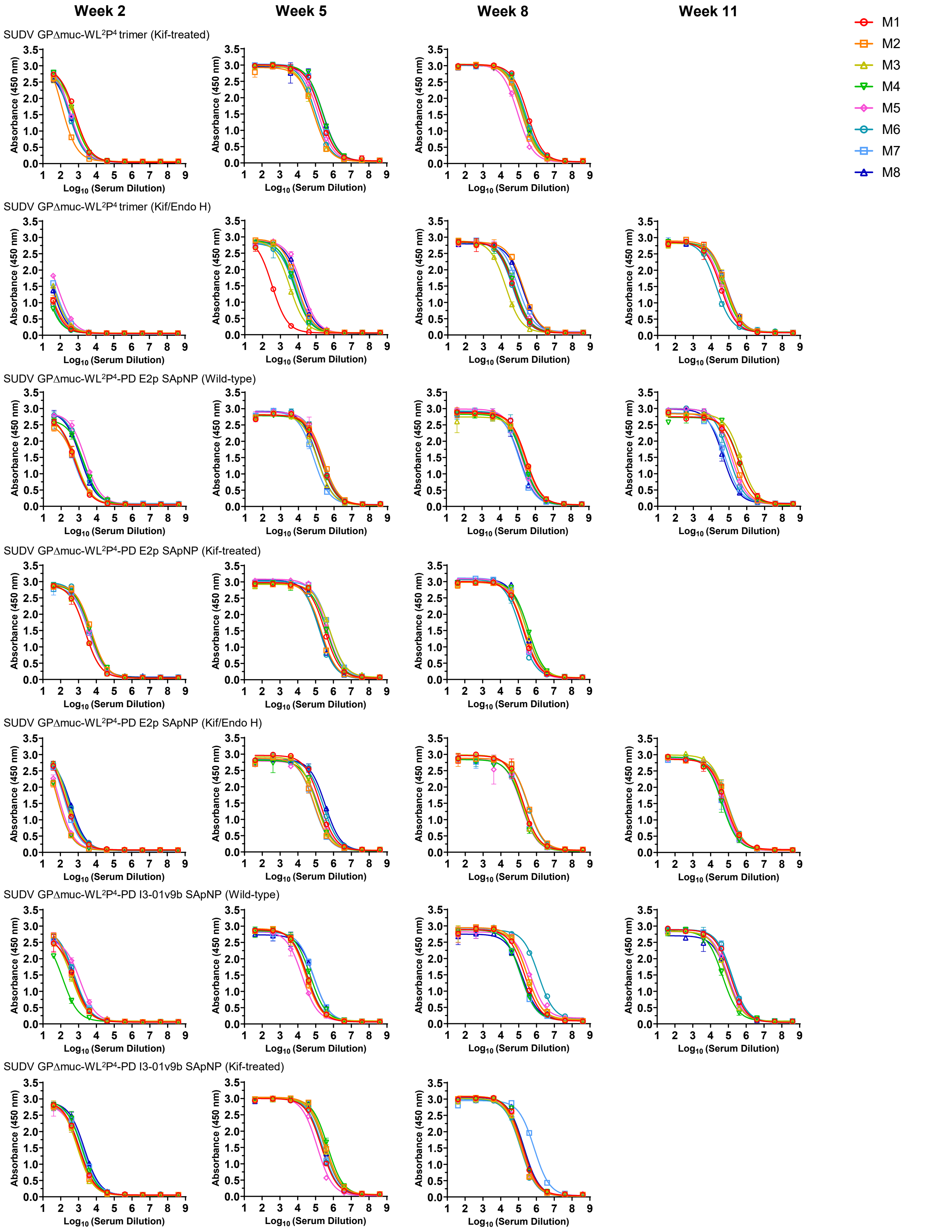

### Figure S11

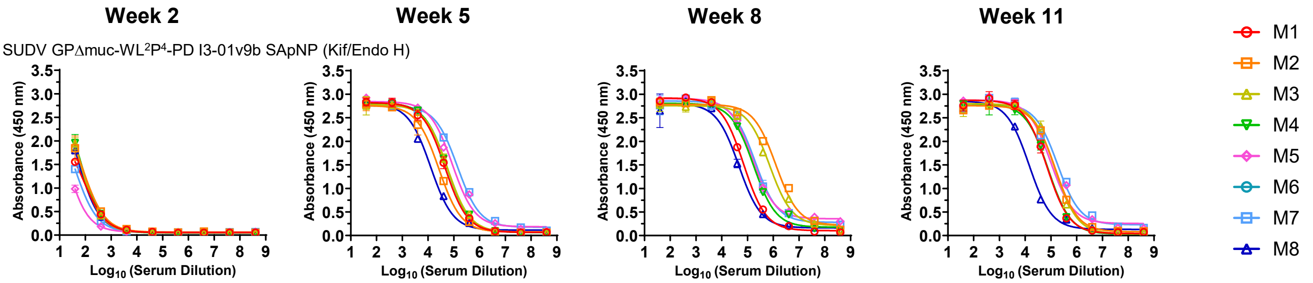


#### f Mouse serum ELISA $EC_{50}$ titers

**Week 2**

| Antigen | $EC_{50}$ titers (week 2) | | | | | | | | Geometric Mean |
|---|---|---|---|---|---|---|---|---|---|
| | M1 | M2 | M3 | M4 | M5 | M6 | M7 | M8 | |
| SUDV GPΔmuc-WL$^2$P$^4$ trimer-Wild-type | 69.8 | 20.2 | 50.0 | 149.7 | 116.9 | 122.4 | 68.1 | 55.0 | 69.8 |
| SUDV GPΔmuc-WL$^2$P$^4$ trimer-Kif-treated | 714.8 | 192.9 | 605.3 | 591.0 | 412.8 | 343.0 | 419.8 | 362.8 | 425.0 |
| SUDV GPΔmuc-WL$^2$P$^4$ trimer-Kif/Endo H | 23.3 | 24.5 | 44.7 | 15.7 | 72.2 | 19.6 | 50.5 | 36.9 | 31.8 |
| SUDV GPΔmuc-WL$^2$P$^4$-PD E2p SApNP-Wild-type | 527.1 | 474.8 | 583.8 | 1483.0 | 2336.0 | 1545.0 | 475.4 | 1316.0 | 914.5 |
| SUDV GPΔmuc-WL$^2$P$^4$-PD E2p SApNP-Kif-treated | 2482.0 | 5568.0 | 2483.0 | 4798.0 | 4273.0 | 4408.0 | 3878.0 | 4312.0 | 3884.6 |
| SUDV GPΔmuc-WL$^2$P$^4$-PD E2p SApNP-Kif/Endo H | 271.0 | 99.3 | 330.9 | 99.8 | 124.0 | 375.7 | 234.7 | 410.6 | 211.4 |
| SUDV GPΔmuc-WL$^2$P$^4$-PD I3-01v9b SApNP-Wild-type | 469.6 | 479.9 | 387.3 | 115.2 | 890.0 | 616.4 | 586.8 | 503.1 | 448.2 |
| SUDV GPΔmuc-WL$^2$P$^4$-PD I3-01v9b SApNP-Kif-treated | 1206.0 | 1067.0 | 961.1 | 1644.0 | 1098.0 | 1237.0 | 1839.0 | 2248.0 | 1355.8 |
| SUDV GPΔmuc-WL$^2$P$^4$-PD I3-01v9b SApNP-Kif/Endo H | 49.8 | 73.8 | 77.8 | 80.5 | 20.8 | N/A | 38.4 | 65.2 | 53.1 |

**Week 5**

| Antigen | $EC_{50}$ titers (week 5) | | | | | | | | Geometric Mean |
|---|---|---|---|---|---|---|---|---|---|
| | M1 | M2 | M3 | M4 | M5 | M6 | M7 | M8 | |
| SUDV GPΔmuc-WL$^2$P$^4$ trimer-Wild-type | 16436 | 19941 | 7907 | 60277 | 25656 | 19754 | 9992 | 15619 | 18259.2 |
| SUDV GPΔmuc-WL$^2$P$^4$ trimer-Kif-treated | 187796 | 83838 | 82374 | 256237 | 143796 | 100245 | 199748 | 272537 | 150326.2 |
| SUDV GPΔmuc-WL$^2$P$^4$ trimer-Kif/Endo H | 337 | 8926 | 3406 | 6542 | 16601 | 6364 | 8124 | 13307 | 5439.3 |
| SUDV GPΔmuc-WL$^2$P$^4$-PD E2p SApNP-Wild-type | 194403 | 254967 | 120302 | 189915 | 195340 | 209194 | 72385 | 116111 | 158028.4 |
| SUDV GPΔmuc-WL$^2$P$^4$-PD E2p SApNP-Kif-treated | 328808 | 203797 | 637646 | 429942 | 440079 | 160965 | 596309 | 178438 | 329358.5 |
| SUDV GPΔmuc-WL$^2$P$^4$-PD E2p SApNP-Kif/Endo H | 174902 | 82685 | 108551 | 129398 | 143808 | 261848 | 79238 | 345641 | 146267.1 |
| SUDV GPΔmuc-WL$^2$P$^4$-PD I3-01v9b SApNP-Wild-type | 32933 | 27405 | 30608 | 50359 | 17406 | 34313 | 79188 | 83439 | 39124.1 |
| SUDV GPΔmuc-WL$^2$P$^4$-PD I3-01v9b SApNP-Kif-treated | 213630 | 375263 | 331165 | 484567 | 123702 | 233165 | 246351 | 324809 | 271692.3 |
| SUDV GPΔmuc-WL$^2$P$^4$-PD I3-01v9b SApNP-Kif/Endo H | 45679 | 26310 | 57270 | 55138 | 87196 | N/A | 138210 | 12630 | 47889.1 |

**Week 11 or 8**

| Antigen | $EC_{50}$ titers (week 11 or 8) | | | | | | | | Geometric Mean |
|---|---|---|---|---|---|---|---|---|---|
| | M1 | M2 | M3 | M4 | M5 | M6 | M7 | M8 | |
| SUDV GPΔmuc-WL$^2$P$^4$ trimer-Wild-type (w11) | 51852 | 34819 | 59219 | 58035 | 92570 | 20182 | 108658 | 56856 | 53935.6 |
| SUDV GPΔmuc-WL$^2$P$^4$ trimer-Kif-treated (w8) | 302797 | 144377 | 177306 | 198140 | 88084 | 235081 | 164275 | 185242 | 177101.1 |
| SUDV GPΔmuc-WL$^2$P$^4$ trimer-Kif/Endo H (w8) | 50453 | 172980 | 18107 | 59100 | 58943 | 44125 | 87452 | 156514 | 65346.1 |
| SUDV GPΔmuc-WL$^2$P$^4$-PD E2p SApNP-Wild-type (w8) | 299781 | 269350 | 317854 | 207812 | 165230 | 208403 | 120522 | 131427 | 203221.8 |
| SUDV GPΔmuc-WL$^2$P$^4$-PD E2p SApNP-Kif-treated (w8) | 192720 | 218683 | 292392 | 372944 | 179861 | 130425 | 179178 | 274004 | 219008.7 |
| SUDV GPΔmuc-WL$^2$P$^4$-PD E2p SApNP-Kif/Endo H (w8) | 180099 | 336638 | 152272 | 156367 | 145160 | 325977 | 149103 | N/A | 193576.0 |
| JDV GPΔmuc-WL$^2$P$^4$-PD I3-01v9b SApNP-Wild-type (w8) | 219021 | 293708 | 296644 | 131589 | 450528 | 1270551 | 122274 | 162786 | 270436.4 |
| JDV GPΔmuc-WL$^2$P$^4$-PD I3-01v9b SApNP-Kif-treated (w8) | 166508 | 138485 | 135652 | 171299 | 143035 | 129149 | 754862 | 196507 | 186573.4 |
| IDV GPΔmuc-WL$^2$P$^4$-PD I3-01v9b SApNP-Kif/Endo H (w8) | 71039 | 1261224 | 728092 | 170130 | 156138 | N/A | 198137 | 45999 | 206095.4 |

#### Statistical analysis

| One-way ANOVA with Tukey's multiple comparisons test (w2) | Statistics | Adjusted P Value |
|---|---|---|
| SUDV GPΔmuc-WL$^2$P$^4$ trimer-Wild-type vs. SUDV GPΔmuc-WL$^2$P$^4$ trimer-Kif-treated | **** | <0.0001 |
| SUDV GPΔmuc-WL$^2$P$^4$ trimer-Wild-type vs. SUDV GPΔmuc-WL$^2$P$^4$ trimer-Kif/Endo H | ns | 0.6485 |
| SUDV GPΔmuc-WL$^2$P$^4$ trimerKif-treated vs. SUDV GPΔmuc-WL$^2$P$^4$ trimer-Kif/Endo H | **** | <0.0001 |

| One-way ANOVA with Tukey's multiple comparisons test (w2) | Statistics | Adjusted P Value |
|---|---|---|
| SUDV GPΔmuc-WL$^2$P$^4$-PD E2p SApNP-Wild-type vs. SUDV GPΔmuc-WL$^2$P$^4$-PD E2p SApNP-Kif-treated | **** | <0.0001 |
| SUDV GPΔmuc-WL$^2$P$^4$-PD E2p SApNP-Wild-type vs. SUDV GPΔmuc-WL$^2$P$^4$-PD E2p SApNP-Kif/Endo H | ns | 0.0778 |
| SUDV GPΔmuc-WL$^2$P$^4$-PD E2p SApNP-Kif-treated vs. SUDV GPΔmuc-WL$^2$P$^4$-PD E2p SApNP-Kif/Endo H | **** | <0.0001 |

| One-way ANOVA with Tukey's multiple comparisons test (w2) | Statistics | Adjusted P Value |
|---|---|---|
| SUDV GPΔmuc-WL$^2$P$^4$-PD I3-01v9b SApNP-Wild-type vs. SUDV GPΔmuc-WL$^2$P$^4$-PD I3-01v9b SApNP-Kif-treated | **** | <0.0001 |
| SUDV GPΔmuc-WL$^2$P$^4$-PD I3-01v9b SApNP-Wild-type vs. SUDV GPΔmuc-WL$^2$P$^4$-PD I3-01v9b SApNP-Kif/Endo H | * | 0.0224 |
| SUDV GPΔmuc-WL$^2$P$^4$-PD I3-01v9b SApNP-Kif-treated vs. SUDV GPΔmuc-WL$^2$P$^4$-PD I3-01v9b SApNP-Kif/Endo H | **** | <0.0001 |

| One-way ANOVA with Tukey's multiple comparisons test (w5) | Statistics | Adjusted P Value |
|---|---|---|
| SUDV GPΔmuc-WL$^2$P$^4$ trimer-Wild-type vs. SUDV GPΔmuc-WL$^2$P$^4$ trimer-Kif-treated | **** | <0.0001 |
| SUDV GPΔmuc-WL$^2$P$^4$ trimer-Wild-type vs. SUDV GPΔmuc-WL$^2$P$^4$ trimer-Kif/Endo H | ns | 0.8066 |
| SUDV GPΔmuc-WL$^2$P$^4$ trimerKif-treated vs. SUDV GPΔmuc-WL$^2$P$^4$ trimer-Kif/Endo H | **** | <0.0001 |

| One-way ANOVA with Tukey's multiple comparisons test (w5) | Statistics | Adjusted P Value |
|---|---|---|
| SUDV GPΔmuc-WL$^2$P$^4$-PD E2p SApNP-Wild-type vs. SUDV GPΔmuc-WL$^2$P$^4$-PD E2p SApNP-Kif-treated | * | 0.0103 |
| SUDV GPΔmuc-WL$^2$P$^4$-PD E2p SApNP-Wild-type vs. SUDV GPΔmuc-WL$^2$P$^4$-PD E2p SApNP-Kif/Endo H | ns | 0.9984 |
| SUDV GPΔmuc-WL$^2$P$^4$-PD E2p SApNP-Kif-treated vs. SUDV GPΔmuc-WL$^2$P$^4$-PD E2p SApNP-Kif/Endo H | ** | 0.0091 |

| One-way ANOVA with Tukey's multiple comparisons test (w5) | Statistics | Adjusted P Value |
|---|---|---|
| SUDV GPΔmuc-WL$^2$P$^4$-PD I3-01v9b SApNP-Wild-type vs. SUDV GPΔmuc-WL$^2$P$^4$-PD I3-01v9b SApNP-Kif-treated | **** | <0.0001 |
| SUDV GPΔmuc-WL$^2$P$^4$-PD I3-01v9b SApNP-Wild-type vs. SUDV GPΔmuc-WL$^2$P$^4$-PD I3-01v9b SApNP-Kif/Endo H | ns | 0.903 |
| SUDV GPΔmuc-WL$^2$P$^4$-PD I3-01v9b SApNP-Kif-treated vs. SUDV GPΔmuc-WL$^2$P$^4$-PD I3-01v9b SApNP-Kif/Endo H | **** | <0.0001 |

### Figure S11

#### g Sera of individual mice immunized with wild-type/glycan-modified SUDV vaccines neutralizing a SUDV Gulu strain

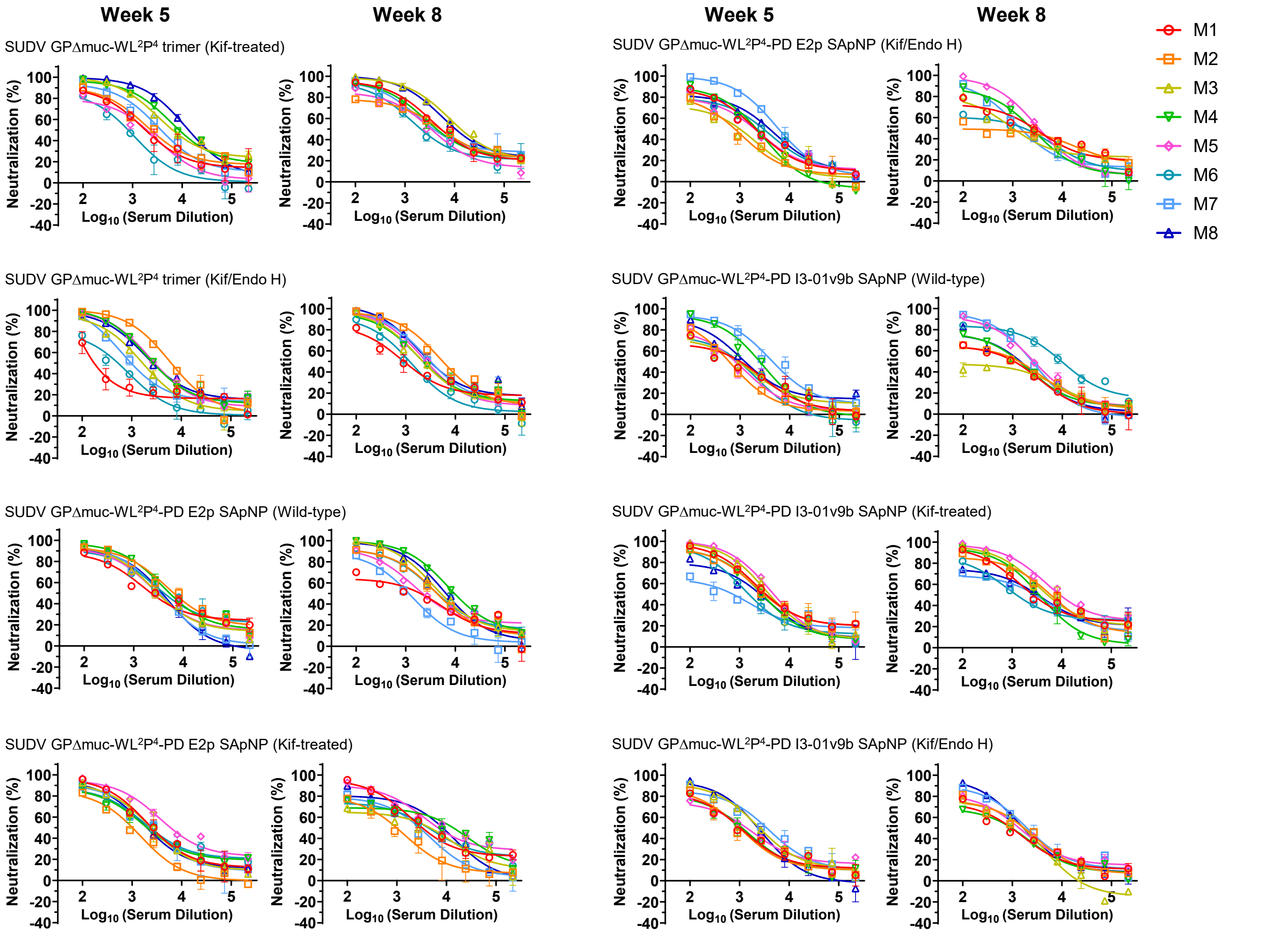


#### h Mouse serum neutralizing $ID_{50}$ titers

**Week 5**

| Antigen | $ID_{50}$ titers (week 5) M1 | M2 | M3 | M4 | M5 | M6 | M7 | M8 | Geometric Mean |
|---|---|---|---|---|---|---|---|---|---|
| SUDV GPΔmuc-WL$^2$P$^4$ trimer-Wild-type | 420 | 3076 | 156.6 | 2417 | 1377 | 1896 | 5149 | 1477 | 1328.6 |
| SUDV GPΔmuc-WL$^2$P$^4$ trimer-Kif-treated | 1986 | 2709 | 8760 | 10520 | 1522 | 736.6 | 4016 | 14183 | 3652.3 |
| SUDV GPΔmuc-WL$^2$P$^4$ trimer-Kif/Endo H | 243.4 | 6460 | 1691 | 3127 | 3197.0 | 463.3 | 1201.0 | 3265.0 | 1623.6 |
| SUDV GPΔmuc-WL$^2$P$^4$-PD E2p SApNP-Wild-type | 3142 | 7835 | 3520 | 6205 | 3431 | 5382 | 3249 | 3187 | 4231.6 |
| SUDV GPΔmuc-WL$^2$P$^4$-PD E2p SApNP-Kif-treated | 2877 | 873.4 | 2512 | 2311 | 7871 | 2706 | 2473 | 2186 | 2530.2 |
| SUDV GPΔmuc-WL$^2$P$^4$-PD E2p SApNP-Kif/Endo H | 2075 | 721.3 | 713.9 | 1791 | 1742 | 2481 | 6384 | 3462 | 1917.5 |
| SUDV GPΔmuc-WL$^2$P$^4$-PD I3-01v9b SApNP-Wild-type | 662.4 | 592.3 | 758.7 | 2268 | 682.9 | 642.8 | 4778 | 1402 | 1089.5 |
| SUDV GPΔmuc-WL$^2$P$^4$-PD I3-01v9b SApNP-Kif-treated | 4252 | 3819 | 4262 | 2863 | 5193 | 1734 | 651.4 | 2141 | 2657.6 |
| SUDV GPΔmuc-WL$^2$P$^4$-PD I3-01v9b SApNP-Kif/Endo H | 1023 | 1000 | 2652 | 1043 | 1287 | N/A | 3262 | 2177 | 1591.5 |

**Week 11 or 8**

| Antigen | $ID_{50}$ titers (week 11 or 8) M1 | M2 | M3 | M4 | M5 | M6 | M7 | M8 | Geometric Mean |
|---|---|---|---|---|---|---|---|---|---|
| SUDV GPΔmuc-WL$^2$P$^4$ trimer-Wild-type (w11) | 13810 | 4694 | 2805 | 3670 | 2858 | 2497 | 14410 | 3069 | 4628.4 |
| SUDV GPΔmuc-WL$^2$P$^4$ trimer-Kif-treated (w8) | 6582 | 5258 | 16937 | 5674 | 3316 | 2649 | 5681 | 13235 | 6204.7 |
| SUDV GPΔmuc-WL$^2$P$^4$ trimer-Kif/Endo H (w8) | 1035 | 6006 | 2560 | 2543 | 2847 | 1139 | 3639 | 3835 | 2557.7 |
| SUDV GPΔmuc-WL$^2$P$^4$-PD E2p SApNP-Wild-type (w8) | 1224 | 4430 | 6202 | 11133 | 2887 | 4953 | 1083 | 6855 | 3757.6 |
| SUDV GPΔmuc-WL$^2$P$^4$-PD E2p SApNP-Kif-treated (w8) | 4414 | 880.9 | 2466 | 7047 | 10493 | 4060 | 2235 | 6092 | 3751.0 |
| SUDV GPΔmuc-WL$^2$P$^4$-PD E2p SApNP-Kif/Endo H (w8) | 2234 | 555.2 | 1224 | 2211 | 3512 | 1040 | 1615 | N/A | 1531.9 |
| SUDV GPΔmuc-WL$^2$P$^4$-PD I3-01v9b SApNP-Wild-type (w8) | 739.1 | 911.5 | 343.7 | 1130 | 2675 | 9196 | 2305 | 1040 | 1407.8 |
| SUDV GPΔmuc-WL$^2$P$^4$-PD I3-01v9b SApNP-Kif-treated (w8) | 3830 | 5923 | 6805 | 3469 | 12386 | 1653 | 2090 | 3103 | 4041.0 |
| SUDV GPΔmuc-WL$^2$P$^4$-PD I3-01v9b SApNP-Kif/Endo H (w8) | 810.8 | 1367 | 1095 | 777.2 | 1388 | N/A | 2057 | 1791 | 1252.0 |

#### Statistical analysis

| One-way ANOVA with Tukey's multiple comparisons test (w5) | Statistics | Adjusted P Value |
|---|---|---|
| SUDV GPΔmuc-WL$^2$P$^4$ trimer-Wild-type vs. SUDV GPΔmuc-WL$^2$P$^4$ trimer-Kif-treated | ns | 0.0934 |
| SUDV GPΔmuc-WL$^2$P$^4$ trimer-Wild-type vs. SUDV GPΔmuc-WL$^2$P$^4$ trimer-Kif/Endo H | ns | 0.9563 |
| SUDV GPΔmuc-WL$^2$P$^4$ trimerKif-treated vs. SUDV GPΔmuc-WL$^2$P$^4$ trimer-Kif/Endo H | ns | 0.1577 |

| One-way ANOVA with Tukey's multiple comparisons test (w5) | Statistics | Adjusted P Value |
|---|---|---|
| SUDV GPΔmuc-WL$^2$P$^4$-PD E2p SApNP-Wild-type vs. SUDV GPΔmuc-WL$^2$P$^4$-PD E2p SApNP-Kif-treated | ns | 0.2678 |
| SUDV GPΔmuc-WL$^2$P$^4$-PD E2p SApNP-Wild-type vs. SUDV GPΔmuc-WL$^2$P$^4$-PD E2p SApNP-Kif/Endo H | ns | 0.0973 |
| SUDV GPΔmuc-WL$^2$P$^4$-PD E2p SApNP-Kif-treated vs. SUDV GPΔmuc-WL$^2$P$^4$-PD E2p SApNP-Kif/Endo H | ns | 0.8296 |

| One-way ANOVA with Tukey's multiple comparisons test (w8) | Statistics | Adjusted P Value |
|---|---|---|
| SUDV GPΔmuc-WL$^2$P$^4$-PD E2p SApNP-Wild-type vs. SUDV GPΔmuc-WL$^2$P$^4$-PD E2p SApNP-Kif-treated | ns | 0.9947 |
| SUDV GPΔmuc-WL$^2$P$^4$-PD E2p SApNP-Wild-type vs. SUDV GPΔmuc-WL$^2$P$^4$-PD E2p SApNP-Kif/Endo H | ns | 0.1006 |
| SUDV GPΔmuc-WL$^2$P$^4$-PD E2p SApNP-Kif-treated vs. SUDV GPΔmuc-WL$^2$P$^4$-PD E2p SApNP-Kif/Endo H | ns | 0.1201 |

| One-way ANOVA with Tukey's multiple comparisons test (w5) | Statistics | Adjusted P Value |
|---|---|---|
| SUDV GPΔmuc-WL$^2$P$^4$-PD I3-01v9b SApNP-Wild-type vs. SUDV GPΔmuc-WL$^2$P$^4$-PD I3-01v9b SApNP-Kif-treated | ns | 0.0604 |
| SUDV GPΔmuc-WL$^2$P$^4$-PD I3-01v9b SApNP-Wild-type vs. SUDV GPΔmuc-WL$^2$P$^4$-PD I3-01v9b SApNP-Kif/Endo H | ns | 0.9009 |
| SUDV GPΔmuc-WL$^2$P$^4$-PD I3-01v9b SApNP-Kif-treated vs. SUDV GPΔmuc-WL$^2$P$^4$-PD I3-01v9b SApNP-Kif/Endo H | ns | 0.1599 |

| One-way ANOVA with Tukey's multiple comparisons test (w8) | Statistics | Adjusted P Value |
|---|---|---|
| SUDV GPΔmuc-WL$^2$P$^4$-PD I3-01v9b SApNP-Wild-type vs. SUDV GPΔmuc-WL$^2$P$^4$-PD I3-01v9b SApNP-Kif-treated | ns | 0.1544 |
| SUDV GPΔmuc-WL$^2$P$^4$-PD I3-01v9b SApNP-Wild-type vs. SUDV GPΔmuc-WL$^2$P$^4$-PD I3-01v9b SApNP-Kif/Endo H | ns | 0.7712 |
| SUDV GPΔmuc-WL$^2$P$^4$-PD I3-01v9b SApNP-Kif-treated vs. SUDV GPΔmuc-WL$^2$P$^4$-PD I3-01v9b SApNP-Kif/Endo H | * | 0.0468 |

### Figure S11

#### i Sera of individual mice immunized with glycan-modified RAVV vaccines binding to RAVV GPΔmuc-P²CT(1TD0) trimer

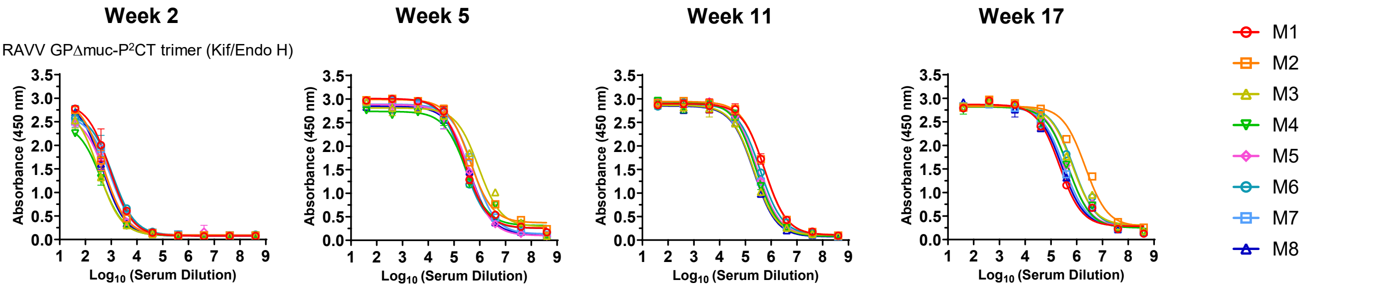


#### j Mouse serum ELISA $EC_{50}$ titers

**Week 2**

| Antigen | $EC_{50}$ titers (week 2) | | | | | | | | Geometric Mean |
|---|---|---|---|---|---|---|---|---|---|
| | M1 | M2 | M3 | M4 | M5 | M6 | M7 | M8 | |
| RAVV GPΔmuc-P²CT trimer-Wild-type | 718.9 | 721.9 | 432.3 | 1030.0 | 946.3 | 262.3 | 481.5 | 958.5 | 635.1 |
| RAVV GPΔmuc-P²CT trimer-Kif/Endo H | 1088.0 | 842.6 | 372.8 | 326.6 | 773.6 | 1455.0 | 1260.0 | 571.6 | 740.6 |

**Week 5**

| Antigen | $EC_{50}$ titers (week 5) | | | | | | | | Geometric Mean |
|---|---|---|---|---|---|---|---|---|---|
| | M1 | M2 | M3 | M4 | M5 | M6 | M7 | M8 | |
| RAVV GPΔmuc-P²CT trimer-Wild-type | 202735 | 1544554 | 746406 | 628189 | 447117 | 44771 | 446178 | 225949 | 362215.2 |
| RAVV GPΔmuc-P²CT trimer-Kif/Endo H | 208318 | 261241 | 222068 | 150370 | 267446 | 198233 | 450054 | 211636 | 234603.9 |

**Week 11 or 8**

| Antigen | $EC_{50}$ titers (week 11 or 8) | | | | | | | | Geometric Mean |
|---|---|---|---|---|---|---|---|---|---|
| | M1 | M2 | M3 | M4 | M5 | M6 | M7 | M8 | |
| RAVV GPΔmuc-P²CT trimer-Wild-type (w11) | 139135 | 218951 | 208199 | 104125 | 998580 | 786127 | 408247 | 853062 | 340466.6 |
| RAVV GPΔmuc-P²CT trimer-Kif/Endo H (w8) | 770620 | 683058 | 388118 | 292655 | 294767 | 621323 | 524297 | 273292 | 446120.6 |

**Week 17 or 11**

| Antigen | $EC_{50}$ titers (week 17 or 11) | | | | | | | | Geometric Mean |
|---|---|---|---|---|---|---|---|---|---|
| | M1 | M2 | M3 | M4 | M5 | M6 | M7 | M8 | |
| RAVV GPΔmuc-P²CT trimer-Wild-type (w17) | 140811 | 110900 | 164905 | 119990 | 336478 | 344895 | 127509 | 141835 | 168457 |
| RAVV GPΔmuc-P²CT trimer-Kif/Endo H (w11) | 125144 | 295308 | 212027 | 173036 | 313400 | 398840 | 295603 | 237268 | 242316 |

##### Statistical analysis

| Unpaired t test (w2) | Statistics | P Value |
|---|---|---|
| RAVV GPΔmuc-P²CT trimer-Wild-type vs. RAVV GPΔmuc-P²CT trimer-Kif/Endo H | ns | 0.4307 |

| Unpaired t test (w5) | Statistics | P Value |
|---|---|---|
| RAVV GPΔmuc-P²CT trimer-Wild-type vs. RAVV GPΔmuc-P²CT trimer-Kif/Endo H | ns | 0.1079 |

#### k Purified IgG of individual mice immunized with RAVV vaccines neutralizing a MARV Angola strain

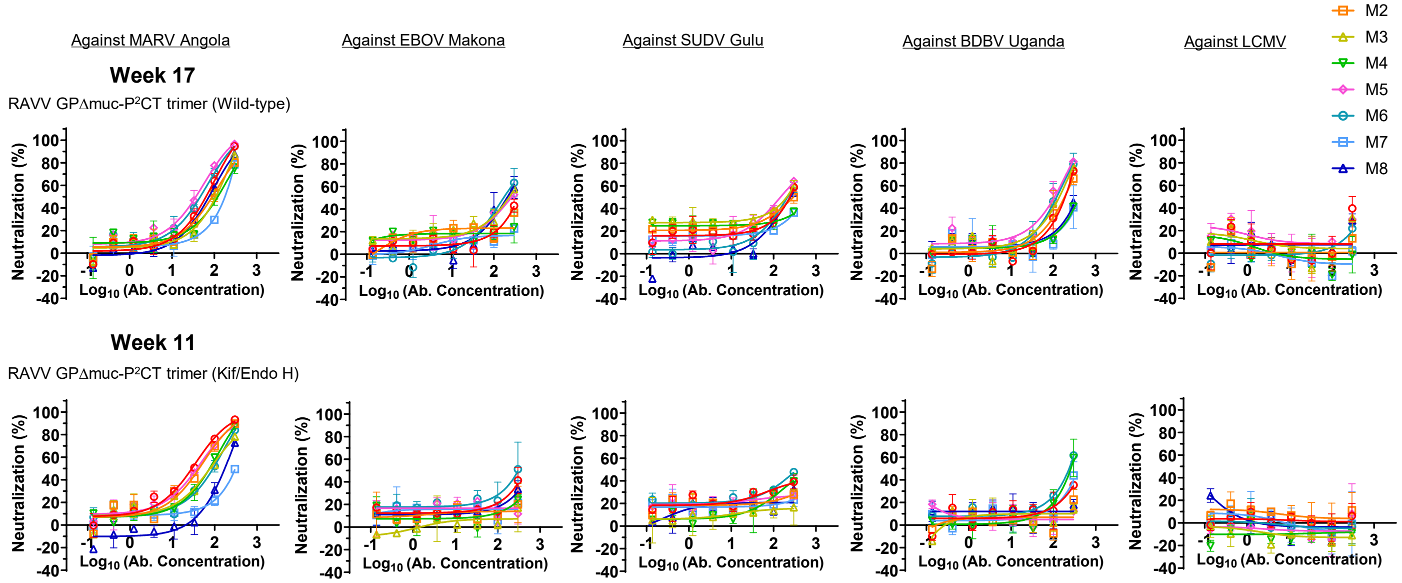

**Fig. S11. Immunogenicity of glycan-modified EBOV, SUDV, and RAVV GPΔmuc vaccines in mice.** (**a**) ELISA curves of mouse sera from glycan-modified EBOV GPΔmuc trimer and SApNP vaccine groups (n = 8 mice/group) binding to EBOV GPΔmuc-WL$^2$P$^4$(1TD0) trimer. (**b**) (Top) Summary of geometric mean $EC_{50}$ titers measured for glycan-modified EBOV GPΔmuc vaccine groups against EBOV GPΔmuc-WL$^2$P$^4$(1TD0) trimer. Color coding indicates $EC_{50}$ levels (green to red: low to high binding). (Bottom) Summary of statistical analysis performed for each timepoint. Note: $EC_{50}$ values at week 2 were derived by setting the minimum/maximum $OD_{450}$ values to 0.0/2.8. (**c**) Neutralization curves of mouse sera from glycan-modified EBOV GPΔmuc trimer and SApNP vaccine groups against EBOV Makona pseudovirus. (**d**) (Top) Summary of geometric means of $ID_{50}$ titers measured for EBOV GPΔmuc vaccine groups against EBOV Makona pseudovirus. Color coding: white (no neutralization), green to red (low to high neutralization). Note: $ID_{50}$ values were calculated using %neutralization constraints of 0.0 (min) and 100.0 (max). (Bottom) Summary of statistical analysis. (**e**) ELISA curves of mouse sera from glycan-modified SUDV GPΔmuc trimer and SApNP vaccine groups binding to SUDV GPΔmuc-WL$^2$P$^4$(1TD0) trimer. (**f**) (Top) Summary of geometric mean $EC_{50}$ titers measured for glycan-modified SUDV GPΔmuc vaccine groups against SUDV GPΔmuc-WL$^2$P$^4$(1TD0) trimer. (Bottom) Summary of statistical analysis. Note: $EC_{50}$ values at week 2 were derived by setting the minimum/maximum $OD_{450}$ values to 0.0/2.9. (**g**) Neutralization curves of mouse sera from glycan-modified SUDV GPΔmuc trimer and SApNP vaccine groups against SUDV Gulu pseudovirus. (**h**) (Top) Summary of geometric means of $ID_{50}$ titers measured for SUDV GPΔmuc vaccine groups against SUDV Gulu pseudovirus. (Bottom) Summary of statistical analysis. (**i**) ELISA curves of mouse sera from glycan modified RAVV GPΔmuc trimer vaccine groups binding to RAVV GPΔmuc-P$^2$CT(1TD0) trimer. (**j**) (Top) Summary of geometric mean $EC_{50}$ titers measured for glycan-modified RAVV GPΔmuc vaccine groups against RAVV GPΔmuc-P$^2$CT(1TD0) trimer. (Bottom) Summary of statistical analysis. Note: $EC_{50}$ values at week 2 were derived by setting the minimum/maximum $OD_{450}$ values to 0.0/2.9. (**k**) Neutralization curves of purified mouse IgG from glycan-modified RAVV GPΔmuc vaccine groups after four immunizations at week 17 or week 11 against MARV Angola or orthoebolavirus strains. Lymphocytic choriomeningitis virus (LCMV) was included as a negative control to confirm the cross-NAb responses measured using purified IgG. Error bars represent the difference between duplicate measurements at each concentration for each sample. $EC_{50}$ and $ID_{50}$ values were calculated using GraphPad Prism version 10.3.1. Data were analyzed using one-way ANOVA, followed by Tukey's multiple comparison post hoc test for each timepoint. Two-tailed unpaired t-tests were used to compare the geometric means between two independent groups. Statistical significance was defined as follows: ns (not significant), $^*p < 0.05$, $^{**}p < 0.01$, $^{***}p < 0.001$, and $^{****}p < 0.0001$.

### Figure S12

a

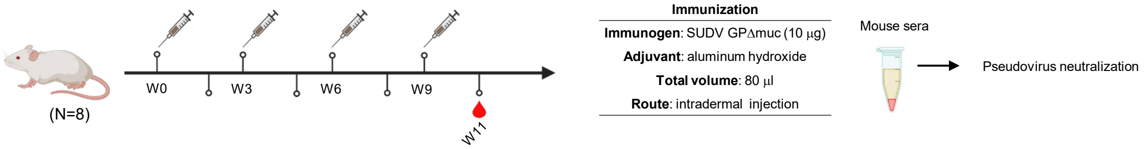


b **Week-11 sera of female and male mice immunized with wild-type SUDV vaccines neutralizing a SUDV Gulu strain**

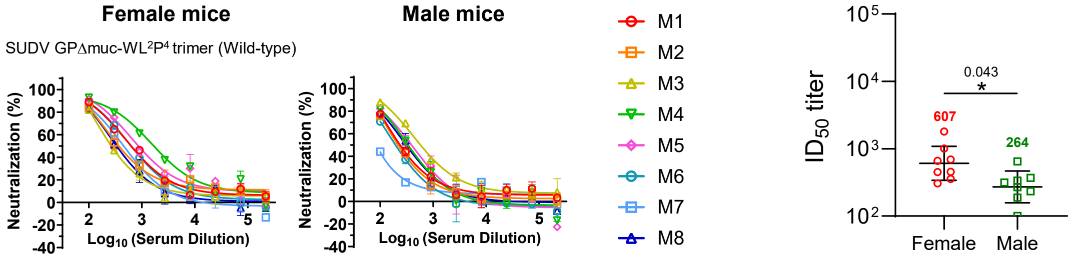


| Sex | ID$_{50}$ titers (week 11) | | | | | | | | Geometric Mean |
|---|---|---|---|---|---|---|---|---|---|
| | M1 | M2 | M3 | M4 | M5 | M6 | M7 | M8 | |
| Female | 690.2 | 454.8 | 307.7 | 1796 | 1007 | 660.1 | 449 | 355.5 | 606.9 |
| Male | 267.7 | 233.5 | 649.3 | 335.9 | 371.4 | 188.1 | <100 | 314.3 | 263.9 |

| Unpaired t test (w11) | Statistics | P Value |
|---|---|---|
| female vs. male | * | 0.043 |

c **Week-11 sera of female and male mice immunized with glycan-modified SUDV vaccines neutralizing a SUDV Gulu strain**

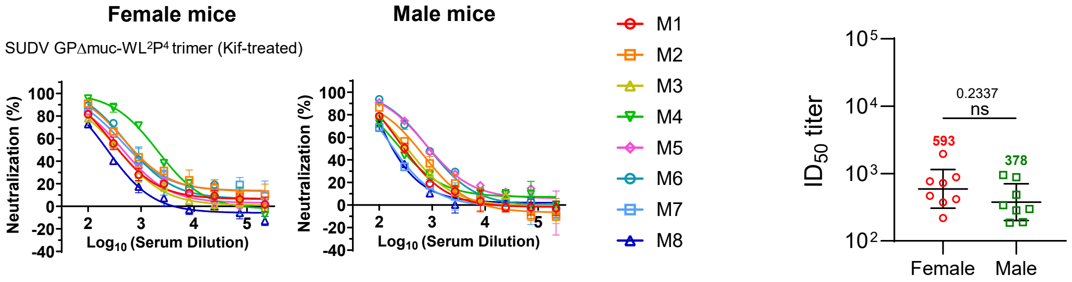


| Sex | ID$_{50}$ titers (week 11) | | | | | | | | Geometric Mean |
|---|---|---|---|---|---|---|---|---|---|
| | M1 | M2 | M3 | M4 | M5 | M6 | M7 | M8 | |
| Female | 417.3 | 885.7 | 374.1 | 1953 | 468 | 769.3 | 726 | 217.8 | 593.4 |
| Male | 297.6 | 485.6 | 340.5 | 283.6 | 954.3 | 880.1 | 185.9 | 190.2 | 377.7 |

| Unpaired t test (w11) | Statistics | P Value |
|---|---|---|
| female vs. male | ns | 0.2337 |

**Fig. S12. Influence of sex on immunogenicity of SUDV GPΔmuc-induced neutralizing antibody responses in mice.** (**a**) Schematic illustration of the mouse immunization regimen for SUDV GPΔmuc trimer vaccines (n = 8 mice per group). Each mouse received 80 µl of a vaccine antigen/aluminum hydroxide (AH) adjuvant mix containing 10 µg of SUDV antigen and 40 µl of AH. Mice were immunized via the intradermal route through four footpad injections at weeks 0, 3, 6, and 9, with 3-week intervals between doses. (**b**) Neutralization curves of female and male mouse sera from the wild-type SUDV GPΔmuc trimer group at week 11 (after four immunizations), tested against SUDV Gulu pseudovirus. (**c**) Neutralization curves of female and male mouse sera from the glycan-modified SUDV GPΔmuc trimer group under the same conditions. Comparison of NAb responses (by ID$_{50}$ titers) between female and male mice is shown, along with a summary of geometric mean ID$_{50}$ values for each group against SUDV Gulu pseudovirus. Error bars represent the difference between duplicate measurements at each concentration for each sample. ID$_{50}$ values were calculated using GraphPad Prism version 10.3.1. Two-tailed unpaired t-tests were used to compare the geometric means between two independent groups. Statistical significance was defined as follows: ns (not significant), *$p < 0.05$.

**Table S1. Data collection and refinement statistics for EBOV Mayinga GPΔmuc-WL²P⁴.**

| Data Collection | **EBOV Mayinga GPΔmuc-WL²P⁴** |
|---|---|
| Beamline | SSRL 12-1 |
| Wavelength (Å) | 0.97946 |
| Resolution (Å)[a] | 40.6 - 3.20 (3.29 - 3.20) |
| Space group | P321 |
| Unit cell (Å) | 114.47,114.47,133.38 |
| ( °) | 90 90 120 |
| Total reflections | 189,316 (12,886) |
| Unique reflections | 17,104 (815) |
| Multiplicity | 11.1 (9.6) |
| Completeness (%) | 99.8 (98.2) |
| Mean (I)/ $\sigma_I$ | 22.2 (1.2) |
| $R_{merge}$ (%) | 6.1 (>100) |
| $R_{meas}$[c] (%) | 12.3 (>100) |
| $R_{pim}$[d] (%) | 3.7 (69.3) |
| $CC_{1/2}$[e] (%) | 99.8 (30.3) |
| Refinement | |
| Refinement resolution (Å)[a] | 40.6 - 3.20 (3.29 - 3.20) |
| # reflections in refinement (work/free) | 17,082 (1,323) |
| $R_{work}$ (%) | 21.5 (36.1) |
| $R_{free}$ (%) | 22.2 (33.0) |
| # Protein atoms | 3,119 |
| # Carbohydrate atoms | 128 |
| # Waters | 0 |
| # Protein residues | 398 |
| Bond r.m.s. deviation (Å) | 0.013 |
| Angle r.m.s. deviation (º) | 1.60 |
| Wilson B (Å$^2$) | 114 |
| Average B protein (Å$^2$) | 119 |
| carbohydrate (Å$^2$) | 156 |
| Ramachandran favored, allowed, outliers (%) | 96.1, 3.9, 0.0, 0.0 |
| Clashscore[f] | 8.2 |
| PDB ID | 9N8E |

[a]Numbers in parentheses are for highest resolution shell

[b]$R_{merge} = \Sigma_{hkl} \Sigma_{i=1,n}|I_i(hkl) - \langle I(hkl)\rangle| / \Sigma_{hkl} \Sigma_{i=1,n} I_i(hkl)$

[c]$R_{meas} = \Sigma_{hkl} \sqrt{(n/n-1)} \Sigma_{i=1,n}|I_i(hkl) - \langle I(hkl)\rangle| / \Sigma_{hkl} \Sigma_{i=1,n} I_i(hkl)$

[d]$R_{pim} = \Sigma_{hkl} \sqrt{(1/n-1)} \Sigma_{i=1,n}|I_i(hkl) - \langle I(hkl)\rangle| / \Sigma_{hkl} \Sigma_{i=1,n} I_i(hkl)$

[e]$CC_{1/2}$ = Pearson correlation coefficient between two random half datasets

[f] Number of unfavorable all-atom steric overlaps ≥ 0.4 Å per 1000 atoms

**Table S2. Cryo-EM data collection information, model building, and refinement statistics.**

| Data collection information | **SUDV Gulu GPΔmuc-WL$^2$P$^4$/CA45 Fab** |
|---|---|
| Microscope | Titan Krios G4 |
| Voltage (keV) | 200 |
| Detector | Falcon 4 camera with a Selectris-X energy filter |
| Recording mode | Counting |
| Magnification | 130,000 x |
| Movie micrograph pixel size | 0.89 |
| Total dose ($e^-$/Å$^2$) | 50 |
| Under focus range (µm) | -1.0 to -2.0 um |
| Number of movie micrographs | 1,824 |
| Model building and refinement statistics | |
| Map Resolution (Å) | 3.13 |
| Residues | |
| Amino-acids | 1473 |
| Carbohydrates | 9 |
| RMSD Bonds (Å) | 0.003 |
| RMSD Angles (º) | 0.746 |
| Ramachandran | |
| Outliers (%) | 0.00 |
| Allowed (%) | 7.2 |
| Favored (%) | 92.8 |
| Rotamer outliers | 1.67 |
| Clash score | 8.98 |
| MolProbity score | 2.10 |
| PDB ID | 9N8F |
| EMDB ID | EMD-49127 |
